# Statistics of eigenvalue dispersion indices: quantifying the magnitude of phenotypic integration

**DOI:** 10.1101/2021.06.19.449119

**Authors:** Junya Watanabe

## Abstract

Quantification of the magnitude of trait covariation plays a pivotal role in the study of phenotypic evolution, for which statistics based on dispersion of eigenvalues of a covariance or correlation matrix—eigenvalue dispersion indices—are commonly used. This study remedies major issues over the use of these statistics, namely, a lack of clear understandings on their statistical justifications and sampling properties. The relative eigenvalue variance of a covariance matrix is known in the statistical literature a test statistic for sphericity, thus is an appropriate measure of eccentricity of variation. The same of a correlation matrix is equal to the average squared correlation, which has a straightforward interpretation as a measure of integration. Expressions for the mean and variance of these statistics are analytically derived under multivariate normality, clarifying the effects of sample size *N*, number of variables *p*, and parameters on sampling bias and error. Simulations confirmed that approximations involved are reasonably accurate with a moderate sample size (*N* ≥ 16–64). Importantly, sampling properties of these indices are not adversely affected by a high *p*:*N* ratio, promising their utility in high-dimensional phenotypic analyses. They can furthermore be applied to shape variables and phylogenetically structured data with appropriate modifications.

## Introduction

Analysis of trait covariation plays a central role in investigations into evolution of quantitative traits. The well-known quantitative genetic theory of correlated traits predicts that evolutionary response in a population under selection is dictated by the additive genetic covariance matrix **G** as well as the selection gradient (Lande, 1979; Lande & Arnold, 1983). Short-term evolutionary changes of a population are expected to be concentrated in major axes of the **G** matrix (Schluter, 1996). Arguably, the structure of the **G** matrix can be approximated by that of the phenotypic covariance matrix for certain types of traits (Cheverud, 1988, 1996; Roff, 1995; Dochtermann, 2011; Sodini et al., 2018), so the latter could be analyzed when accurate estimation of the **G** matrix is not feasible. These theories and conjectures spurred extensive theoretical and empirical explorations on character covariation as an evolutionary constraint (e.g., Steppan et al., 2002; Chenoweth et al., 2010; Pitchers et al., 2014; Hansen et al., 2019 and references therein). Partly fueled by these developments, the study of phenotypic integration has developed as an active field of research, where various aspects of character covariation are investigated with diverse motivations and scopes (e.g., Olson & Miller, 1958; Cheverud, 1982; Goswami, 2006; Hallgrímsson et al., 2009; Armbruster et al., 2014; Felice et al., 2018). In the latter context, many different levels of organismal variation can be subjects of research, such as static, ontogenetic, and evolutionary levels (Klingenberg, 2014). For example, relationships between within-population integration and evolutionary rate and/or trajectories have attained much attention as potential links between micro- and macroevolutionary phenomena (e.g., Klingenberg et al., 2012; Renaud & Auffray, 2013; Bolstad et al., 2014; Goswami et al., 2015; Haber, 2015, 2016).

An obvious target of investigation in these contexts is quantitative analysis of the magnitude of constraint or integration entailed in covariance structures. In particular, this paper concerns the methodology for quantifying the overall magnitude of covariation within a set of traits. Quantification of relative (in)dependence between multiple sets of traits—the modularity–integration spectrum—is another major way of studying integration which has separate methodological frameworks (e.g., Goswami & Polly, 2010; Adams, 2016; Goswami & Finarelli, 2016; Adams & Collyer, 2019a). Demonstrating the presence of integration with a statistically justified measure can be the scope of an empirical analysis, sometimes as a part of testing combined hypotheses (e.g., Brommer, 2014; Watanabe, 2018). A univariate summary statistic for magnitude of integration can conveniently be used in comparative analyses across developmental stages, populations, or phylogeny (e.g., Marroig et al., 2009; Porto et al., 2009; Haber, 2016). A plethora of statistics have been proposed for such purposes from various standpoints (e.g., Van Valen, 1974, 2005; Cheverud et al., 1983, 1989; Wagner, 1984; Cane, 1993; Hansen & Houle, 2008; Agrawal & Stinchcombe, 2009; Kirkpatrick, 2009; Pavlicev et al., 2009; Armbruster et al., 2009, 2014; Haber, 2011). One of the most popular classes of such statistics is based on the dispersion of eigenvalues of a covariance or correlation matrix. These statistics have the forms

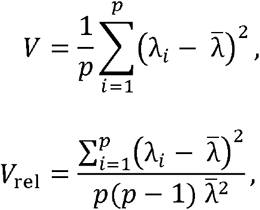

where *p* is the number of variables (traits), λ_*i*_ is the *i*th eigenvalue of the covariance or correlation matrix under analysis, and 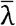 is the average of the eigenvalues. Here, *V* is the most naïve form of eigenvalue dispersion, and *V*_rel_ is a scaled version which ranges between 0 and 1. Formal definitions are given below with distinction between population and sample quantities. Some authors use square root or a constant multiple of these forms, but such variants essentially bear identical information when calculated from the same matrix. Alternative terms for this class of statistics include the tightness (for *V*_rel_; Van Valen, 1974; later used for 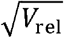 by Van Valen, 2005), integration coefficient of variation (for 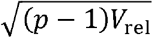; Shirai & Marroig, 2010), and phenotypic integration index (for *V*; Torices & Muñoz-Pajares, 2015). In this paper, *V* and *V*_rel_ are called the eigenvalue variance and relative eigenvalue variance, respectively, to take a balance between brevity and descriptiveness. These quantities are not to be confused with the sampling variance associated with eigenvalues in a sample (see below).

Since eigenvalues of a covariance or correlation matrix correspond to the variance along the corresponding eigenvectors (principal components), these statistics are supposed to represent eccentricity of variation across directions in a trait space (Fig. 1; Wagner, 1984). Cheverud et al. (1983) and Wagner (1984) were the first to propose using *V* of a correlation matrix for quantifying magnitude of integration. Pavlicev et al. (2009) devised *V*_rel_ for a correlation matrix, and explored its relationships to correlation structures in certain biologically relevant conditions. Haber (2011) pointed out similarity between these indices and Van Valen’s (1974) tightness index for a covariance matrix, and proposed that these indices can be applied to either covariance or correlation matrices with slightly different interpretations. Eigenvalue dispersion indices are frequently used in empirical analyses of phenotypic integration at various levels of organismal variation, from phenotypic covariance at the population level to evolutionary covariance at the interspecific level (e.g., Ordano et al., 2008; Torices & Mendez, 2014; Haber, 2016; Haber & Dworkin, 2017; Watanabe, 2018; Arlegi et al., 2020). However, use of these indices has been criticized for a lack of clear statistical justifications; it has not been known—or not widely appreciated by biologists— exactly what they are designed to measure, beyond the intuitive allusion to eccentricity mentioned above (Hansen & Houle, 2008; Blows & McGuigan, 2015; Hansen et al., 2019).

**Figure 1.**
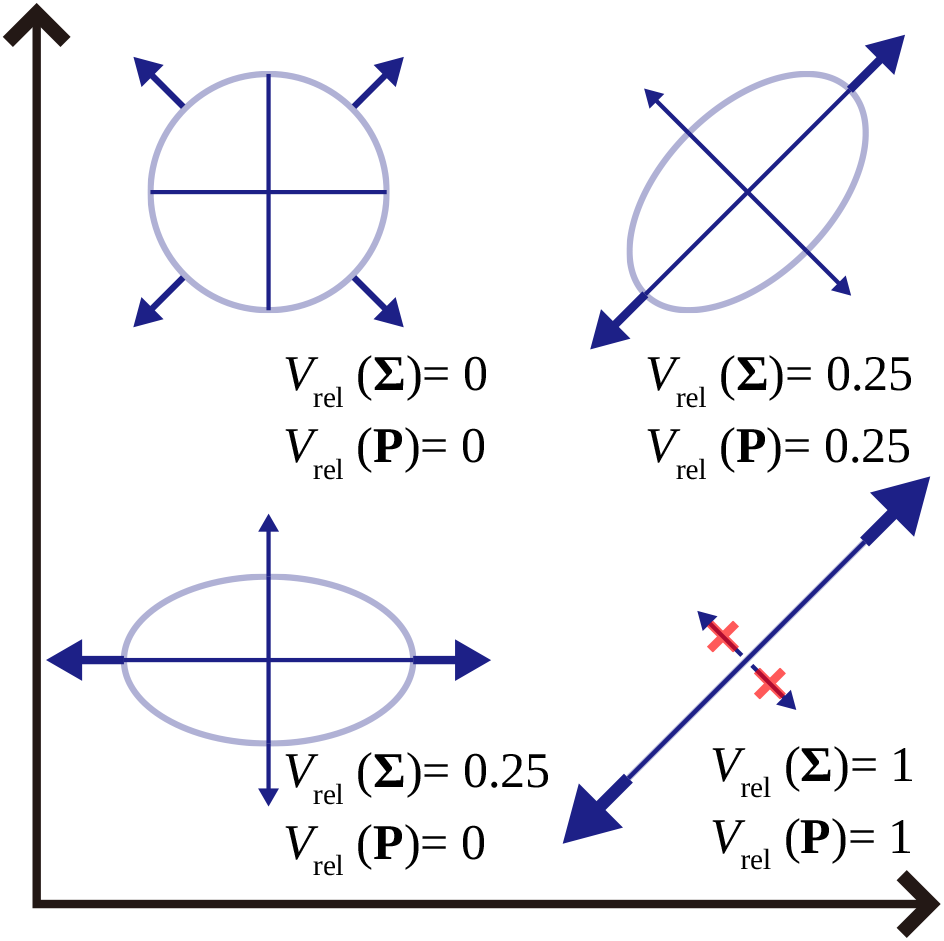
Schematic illustration of eigenvalue dispersion indices in bivariate cases. Ellipses representing equiprobability contours are shown on a Cartesian space of two hypothetical variables for four conditions, as well as the relative eigenvalue variance of the corresponding covariance and correlation matrices (*V*_rel_(**Σ**) and *V*_rel_(**P**), respectively). The scale is arbitrary but identical for the two axes. The axes of each ellipse are proportional to square roots of the two eigenvalues of the respective covariance matrix. *V*_rel_(**Σ**) represents eccentricity of variation and is sensitive to differing scale changes between axes but not to rotation (change of eigenvectors), whereas *V*_rel_(**P**) represents magnitude of correlation and is insensitive to scale changes. Arrows schematically represent variation along major axes (whose directions are arbitrary when *V*_rel_(**Σ**) = 0).

Another fundamental issue over the eigenvalue dispersion indices is a virtual lack of systematic understanding of their sampling properties. In empirical analyses, eigenvalue dispersion indices are calculated from sample covariance or correlation matrices, but interests will be in making inferences for the underlying populations. For example, interest may be in detecting the presence of constraint in a population, i.e., testing the null hypothesis of sphericity. As detailed below, however, sample eigenvalues are always estimated with error, so that *V* and *V*_rel_ calculated from them take a positive value even if the corresponding population values are 0. In other words, empirical eigenvalue dispersion indices are biased estimators of the corresponding population values under the null hypothesis. For statistically justified inferences, it is crucial to capture essential aspects of their sampling distributions, e.g., expectation and variance.

The presence of sampling bias in eigenvalue dispersion indices has been well known in the literature (Wagner, 1984; Cheverud et al., 1989; Grabowski & Porto, 2017; see also Marroig et al., 2012; Björklund, 2019). Simulation-based studies have been undertaken to sketch sampling distributions of eigenvalue dispersion indices and related statistics (Haber, 2011; Grabowski & Porto, 2017; Machado et al., 2019; Jung et al. 2020). However, these approaches hardly give any systematic insight beyond the specific conditions considered. Analytic results should preferably be sought to comprehend the sampling bias and error. In this regard, it is notable that Wagner (1984) derived the first two moments of eigenvalues of sample covariance and correlation matrices under the null conditions. The variance of sample eigenvalues obtained from these moments has later been used as an estimate of sampling bias in these conditions (e.g., Cheverud et al., 1989). Strictly speaking, however, the variance of a sample eigenvalue is fundamentally different from the expectation of the eigenvalue variance *V*. These quantities are identical for correlation matrices under the null hypothesis, but this is not the case for covariance matrices where the covariances between sample eigenvalues cannot be ignored (see below). Furthermore, Wagner’s (1984) results have a few restrictive conditions: variables to have the means of 0, or equivalently, to be centered at the population mean rather than the sample mean as is done in most empirical analyses (although this was probably appropriate in the strict context of his theoretical model); and the sample size *N* to be equal to or larger than the number of variables *p*, so their applicability to *p* > *N* conditions has not been demonstrated.

In addition to the naïve null condition of no integration, moments under arbitrary conditions are also desired. Such would be useful in testing hypotheses about the magnitude (rather than the mere presence/absence) of integration (Harder, 2009; Fornoni et al., 2009) and comparing the magnitudes between different samples (Cheverud et al., 1989). Also, the assumption of no covariation is intrinsically inappropriate as a null hypothesis for shape variables where raw data are transformed in such a way that individual “variables” are necessarily dependent on one another (e.g., Mitteroecker et al., 2012). For this type of data, a covariance matrix with an appropriate structure needs to be specified as the null model representing the intrinsic covariation.

This paper addresses the issues over the eigenvalue dispersion indices mentioned above. It first gives a theoretical overview of these statistics to clarify their statistical justifications, particularly in connection to the sphericity test in multivariate analysis. Then exact and approximate expressions are analytically derived for the expectation and variance of *V* and *V*_rel_ of sample covariance and correlation matrices under the null and arbitrary conditions, assuming the multivariate normality of variables. These expressions are derived without any assumption on *p* or *N*, except for the variance of *V* and *V*_rel_ of a correlation matrix under arbitrary conditions, which is based on large-sample asymptotic theories. Simulations were subsequently conducted to obtain systematic insights into sampling properties and to evaluate the accuracy of the approximate expressions. Extensions into shape variables and phylogenetically structured data are briefly discussed.

## Theory

### Preliminaries

For the purpose here, the distinction between population and sample quantities is essential. Corresponding Greek and Latin letters are used as symbols for the former and latter, respectively. Let **Σ** be the *p* × *p* population covariance matrix, whose (*i, j*)-th element σ_*ij*_ is the population variance (*i* = *j*) or covariance (*i* ≠ *j*). It is a symmetric, nonnegative definite matrix with the eigendecomposition

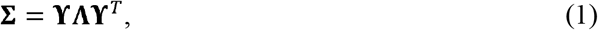

where the superscript ^*T*^ denotes matrix transposition, **ϒ** is an orthogonal matrix of eigenvectors **(ϒϒ**^*T*^ = **ϒ**^*T*^**ϒ** = **I**_*p*_ where **I**_*p*_ is the *p* × *p* identity matrix), and **Λ** is a diagonal matrix whose diagonal elements are the eigenvalues λ_1_, λ_2_, …, λ_*p*_ of **Σ** (population eigenvalues). For convenience, the eigenvalues are arranged in the non-increasing order: λ_1_ ≥ λ_2_ ≥ ··· ≥ λ_*p*_ ≥ 0. Let **μ** be the *p* × 1 population mean vector.

Let **X** be an *N* × *p* observation matrix consisting of *p*-variate observations, which are individually denoted as **x**_*i*_ (*p* × 1 vector; transposed in the rows of **X**). (No strict notational distinction is made between a random variable and its realization.) At this point, *N* observations are assumed to be identically and independently distributed (i.i.d.). The sample covariance matrix **S** and cross-product matrix **A** are defined as

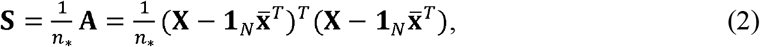

where **1**_*N*_ is a *N* × 1 column vector of 1’s, 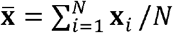 is the sample mean vector, and *n*_*_ denotes an appropriate devisor; e.g., *n*_*_ = *N* − 1 for the ordinary unbiased estimator, and *n*_*_ = *N* for the maximum likelihood estimator under the normal distribution. The (*i, j*)-th element of **S**, denoted *S_ij_*, is the sample variance or covariance. The eigendecomposition of **S** is constructed in the same way as above:

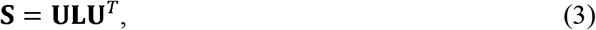

where **U** is an orthogonal matrix of sample eigenvectors and **L** is a diagonal matrix whose elements are the sample eigenvalues *l*_1_, *l*_2_, …, *l_p_*.

In what follows, the following identity entailed by the orthogonality of **U** is frequently utilized:

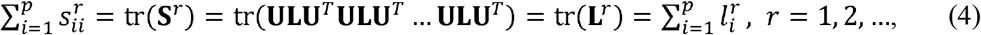

where tr(·) denotes the matrix trace operator, i.e., summation of the diagonal elements; the parentheses are omitted for visual clarity when little ambiguity exists. The sum of variances tr **S** = tr **L** is called total variance. Note that equation 4 holds even when *N* − 1 < *p*, in which case *l_i_* = 0 for some *i*. In other words, when *N* − 1 < *p*, the sample total variance is in a way concentrated in a subspace with fewer dimensions than the full space.

The population and sample correlation matrices **P** and **R**, whose (*i, j*)-th elements are the population and sample correlation coefficients ρ_*ij*_ and *r_ij_*, respectively, are obtained by standardizing **Σ** and **S**:

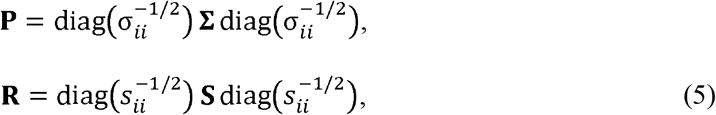

where diag(·) stands for the *p* × *p* diagonal matrix with the designated *i*th elements. Their eigendecomposition is defined as for covariance matrices, and the eigenvalues are denoted with the same symbols. For any *i*, ρ_*ii*_ = *r_ii_* = 1, and hence, for correlation matrices

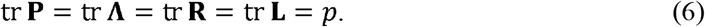

In what follows, the notations E(·), Var(·), and Cov(·,·) are used for the expectation (mean), variance, and covariance of random variables, respectively.

### Eigenvalue dispersion

The eigenvalue variance *V* is defined as:

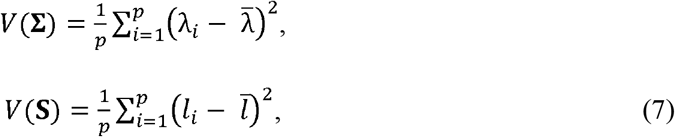

where 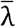 and 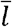 are the averages of the population and sample eigenvalues, respectively 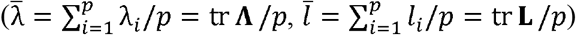. Note that *V*(**Σ**) is a quantity pertaining to the population, whereas *V*(**S**) is a sample statistic. The definition here follows the convention in the literature that *p*, rather than *p* − 1, is used as the divisor (e.g., Cheverud et al., 1983; Pavlicev et al., 2009; Haber, 2011). The latter might be more suitable for *V*(**S**) because the sum of squares is taken around the average sample eigenvalue which is a random variable. After all, however, the choice of *p* − 1 is not so useful because *V*(**S**) cannot be an unbiased estimator of *V*(**Σ**) even with that choice (below).

Note that the average and sum of squares are taken across all *p* eigenvalues, even if some eigenvalues are zero due to the condition *N* − 1 < *p*. This is reasonable given that sums of moments across all *p* sample eigenvalues are comparable in magnitude to those of population eigenvalues (see below). One could alternatively use eigenvalue standard deviation 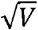 (Pavlicev et al., 2009; Haber, 2011), but this study concentrates on *V* rather than 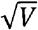, because the former is much more tractable for the purposes of characterizing distributions.

It is obvious that *V*(**Σ**) takes a single minimum of 0 at 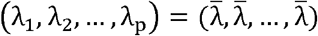. On the other hand, for a fixed 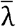, it takes a single maximum of 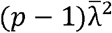 at 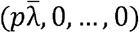 (e.g., Van Valen, 1974; Machado et al., 2019). Hence, not only is *V*(**Σ**) scale-variant, but also its range depends on *p* − 1. Therefore, it is often useful to standardize *V* by division with this maximum to obtain the relative eigenvalue variance *V*_rel_:

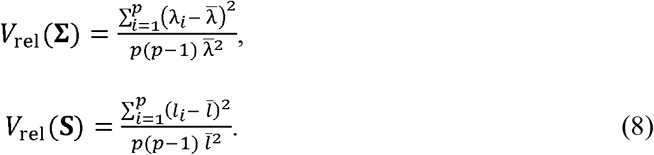

Because of the standardization, *V*_rel_ ranges between 0 and 1. This is a heuristic introduction of *V*_rel_ from *V*, but it will be seen below that *V*_rel_(**S**) has a clearer theoretical justification.

These indices are similarly defined for correlation matrices. By noting 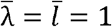 (eq. 6), these are

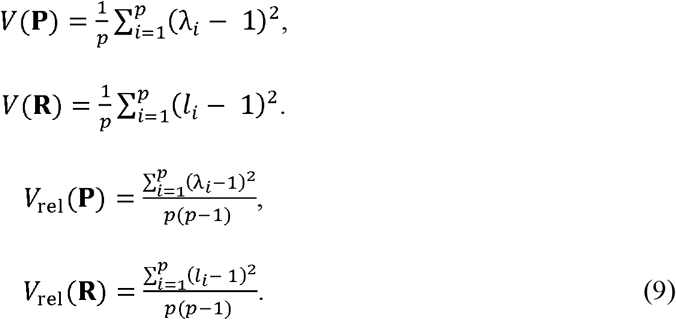

In the following discussions, we will concentrate on *V*_rel_ for correlation matrices, because *V*(**R**) and *V*_rel_(**R**) are proportional to each other by the factor *p* − 1, and hence their distributions are identical up to this scaling. This is in contrast to those of covariance matrices, where 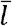 in the denominator in *V*_rel_(**S**) is a random variable and affects sampling properties.

Importantly, a single value of *V*_rel_ in general corresponds to multiple combinations of eigenvalues even if the average eigenvalue is fixed, except when *p* = 2 or under the extreme conditions *V*_rel_ = 0 and *V*_rel_ = 1 (Fig. 1). As such, it is not always straightforward to discern how intermediate values of *V*_rel_ are translated into actual covariance structures when *p* > 2. Nevertheless, it is possible to show that *V*_rel_ > 0.5 cannot happen when multiple leading eigenvalues are of the same magnitude (Appendix A); in other words, such a large value indicates dominance of the first principal component.

As would be obvious from the definition, *V* and *V*_rel_ of covariance matrices only describe the (relative) magnitudes of eigenvalues—proportions of the axes of variation—and do not reflect any information of eigenvectors—directions of the axes. A large eigenvalue of a covariance matrix can represent, e.g., strong covariation between equally varying traits or large variation of a single trait uncorrelated with others; either of these cases describes eccentricity of variation in the multivariate space. By contrast, a large eigenvalue of a correlation matrix can only happen in the presence of correlation. Therefore, a large eigenvalue dispersion in a correlation matrix constrains conformation of eigenvectors to a certain extent. The correlations can nevertheless be realized in various ways depending on eigenvectors, whose conformation does influence the sampling distribution of *V*_rel_(**R**) (see below).

For covariance matrices, *V*_rel_(**S**) has a natural relation to the test of sphericity, i.e., test of the null hypothesis that **Σ** = σ^2^**I**_*p*_ for an arbitrary positive constant σ^2^. Simple transformations from equation 8 lead to

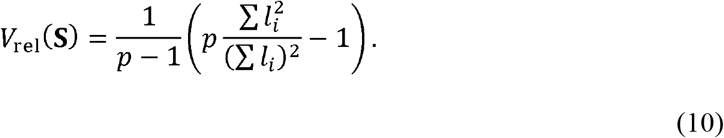

By noting 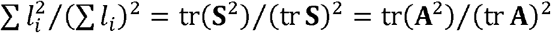 (see eqs. 2 and 4), *V*_rel_(**S**) in the form of equation 10 is exactly John’s (1972) *T* statistic for the test of sphericity (see also Ledoit & Wolf, 2002). Beyond the intuition that it measures eccentricity of variation along principal components, this statistic (and its linear functions) can be justified as the most powerful test statistic in the proximity of the null hypothesis under multivariate normality, among the class of such statistics that are invariant against translation by a constant vector, uniform scaling, and orthogonal rotation (John, 1971, 1972; Sugiura, 1972; Nagao, 1973). On the other hand, *V*(**S**) does not seem to have as much theoretical justification, but rather has a practical advantage in the tractability of its moments and ease of correcting sampling bias (see below).

For a correlation matrix, *V*_rel_ is a measure of association between variables. Following similar transformations, it is straightforward to see

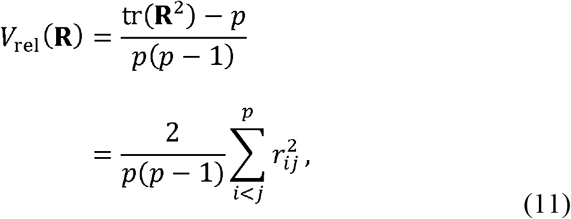

because 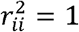 for all *i*. This relationship has been known in the statistical literature (e.g., Gleason & Staelin, 1975; Durand & Le Roux, 2017), and empirically confirmed by Haber (2011). This statistic is used as a measure of overall association between variables (e.g., Schott, 2005; Durand & Le Roux, 2017), with the corresponding null hypothesis being **P** = **I**_*p*_.

### Sampling properties of eigenvalues

The distribution of eigenvalues of **S**, or equivalently those of **A** (which are *n*_*_ times those of **S**), has been extensively investigated in the literature of multivariate analysis (see, e.g., Jolliffe, 2002; Anderson, 2003). Unfortunately, however, most of such results are of limited value for the present purposes. On the one hand, forms of the exact joint distribution of the eigenvalues of **A** are known under certain assumptions on the population eigenvalues (e.g., Muirhead, 1982: pp. 107, 388), but they do not allow for much intuitive interpretation (let alone direct evaluation of moments), apart from the following points: 1) sample eigenvalues are *not* stochastically independent from one another; and 2) the distribution of sample eigenvalues are only dependent on the population eigenvalues, but not on the population eigenvectors. On the other hand, a substantial body of results is available for large-sample asymptotic distributions of sample eigenvalues (assuming *n* → ∞, *p* being constant; e.g., Anderson, 1963, 2003), but their accuracy under finite *n* conditions is questionable. For example, a well-known asymptotic result states that 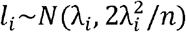 and Cov(*l_i_*, *l_j_*) ≈ 0 for *i* ≠ *j*, assuming λ_*i*_ ≠ λ_*j*_ and *n*_*_ = *n* (Girshick, 1939; Anderson, 1963; Srivastava & Khatri, 1979). However, these expressions ignore terms of order higher than *O*(*n*^−1^)—that is, all terms with >1 powers of *n* in the denominator—whose magnitude can be substantial for finite *n*. Indeed, with further evaluation of higher-order terms, it becomes evident that sample eigenvalues are biased estimators of the population equivalents, where large eigenvalues are prone to overestimation and small ones are prone to underestimation, and that Cov(*l_i_*, *l_j_*) = 2λ_*i*_λ_*j*_/[(λ_*i*_ − λ_*j*_)*n*]^2^ + *O*(*n*^−3^) for *i* ≠ *j*, again assuming λ_*i*_ ≠ λ_*j*_ (Lawley, 1956; Srivastava & Khatri, 1979). In general, the covariance between sample eigenvalues is nonzero.

Sampling properties of eigenvalues have also been investigated under high-dimensional asymptotic conditions (assuming *n* → ∞, *p* → ∞, and *p/n* typically held finite and constant; e.g., Johnstone, 2007; Johnstone & Paul, 2018). Attempts have been made to apply some of these results to **G** matrices (Blows & McGuigan, 2015; Sztepanacz & Blows, 2017), although these remain highly heuristic in nature and rely more heavily on simulations than analytic results. The theories primarily concern limiting distributions of largest sample eigenvalues or the entire bulk of sample eigenvalues, and do not seem directly applicable to the present analysis, which requires accurate expressions of certain higher-order moments of sum of eigenvalues (see below).

Much less is known about eigenvalues of a sample correlation matrix **R** (Jolliffe, 2002), whose distribution seems intractable except under certain special conditions (Anderson, 1963). Large-sample asymptotic results indicate that the limiting distribution *n* → ∞ of an eigenvalue of **R** is centered around the corresponding population eigenvalue but its variance depends on population eigenvectors (Anderson, 1963; Konishi, 1979), unlike that of a covariance matrix where the distribution does not depend on population eigenvectors (above).

It is often of practical interest to detect the presence of eccentricity or integration, i.e., to test the null hypothesis of sphericity **Σ** = σ^2^**I**_*p*_ or no correlation **P** = **I**_*p*_. These hypotheses are equivalent to *V*(**Σ**) = *V*_rel_(**Σ**) = 0 and *V*_rel_(**P**) = 0, respectively. Even under these conditions, nonzero sampling variance in sample eigenvalues renders *V*(**S**) > 0, *V*_rel_(**S**) > 0, and *V*_rel_(**R**) > 0 with probability 1, because these statistics are calculated from sum of squares. The primary aim here is to derive explicit expressions for this sampling bias (expectation), as well as sampling variance.

It should be remembered that the expectation of the eigenvalue variance E[*V*(**S**)] is fundamentally different from the variance of eigenvalues Var(*l_i_*). This point will be clarified by the following transformation:

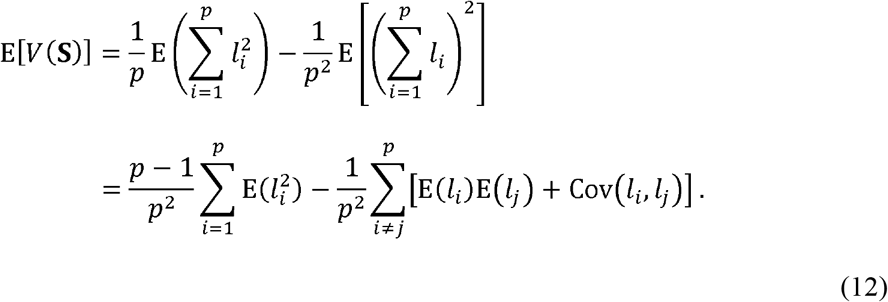

Under the null hypothesis, the moments are equal across all *i*, and the above simplifies into

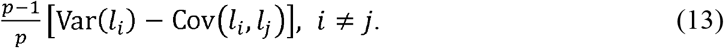

If Cov(*l_i_, l_j_*) were zero, the expectation would coincide with (*p* − 1)Var(*l_i_*)/*p*, which can be evaluated from, e.g., Wagner’s (1984) results. As already mentioned, however, this covariance is nonzero in general and hence cannot be ignored for covariance matrices. This is unlike the case for correlation matrices, where E [*V*(**R**)] = Var(*l_i_*) holds under the null hypothesis, because 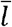 is a constant and equals E(*l_i_*) = 1.

In the following discussions on moments of eigenvalue dispersion indices, observations are assumed to be i.i.d. multivariate normal variables. If the *N* × *p* matrix **X** consists of *N* i.i.d. *p*-variate normal variables **x**_*i*_~*N_p_* (**μ, Σ**), then the distribution of the sample-mean-centered cross-product matrix **A** (eq. 2) is said to be the (central) Wishart distribution *W_p_*(**Σ**, *n*), where *n* = *N* − 1 is the degree of freedom. It is well known that this is identical to the distribution of **Z**^*T*^**Z**, where the *n* × *p* matrix **Z** consists of *n* i.i.d. *p*-variate normal variables **z**_*i*_~*N_p_*(**0**_*p*_, **Σ**) with **0**_*p*_ being the *p* × 1 column vector of 0’s (e.g., Anderson, 2003). Therefore, for analyzing statistics associated with sample covariance or correlation matrices, we can conveniently consider

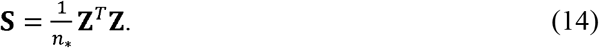

without loss of generality, by bearing in mind the distinction between the degree of freedom *n* and sample size *N*. From elementary moments of the normal distribution, we have

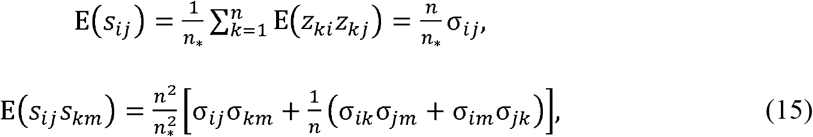

where *Z_ij_, S_ij_*, σ_*ij*_ and the like are the (*i, j*)-th elements of **Z**, **S**, and **Σ**, respectively.

### Moments under null hypotheses

#### Covariance matrix

Before proceeding to arbitrary conditions, let us consider the null hypothesis of sphericity: **Σ** = σ^2^**I**_*p*_. For the expectation of *V*(**S**), we need Var(*l_i_*) and Cov(*l_i_, l_j_*) or equivalently 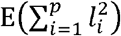 and 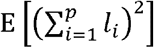 (see eqs. 12 and 13); we will proceed with the latter here. By use of equations 4 and 15, we have

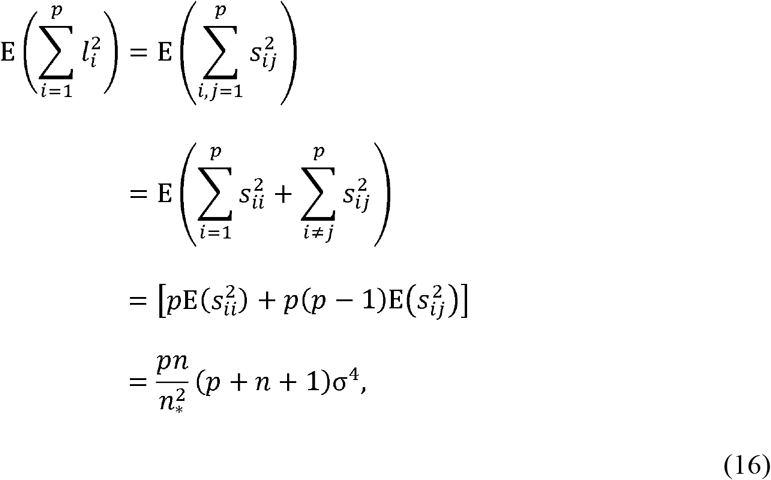

and similarly

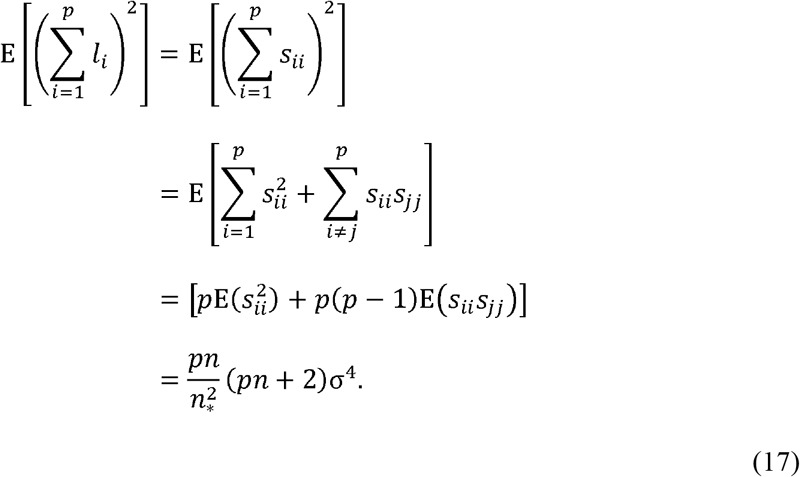

Then, inserting these results into equation 12,

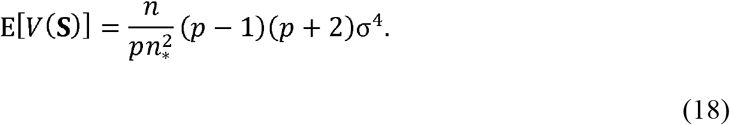

Alternatively, it could be seen that 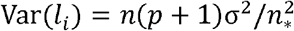 and 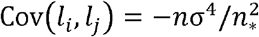 for *i* ≠ *j* (see also Girshick, 1939), with which equation 13 yields the identical result.

The variance of *V*(**S**) is, by equation 12,

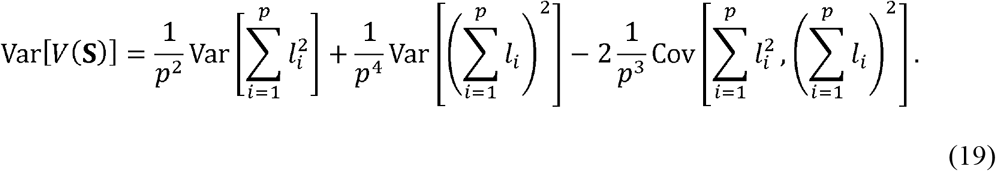

The relevant moments can most conveniently be found as a special case of general expressions under arbitrary **Σ** (see below and Appendix B), although direct derivation using normal moments is possible:

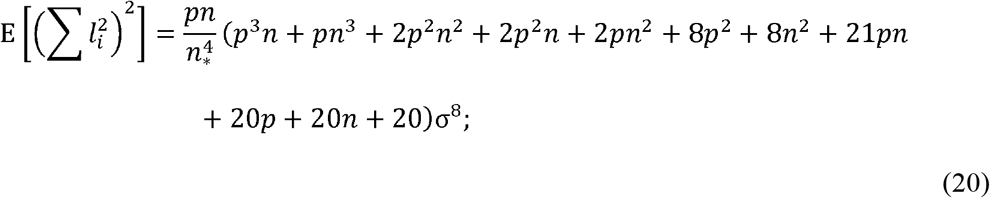

Inserting these into equation 19 yields

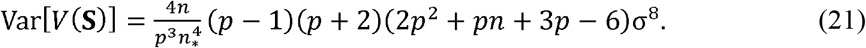

Next, consider the first two moments of *V*_rel_(**S**) under the null hypothesis (which have previously been derived by John [1972]). Recalling the form of equation 10,

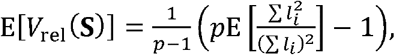

and

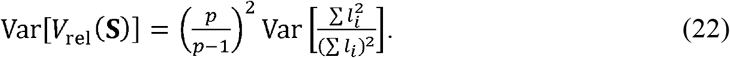

In general, moments of the ratio 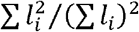 do not coincide with the ratios of the moments of the numerator and denominator. Specifically under the null hypothesis, however,

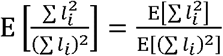

and

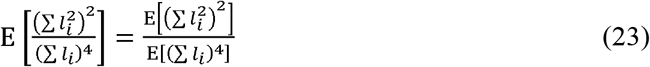

hold because of the stochastic independence between 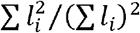 and ∑ *l_i_* in this special condition (which follows from inspection of the density; John, 1972). Therefore, by use of equations 16, 17, and 20,

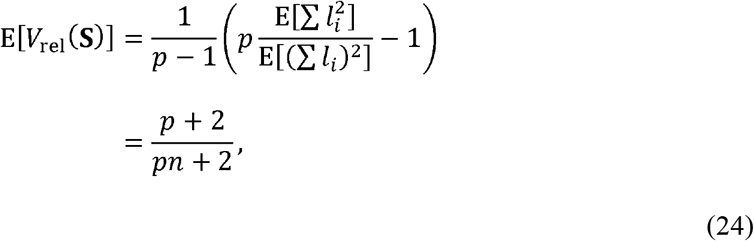

and

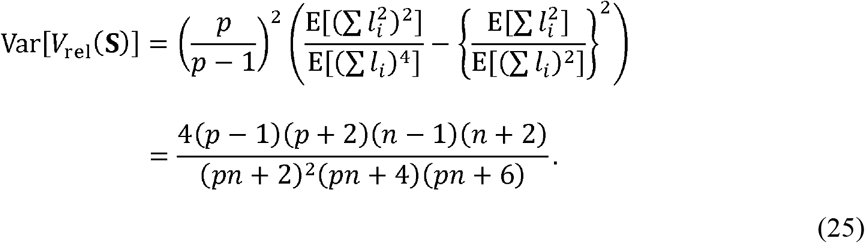

These results are exact (valid across any *p* and *n*) under multivariate normality.

#### Correlation matrix

Consider the null hypothesis **P** = **I**_*p*_ or ρ_*ij*_ = 0 for *i* ≠ *j*. The moments can conveniently be obtained from the form of average squared correlation (eq. 11). It is well known that, under the assumptions of normality and ρ_*ij*_ = 0 (for *i* ≠ *j*), 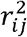 is distributed as Beta(1/2, (*n* − 1)/2), where *n* is the degree of freedom (e.g., Anderson, 2003). Therefore, under the null hypothesis,

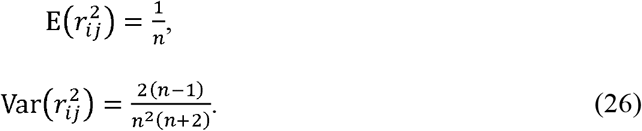

The expectation of *V*_rel_(**R**) is simply the average:

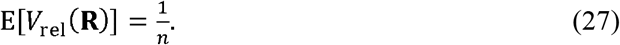

This expression is identical to (*p* − l)^−1^Var(*l_i_*) obtainable from Wagner’s (1984) results, except for having the degree of freedom *n* rather than the sample size *N* in the denominator. This is because Wagner (1984) considered *N* uncentered observations with mean 0 without explicitly distinguishing *n* and *N*. Most practical analyses would concern data centered at the sample mean, thus should use *n* rather than *N*.

Derivation of the variance is more complicated than it may seem, because, in principle,

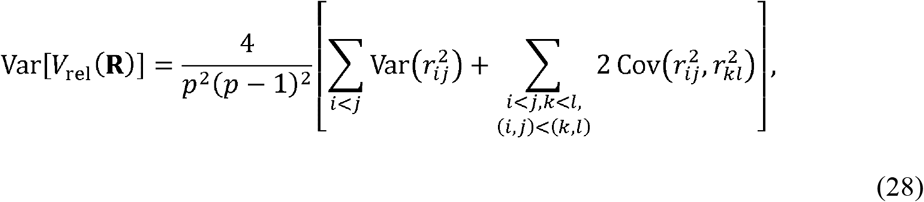

where the latter summation is across all non-redundant pairs. However, it is possible to show 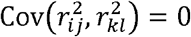 under the null hypothesis (Appendix C). Therefore, from equations 26 and 28,

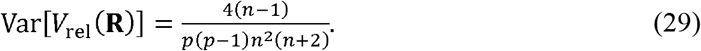

These expressions are exact for any *p* and *n*. Schott (2005) proposed a test for independence between sets of normal variables based on these moments.

### Moments under arbitrary conditions

#### Covariance matrix

This section considers moments of eigenvalue dispersion indices under arbitrary covariance/correlation structures and multivariate normality. It is straightforward to obtain the first two moments of *V*(**S**) under arbitrary **Σ**, provided that moments of relevant terms in equation 12 are available. The results are (Appendix B)

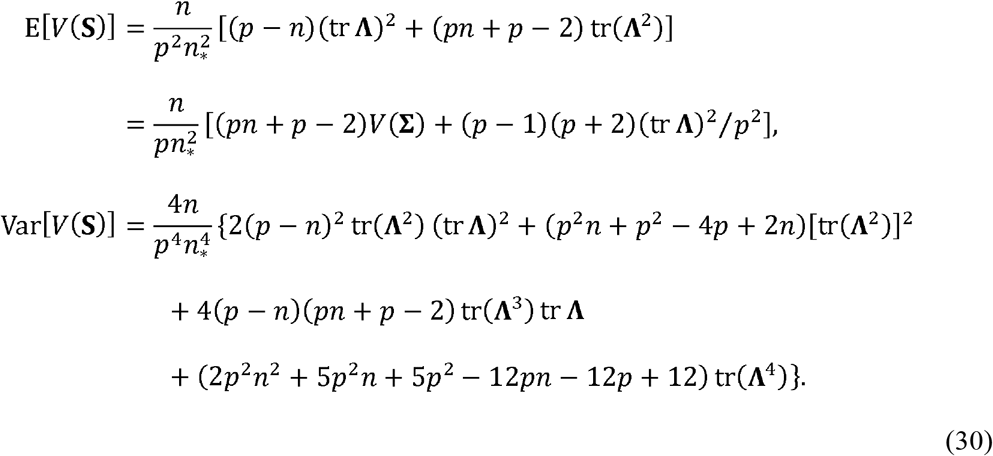

The second expression for the expectation comes from the fact *V*(**Σ**) =[*p* tr(**Λ**^2^) − (tr **Λ**)^2^]/*p*^2^, and clarifies that the expectation is a linear function of *V*(**Σ**). These results are exact, and it is easily verified that they reduce to equations 18 and 21 under the null hypothesis. Profiles of E[*V*(**S**)] across a range of *V*(**Σ**) are shown in Figure 2 (top row), under single large eigenvalue conditions with varying *p* and *N*, and fixed tr **Σ** (detailed conditions are described under simulation settings below).

**Figure 2.**
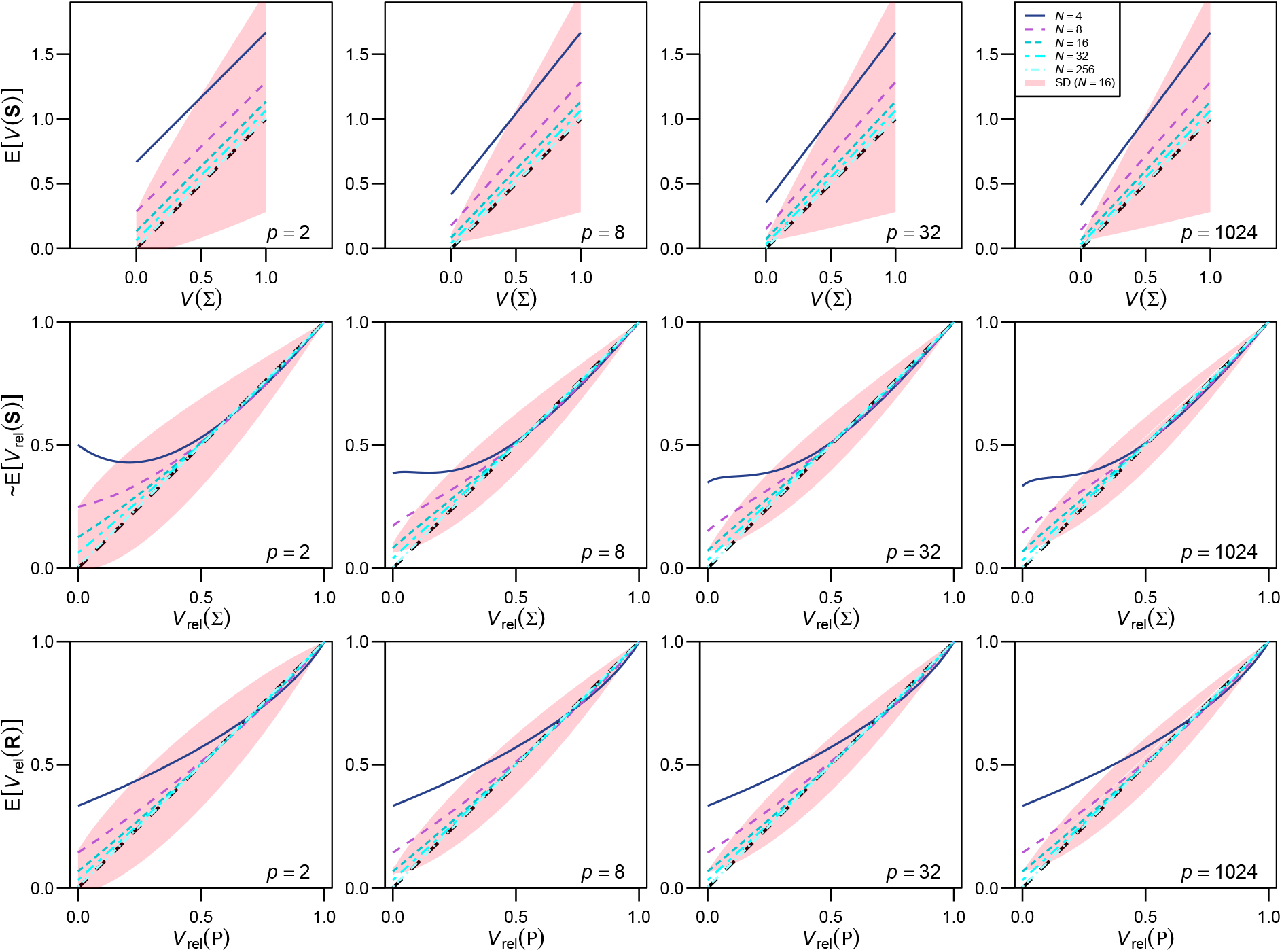
Profiles of the expectations of eigenvalue dispersion measures in selected conditions. The expectations of *V*(**S**) (top row), *V*_rel_(**S**) (approximate; middle row), and *V*_rel_(**R**) (bottom row) are drawn with solid lines, for *p* = 2, 8, 32, and 1024 (from left to right) and for *N* = 4, 8, 16, 32, and 256 (from top to bottom on the left end of each box). In all cases, *n* = *N* − 1. The breadth of one standard deviation at *N* = 16 is also shown around the mean profiles with pink fills; these are approximations for *V*_rel_(**S**) and for *V*_rel_(**R**) with *P* > 2 (the latter is from eq. 39; eqs. 36–38 yielded similar values under these conditions). Note that actual distributions might be skewed unlike these fills. There are generally many suites of eigenvalues corresponding to a single value of *V*_rel_ and E[*V*_rel_(**R**)] can also depend on eigenvector configurations; the profiles shown here are from such eigenvalue configurations that there is one large eigenvalue, with the rest being equally small, in which case E[*V*(**R**)| does not depend on eigenvector configurations. The population covariance matrix **Σ** is scaled so that tr **Σ** = *p*(*p* − 1)^−1/2^. The initial decrease of the E[*V*_rel_(**S**)] profiles in some cases seems to be an artifact of approximation. See text for further technical details.

Moments of *V*_rel_(**S**) are more difficult to obtain, as moments of the ratio 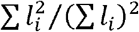 do not coincide with the ratios of moments under arbitrary **Σ**. Here we utilize the following approximations based on the delta method (e.g., Stuart & Ord, 1994: chapter 10):

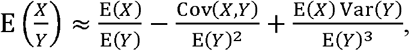

and

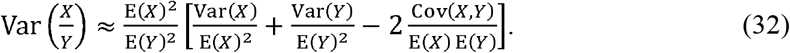

The approximate moments are (Appendix B):

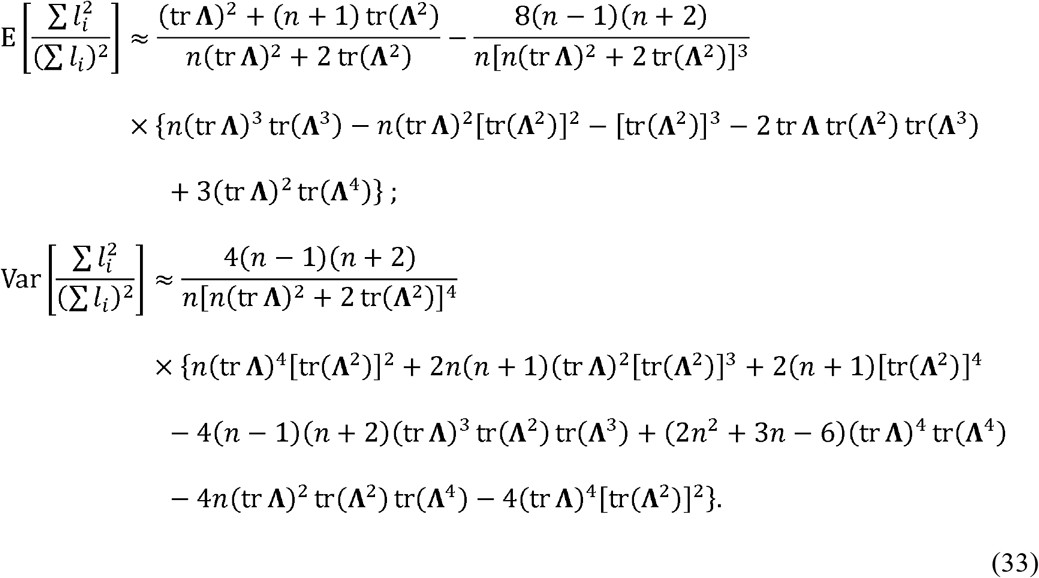

Inserting these into equation 22 yields the desired moments. The approximate expectation reduces to equation 24 under the null hypothesis, as the higher-order terms cancel out, whereas this is not the case for the approximate variance. Because these expressions are specified by the population eigenvalues alone, they are invariant with respect to orthogonal rotations, as expected from theoretical considerations above. Also, it is easily discerned that these expressions are invariant with respect to uniform scaling of the variables. The accuracy of these approximations will be examined in simulations below.

Profiles of the approximation of E[*V*_rel_(**S**)] across a range of *V*_rel_(**Σ**) are shown in Figure 2 (middle row) for the same conditions as above. The profiles are nonlinear; *V*_rel_(**S**) tends to overestimate *V*_rel_(**Σ**) when the latter is small, but tends to slightly underestimate when the latter is large. The initial decrease of E[*V*_rel_(**S**)] observed in some profiles appears to be an artifact of the approximation.

#### Correlation matrix

The expectation of *V*_rel_(**R**) under arbitrary conditions can be obtained from equation 11 with 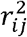 replaced by its expectations, which is known to be (e.g., Soper et al., 1917; Ghosh, 1966; Muirhead, 1982)

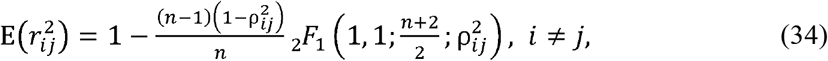

where

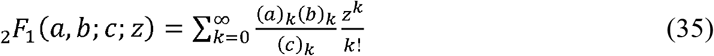

is the hypergeometric function, with (*x*)_*k*_ = *x*(*x* + 1) … (*x* + *k* − 1) denoting rising factorial (formally, (*x*)_*k*_ = Γ(*x* + *k*)/Γ(*x*) with the gamma function Γ(·)). Taking the average of equation 34 across all pairs of variables gives the desired expectation, which is non-asymptotic and exact. It is seen that equation 34 reduces to equation 26 under the null hypothesis.

When *p* = 2, the exact variance of *V*_rel_(**R**) is equal to that of the single squared correlation coefficient (Ghosh, 1966):

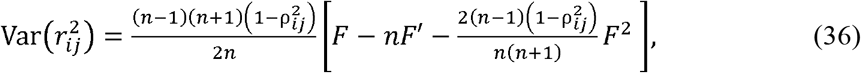

where 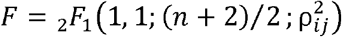 and 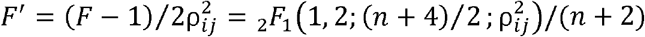; this last form is preferred to avoid numerical instability when 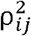 is close to 0. This expression reduces to equation 26 under the null hypothesis. When *p* > 2, we cannot ignore covariances between squared correlation coefficients (see eq. 28). Unfortunately, no exact expression seems available for this quantity in the literature, so we resort to approximation. One option is to utilize the following heuristic approximation from Pan & Frank’s (2004) approach (Appendix D):

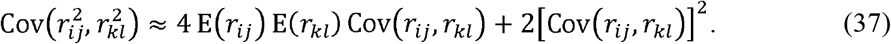

Exact and large-sample asymptotic expressions under multivariate normality are available for E(*r_ij_*) and Cov(*r_ij_, r_kl_*) respectively (e.g., Soper et al., 1917; Ghosh, 1966; Olkin & Siotani, 1976):

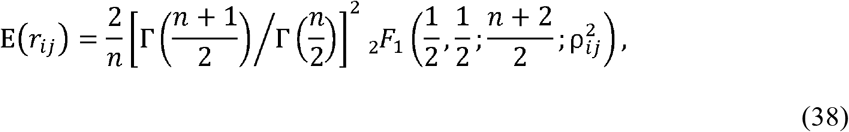

Inserting equations 36–38 to equation 28 yields an approximation for Var[*V*_rel_(**R**)] in arbitrary conditions. In terms of consistency, E(*r_ij_*) and 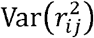 could be replaced by respective asymptotic expressions (e.g., Ghosh, 1966; Olkin & Siotani, 1976), but this does not seem to yield improved accuracy or substantial computational gain.

In practice, the variance based on equation 28 can be difficult to calculate for large *p*,because there are ~*p*^4^/4 pairs of correlation coefficients to evaluate. For such cases, the following asymptotic expression based on Konishi’s (1979) theory may be more useful (Appendix E):

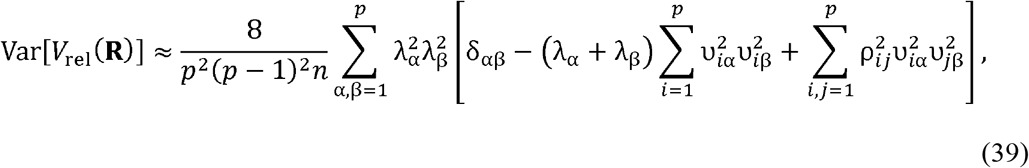

where δ_*ij*_ is the Kronecker delta (equals 1 for *i* = *j* and 0 otherwise) and υ_*i*α_ is the (*i*, α)-th element of the population eigenvector matrix **ϒ**. For *p* = 2, the accuracy of this expression can be compared with the exact expression (Fig. 3); visual inspection of the profiles suggest that the accuracy is satisfactory past *N* = 32– 64, except around *V*_rel_(**P**) = 0 where the asymptotic expression diminishes to 0 (as expected from its formula; Appendix E). For *p* > 2, accuracies of these approximate expressions are to be evaluated with simulations below.

**Figure 3.**
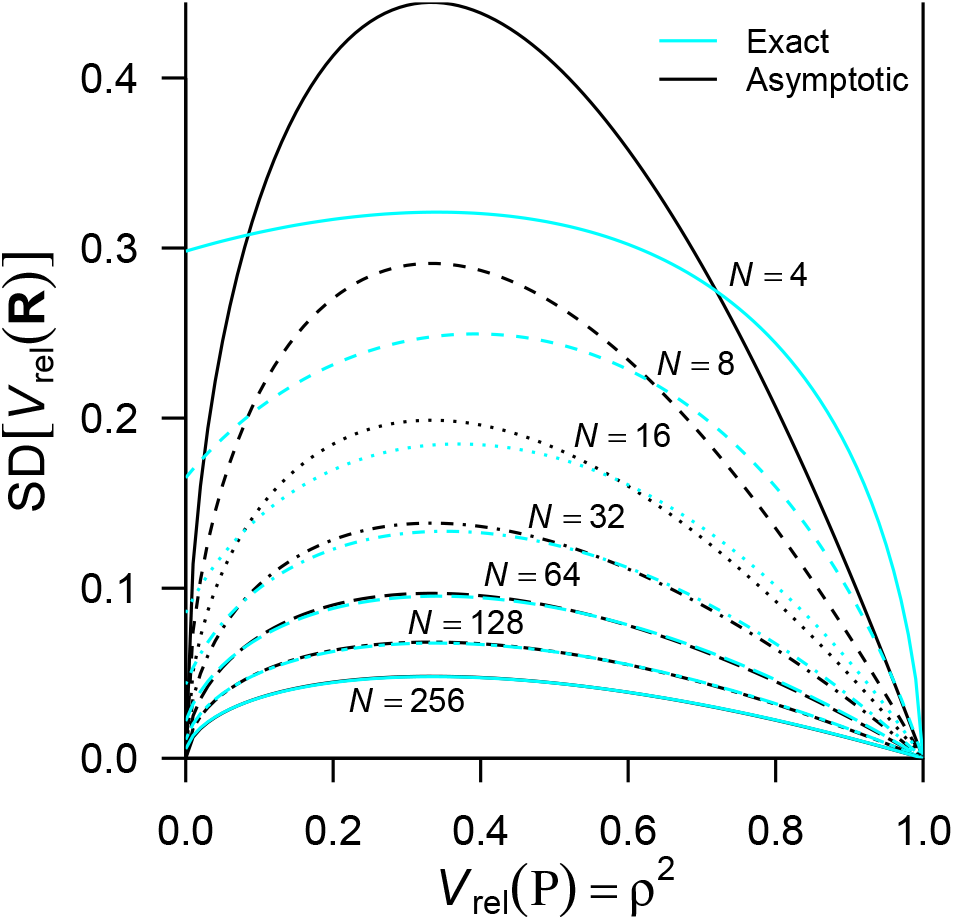
Comparison of exact and asymptotic standard deviations of *V*_rel_(**R**). Profiles of the exact (cyan lines; eq. 36) and asymptotic (black lines; eq. 39) standard deviations for *p* = 2 are shown across the entire range of the population value *V*_rel_(**P**), for *N* = 4, 8, 16, 32, 64, 128, and 256 (from top to bottom as labeled; shown with different line styles). Note that the asymptotic profiles converge to 0 when *V*_rel_(**P**) = 0.

Importantly, the expectation of *V*_rel_(**R**) is a function of ρ^2^’s rather than **Λ**, and cannot be specified by the latter alone in general. For instance, consider 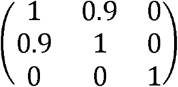 and 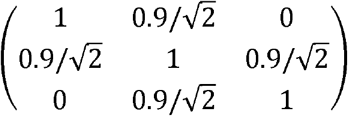, both of which are valid correlation matrices. These matrices have identical eigenvalues **Λ** = diag(1.9, 1.0, 0.1) and hence an identical value of *V*_rel_(**P**) (= 0.27), but E[*V*_rel_(**R**)] with *n* = 10 are 0.3326 and 0.3156, respectively. Although the difference diminishes as *n* increases, this example highlights that the distribution of *V*_rel_(**R**) is partly dependent on population eigenvectors.

Profiles of E[*V*_rel_(**R**)] across a range of *V*_rel_(**P**) are shown in Figure 2 (bottom row), under the same conditions as above. These conditions with single large eigenvalues are special cases in which E[*V*_rel_(**R**)| can be specified by *V*_rel_(**P**) regardless of eigenvectors (detailed in Appendix A). Indeed, the profiles of the expectations are invariant across *p* in these special conditions. In some way similar to *V*_rel_(**S**), *V*_rel_(**R**) tends to overestimate and underestimate small and large values of *V*_rel_(**P**), respectively.

### Bias correction

Some authors (Cheverud et al., 1989; Torices & Muñoz-Pajares, 2015) have suggested correcting the sampling bias in eigenvalue dispersion indices by means of subtracting Wagner’s (1984) null expectation from empirical values. This method could potentially be used for *V* and *V*_rel_ with the correct null expectations derived above, to obtain estimators that is unbiased under the null hypothesis. For *V*_rel_(**S**) and *V*_rel_(**R**), however, the subtraction truncates the upper end of the range, potentially compromising interpretability. To avoid this, it might be desirable to scale these indices in a way analogous to the adjusted coefficient of determination in regression analysis (e.g., Cramer, 1987):

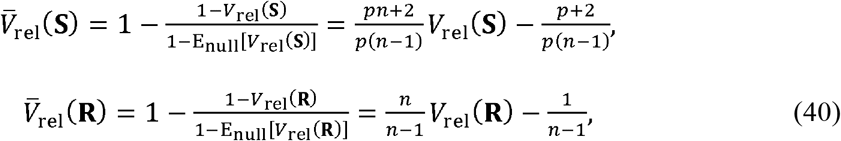

where E_null_(·) denotes expectation under the null conditions (eqs. 24 and 27). This adjustment inflates the variance by the factor of 1/[1 − E_null_(*V*_rel_)]^2^. Furthermore, these adjusted indices are unbiased only under the null hypothesis (Armbruster et al., 2009; and trivially the case of complete integration), and uniformly underestimate the corresponding population values otherwise (Fig. S2). As the population value gets away from 0, the adjusted index is outperformed by the unadjusted one in both precision and bias (Fig. S3). It should also be borne in mind that the profiles of expectations are nonlinear and dependent on *N* (Fig. 2). As the adjusted indices will be increasingly conservative for small *N*, it is questionable whether they can be used for comparing samples with different *N*, as originally intended by Cheverud et al. (1989). For these reasons, use of this adjustment would be restricted to estimation of the population value near 0 (up to 0.1–0.2, depending on *p* and *N*).

On the other hand, a global unbiased estimator of *V*(**Σ**) can be derived from above results:

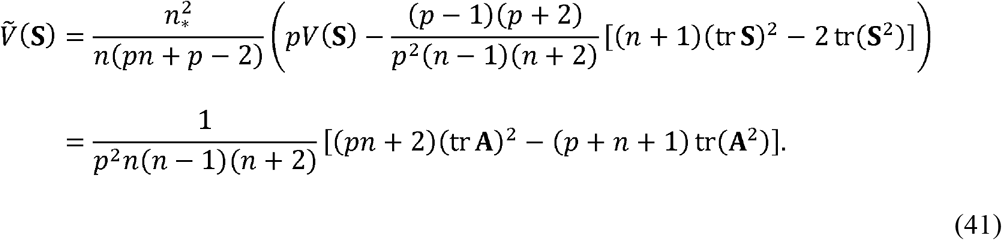

Its variance can be similarly obtained as

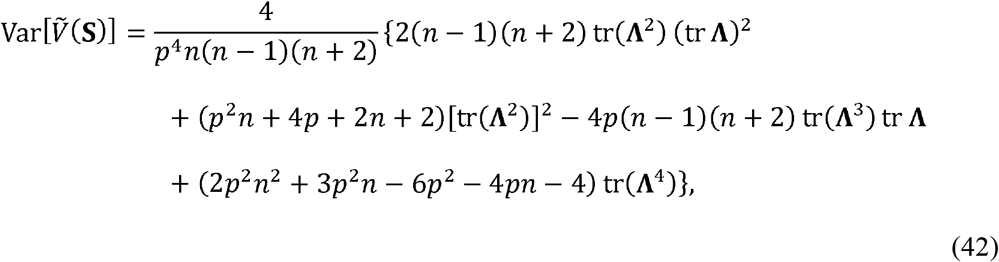

which reduces to 4(*p* − 1)(*p* + 2)σ^8^/*p*^3^*n*(*n* − 1)(*n* + 2) under the null hypothesis. Comparison with equations 21 and 30 suggests that this variance is smaller than that of *V*(**S**), especially under the null hypothesis. Therefore, 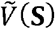 seems superior in both precision and bias and can be used when estimation of *V*(**Σ**) is desired. It can be used to compare multiple samples, provided that its sensitivity to overall scaling is not of concern, e.g., comparison between closely related taxa.

## Simulation

### Methods

Simulations were conducted under various conditions in order to understand sampling properties of the eigenvalue dispersion indices. All simulations were done assuming multivariate normality, with varying population covariance matrix **Σ**, number of variables *p* (= 2, 4, 8, 16, 32, 64, 128, 256, and 1024), and sample size *N* (= 4, 8, 16, 32, 64, 128, and 256).

For every *p*, the following population eigenvalue conformations were considered: 1) the null condition, 2) *q*-large λ conditions, 3) a linearly decreasing λ condition, and 4) a quadratically decreasing λ condition (see Fig. 4 for examples). The null condition is where all population eigenvalues are equal in magnitude (*V*_rel_(**Σ**) = 0), corresponding to the null hypothesis of sphericity (Fig. 4A). The *q*-large λ conditions are where the first *q* (= 1, 2, and 4, provided *p* > *q*) population eigenvalues are equally large and the remaining *p* − *q* ones are equally small (λ_1_ = ··· = λ_*q*_ > λ_*q*+1_ = ··· = λ_*p*_), with varying *V*_rel_(**Σ**) (= 0.1, 0.2, 0.4, 0.6, and 0.8; Fig. 4B–G). The necessary condition λ_*p*_ ≥ 0 constrains possible combinations of *q* and *V*_rel_(**Σ**): the possible choices of *V*_rel_(**Σ**) are 0.1–0.8, 0.1–0.4, and 0.1–0.2 for *q* = 1, 2, and 4, respectively (Appendix A). These conditions are intended to represent hypothetical situations where only a few components of meaningful signals are present in the population. Individual eigenvalues were calculated for each combination of *p, q*, and *V*_rel_(**Σ**) as described in Appendix A. The linearly and quadratically decreasing λ conditions are where the population eigenvalues are linearly and quadratically, respectively, decreasing in magnitude (Fig. 4H; Appendix A), in which cases the value of *V*_rel_(**Σ**) is fixed for a given *p*. These conditions are intended to represent covariance structures with gradually decreasing signals. One might claim that some of these situations, especially *q*-large λ conditions, are too simplistic and biologically unrealistic, but these simple settings enable us to clarify systematic relationships between parameters and sampling properties. The primary aim here is to explore sampling properties across a wide range of parameters, rather than focusing on biologically “realistic” regions (which would depend on specific organismal systems). It should also be recalled that sampling error alone can yield gradually decreasing patterns of sample eigenvalues typically observed in empirical datasets (see above and below).

**Figure 4.**
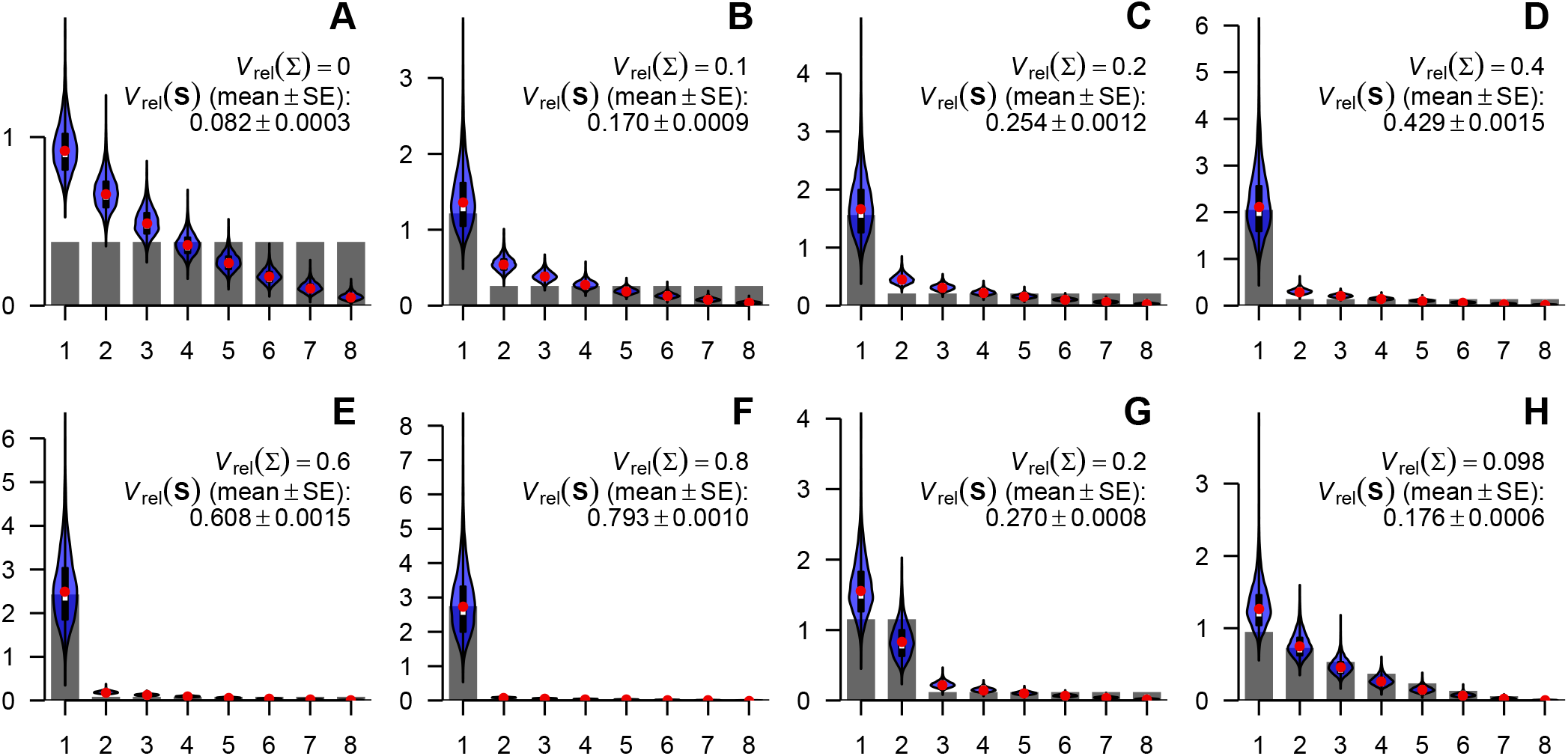
Selected population eigenvalue structures used in simulations and distributions of sample eigenvalues, examples for *p* = 8. The eigenvalues of population covariance matrix are shown as scree plots, and distributions of sample eigenvalues with *N* = 16 are shown as violin plots. **A**, null condition; **B–G**, *q*-large λ conditions, *q* = 1 (**B–F**) or 2 (**G**), with *V*_rel_(**Σ**) = 0.1, 0.2, 0.4, 0.6, 0.8, and 0.2, respectively; **H**, quadratically decreasing λ condition. Red dots denote empirical means of sample eigenvalues, whereas white bars (mostly overlapping with red dots) denote medians. Thick black bars within violins denote interquartile ranges. Note different scales of vertical axes.

For sake of simplicity, all population covariance matrices were scaled to ensure *V*(**Σ**) = *V*_rel_(**Σ**); that is, tr **Σ** = *p*(*p* − 1)^−1/2^. This scaling also makes the magnitude of *V*(**Σ**) comparable across varying *p*. In addition, a population covariance matrix **Σ** was constructed from a predefined set of eigenvalues such that its diagonal elements are equal: 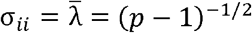 for all *i*, thereby enforcing **Σ** = (*p* − 1)^−1/2^**P**. This construction allows for examining both covariance and correlation matrices with the same population *V*_rel_ from a single simulated dataset, saving computational resources. **Σ** was constructed from **Λ** by the iterative Givens rotation algorithm of Davies & Higham (2000), which is guaranteed to converge within *p* − 1 iterations. This algorithm was implemented as coded by Waller (2020), but with the following modifications for reproducibility: no random orthogonal rotation was involved at the initial stage, and rotation axes were chosen in a fixed order. It should be noted that the rotations involved—choice of eigenvectors—would in general influence distributions of *V*_rel_(**R**), except for certain special cases including the 1-large λ condition (Appendix A). It is impractical to exhaustively examine numerous possible conformations of eigenvectors, so only the single conformation generated by this algorithm was used for each combination of parameters.

The eigenvalues of a sample covariance matrix were obtained from singular value decomposition of the data matrix, as the singular values squared and then divided by *n*_*_ (see, e.g., Jolliffe, 2002). When *p* > *N* − 1, 0’s were appended to this vector so that *p* sample eigenvalues were present. Data were centered at the sample mean before the decomposition, therefore *n* = *N* − 1. It was chosen that *n*_*_ = *n*. The eigenvalues for a sample correlation matrix were obtained similarly from the sample-mean-centered data scaled with the sample standard deviation for each variable.

To summarize, each set of simulations consists of the following steps: 1) define a desired set of eigenvalues **Λ**; 2) construct the population covariance matrix **Σ** with the rotation algorithm explained above; 3) generate *N* i.i.d. normal observations from *N_p_*(**0, Σ**); 4) eigenvalues of sample covariance and correlation matrices were obtained from singular value decomposition of the sample-mean-centered data; 5) *V*(**S**), *V*_rel_(**S**), and *V*_rel_(**R**) were calculated from the eigenvalues; 6) the steps 3–5 were iterated for 5,000 times in total with the same *N* and **Σ**. The simulations were conducted on the R environment version 3.5.3 (R Core Team, 2019). The function “genhypergeo” of the package “hypergeo” (Hankin, 2015) was used to evaluate the hypergeometric function in the moments of *V*_rel_(**R**). The time-consuming calculation of Var[*V*_rel_(**R**)] from equations 28 and 36–38 was aided by a C++ code via the package “Rcpp” (Eddelbuettel & Balamuta, 2018). The codes are provided as Supplementary Material.

### Results

Examined individually, sample eigenvalues were biased estimators of population eigenvalues, as expected. Selected eigenvalue distributions of sample covariance and correlation matrices are shown in Figures 4 and S1, respectively. Typically, the first few eigenvalues were overestimated, with the rest being underestimated. Note that gradually decreasing scree-like profiles of sample eigenvalues typical of empirical datasets can arise even when most population eigenvalues are identical in magnitude. The sampling biases decreases as *N* increases. These overall trends were similarly observed for correlation matrices, although the upper tail of the largest eigenvalue tended to be truncated for correlation matrices because of the constraint tr **R** = *p*, effectively cancelling the tendency of overestimation in this eigenvalue (Fig. S1).

Sampling distributions of *V*(**S**) are shown in Figures 5 and S4–S6, and their summary statistics are shown in Tables 1 and S1. Distributions were unimodal but highly skewed with long upper tails, especially when *N* or *p* is small. As expected, sampling dispersion decreases consistently with increasing *N*, with skewness decreasing at the same time. Interestingly, the shape of distribution does not visibly change with increasing *p*, at least with moderately large *N* (≥ 32, say). In all conditions, *V*(**S**) tended to overestimate the population value *V*(**Σ**). Increasing *V*(**Σ**) drastically increased sampling dispersion and skewness, whereas increasing *q* with a fixed *V*(**Σ**) decreased sampling dispersion without changing the mean as much. Sampling distributions of *V*(**S**) under linearly and quadratically decreasing λ conditions look similar to those under *q*-large λ conditions with similar *V*(**Σ**) values for the respective *p*. The expressions of the expectation and variance of *V*(**S**) almost always coincided with the sampling mean and variance within a reasonable range of random fluctuations (as expected, since those results are exact).

**Figure 5.**
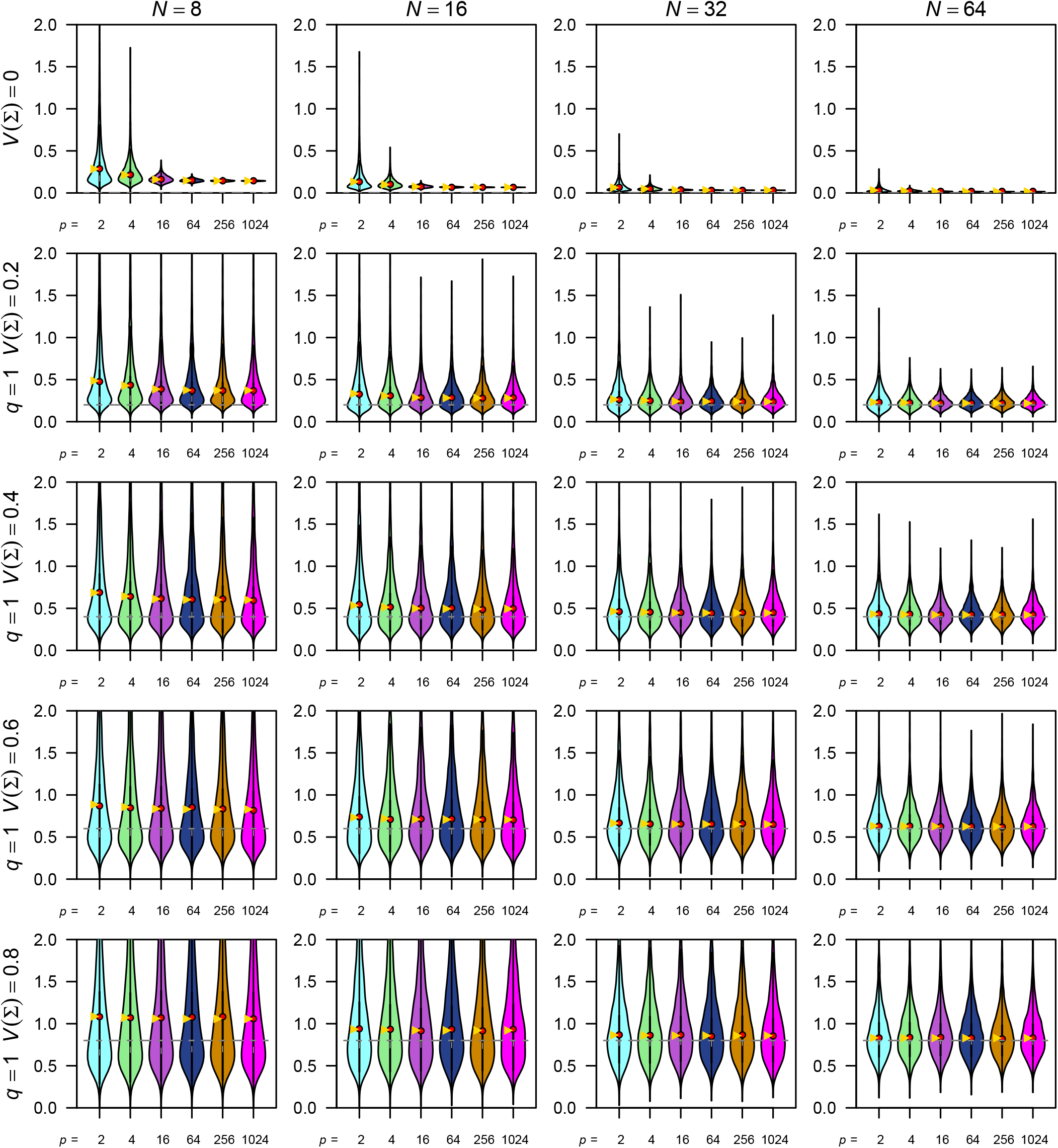
Selected simulation results for the eigenvalue variance of covariance matrix *V*(**S**). Empirical distributions of simulated *V*(**S**) values are shown as violin plots, whose tails extend to the extreme values. Red dots denote empirical means, whereas yellow triangles denote expectations (which are exact). Thick black bars within violins denote interquartile ranges, with white bars near the center (in some cases overlapping with red dots) denote medians. Rows of panels correspond to varying population values of *V*(**Σ**) (under 1-large λ conditions), whereas columns correspond to varying sample size *N*. Columns within each panel correspond to varying number of variables *p*. Note that extreme values in some panels are cropped for visual clarity. See Figure S4–S6 for full results.

**Table 1.**
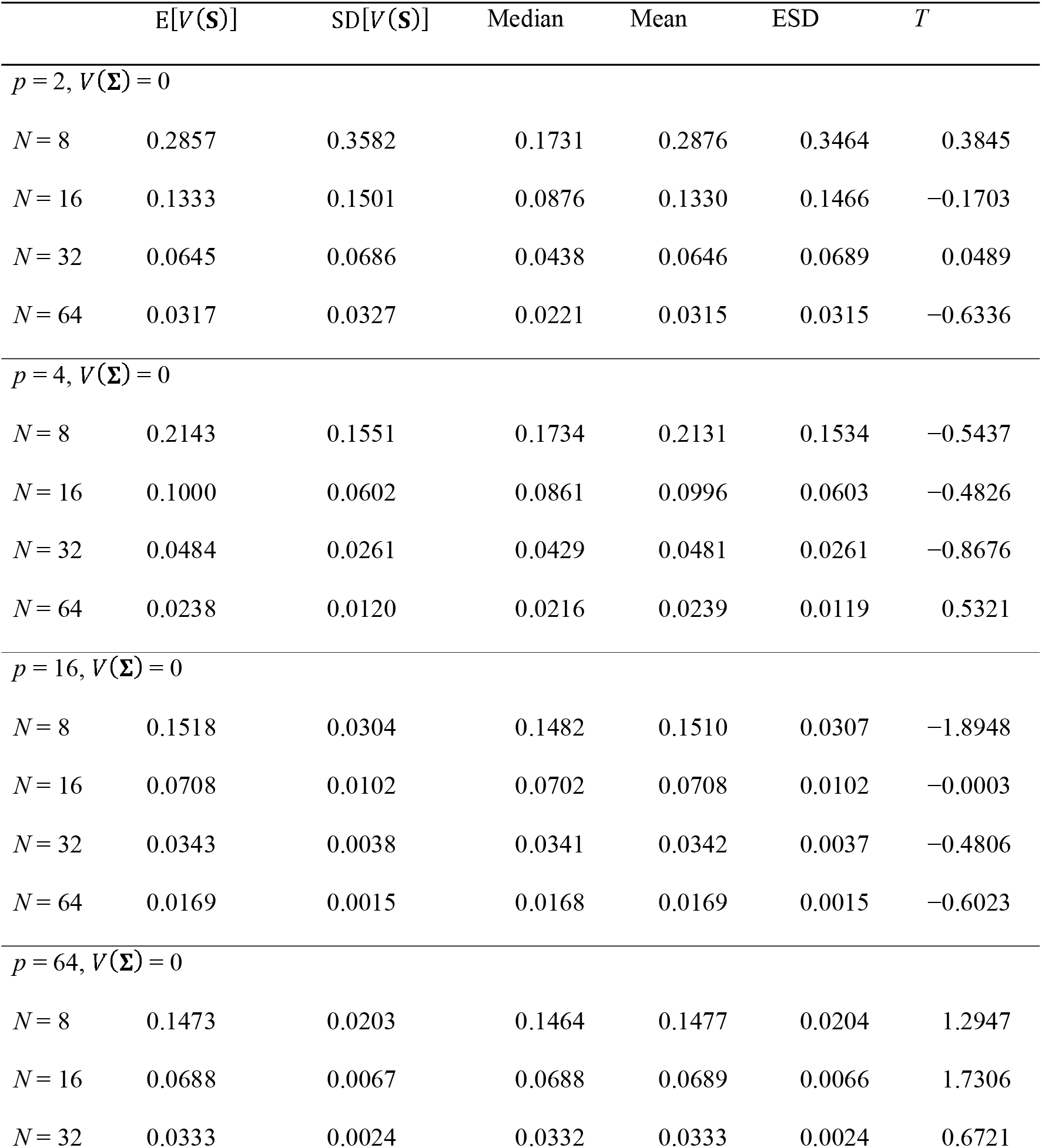
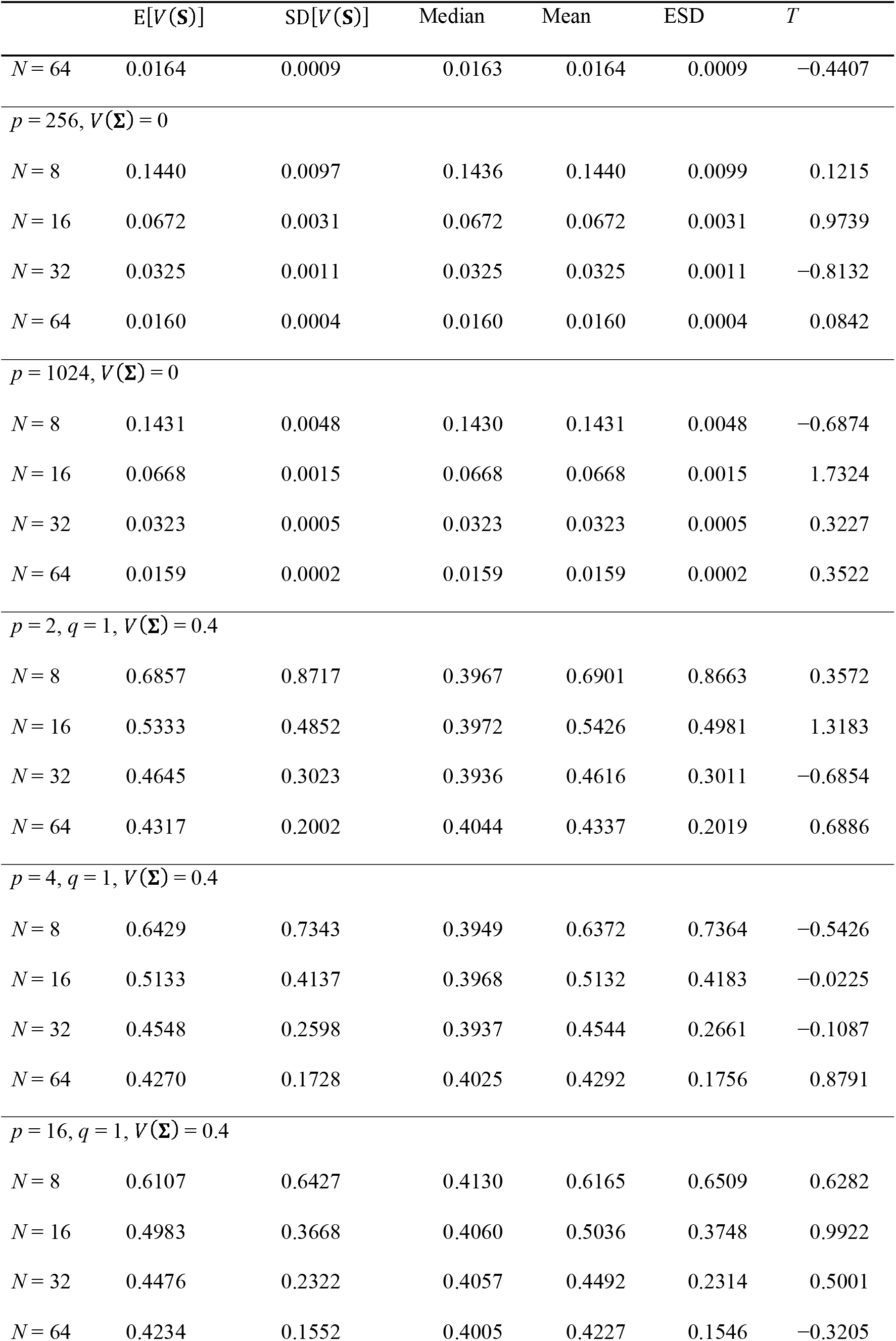
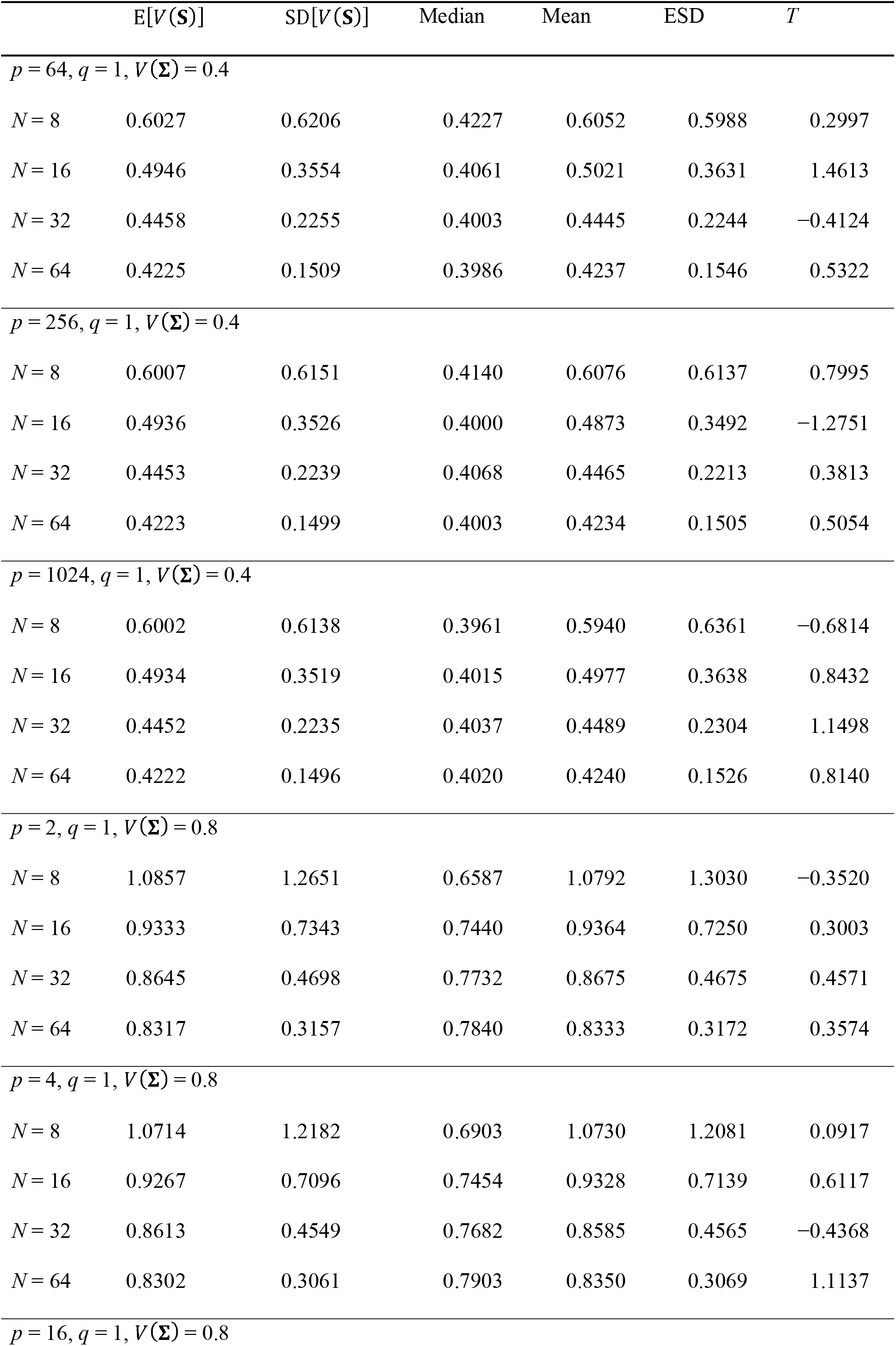
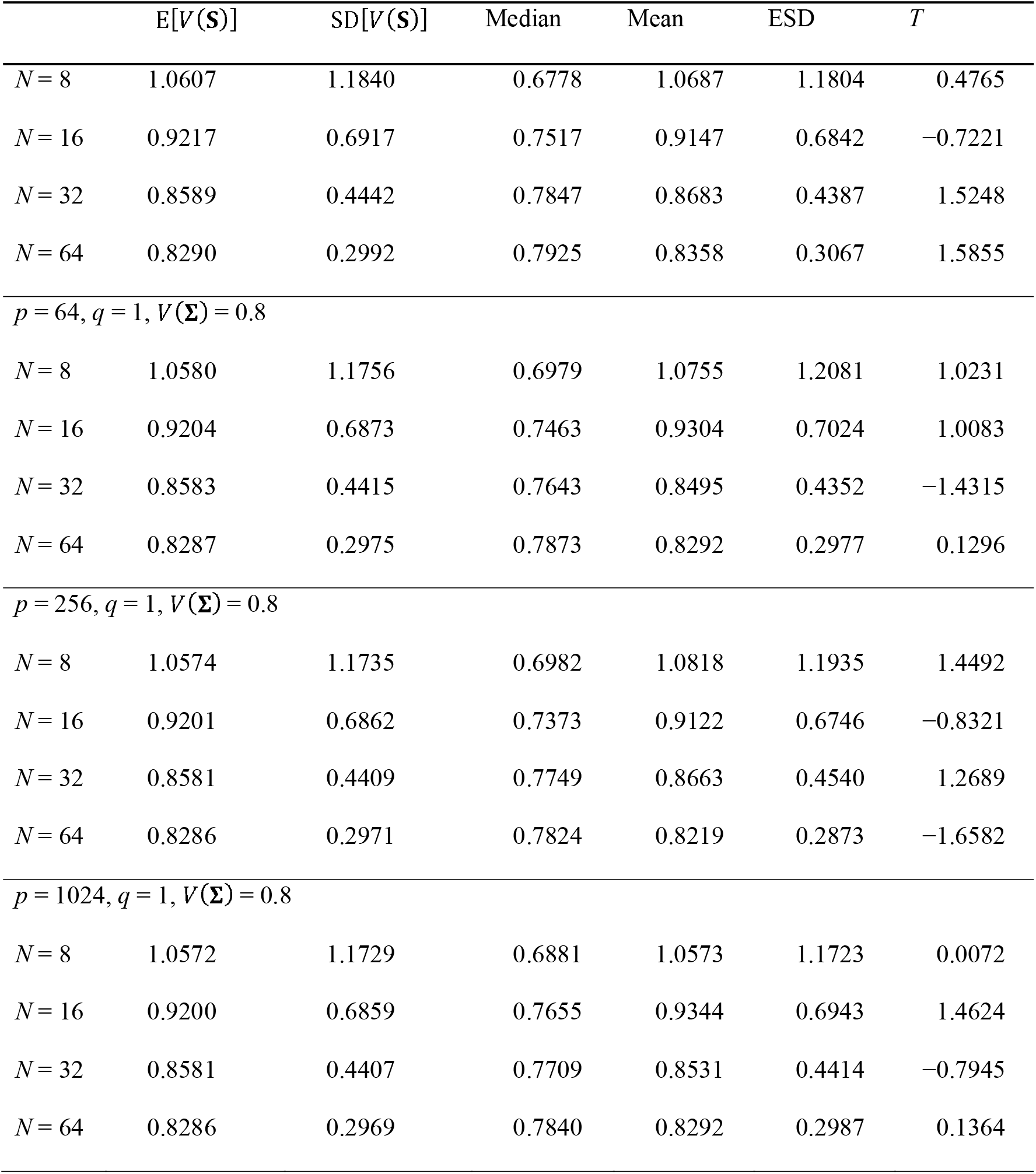
Summary of selected simulation results for eigenvalue variance of covariance matrix *V*(**S**). Theoretical expectation (E[*V*(**S**)]) and standard deviation (SD[*V*(**S**)]), as well as empirical median, mean, standard deviation (ESD), and bias of mean in standard error unit (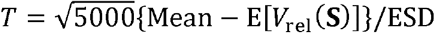, which should roughly follow *t* distribution with 4999 degrees of freedom if the expectation is exact) from 5000 simulation runs are shown for selected conditions. See Table S1 for full results.

Results for *V*_rel_(**S**) are summarized in Figures 6 and S6–S8 and Tables 2 and S2. Distributions were unimodal within the range (0, 1), except when *N* = 4 and *p* = 2 where the distribution was essentially uniform. As was the case for *V*_rel_(**S**), the sampling dispersion of *V*_rel_(**S**) decreased drastically with increasing *N*, and to some extent with increasing *p*, while the shape of distribution does not seem to change remarkably with increasing *p* past certain *N*. *V*_rel_(**S**) tended to overestimate the population value *V*_rel_(**Σ**), except when the latter is rather large (= 0.8) where slight underestimation was observed. With increasing *q* for a fixed *V*_rel_(**Σ**) the distributions tended to shrink, but the sampling bias remained virtually unchanged or slightly increased. In the null conditions, the exact expressions of the expectation and variance performed perfectly (as expected). The approximate expectation for arbitrary conditions derived above yielded substantially smaller values than the empirical means when *N* is small; however, the approximation worked satisfactorily with moderate *N* (≥ 16–32), with the deviations from empirical means mostly falling within 2 standard error units. In addition, the approximate expectation worked rather well, even with small *N*, under either A) the *q*-large λ conditions with *q* = 2 and *V*_rel_(**Σ**) = 0.4, B) same with *q* = 4, or C) linearly and quadratically decreasing λ conditions with moderately large *p* (≥ 16). Other conditions held constant, the accuracy of the approximate expectation in absolute scale tended to slightly improve with increasing *p*, effectively balancing with the decreasing sampling dispersion, so that the relative bias in standard error unit remains almost invariant across varying *p*. The approximate variance for arbitrary conditions derived above yielded substantially larger values than the empirical variance, except under the *q*-large λ conditions with *q* = 1 and *V*_rel_(**Σ**) = 0.8 where it yielded smaller values. Nevertheless, with moderately large *N* (≥ 64), the inaccuracy typically decreased to <5% in the scale of standard deviation (SD scale hereafter), except under the linearly and quadratically decreasing λ conditions with moderately large *p* (≥ 16), where it was more accurate.

**Figure 6.**
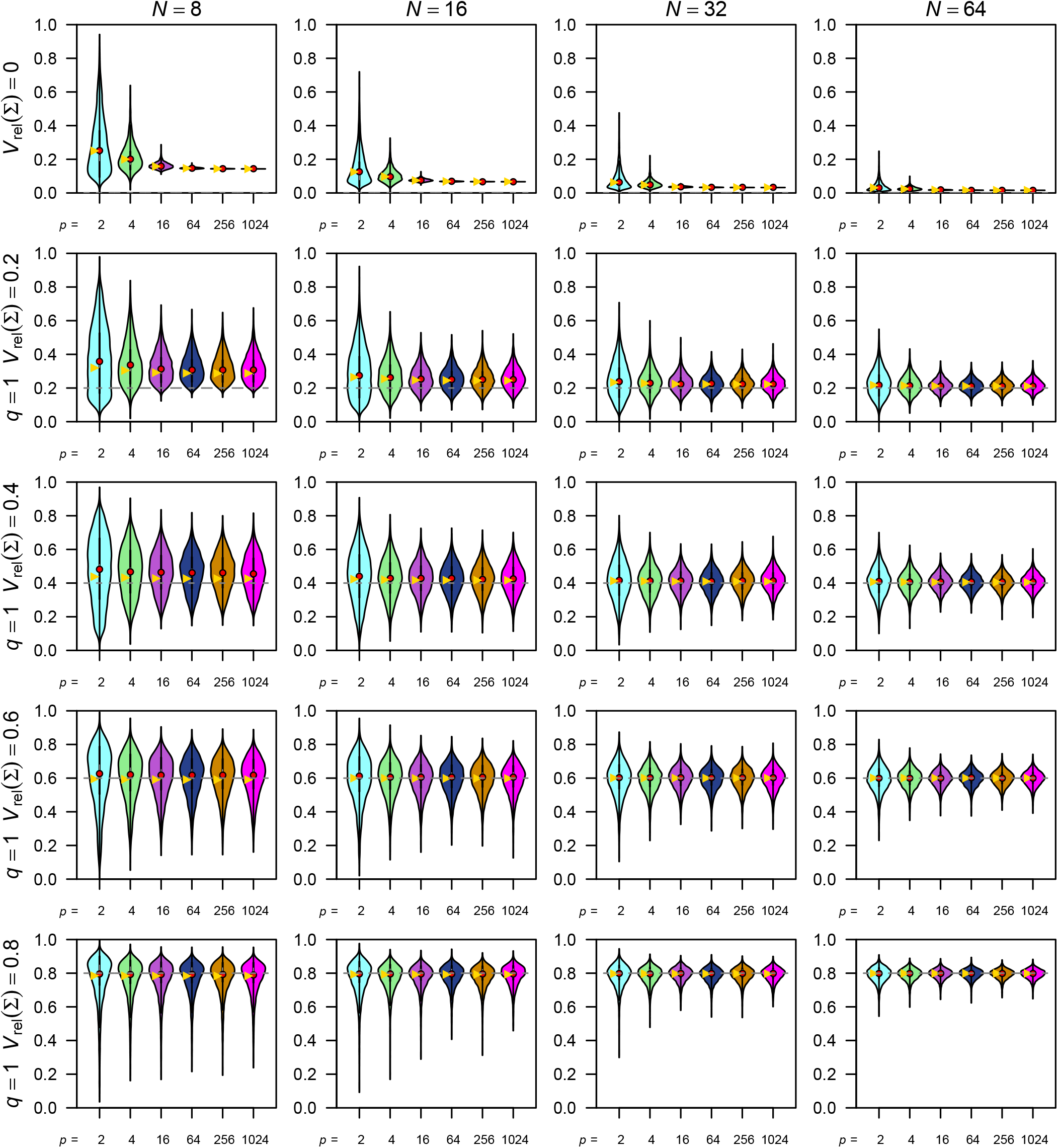
Selected simulation results for the relative eigenvalue variance of covariance matrix *V*_rel_(**S**). Empirical distributions of simulated *V*_rel_(**S**) values are shown as violin plots. Yellow triangles denote expectations (which are approximate except under the null condition). Rows of panels correspond to varying population values of *V*_rel_(**Σ**) (under 1-large λ conditions). Other legends are as in Fig. 5. See Figure S6–S8 for full results.

**Table 2.** Summary of selected simulation results for relative eigenvalue variance of covariance matrix *V*_rel_(**S**). (Approximate) theoretical expectation (E[*V*_rel_(**S**)]) and standard deviation (SD[*V*_rel_(**S**)]), as well as empirical median, mean, standard deviation (ESD), and bias of mean in standard error unit (*T*) from 5000 simulation runs are shown for selected conditions. See Table 1 for further information and Table S2 for full results.

| | $\approx E[V_{\text{rel}}(\mathbf{S})]$ | $\approx SD[V_{\text{rel}}(\mathbf{S})]$ | Median | Mean | ESD | $T$ |
| --- | --- | --- | --- | --- | --- | --- |
| $p = 2, V_{\text{rel}}(\mathbf{\Sigma}) = 0$ | | | | | | |
| $N = 8$ | 0.2500 | 0.1936 | 0.2079 | 0.2514 | 0.1924 | 0.4961 |
| $N = 16$ | 0.1250 | 0.1102 | 0.0953 | 0.1259 | 0.1098 | 0.6000 |
| $N = 32$ | 0.0625 | 0.0587 | 0.0448 | 0.0628 | 0.0597 | 0.3234 |
| $N = 64$ | 0.0313 | 0.0303 | 0.0224 | 0.0309 | 0.0289 | -0.8301 |
| $p = 4, V_{\text{rel}}(\mathbf{\Sigma}) = 0$ | | | | | | |
| $N = 8$ | 0.2000 | 0.0840 | 0.1856 | 0.2002 | 0.0858 | 0.1644 |
| $N = 16$ | 0.0968 | 0.0433 | 0.0907 | 0.0968 | 0.0434 | 0.0620 |
| $N = 32$ | 0.0476 | 0.0219 | 0.0438 | 0.0474 | 0.0221 | -0.6146 |
| $N = 64$ | 0.0236 | 0.0110 | 0.0218 | 0.0237 | 0.0110 | 0.6544 |
| $p = 16, V_{\text{rel}}(\mathbf{\Sigma}) = 0$ | | | | | | |
| $N = 8$ | 0.1579 | 0.0193 | 0.1562 | 0.1580 | 0.0197 | 0.3764 |
| $N = 16$ | 0.0744 | 0.0091 | 0.0738 | 0.0746 | 0.0094 | 1.4158 |
| $N = 32$ | 0.0361 | 0.0044 | 0.0357 | 0.0361 | 0.0044 | -1.2604 |
| $N = 64$ | 0.0178 | 0.0022 | 0.0177 | 0.0178 | 0.0022 | 0.4338 |
| $p = 64, V_{\text{rel}}(\mathbf{\Sigma}) = 0$ | | | | | | |
| $N = 8$ | 0.1467 | 0.0047 | 0.1462 | 0.1466 | 0.0046 | -0.6909 |
| $N = 16$ | 0.0686 | 0.0022 | 0.0685 | 0.0686 | 0.0022 | 0.5618 |
| $N = 32$ | 0.0332 | 0.0010 | 0.0332 | 0.0332 | 0.0011 | 0.0446 |
| $N = 64$ | 0.0164 | 0.0005 | 0.0163 | 0.0164 | 0.0005 | -0.5869 |

Table 2 (continued)
| | $\approx E[V_{\text{rel}}(\mathbf{S})]$ | $\approx \text{SD}[V_{\text{rel}}(\mathbf{S})]$ | Median | Mean | ESD | $T$ |
| --- | --- | --- | --- | --- | --- | --- |
| $p = 256, V_{\text{rel}}(\mathbf{\Sigma}) = 0$ | | | | | | |
| $N = 8$ | 0.1438 | 0.0012 | 0.1437 | 0.1438 | 0.0012 | -0.0173 |
| $N = 16$ | 0.0672 | 0.0005 | 0.0671 | 0.0672 | 0.0005 | 1.4545 |
| $N = 32$ | 0.0325 | 0.0003 | 0.0325 | 0.0325 | 0.0003 | -0.7481 |
| $N = 64$ | 0.0160 | 0.0001 | 0.0160 | 0.0160 | 0.0001 | 0.3608 |
| $p = 1024, V_{\text{rel}}(\mathbf{\Sigma}) = 0$ | | | | | | |
| $N = 8$ | 0.1431 | 0.0003 | 0.1431 | 0.1431 | 0.0003 | -0.5105 |
| $N = 16$ | 0.0668 | 0.0001 | 0.0668 | 0.0668 | 0.0001 | 0.0231 |
| $N = 32$ | 0.0323 | 0.0001 | 0.0323 | 0.0323 | 0.0001 | -1.3055 |
| $N = 64$ | 0.0159 | 0.0000 | 0.0159 | 0.0159 | 0.0000 | -0.0022 |
| $p = 2, q = 1, V_{\text{rel}}(\mathbf{\Sigma}) = 0.4$ | | | | | | |
| $N = 8$ | 0.4377 | 0.2825 | 0.4939 | 0.4804 | 0.2334 | 12.9356 |
| $N = 16$ | 0.4232 | 0.1982 | 0.4457 | 0.4384 | 0.1764 | 6.0670 |
| $N = 32$ | 0.4132 | 0.1378 | 0.4169 | 0.4152 | 0.1297 | 1.1224 |
| $N = 64$ | 0.4070 | 0.0963 | 0.4108 | 0.4083 | 0.0945 | 0.9214 |
| $p = 4, q = 1, V_{\text{rel}}(\mathbf{\Sigma}) = 0.4$ | | | | | | |
| $N = 8$ | 0.4319 | 0.2065 | 0.4730 | 0.4675 | 0.1658 | 15.1734 |
| $N = 16$ | 0.4200 | 0.1444 | 0.4316 | 0.4288 | 0.1272 | 4.8902 |
| $N = 32$ | 0.4113 | 0.1001 | 0.4147 | 0.4125 | 0.0944 | 0.9058 |
| $N = 64$ | 0.4060 | 0.0699 | 0.4073 | 0.4067 | 0.0682 | 0.7503 |
| $p = 16, q = 1, V_{\text{rel}}(\mathbf{\Sigma}) = 0.4$ | | | | | | |
| $N = 8$ | 0.4275 | 0.1691 | 0.4657 | 0.4626 | 0.1346 | 18.4219 |
| $N = 16$ | 0.4174 | 0.1175 | 0.4302 | 0.4270 | 0.1049 | 6.4710 |
| $N = 32$ | 0.4098 | 0.0813 | 0.4161 | 0.4129 | 0.0762 | 2.8682 |
| $N = 64$ | 0.4052 | 0.0566 | 0.4070 | 0.4058 | 0.0548 | 0.7118 |
| $p = 64, q = 1, V_{\text{rel}}(\mathbf{\Sigma}) = 0.4$ | | | | | | |

Table 2 (continued)
| | $\approx E[V_{\text{rel}}(\mathbf{S})]$ | $\approx \text{SD}[V_{\text{rel}}(\mathbf{S})]$ | Median | Mean | ESD | $T$ |
| --- | --- | --- | --- | --- | --- | --- |
| $N = 8$ | 0.4264 | 0.1613 | 0.4670 | 0.4599 | 0.1273 | 18.6017 |
| $N = 16$ | 0.4168 | 0.1119 | 0.4286 | 0.4266 | 0.0996 | 6.9651 |
| $N = 32$ | 0.4095 | 0.0773 | 0.4125 | 0.4109 | 0.0726 | 1.3497 |
| $N = 64$ | 0.4050 | 0.0538 | 0.4063 | 0.4056 | 0.0532 | 0.7287 |
| $p = 256, q = 1, V_{\text{rel}}(\mathbf{\Sigma}) = 0.4$ | | | | | | |
| $N = 8$ | 0.4261 | 0.1594 | 0.4637 | 0.4606 | 0.1249 | 19.5222 |
| $N = 16$ | 0.4166 | 0.1105 | 0.4262 | 0.4228 | 0.0977 | 4.4222 |
| $N = 32$ | 0.4094 | 0.0763 | 0.4148 | 0.4120 | 0.0718 | 2.5791 |
| $N = 64$ | 0.4050 | 0.0531 | 0.4066 | 0.4058 | 0.0520 | 1.1405 |
| $p = 1024, q = 1, V_{\text{rel}}(\mathbf{\Sigma}) = 0.4$ | | | | | | |
| $N = 8$ | 0.4261 | 0.1590 | 0.4566 | 0.4548 | 0.1261 | 16.1041 |
| $N = 16$ | 0.4166 | 0.1102 | 0.4272 | 0.4246 | 0.0990 | 5.7314 |
| $N = 32$ | 0.4094 | 0.0761 | 0.4140 | 0.4123 | 0.0718 | 2.8690 |
| $N = 64$ | 0.4050 | 0.0530 | 0.4068 | 0.4058 | 0.0521 | 1.1494 |
| $p = 2, q = 1, V_{\text{rel}}(\mathbf{\Sigma}) = 0.8$ | | | | | | |
| $N = 8$ | 0.7847 | 0.1153 | 0.8314 | 0.7947 | 0.1473 | 4.8016 |
| $N = 16$ | 0.7929 | 0.0866 | 0.8164 | 0.7963 | 0.1004 | 2.4327 |
| $N = 32$ | 0.7969 | 0.0625 | 0.8090 | 0.7990 | 0.0672 | 2.2207 |
| $N = 64$ | 0.7986 | 0.0445 | 0.8030 | 0.7983 | 0.0470 | -0.3966 |
| $p = 4, q = 1, V_{\text{rel}}(\mathbf{\Sigma}) = 0.8$ | | | | | | |
| $N = 8$ | 0.7841 | 0.0920 | 0.8207 | 0.7927 | 0.1170 | 5.1526 |
| $N = 16$ | 0.7927 | 0.0692 | 0.8086 | 0.7945 | 0.0793 | 1.6175 |
| $N = 32$ | 0.7968 | 0.0498 | 0.8039 | 0.7967 | 0.0537 | -0.1966 |
| $N = 64$ | 0.7985 | 0.0354 | 0.8026 | 0.7993 | 0.0368 | 1.5352 |
| $p = 16, q = 1, V_{\text{rel}}(\mathbf{\Sigma}) = 0.8$ | | | | | | |
| $N = 8$ | 0.7838 | 0.0810 | 0.8153 | 0.7913 | 0.1049 | 5.0688 |

Table 2 (continued)
| | $\approx E[V_{\text{rel}}(\mathbf{S})]$ | $\approx \text{SD}[V_{\text{rel}}(\mathbf{S})]$ | Median | Mean | ESD | $T$ |
| --- | --- | --- | --- | --- | --- | --- |
| $N = 16$ | 0.7926 | 0.0609 | 0.8067 | 0.7943 | 0.0696 | 1.7415 |
| $N = 32$ | 0.7968 | 0.0437 | 0.8044 | 0.7989 | 0.0452 | 3.3474 |
| $N = 64$ | 0.7985 | 0.0310 | 0.8019 | 0.7990 | 0.0323 | 1.0675 |
| $p = 64, q = 1, V_{\text{rel}}(\mathbf{\Sigma}) = 0.8$ | | | | | | |
| $N = 8$ | 0.7837 | 0.0787 | 0.8157 | 0.7915 | 0.1022 | 5.4258 |
| $N = 16$ | 0.7926 | 0.0592 | 0.8072 | 0.7956 | 0.0669 | 3.2349 |
| $N = 32$ | 0.7967 | 0.0425 | 0.8021 | 0.7967 | 0.0447 | -0.0106 |
| $N = 64$ | 0.7985 | 0.0301 | 0.8018 | 0.7989 | 0.0308 | 0.8231 |
| $p = 256, q = 1, V_{\text{rel}}(\mathbf{\Sigma}) = 0.8$ | | | | | | |
| $N = 8$ | 0.7836 | 0.0781 | 0.8150 | 0.7930 | 0.0994 | 6.6508 |
| $N = 16$ | 0.7926 | 0.0588 | 0.8058 | 0.7944 | 0.0663 | 1.9392 |
| $N = 32$ | 0.7967 | 0.0422 | 0.8033 | 0.7976 | 0.0455 | 1.3138 |
| $N = 64$ | 0.7985 | 0.0299 | 0.8010 | 0.7983 | 0.0303 | -0.5863 |
| $p = 1024, q = 1, V_{\text{rel}}(\mathbf{\Sigma}) = 0.8$ | | | | | | |
| $N = 8$ | 0.7836 | 0.0780 | 0.8140 | 0.7904 | 0.1011 | 4.7652 |
| $N = 16$ | 0.7926 | 0.0587 | 0.8081 | 0.7959 | 0.0665 | 3.5530 |
| $N = 32$ | 0.7967 | 0.0421 | 0.8026 | 0.7967 | 0.0448 | -0.0218 |
| $N = 64$ | 0.7985 | 0.0299 | 0.8012 | 0.7987 | 0.0307 | 0.4338 |

Results for *V*_rel_(**R**) are summarized in Figures 7, S6, S9, and S10 and Tables 3 and S3. Distributions were unimodal within the range (0, 1), except when *p* = 2 and *N* ≤ 8 where an additional peak is usually present near 0. The overall response to varying *p* and *N* is largely similar to that of *V*_rel_(**S**), although the shape of distribution was substantially different for small *N*. As expected from the theoretical expectations noted above, *V*_rel_(**R**) tends to overestimate the population value *V*_rel_(**P**) when the latter is small but tends to underestimate it when *V*_rel_(**P**) = 0.8. The expressions of expectation for the null and arbitrary conditions and variance for the null condition derived above showed almost perfect match with the empirical means and variances (as expected). The heuristic approximation of the variance of *V*_rel_(**R**) for arbitrary conditions using equations 28 and 36–38 yielded larger values than the empirical variances in all cases, except when *V*_rel_(**P**) = 0.8 where slight underestimation was observed. In all cases, the error of this expression decreased to ~0–2% in the SD scale—statistically indistinguishable from random fluctuation with 5000 iterations—for moderately large *N* (typically ≥64–128, occasionally ≥16–32). The asymptotic variance of *V*_rel_(**R**) with equation 39 behaved more idiosyncratically. It yielded similar values to the previous expression, overestimating the true values, under A) the *q*-large λ conditions with *q* = 1 and *V*_rel_(**P**) = 0.1–0.6, B) same with *q* = 2 and *V*_rel_(**P**) = 0.1–0.2 except when *p* = 4, and C) the quadratically decreasing λ conditions with *p* = 4; whereas it yielded smaller values than the true values under a) the *q*-large λ conditions with *q* = 1 and *V*_rel_(**P**) = 0.8, b) same with *q* = 2 and *V*_rel_(**P**) = 0.4, c) same with *q* = 4, d) same with *q* = 2 and *p* = 4, e) the linearly decreasing λ conditions, and f) the quadratically decreasing λ conditions except when *p* = 4. This expression was more accurate than the previous one in the cases a and b, but more inaccurate in other cases. Under some conditions, relative error can be extremely large (10–300% in SD scale with *N* = 256), especially when the smallest population eigenvalue was small in magnitude (<0.1). A practical advantage of this approximation over the previous one may lie in the computational resource required for large *p*. With the present R implementation, evaluation of this expression is faster by a factor of thousands than that of the previous one for a correlation matrix with *p* = 1024 (~1 versus ~1500 CPU seconds on a regular desktop PC), as the amount of computation increases drastically as *p* grows (the latter took only ~5 CPU seconds for *p* = 256).

**Figure 7.**
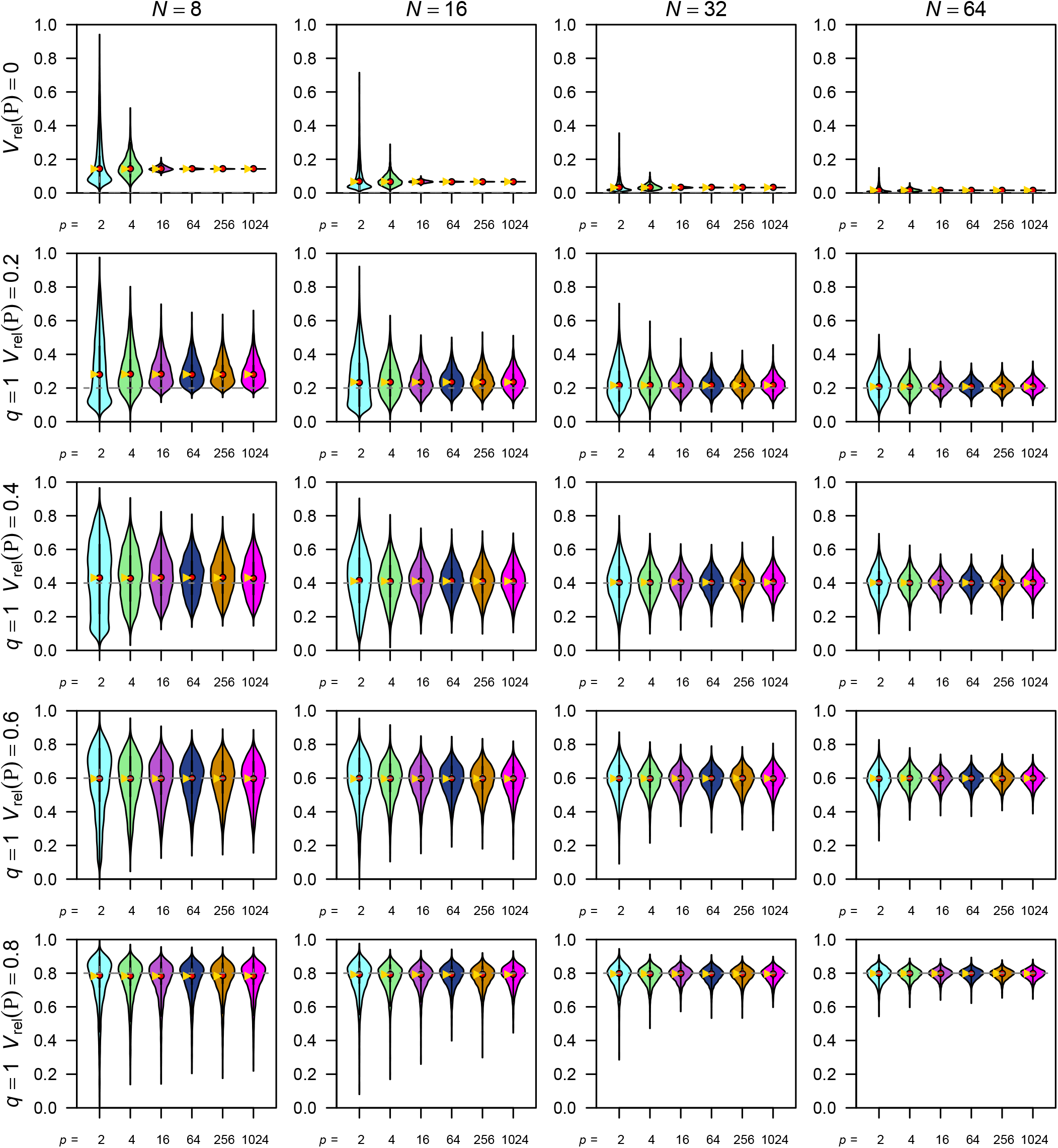
Selected simulation results for the relative eigenvalue variance of correlation matrix *V*_rel_(**R**). Empirical distributions of simulated *V*_rel_(**R**) values are shown as violin plots. Yellow triangles denote expectations (which are exact). Rows of panels correspond to varying population values of *V*_rel_(**P**) (under 1-large λ conditions). Other legends are as in Fig. 5. See Figure S6, S9, and S10 for full results.

**Table 3.** Summary of selected simulation results for relative eigenvalue variance of correlation matrix *V*_rel_(*R*). Theoretical expectation (E[*V*_rel_(**R**)]) and (approximate) standard deviation (SD[*V*_rel_(**R**)]), as well as empirical median, mean, standard deviation (ESD), and bias of mean in standard error unit (*T*) from 5000 simulation runs are shown for selected conditions. The theoretical standard deviations shown in this table are from heuristic approximation (eqs. 28, 36–38) except under the null conditions. See Table 1 for further information and Table S3 for full results.

| | $E[V_{\text{rel}}(\mathbf{R})]$ | $\approx SD[V_{\text{rel}}(\mathbf{R})]$ | Median | Mean | ESD | $T$ |
| --- | --- | --- | --- | --- | --- | --- |
| $p = 2, V_{\text{rel}}(\mathbf{P}) = 0$ | | | | | | |
| $N = 8$ | 0.1429 | 0.1650 | 0.0792 | 0.1429 | 0.1635 | 0.0363 |
| $N = 16$ | 0.0667 | 0.0856 | 0.0336 | 0.0679 | 0.0862 | 1.0235 |
| $N = 32$ | 0.0323 | 0.0435 | 0.0163 | 0.0326 | 0.0427 | 0.6113 |
| $N = 64$ | 0.0159 | 0.0219 | 0.0075 | 0.0156 | 0.0205 | −1.0380 |
| $p = 4, V_{\text{rel}}(\mathbf{P}) = 0$ | | | | | | |
| $N = 8$ | 0.1429 | 0.0673 | 0.1332 | 0.1432 | 0.0673 | 0.3148 |
| $N = 16$ | 0.0667 | 0.0349 | 0.0608 | 0.0661 | 0.0344 | −1.0696 |
| $N = 32$ | 0.0323 | 0.0178 | 0.0292 | 0.0321 | 0.0175 | −0.6906 |
| $N = 64$ | 0.0159 | 0.0090 | 0.0142 | 0.0157 | 0.0087 | −1.0749 |
| $p = 16, V_{\text{rel}}(\mathbf{P}) = 0$ | | | | | | |
| $N = 8$ | 0.1429 | 0.0151 | 0.1411 | 0.1428 | 0.0151 | −0.0917 |
| $N = 16$ | 0.0667 | 0.0078 | 0.0663 | 0.0669 | 0.0080 | 1.7743 |
| $N = 32$ | 0.0323 | 0.0040 | 0.0319 | 0.0322 | 0.0039 | −1.7517 |
| $N = 64$ | 0.0159 | 0.0020 | 0.0158 | 0.0159 | 0.0020 | 0.7127 |
| $p = 64, V_{\text{rel}}(\mathbf{P}) = 0$ | | | | | | |
| $N = 8$ | 0.1429 | 0.0037 | 0.1425 | 0.1428 | 0.0036 | −0.4617 |
| $N = 16$ | 0.0667 | 0.0019 | 0.0666 | 0.0667 | 0.0019 | 0.4407 |

| | $E[V_{\text{rel}}(\mathbf{R})]$ | $\approx \text{SD}[V_{\text{rel}}(\mathbf{R})]$ | Median | Mean | ESD | $T$ |
| --- | --- | --- | --- | --- | --- | --- |
| $N = 32$ | 0.0323 | 0.0010 | 0.0322 | 0.0323 | 0.0010 | -0.0725 |
| $N = 64$ | 0.0159 | 0.0005 | 0.0159 | 0.0159 | 0.0005 | -0.6525 |
| $p = 256, V_{\text{rel}}(\mathbf{P}) = 0$ | | | | | | |
| $N = 8$ | 0.1429 | 0.0009 | 0.1428 | 0.1429 | 0.0009 | 0.0105 |
| $N = 16$ | 0.0667 | 0.0005 | 0.0667 | 0.0667 | 0.0005 | 1.8676 |
| $N = 32$ | 0.0323 | 0.0002 | 0.0323 | 0.0323 | 0.0002 | -0.9701 |
| $N = 64$ | 0.0159 | 0.0001 | 0.0159 | 0.0159 | 0.0001 | 0.3244 |
| $p = 1024, V_{\text{rel}}(\mathbf{P}) = 0$ | | | | | | |
| $N = 8$ | 0.1429 | 0.0002 | 0.1428 | 0.1429 | 0.0002 | -0.9639 |
| $N = 16$ | 0.0667 | 0.0001 | 0.0667 | 0.0667 | 0.0001 | 0.5220 |
| $N = 32$ | 0.0323 | 0.0001 | 0.0323 | 0.0323 | 0.0001 | -1.8393 |
| $N = 64$ | 0.0159 | 0.0000 | 0.0159 | 0.0159 | 0.0000 | -0.0357 |
| $p = 2, q = 1, V_{\text{rel}}(\mathbf{P}) = 0.4$ | | | | | | |
| $N = 8$ | 0.4318 | 0.2495 | 0.4362 | 0.4294 | 0.2516 | -0.6571 |
| $N = 16$ | 0.4111 | 0.1844 | 0.4230 | 0.4147 | 0.1832 | 1.3640 |
| $N = 32$ | 0.4046 | 0.1326 | 0.4056 | 0.4041 | 0.1322 | -0.2806 |
| $N = 64$ | 0.4021 | 0.0944 | 0.4058 | 0.4025 | 0.0954 | 0.3294 |
| $p = 4, q = 1, V_{\text{rel}}(\mathbf{P}) = 0.4$ | | | | | | |
| $N = 8$ | 0.4318 | 0.1950 | 0.4314 | 0.4287 | 0.1750 | -1.2271 |
| $N = 16$ | 0.4111 | 0.1380 | 0.4141 | 0.4093 | 0.1317 | -0.9769 |
| $N = 32$ | 0.4046 | 0.0975 | 0.4054 | 0.4034 | 0.0960 | -0.8743 |
| $N = 64$ | 0.4021 | 0.0688 | 0.4033 | 0.4022 | 0.0687 | 0.1270 |
| $p = 16, q = 1, V_{\text{rel}}(\mathbf{P}) = 0.4$ | | | | | | |
| $N = 8$ | 0.4318 | 0.1621 | 0.4321 | 0.4329 | 0.1391 | 0.5780 |
| $N = 16$ | 0.4111 | 0.1130 | 0.4149 | 0.4120 | 0.1073 | 0.5980 |
| $N = 32$ | 0.4046 | 0.0793 | 0.4084 | 0.4055 | 0.0773 | 0.7996 |
| $N = 64$ | 0.4021 | 0.0559 | 0.4030 | 0.4021 | 0.0552 | -0.0283 |
| $p = 64, q = 1, V_{\text{rel}}(\mathbf{P}) = 0.4$ | | | | | | |
| $N = 8$ | 0.4318 | 0.1542 | 0.4366 | 0.4332 | 0.1304 | 0.7802 |
| $N = 16$ | 0.4111 | 0.1074 | 0.4144 | 0.4129 | 0.1015 | 1.2079 |
| $N = 32$ | 0.4046 | 0.0754 | 0.4052 | 0.4039 | 0.0733 | -0.6457 |
| $N = 64$ | 0.4021 | 0.0531 | 0.4029 | 0.4021 | 0.0535 | 0.0493 |
| $p = 256, q = 1, V_{\text{rel}}(\mathbf{P}) = 0.4$ | | | | | | |
| $N = 8$ | 0.4318 | 0.1522 | 0.4348 | 0.4343 | 0.1278 | 1.3865 |
| $N = 16$ | 0.4111 | 0.1061 | 0.4118 | 0.4091 | 0.0995 | -1.4526 |
| $N = 32$ | 0.4046 | 0.0744 | 0.4078 | 0.4052 | 0.0726 | 0.5664 |
| $N = 64$ | 0.4021 | 0.0524 | 0.4033 | 0.4024 | 0.0522 | 0.4500 |
| $p = 1024, q = 1, V_{\text{rel}}(\mathbf{P}) = 0.4$ | | | | | | |
| $N = 8$ | 0.4318 | | 0.4266 | 0.4286 | 0.1287 | -1.7550 |
| $N = 16$ | 0.4111 | | 0.4127 | 0.4111 | 0.1008 | -0.0454 |
| $N = 32$ | 0.4046 | | 0.4072 | 0.4055 | 0.0726 | 0.8769 |
| $N = 64$ | 0.4021 | | 0.4035 | 0.4024 | 0.0524 | 0.4578 |
| $p = 2, q = 1, V_{\text{rel}}(\mathbf{P}) = 0.8$ | | | | | | |
| $N = 8$ | 0.7831 | 0.1596 | 0.8250 | 0.7853 | 0.1571 | 0.9790 |
| $N = 16$ | 0.7918 | 0.1015 | 0.8141 | 0.7928 | 0.1033 | 0.6253 |
| $N = 32$ | 0.7961 | 0.0674 | 0.8076 | 0.7976 | 0.0680 | 1.5536 |
| $N = 64$ | 0.7981 | 0.0462 | 0.8024 | 0.7976 | 0.0473 | -0.6617 |
| $p = 4, q = 1, V_{\text{rel}}(\mathbf{P}) = 0.8$ | | | | | | |
| $N = 8$ | 0.7831 | 0.1115 | 0.8149 | 0.7840 | 0.1249 | 0.4605 |
| $N = 16$ | 0.7918 | 0.0749 | 0.8059 | 0.7913 | 0.0815 | -0.4682 |
| $N = 32$ | 0.7961 | 0.0516 | 0.8027 | 0.7953 | 0.0543 | -1.0173 |
| $N = 64$ | 0.7981 | 0.0360 | 0.8019 | 0.7987 | 0.0370 | 1.2077 |
| $p = 16, q = 1, V_{\text{rel}}(\mathbf{P}) = 0.8$ | | | | | | |
| $N = 8$ | 0.7831 | 0.0927 | 0.8102 | 0.7835 | 0.1116 | 0.2370 |
| $N = 16$ | 0.7918 | 0.0638 | 0.8045 | 0.7913 | 0.0716 | -0.5326 |
| $N = 32$ | 0.7961 | 0.0445 | 0.8035 | 0.7976 | 0.0457 | 2.4043 |
| $N = 64$ | 0.7981 | 0.0313 | 0.8013 | 0.7984 | 0.0325 | 0.7108 |
| $p = 64, q = 1, V_{\text{rel}}(\mathbf{P}) = 0.8$ | | | | | | |
| $N = 8$ | 0.7831 | 0.0896 | 0.8098 | 0.7841 | 0.1083 | 0.6205 |
| $N = 16$ | 0.7918 | 0.0619 | 0.8046 | 0.7928 | 0.0687 | 0.9579 |
| $N = 32$ | 0.7961 | 0.0432 | 0.8009 | 0.7955 | 0.0452 | -0.9741 |
| $N = 64$ | 0.7981 | 0.0303 | 0.8012 | 0.7983 | 0.0309 | 0.4419 |
| $p = 256, q = 1, V_{\text{rel}}(\mathbf{P}) = 0.8$ | | | | | | |
| $N = 8$ | 0.7831 | 0.0890 | 0.8098 | 0.7857 | 0.1056 | 1.7052 |
| $N = 16$ | 0.7918 | 0.0614 | 0.8031 | 0.7915 | 0.0681 | -0.3408 |
| $N = 32$ | 0.7961 | 0.0429 | 0.8022 | 0.7963 | 0.0460 | 0.3738 |
| $N = 64$ | 0.7981 | 0.0301 | 0.8004 | 0.7977 | 0.0305 | -0.9565 |
| $p = 1024, q = 1, V_{\text{rel}}(\mathbf{P}) = 0.8$ | | | | | | |
| $N = 8$ | 0.7831 | 0.0888 | 0.8088 | 0.7830 | 0.1074 | -0.0831 |
| $N = 16$ | 0.7918 | 0.0613 | 0.8057 | 0.7930 | 0.0683 | 1.2443 |
| $N = 32$ | 0.7961 | 0.0428 | 0.8014 | 0.7954 | 0.0454 | -0.9758 |
| $N = 64$ | 0.7981 | 0.0301 | 0.8006 | 0.7981 | 0.0308 | 0.0710 |

## Discussion

Eigenvalue dispersion indices can be calculated for covariance or correlation matrices in similar ways, but implications are rather different. On the one hand, the relative eigenvalue variance of a sample covariance matrix *V*_rel_(**S**) is a test statistic for sphericity (John, 1972; Sugiura, 1972; Nagao, 1973), and is thus interpreted as a measure of eccentricity of variation, be it due to large variation of a single trait or covariation between traits. Interpretation of the unstandardized eigenvalue variance of a sample covariance matrix *V*(**S**) is less straightforward, but it can potentially be useful in comparing eccentricity between samples when the sensitivity to overall scaling is not of concern, primarily for the presence of an unbiased estimator of the corresponding population value with a known variance (eq. 41). On the other hand, the relative eigenvalue variance of a sample correlation matrix *V*_rel_(**R**) is identical to the average of the squared correlation coefficients across all pairs of traits (Durand & Le Roux, 2017; see above). The average squared correlation is another commonly used index of phenotypic integration (e.g., Cheverud et al., 1983), but its equivalence to *V*_rel_(**R**) seems to have been overlooked, apart from an empirical confirmation by Haber’s (2011) simulations. Obviously, the choice between covariance and correlation should be made according to the scope of individual analyses (Klingenberg, 1996; Hansen & Houle, 2008; Pavlicev et al., 2009; Goswami & Polly, 2010; see also Machado et al., 2019 for an interesting discussion). Usual caveats for the choice between covariance and correlation is also pertinent here (Jolliffe, 2002): covariance between traits have clear interpretability only if all traits are in the same unit. This is despite that *V*_rel_(**S**) is dimensionless and independent of the overall scaling of traits.

Perhaps the most remarkable finding of this study is that the distributions of *V*_rel_(**S**) and *V*_rel_(**R**) do not seem to vary much with the number of variables *p* itself. The above expressions for the (approximate) mean and variance can be calculated for any *p*, and simulation results indicate that their accuracy are not compromised by large *p* (Figs. 5–7 and S4–S10; Tables 1–3 and S1–S3). This finding highlights potential applicability of these measures to high-dimensional phenotypic data. Nevertheless, it should be remembered that, when *p* exceeds the degree of freedom *n, p* − *n* of the sample eigenvalues are 0 and hence the corresponding population eigenvalues are not estimable. In addition, the first sample eigenvector tends to be consistently diverged from the first population eigenvector in high-dimensional settings (Johnstone, 2007; Johnstone & Paul, 2018).

### Applications and limitations

The present analytic results assume simple independent sampling from a multivariate normal population and the Wishart-ness of the cross-product matrix. For some biological datasets, certain modifications would be required. A simple example is data consisting of multiple groups with potentially heterogeneous means, e.g., intraspecific variation calculated from multiple geographic populations or sexes. If uniform **Σ** across groups can be assumed, cross-product matrices from the data centered at the respective group’s sample mean can be summed across groups to obtain a pooled cross-product matrix, which is, by the additivity of Wishart variables, distributed as *W_p_*(**Σ**, *N* − *g*), where *N* is the total sample size and *g* is the number of groups. That is, all above expressions can be applied by simply using the degree of freedom *N* − *g*. A similar correction is required when eigenvalue dispersion indices are applied to partial correlation matrices (Torices & Méndez, 2014; Torices & Muñoz-Pajares, 2015). The distribution of sample partial correlation coefficients in *p*_1_ variables conditionalized on *p*_2_ other variables based on *N* observations is the same as that of ordinary correlation coefficients based on *N* − *p*_2_ observations with the same corresponding parameters (e.g., Anderson, 2003: p. 143), so the appropriate degree of freedom is *n* − *p*_2_. Both of these procedures are essentially to examine the covariance/correlation matrix of residuals after conditionalizing on covariates.

Present analytical results may not be applicable to those empirical covariance or correlation matrices that are not based on a Wishart matrix. Primary examples are the empirical **G** matrices estimated from variance components in MANOVA designs or obtained as likelihood-based estimators in mixed models (e.g., Lynch & Walsh, 1998; Meyer & Kirkpatrick, 2005). Mean-standardization, a method recommended for analyzing **G** matrices (Houle, 1992; Hereford et al., 2004; Hansen & Houle, 2008), can also violate the distributional assumption if sample means are used in the standardization. If eigenvalue dispersion indices are to be used with any of these methods, their sampling properties need to be critically assessed (see also Sztepanacz & Blows, 2017).

The assumption of multivariate normality may be intrinsically inappropriate for some types of data, including meristic (count) data, compositional or proportional data, angles, and directional data. Application of eigenvalue dispersion indices (or indeed covariance/correlation itself) to such data types would require special treatments, which are beyond the scope of this study. Needless to say, the appropriateness of multivariate normality should be critically assessed in every empirical dataset when the present analytic results are to be applied, even if the data type is conformable with normality. Robustness of the above results against non-normality may deserve some investigations.

### Shape variables

The application to traditional morphometric datasets, in which all variables are typically measured in the same unit, is rather straightforward, as covariance/correlation in such variables has full interpretability in the Euclidean trait space. Quite often, component(s) of little interest, e.g., size, are removed by transforming raw data, inducing covariation in resultant variables that needs to be taken into account in hypothesis tests. The most typical transformation is the division by an isometric or allometric size variable (Jolicoeur, 1963; Mosimann, 1970; Mosimann & James, 1979; Darroch & Mosimann, 1985; Klingenberg, 1996, 2016), which can conveniently be done by orthogonal projection in the space of log-transformed variables. The projection of objects onto the hyperplane orthogonal to a subspace, say, the column space of **H** (*p* × *k* full-column-rank matrix; for the isometric size vector, **H** = *p*^1/2^**1**_*p*_), can be done by right-multiplying the data by the projection matrix (e.g., Burnaby, 1966):

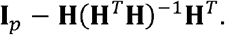

Therefore, the covariance matrix in the resultant space can be obtained from that in the original space **Σ** as

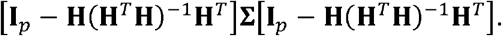

Under the null condition (**Σ** = σ^2^**I**_*p*_) specifically, this becomes

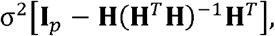

because the projection matrix is symmetric and idempotent. This transformation renders *k* eigenvalues to be 0 by construction. When the focus is on covariance rather than correlation, these null eigenvalues can optionally be dropped from calculation of eigenvalue mean and dispersion, so that the resultant dispersion index quantifies eccentricity of variation in the subspace of interest. Theories derived above can be applied with minimal modifications, although the asymptotic variance of *V*_rel_(**R**) (eq. 39) may not work well due to singularity. These discussions assume that independence between the raw variables can at least hypothetically be conceived, e.g., when measurements are taken from non-overlapping parts of an organism. If measurements are taken from overlapping parts of an organism, then the dependence between variables due to the geometric configuration needs to be taken into consideration on a case-by-case basis (Mitteroecker et al., 2012).

Application to landmark-based geometric morphometric data is more complicated, primarily because the shape space of Procrustes-aligned landmark configurations is (typically a restricted region of) the surface of a hyper(hemi)sphere (e.g., Slice, 2001). In practice, however, empirical analyses are usually conducted on a Euclidean tangent space instead of the shape space itself, assuming that the former gives a satisfactory metric approximation of the latter (e.g., Rohlf, 1999; Marcus et al., 2000; Klingenberg, 2020). It will in principle be possible to obtain an approximate population covariance matrix of landmark coordinates in this tangent space from a hypothetical covariance matrix of raw landmark coordinates before alignment, by using the orthogonal projection method mentioned above with such an **H** whose columns represent the non-shape components. Such a set of vectors can be obtained either as a basis of the complement of the tangent space (see Rohlf & Bookstein, 2003) or directly from the consensus configuration (Klingenberg, 2020). The stereographic projection might potentially be preferred over the orthogonal projection in projecting aligned empirical configurations in the shape space to the tangent space—not to be confused with the projection from the raw space to the tangent space—for purposes of analysing eccentricity of variation. This is because the stereographic projection tends to approximately preserve multivariate normality of the raw coordinates into the resultant tangent space, provided that the variation in the raw coordinates is sufficiently small and that the mean configuration is taken as the point of tangency (Rohlf, 1999). It should be noted that Procrustes superimposition changes perceived patterns of variation in landmark coordinates, often rather drastically (Rohlf & Slice, 1990; Walker, 2000). Such phenomena are probably to be seen as properties of shape variables, rather than necessarily nuisance artifacts (Klingenberg, 2021). Whether these can be of concern or not would depend on the scope of individual analyses (see also Machado et al., 2019).

### Phylogenetic data

So far data were assumed to be i.i.d. multivariate normal variables. Important applications in evolutionary biology involve non-i.i.d. observations, most notably phylogenetically structured data in which *N* observations (typically species) have covariance due to shared evolutionary histories. Trait covariation at the interspecific level may have interpretations under certain evolutionary models (Felsenstein, 1988; Hansen & Martins, 1996; Revell & Harmon, 2008; Uyeda & Harmon, 2014; Caetano & Harmon, 2019). Under the assumption that trait evolution along phylogeny can be described by (potentially a mixture of) linear invariant Gaussian models, such as the Brownian motion (BM), accelerating–decelerating (ACDC; or early burst), and Ornstein–Uhlenbeck (OU) processes, the joint distribution of the observations is known to be multivariate normal (Hansen & Martins, 1996; Manceau et al., 2017; Mitov et al., 2020). A brief overview is given below for potential applications of the present analytic results to phylogenetically structured data.

For BM and its modifications, including BM with a trend, Pagel’s λ, and ACDC models, the covariance matrix of the *N* × *p* dimensional data **X** can be factorized into the intertrait and interspecific components in the form of Kronecker product: **Σ** ⊗ **Ψ**, where **Ψ** is the *N* × *N* interspecific covariance matrix specified by the underlying phylogeny and parameter(s) specific to the evolutionary model (see Hansen & Martins, 1996; Freckleton et al., 2002; Blomberg et al., 2003; Clavel et al., 2015; Mitov et al., 2020). In this case the data can conveniently be considered as a matrix-variate normal variable (see Gupta & Nagar, 1999): **X** ~ *N_N,p_*(**M**, **Σ** ⊗ **Ψ**), where **M** is a *N* × *p* matrix of means. If **Ψ** is known a priori—that is, we have an accurate phylogenetic hypothesis and parameters—the change of variables **Y** = **Ψ**^−1/2^**X** leads to **Y** ~ *N_n,p_*(**Ψ**^-1/2^**M**, **Σ** ⊗ **I**_*N*_), thereby essentially avoiding the complication of dependence between observations. This procedure is widely recognized as the (phylogenetic) generalized least squares (GLS; e.g., Grafen, 1989; Martins & Hansen, 1997; Rohlf, 2001; Symonds & Blomberg, 2014). If we know the population mean **M** in addition, then the cross-product matrix centered at it,

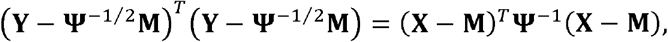

is distributed as *W_p_*(**Σ**, *N*). If we don’t exactly know **M** yet still assume **M** = **1**_*N*_**μ**^*T*^ with the unknown but uniform *p* × 1 mean vector **μ**, then the GLS estimate of the mean 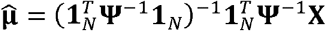 (e.g., Martins & Hansen, 1997) can be used to obtain a sample-mean-centered cross-product matrix

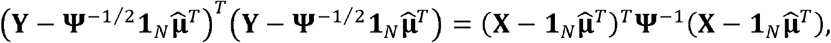

which can be shown to be distributed as *W_p_*(**Σ**, *N* − 1). If there are multiple blocks of species with different means (regimes), then cross-product matrices calculated separately for each of these can be summed to obtain a Wishart matrix with a modified degree of freedom as mentioned above, although it would need to be asked first whether those regimes share the same **Σ** (Revell & Collar, 2009; Caetano & Harmon, 2019). The present analytic results can directly be applied to these Wishart matrices. Estimation of **Σ** based on this transformation has previously been devised (Revell & Harmon, 2008; see also Huelsenbeck & Rannala, 2003, Revell & Harrison, 2008; Adams & Felice, 2014), and has been shown to have superior accuracy in estimating eigenvalues over estimation ignoring phylogenetic structures under model conditions (Revell, 2009). Variants of this method have already been applied to analyze eccentricity of interspecific covariation (Haber, 2016; Watanabe, 2018). In practice, however, **Ψ** is virtually never known exactly because phylogeny and parameters of evolutionary models are generally estimated with error, so empirical cross-product matrices may not be strictly Wishart. This source of error is inherent to any phylogenetic comparative analysis. Unlike the GLS estimate of the mean, which remains unbiased even when **Ψ** is misspecified, the GLS estimate of trait (co)variance is in general biased in this case (see Rohlf, 2006). Although there are certain ways to incorporate phylogenetic uncertainty into statistical inferences (e.g., Huelsenbeck & Rannala, 2003; Garamszegi & Mundry, 2014; Nakagawa & de Villemereuil, 2019), potential consequences of the uncertainty over the distributions of derived statistics require further investigation (see also Revell et al., 2018). Nevertheless, the GLS estimation with slightly inaccurate **Ψ** is supposed to yield a better estimate of trait (co)variance than the estimation ignoring phylogenetic covariation altogether (Rohlf, 2006). It should be noted that uniform scaling of **Ψ** translates to the reciprocal scaling of the cross-product matrix; *V*(**S**) is sensitive to this scaling, whereas *V*_rel_(**S**) and *V*_rel_(**R**) are not. Therefore, specifically under the BM model, the phylogenetic uncertainty would be the only major concern for the latter two indices.

Unfortunately, the GLS estimation of trait covariance does not seem feasible for multivariate OU models, where the joint covariance matrix cannot in general be factorized into intertrait and interspecific components (Bartoszek et al., 2012; Mitov et al., 2020). This is notably except when the selection strength matrix is spherical and the tree is ultrametric, in which case a factorization of the form **Σ** ⊗ **Ψ** is possible (the scalar OU model; Bastide et al., 2018) and hence the GLS cross-product matrix can in principle be calculated, assuming that the relevant parameters are known. Otherwise, the random drift/diffusion matrix of the OU model estimated in one or other criteria can potentially be analyzed, although little is known about its sampling properties under various implementations, other than that accurate estimation is notoriously difficult (e.g., Ho & Ané, 2014; Clavel et al., 2015). Further studies are required on technical aspects of quantifying trait covariation in phylogenetically structured data under such complex models, as well as its biological implications (e.g., Adams & Collyer, 2018, 2019b; Mitov et al., 2019, 2020; Clavel et al., 2019; Clavel & Morlon, 2020).

### Concluding remarks

Eigenvalue dispersion indices of covariance or correlation matrices are commonly used as measures of trait covariation, but their statistical implications have not been well appreciated by biologists, against which criticism has reasonably been directed (Hansen & Houle, 2008; Blows & McGuigan, 2015; Hansen et al., 2019). As discussed above, *V*_rel_(**S**) and *V*_rel_(**R**) have clear statistical justifications as test statistics for sphericity and no correlation, respectively. However, sample eigenvalue dispersion indices are biased estimators of the corresponding population values. This paper derived (or restated) exact and approximate expressions for the expectation and variance of *V*(**S**), *V*_rel_(**S**), and *V*_rel_(**R**) under the respective null and arbitrary conditions, with which empirical values can be compared. All null moments are exact, as well as both moments of *V*_rel_(**S**) and the expectation of *V*_rel_(**R**) under arbitrary conditions. Moments of *V*_rel_(**S**) under arbitrary conditions are approximations based on the delta method; the approximate expectation was shown to work reasonably well with a moderate sample size (*N* ≥ 16– 32), whereas the approximate variance requires a larger sample size to be reliable (e.g., *N* ≥ 64, depending on other conditions). Two approximate expressions were given for the variance of *V*_rel_(**R**) under arbitrary conditions. The one with equations 28 and 36–38 works reasonably well with a moderate to large sample size (*N* ≥ 16– 128) but requires nontrivial computational time for a large matrix (e.g., *p* = 1024), whereas the one with equation 39 tends to be more inaccurate in some conditions but can be evaluated almost instantly. Under such conditions where these expressions work, they can be used for (approximate) statistical inferences and hypothesis tests for the magnitude of integration, as well as for determination of appropriate sample sizes in empirical analyses, essentially replacing qualitative thresholds proposed earlier (e.g., Haber, 2011; Jung et al., 2020).

There are several conceivable ways for statistical inferences and hypothesis testing for eigenvalue dispersion indices. When sample size is so large that distributions of the indices are virtually symmetric (*N* ≥ 16– 128, depending on other conditions), the moments derived above may potentially be used to construct approximate confidence intervals. If multivariate normality (or any other explicit distribution) can be assumed, then it is straightforward to obtain empirical distributions under appropriate conditions with Monte Carlo simulations. Critical points of the null distributions and empirical power at α = 0.05 and 0.01 based on the present simulations are presented in Table S1–S3 as a quick guide for sampling design. Several limiting and approximate distributions have been proposed for related statistics (e.g., John, 1972; Nagao, 1973; Ledoit & Wolf, 2002; Schott, 2005), which could be used for simple null hypothesis testing with large *N*. Resampling-based tests are another potential avenue of development. Applicability and performance of these alternative methods would deserve further investigations.

## Supporting information

Supplemental codes

## Acknowledgements

The author would like to thank Carmelo Fruciano and Christian P. Klingenberg for encouragements and constructive comments in an early stage of the study. This work was partly supported by the Newton International Fellowships by the Royal Society (NIF\R1\180520) and the Overseas Research Fellowships by the Japan Society for the Promotion of Science (202160529). The author declares no conflict of interest.

## Appendix A

In this part, relationships between eigenvalue dispersion indices and individual eigenvalues are derived under certain restrictive conditions, in order to facilitate interpretation and to clarify algorithms used in simulations. For simplicity, it is assumed 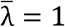 in the following discussions; general cases easily follow by scaling.

Let us first consider the simple conditions where the first *q* (<*p*) population eigenvalues are equally large and the rest *p* − *q* eigenvalues are equally small: λ_1_ = ··· = λ_*q*_ ≥ λ_*q*+1_ = ··· = λ_*p*_ (“*q*-large λ conditions” in simulations). By noting ∑λ_*i*_ = *qλ*_1_ + (*p* − *q*)λ_*p*_ = *p*, it is seen that

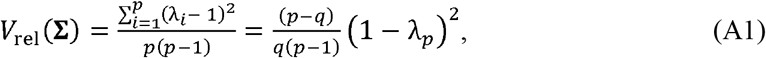

and hence

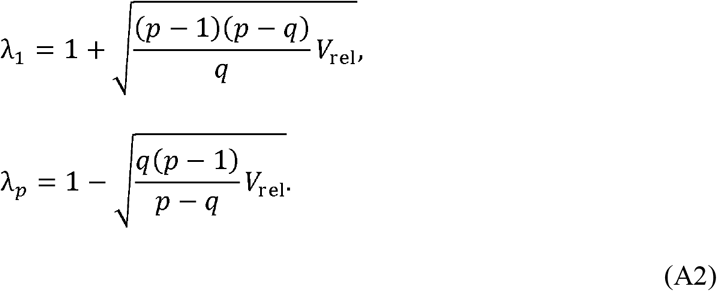

By noting the constraint 0 ≤ λ_*p*_ ≤ 1, an upper limit of *V*_rel_ can be seen from equation A1:

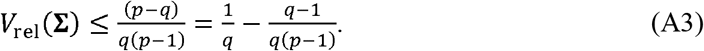

It is then obvious that, under these constraints, a value of *V*_rel_ greater than 0.5 cannot happen when *q* > 1; that is, such a large value implies the dominance of a single component of variance. The same arguments equally apply to correlation matrices.

When *q* = 1 for the correlation matrix, *V*_rel_(**P**) completely specifies the magnitude of correlation in every pair of variables. This point can be seen from the definition of eigendecomposition:

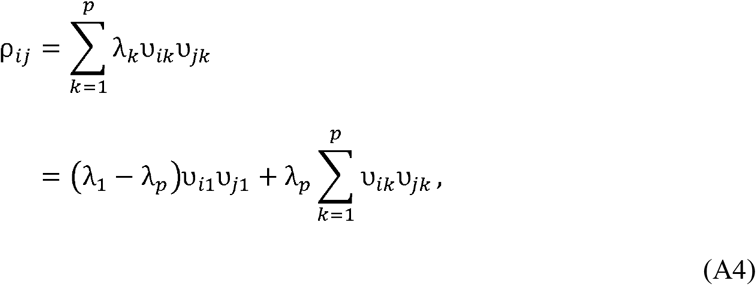

where υ_*ij*_ is the (*i, j*)-th element of the eigenvector matrix. By remembering 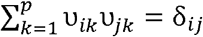, the Kronecker delta, and noting equation A2 with *q* = 1,

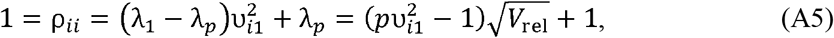

therefore 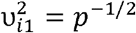 for any *i* (that is, the coefficients of the first eigenvector are equal in magnitude). Finally, we have

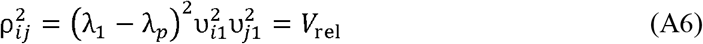

for any combination of *i* and *j* (*i* ≠ *j*); the magnitude of correlation is identical across all pairs. Taken differently, λ_2_ = ··· = λ_*p*_ = |ρ|. These relationships have previously been noted by Anderson (1963) and Pavlicev et al. (2009).

The population eigenvalues of the linearly and quadratically decreasing λ conditions used in simulations are defined as λ_*i*_ = (*p* − *i* + 1)λ_*p*_ and λ_*i*_ = (*p* − *i* + 1)^2^λ_*p*_ (*i* = 1, 2,…, *p*) for linearly and quadratically decreasing conditions, respectively. Under the assumption of a constant average eigenvalue, simple algebra yields the actual values of λ_*p*_ and *V*_rel_(**Σ**) as functions of *p*. The latter equals 1/3(*p* + 1) and (8*p* + 11)/5(*p* + 1)(2*p* + 1) for the linearly and quadratically decreasing λ conditions, respectively.

## Appendix B

In this part, the first two moments of *V*_rel_(**S**) and *V*_rel_(**S**) under the arbitrary **Σ** are derived, assuming multivariate normality. Derivation of the moments of the latter requires evaluation of moments of the ratio 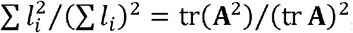, which are not guaranteed to coincide with the ratios of moments except under the null hypothesis. Here we utilize the approximation based on the delta method (eq. 32). In turn, we need E[tr(**A**^2^)], E[(tr **A**)^2^], Var[tr(**A**^2^)], Var[(tr **A**)^2^], and Cov[tr(**A**^2^), (tr **A**)^2^].

We will follow Srivastava & Yanagihara’s (2010) approach to obtain these moments. As in the text, let the *n* × *p* matrix **Z** be (**z**_1_, **z**_2_,…, **z**_*n*_)^*T*^, where **z**_*i*_~*N_p_*(**0**_*p*_, **Σ**) for *i* = 1, 2,…, *n*. Consider the cross-product matrix

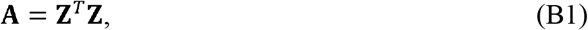

such that **A** ~ *W_p_*(**Σ**, *n*). Let the spectral decomposition of **Σ**:

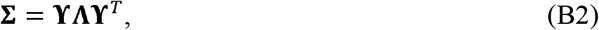

with the orthogonal matrix of eigenvectors **ϒ** and the diagonal matrix of eigenvalues **Λ**. Let the *n* × *p* matrix **J** be (**j**_1_, **j**_2_,…, **j**_*n*_)^*T*^, where are **j**_*i*_ i.i.d. *N_p_*(**0, I**_*p*_), such that **Z** = **JΣ**^1/2^ with **Σ**^1/2^ = **ϒΛ**^1/2^**ϒ**^*T*^. Then, it is possible to write

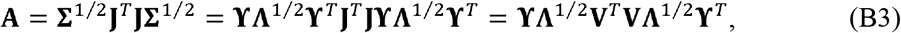

where **V** = **Jϒ** = (**v**_1_, **v**_2_, …, **v**_*p*_) with **v**_*i*_ being i.i.d. *N_n_*(**0, I**_*n*_). Furthermore, let 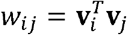, such that *w_ii_* are i.i.d. chi-square variables with *n* degrees of freedom 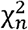. Obviously *w_ij_* = *w_ji_*. Note that

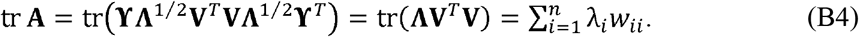

From well-known results on normal and chi-square moments, we have the following:

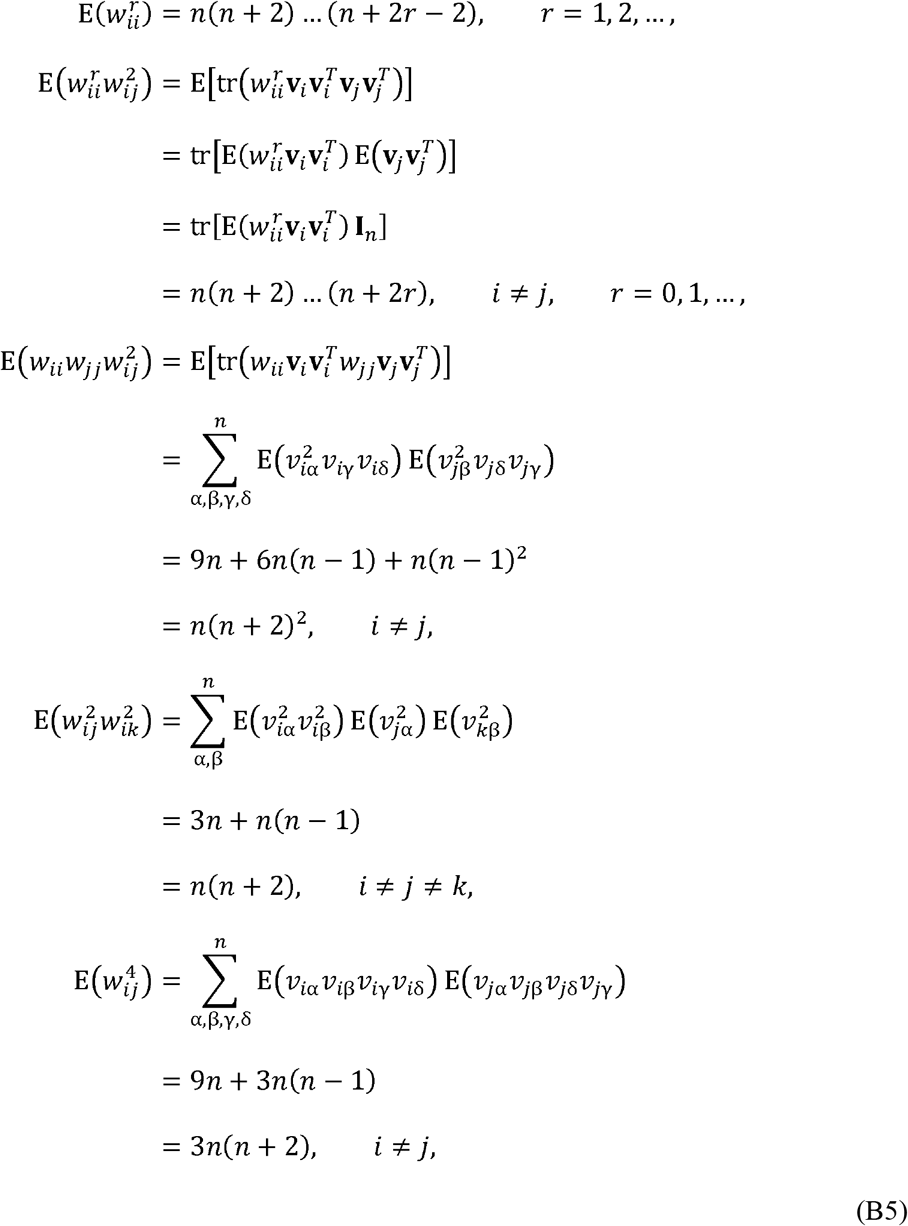

where intervening equations can be confirmed by direct enumeration of the nonzero moments.

From the above expectations, one can evaluate the desired moments as follows:

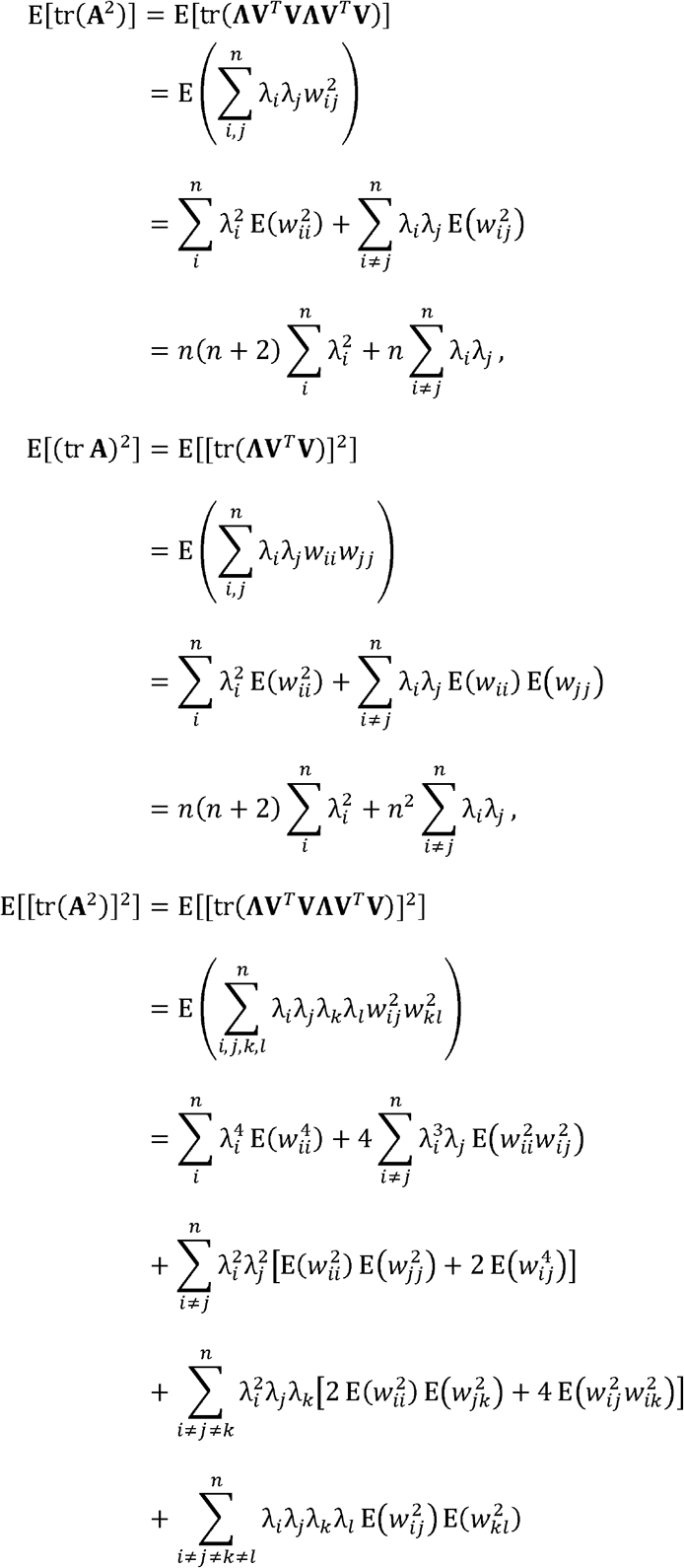

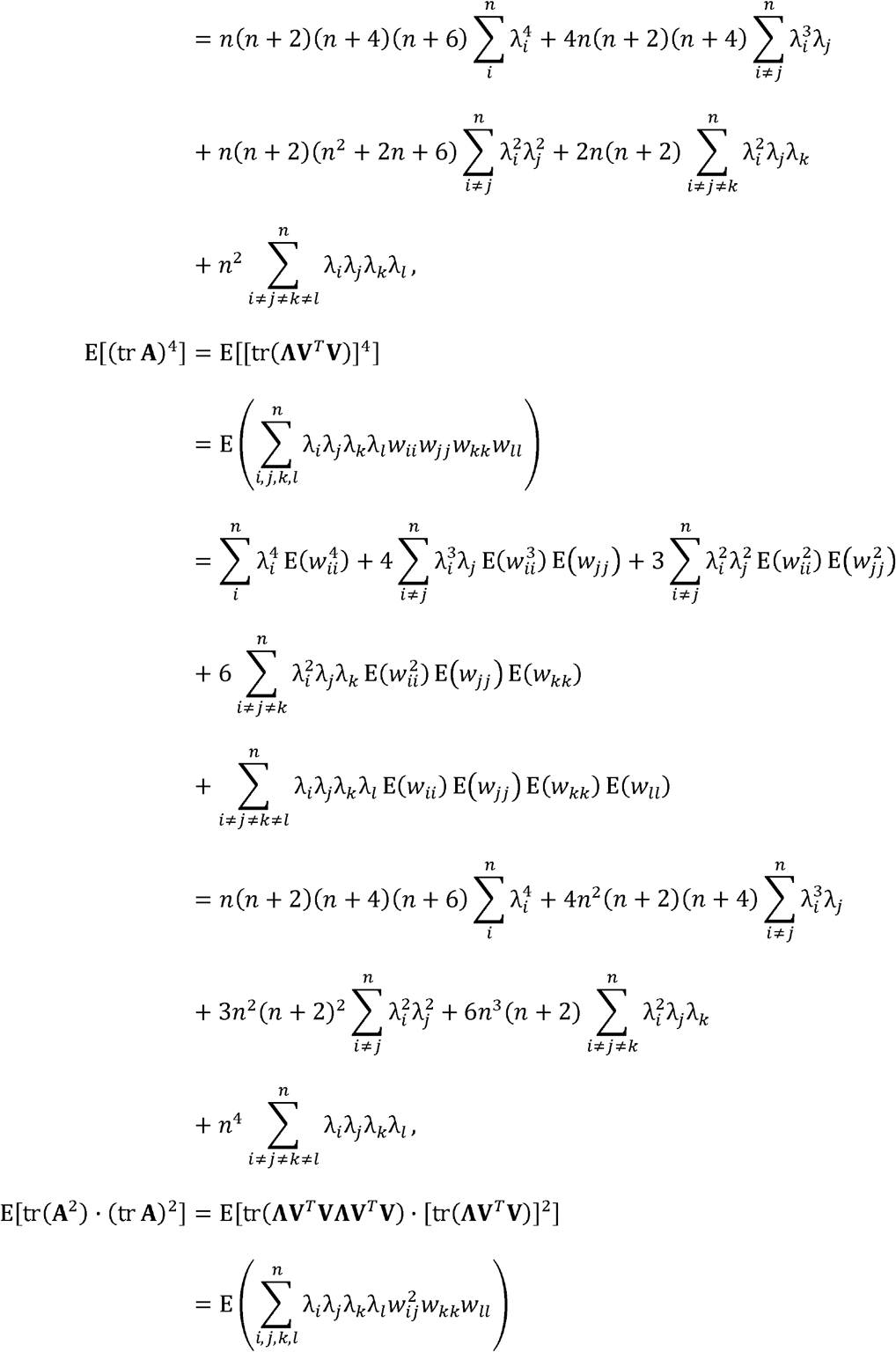

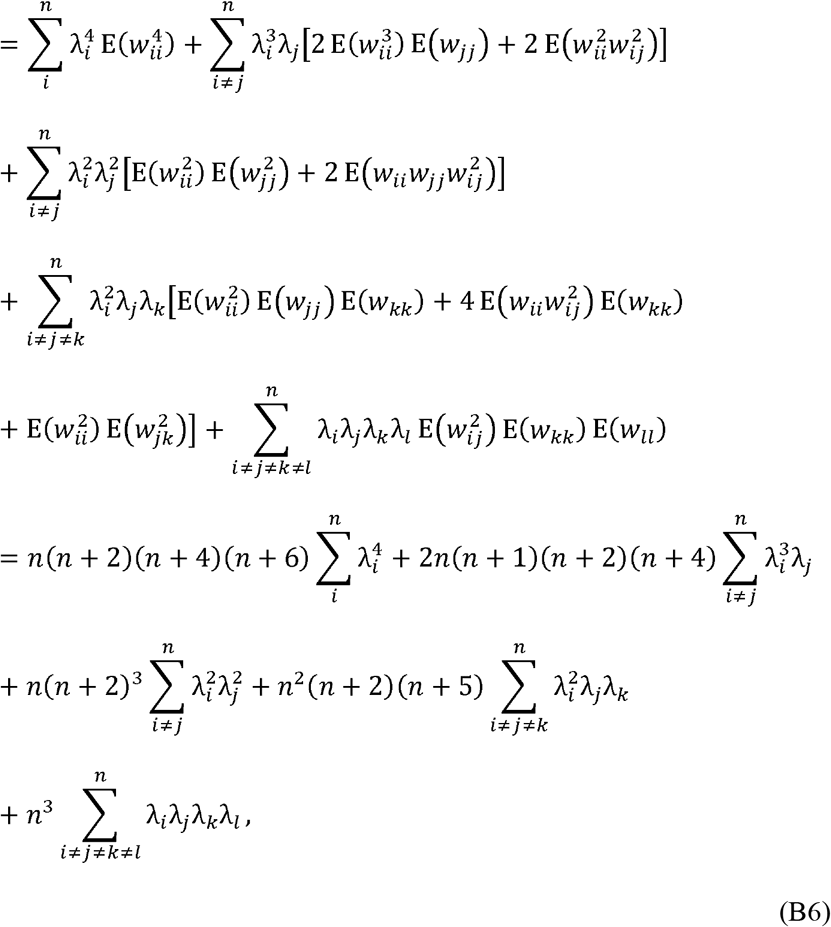

where notations of the form *i* ≠ *j* ≠ *k* ≠ *l* represent inequality of every pairwise combination of the subscripts concerned.

Although equations B6 can be evaluated for any **Σ**, calculating the product of all possible combinations of eigenvalues is rather cumbersome when *p* is large. For this reason, it would be preferable to simplify these expressions by noting

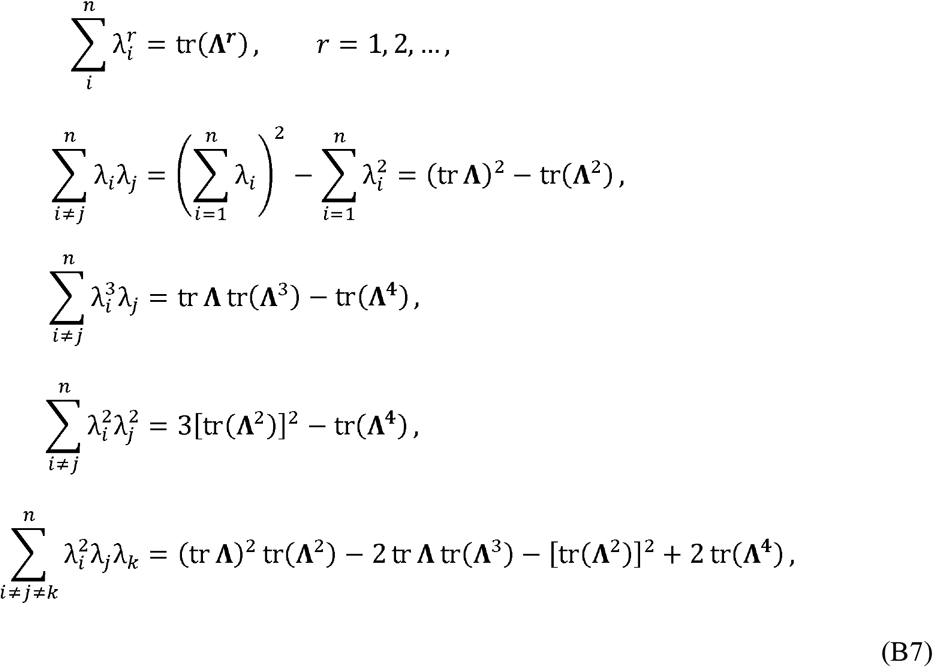

Then, equations B6 can be written as follows:

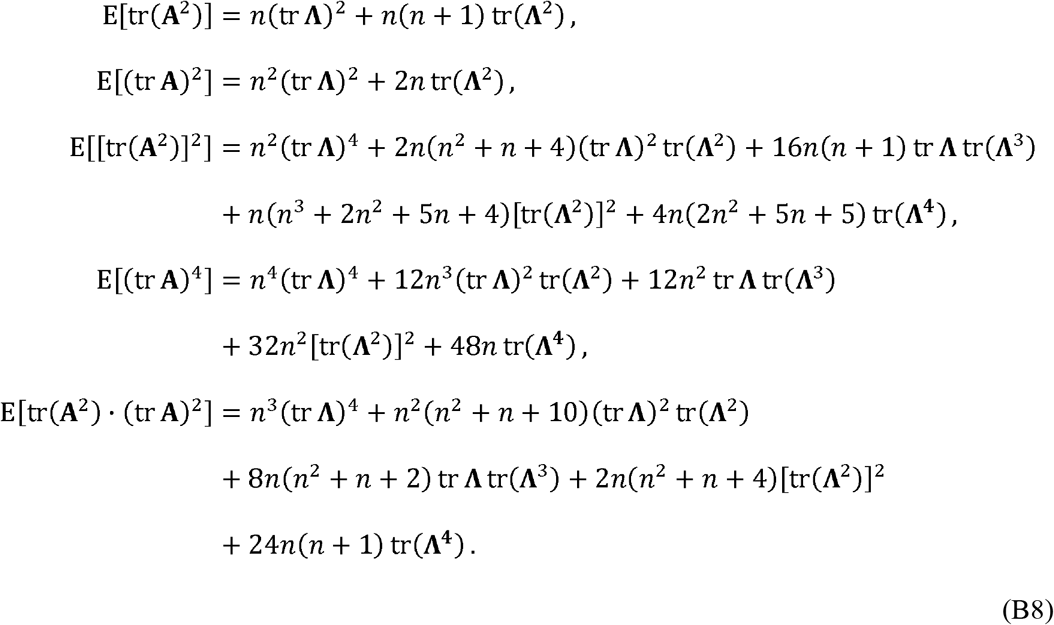

Finally,

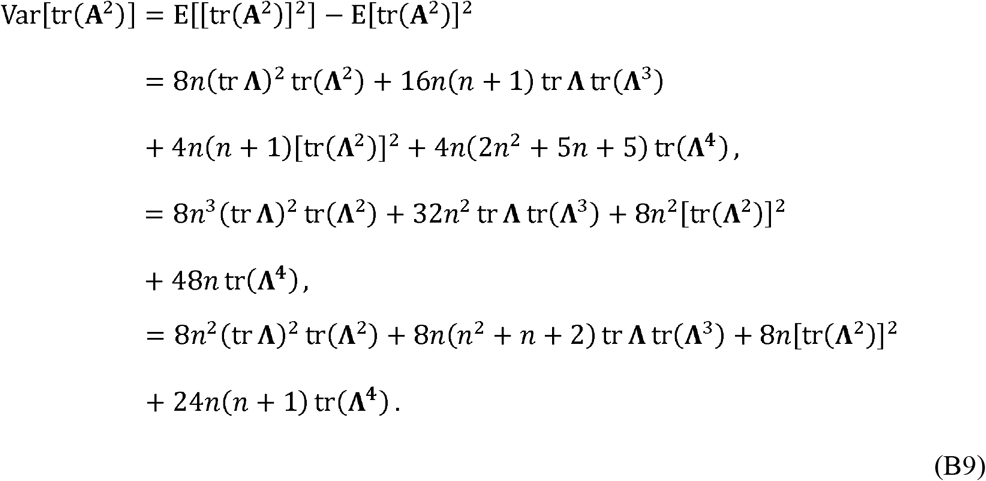

Inserting equations B8 and B9 into equations 12, 19, and 32 yields the desired results.

Identical results can be derived from del Waal & Nel’s (1973) results on the expectations of elementary symmetric functions of eigenvalues and their products for a Wishart matrix. However, these results appear to have been proved only under the condition *n* > *p* − 1 (see also Constantine, 1963; Muirhead, 1982: chapter 7). The above derivation is valid for any combination of *p* and *n*.

## Appendix C

This part demonstrates that 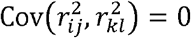 for (*i, j*) ≠ (*k, l*) under the condition **P** = **I**_*p*_, as cursorily mentioned by Schott (2005). Under this condition, a sample covariance can be written as 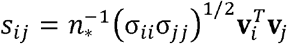, with **v**_*i*_ and **v**_*j*_ being i.i.d. *N_n_*(**0, I**_*n*_). Therefore, a sample correlation coefficient can be written as 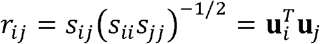, where 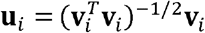 are uniformly distributed on the surface of the unit hypersphere in the *n*-dimensional space. By noting 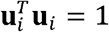, it is possible to see 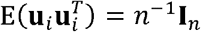 for any *i*, because the elements of **u**_*i*_ are symmetric and uncorrelated with one another (a formal demonstration probably requires introduction of the density function; see Anderson, 2003: p. 49). With these preliminaries, it is easily seen, for *i* ≠ *j* ≠ *k*,

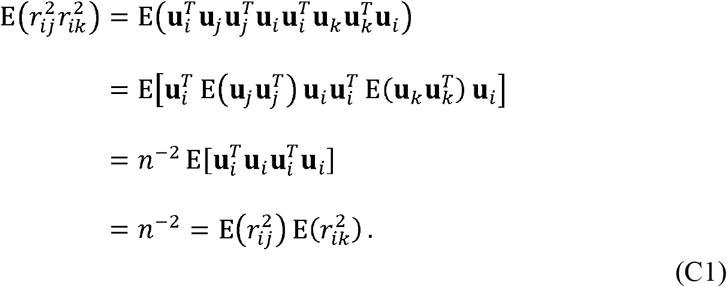

The second equation is valid because **u**_*i*_, **u**_*j*_, and **u**_*k*_ are stochastically independent from one another. Therefore, 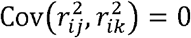 for partly overlapping subscripts. Similarly, 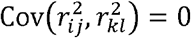 for non-overlapping subscripts, although this could also be seen as a direct consequence of the independence between *r_ij_* and *r_kl_* in this case.

## Appendix D

This part outlines assumptions under the heuristic approximation of 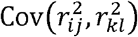 (eq. 37). Pan & Frank (2004) derived approximate moments for the product of two correlation coefficients with overlapping subscripts (i.e., *r_ij_* and *r_ik_*). Equivalent results for general combinations of subscripts can be obtained by following the derivation up to their equation 3.8. The relevant result is:

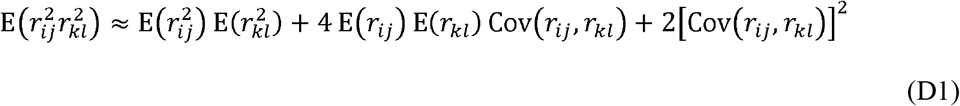

Derivation of this result only requires the assumption that the third and fourth moments of correlation coefficients approximately behaves like normal moments; namely, E([*r*_α_ − E(*r*_α_)][*r*_β_ − E(*r*_β_)][*r*_γ_ − E(*r*_γ_)]) ≈ 0 and E([*r*_α_ − E(*r*_α_)][*r*_β_ − E(*r*_β_)][*r*_γ_ − E(*r*_γ_)][*r*_δ_ − E(*r*_δ_)]) ≈ Cov(*r*_α_, *r*_β_) Cov(*r*_γ_, *r*_δ_) + Cov(*r*_α_, *r*_γ_) Cov(*r*_β_, *r*_δ_) + Cov(*r*_α_, *r*_δ_) Cov(*r*_β_, *r*_γ_), where α, β, γ, and δ are arbitrary pairs of subscripts. Of course these do not strictly hold, except asymptotically under *n* → ∞ (e.g., Olkin & Siotani, 1976; Konishi, 1979). Equation 37 is immediate from equation D1.

## Appendix E

In this part, an asymptotic expression for the variance of *V*_rel_(**R**) is derived for arbitrary non-null conditions with *p* > 2. Konishi (1979) gave an asymptotic theory for the distribution of an arbitrary function of eigenvalues of a sample correlation matrix *f*(*l*_1_, …, *l_p_*) under multivariate normality. In particular, when *n* → ∞, 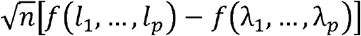 was shown to be normally distributed with mean 0 and variance

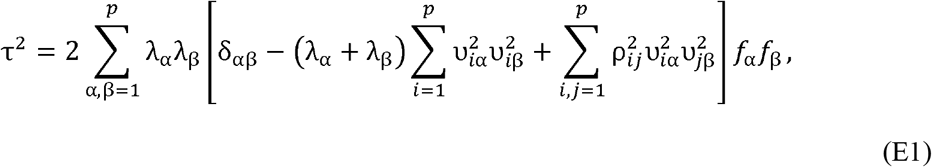

where the summations are over all combinations of subscripts, δ_αβ_ is the Kronecker delta, υ_*i*α_ is the (*i*, α)-th element of the population eigenvector matrix **ϒ**, and *f*_α_ = ∂*f* / ∂*l*_α_|_(*l*_1_,…,*l_p_*)=(λ_1_,…,λ_*p*_)_ the partial derivative of *f* with respect to *l*_α_ evaluated at (*l*_1_, …, *l_p_*) = (λ_1_,…, λ_*p*_). Note that Konishi’s (1979; corollary 2.2) original notation also concerned potential multiplicity of population eigenvalues, which is ignored here for simplicity; the population eigenvectors corresponding to multiplicated eigenvalues can in practice be chosen arbitrarily as a suite of orthogonal vectors in the appropriate subspace, as is done in numerical determination of eigenvectors. The derivative of *V*_rel_(**R**) is simply

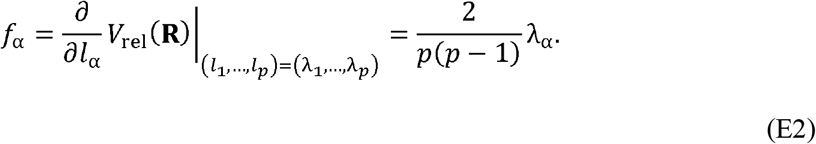

Inserting equation E2 into equation E1, we obtain τ^2^/*n* as an asymptotic expression of the variance of *V*_rel_(**R**) (eq. 39). It should be noted that this expression will be inaccurate around **P** = **Λ** = **I**_*p*_. This is because, under this condition, we can arbitrarily put **ϒ** = **I**_*p*_ to obtain τ^2^ = 0 from equation E1, which is clearly untrue for finite *n*.

An empirically equivalent result can be obtained from the alternative expression of *V*_rel_(**R**) as average squared correlation coefficients (eq. 11), from a similar theory for functions of a sample correlation matrix by Konishi (1979: theorem 6.2). However, that alternative expression does not seem to bear much practical advantage, for it typically takes substantially more computational time to evaluate as *p* grows.

