## Supplemental tables and figures for "Statistics of eigenvalue dispersion indices: quantifying the magnitude of phenotypic integration"

**Table S1.** Summary of simulation results for $V(\mathbf{S})$. Theoretical expectation (E[$V(\mathbf{S})$]) and standard deviation (SD[$V(\mathbf{S})$]), as well as empirical median, mean, standard deviation (ESD), bias in standard error unit ($T = \sqrt{5000}\{\text{Mean} - \text{E}[V(\mathbf{S})]\}$ / ESD, which should roughly follow $t_{4999}$ if the expectation is exact), and critical points or power (for null and non-null conditions, respectively) at α = 0.05 and 0.01 (CP/Pow. 5% and 1%; the latter obtained from the former with same $N$ and $p$) from 5000 simulation runs are shown.

| | E[$V(\mathbf{S})$] | SD[$V(\mathbf{S})$] | Median | Mean | ESD | $T$ | CP 5% | CP 1% |
|---|---|---|---|---|---|---|---|---|
| $p$ = 2, $V(\mathbf{\Sigma})$ = 0 | | | | | | | | |
| $N$ = 4 | 0.6667 | 1.0184 | 0.3299 | 0.6857 | 1.0338 | 1.2996 | 2.5760 | 5.0374 |
| $N$ = 8 | 0.2857 | 0.3582 | 0.1731 | 0.2876 | 0.3464 | 0.3845 | 0.9689 | 1.6625 |
| $N$ = 16 | 0.1333 | 0.1501 | 0.0876 | 0.1330 | 0.1466 | −0.1703 | 0.4143 | 0.6728 |
| $N$ = 32 | 0.0645 | 0.0686 | 0.0438 | 0.0646 | 0.0689 | 0.0489 | 0.1959 | 0.3342 |
| $N$ = 64 | 0.0317 | 0.0327 | 0.0221 | 0.0315 | 0.0315 | −0.6336 | 0.0947 | 0.1414 |
| $N$ = 128 | 0.0157 | 0.0160 | 0.0109 | 0.0155 | 0.0152 | −1.3106 | 0.0463 | 0.0692 |
| $N$ = 256 | 0.0078 | 0.0079 | 0.0054 | 0.0079 | 0.0079 | 0.5012 | 0.0234 | 0.0367 |
| $p$ = 4, $V(\mathbf{\Sigma})$ = 0 | | | | | | | | |
| $N$ = 4 | 0.5000 | 0.4811 | 0.3536 | 0.5028 | 0.4889 | 0.4064 | 1.4161 | 2.3918 |
| $N$ = 8 | 0.2143 | 0.1551 | 0.1734 | 0.2131 | 0.1534 | −0.5437 | 0.5055 | 0.7688 |
| $N$ = 16 | 0.1000 | 0.0602 | 0.0861 | 0.0996 | 0.0603 | −0.4826 | 0.2148 | 0.3091 |
| $N$ = 32 | 0.0484 | 0.0261 | 0.0429 | 0.0481 | 0.0261 | −0.8676 | 0.0972 | 0.1385 |
| $N$ = 64 | 0.0238 | 0.0120 | 0.0216 | 0.0239 | 0.0119 | 0.5321 | 0.0461 | 0.0605 |
| $N$ = 128 | 0.0118 | 0.0058 | 0.0107 | 0.0118 | 0.0059 | −0.1839 | 0.0229 | 0.0297 |
| $N$ = 256 | 0.0059 | 0.0028 | 0.0054 | 0.0058 | 0.0028 | −1.0280 | 0.0110 | 0.0141 |
| $p$ = 8, $V(\mathbf{\Sigma})$ = 0 | | | | | | | | |
| $N$ = 4 | 0.4167 | 0.2651 | 0.3540 | 0.4186 | 0.2654 | 0.5067 | 0.9252 | 1.2858 |
| $N$ = 8 | 0.1786 | 0.0811 | 0.1654 | 0.1793 | 0.0806 | 0.6594 | 0.3282 | 0.4431 |
| $N$ = 16 | 0.0833 | 0.0297 | 0.0790 | 0.0835 | 0.0296 | 0.4185 | 0.1382 | 0.1726 |
| $N$ = 32 | 0.0403 | 0.0121 | 0.0390 | 0.0407 | 0.0124 | 1.9188 | 0.0629 | 0.0766 |
| $N$ = 64 | 0.0198 | 0.0054 | 0.0193 | 0.0199 | 0.0053 | 0.2683 | 0.0297 | 0.0351 |
| $N$ = 128 | 0.0098 | 0.0025 | 0.0096 | 0.0098 | 0.0025 | −0.6006 | 0.0144 | 0.0167 |
| $N$ = 256 | 0.0049 | 0.0012 | 0.0048 | 0.0049 | 0.0012 | 0.5840 | 0.0071 | 0.0083 |
| $p$ = 16, $V(\mathbf{\Sigma})$ = 0 | | | | | | | | |
| $N$ = 4 | 0.3750 | 0.1616 | 0.3480 | 0.3792 | 0.1663 | 1.7786 | 0.6905 | 0.9197 |
| $N$ = 8 | 0.1607 | 0.0477 | 0.1559 | 0.1614 | 0.0478 | 1.0293 | 0.2488 | 0.3007 |
| $N$ = 16 | 0.0750 | 0.0166 | 0.0737 | 0.0752 | 0.0168 | 0.8714 | 0.1055 | 0.1231 |
| $N$ = 32 | 0.0363 | 0.0064 | 0.0358 | 0.0362 | 0.0064 | −0.6019 | 0.0478 | 0.0545 |
| $N$ = 64 | 0.0179 | 0.0027 | 0.0177 | 0.0179 | 0.0027 | 0.9825 | 0.0228 | 0.0253 |
| $N$ = 128 | 0.0089 | 0.0012 | 0.0088 | 0.0088 | 0.0012 | −0.6922 | 0.0110 | 0.0120 |
| $N$ = 256 | 0.0044 | 0.0006 | 0.0044 | 0.0044 | 0.0006 | 1.3830 | 0.0054 | 0.0059 |
| $p$ = 32, $V(\mathbf{\Sigma})$ = 0 | | | | | | | | |
| $N$ = 4 | 0.3542 | 0.1053 | 0.3436 | 0.3559 | 0.1051 | 1.1876 | 0.5454 | 0.6531 |
| $N$ = 8 | 0.1518 | 0.0304 | 0.1482 | 0.1510 | 0.0307 | −1.8948 | 0.2065 | 0.2322 |
| $N$ = 16 | 0.0708 | 0.0102 | 0.0702 | 0.0708 | 0.0102 | −0.0003 | 0.0890 | 0.0971 |
| $N$ = 32 | 0.0343 | 0.0038 | 0.0341 | 0.0342 | 0.0037 | −0.4806 | 0.0405 | 0.0437 |
| $N$ = 64 | 0.0169 | 0.0015 | 0.0168 | 0.0169 | 0.0015 | −0.6023 | 0.0194 | 0.0206 |
| $N$ = 128 | 0.0084 | 0.0006 | 0.0083 | 0.0084 | 0.0006 | 1.0563 | 0.0094 | 0.0100 |
| $N$ = 256 | 0.0042 | 0.0003 | 0.0042 | 0.0042 | 0.0003 | 0.0593 | 0.0046 | 0.0049 |

(*continued*)

**Table S1.** (*continued*)

| | E[$V$(**S**)] | SD[$V$(**S**)] | Median | Mean | ESD | $T$ | CP 5% | CP 1% |
|---|---|---|---|---|---|---|---|---|
| $p = 64$, $V(\boldsymbol{\Sigma}) = 0$ | | | | | | | | |
| $N = 4$ | 0.3438 | 0.07123 | 0.3388 | 0.3445 | 0.07210 | 0.7602 | 0.4670 | 0.5373 |
| $N = 8$ | 0.1473 | 0.02028 | 0.1464 | 0.1477 | 0.02043 | 1.2947 | 0.1836 | 0.2027 |
| $N = 16$ | 0.0688 | 0.00665 | 0.0688 | 0.0689 | 0.00663 | 1.7306 | 0.0801 | 0.0856 |
| $N = 32$ | 0.0333 | 0.00236 | 0.0332 | 0.0333 | 0.00238 | 0.6721 | 0.0374 | 0.0391 |
| $N = 64$ | 0.0164 | 0.00089 | 0.0163 | 0.0164 | 0.00088 | −0.4407 | 0.0178 | 0.0184 |
| $N = 128$ | 0.0081 | 0.00036 | 0.0081 | 0.0081 | 0.00036 | −1.3996 | 0.0087 | 0.0090 |
| $N = 256$ | 0.0040 | 0.00015 | 0.0040 | 0.0040 | 0.00016 | 0.6570 | 0.0043 | 0.0044 |
| $p = 128$, $V(\boldsymbol{\Sigma}) = 0$ | | | | | | | | |
| $N = 4$ | 0.3385 | 0.04924 | 0.3373 | 0.3390 | 0.04854 | 0.7364 | 0.4213 | 0.4614 |
| $N = 8$ | 0.1451 | 0.01392 | 0.1446 | 0.1451 | 0.01379 | 0.1097 | 0.1684 | 0.1793 |
| $N = 16$ | 0.0677 | 0.00450 | 0.0676 | 0.0677 | 0.00450 | 0.4354 | 0.0753 | 0.0788 |
| $N = 32$ | 0.0328 | 0.00156 | 0.0328 | 0.0328 | 0.00155 | 2.7368 | 0.0354 | 0.0366 |
| $N = 64$ | 0.0161 | 0.00057 | 0.0161 | 0.0161 | 0.00057 | −0.2231 | 0.0171 | 0.0175 |
| $N = 128$ | 0.0080 | 0.00022 | 0.0080 | 0.0080 | 0.00022 | −1.7313 | 0.0084 | 0.0085 |
| $N = 256$ | 0.0040 | 0.00009 | 0.0040 | 0.0040 | 0.00009 | −0.0648 | 0.0041 | 0.0042 |
| $p = 256$, $V(\boldsymbol{\Sigma}) = 0$ | | | | | | | | |
| $N = 4$ | 0.3359 | 0.03442 | 0.3340 | 0.3354 | 0.03447 | −1.1403 | 0.3932 | 0.4240 |
| $N = 8$ | 0.1440 | 0.00969 | 0.1436 | 0.1440 | 0.00993 | 0.1215 | 0.1608 | 0.1691 |
| $N = 16$ | 0.0672 | 0.00311 | 0.0672 | 0.0672 | 0.00307 | 0.9739 | 0.0722 | 0.0744 |
| $N = 32$ | 0.0325 | 0.00106 | 0.0325 | 0.0325 | 0.00106 | −0.8132 | 0.0343 | 0.0350 |
| $N = 64$ | 0.0160 | 0.00038 | 0.0160 | 0.0160 | 0.00038 | 0.0842 | 0.0166 | 0.0169 |
| $N = 128$ | 0.0079 | 0.00014 | 0.0079 | 0.0079 | 0.00014 | 0.0262 | 0.0082 | 0.0083 |
| $N = 256$ | 0.0040 | 0.00005 | 0.0040 | 0.0040 | 0.00005 | −0.3846 | 0.0040 | 0.0041 |
| $p = 1024$, $V(\boldsymbol{\Sigma}) = 0$ | | | | | | | | |
| $N = 4$ | 0.3340 | 0.01706 | 0.3342 | 0.3343 | 0.01717 | 1.2126 | 0.3628 | 0.3745 |
| $N = 8$ | 0.1431 | 0.00479 | 0.1430 | 0.1431 | 0.00480 | −0.6874 | 0.1510 | 0.1550 |
| $N = 16$ | 0.0668 | 0.00153 | 0.0668 | 0.0668 | 0.00151 | 1.7324 | 0.0694 | 0.0705 |
| $N = 32$ | 0.0323 | 0.00052 | 0.0323 | 0.0323 | 0.00052 | 0.3227 | 0.0332 | 0.0336 |
| $N = 64$ | 0.0159 | 0.00018 | 0.0159 | 0.0159 | 0.00018 | 0.3522 | 0.0162 | 0.0163 |
| $N = 128$ | 0.0079 | 0.00006 | 0.0079 | 0.0079 | 0.00006 | −1.1216 | 0.0080 | 0.0080 |
| $N = 256$ | 0.0039 | 0.00002 | 0.0039 | 0.0039 | 0.00002 | 0.7640 | 0.0040 | 0.0040 |

(*continued*)

**Table S1.** (*continued*)

| | E[$V$(**S**)] | SD[$V$(**S**)] | Median | Mean | ESD | $T$ | Pow. 5% | Pow. 1% |
|---|---|---|---|---|---|---|---|---|
| $p = 2, q = 1, V(\mathbf{\Sigma}) = 0.1$ | | | | | | | | |
| $N = 4$ | 0.7667 | 1.2893 | 0.3299 | 0.7801 | 1.2981 | 0.7327 | 0.0658 | 0.0182 |
| $N = 8$ | 0.3857 | 0.5163 | 0.2079 | 0.3837 | 0.5087 | −0.2776 | 0.0888 | 0.0314 |
| $N = 16$ | 0.2333 | 0.2592 | 0.1451 | 0.2223 | 0.2419 | −3.2291 | 0.1566 | 0.0528 |
| $N = 32$ | 0.1645 | 0.1496 | 0.1244 | 0.1667 | 0.1497 | 1.0124 | 0.3126 | 0.1190 |
| $N = 64$ | 0.1317 | 0.0943 | 0.1124 | 0.1327 | 0.0954 | 0.6805 | 0.5860 | 0.3762 |
| $N = 128$ | 0.1157 | 0.0626 | 0.1055 | 0.1145 | 0.0623 | −1.3981 | 0.8884 | 0.7414 |
| $N = 256$ | 0.1078 | 0.0429 | 0.1042 | 0.1090 | 0.0429 | 1.9180 | 0.9972 | 0.9806 |
| $p = 2, q = 1, V(\mathbf{\Sigma}) = 0.2$ | | | | | | | | |
| $N = 4$ | 0.8667 | 1.5241 | 0.3716 | 0.9230 | 1.6659 | 2.3912 | 0.0866 | 0.0248 |
| $N = 8$ | 0.4857 | 0.6465 | 0.2615 | 0.4777 | 0.6313 | −0.8980 | 0.1372 | 0.0500 |
| $N = 16$ | 0.3333 | 0.3429 | 0.2311 | 0.3257 | 0.3300 | −1.6266 | 0.2760 | 0.1186 |
| $N = 32$ | 0.2645 | 0.2067 | 0.2101 | 0.2633 | 0.2065 | −0.4030 | 0.5362 | 0.2804 |
| $N = 64$ | 0.2317 | 0.1341 | 0.2066 | 0.2312 | 0.1350 | −0.2979 | 0.8704 | 0.7194 |
| $N = 128$ | 0.2157 | 0.0907 | 0.2048 | 0.2160 | 0.0899 | 0.1913 | 0.9952 | 0.9814 |
| $N = 256$ | 0.2078 | 0.0627 | 0.2028 | 0.2084 | 0.0625 | 0.5752 | 1.0000 | 1.0000 |
| $p = 2, q = 1, V(\mathbf{\Sigma}) = 0.4$ | | | | | | | | |
| $N = 4$ | 1.0667 | 1.9368 | 0.4120 | 1.0723 | 1.8519 | 0.2145 | 0.1132 | 0.0376 |
| $N = 8$ | 0.6857 | 0.8717 | 0.3967 | 0.6901 | 0.8663 | 0.3572 | 0.2302 | 0.0964 |
| $N = 16$ | 0.5333 | 0.4852 | 0.3972 | 0.5426 | 0.4981 | 1.3183 | 0.4828 | 0.2758 |
| $N = 32$ | 0.4645 | 0.3023 | 0.3936 | 0.4616 | 0.3011 | −0.6854 | 0.8316 | 0.5958 |
| $N = 64$ | 0.4317 | 0.2002 | 0.4044 | 0.4337 | 0.2019 | 0.6886 | 0.9944 | 0.9722 |
| $N = 128$ | 0.4157 | 0.1369 | 0.3983 | 0.4142 | 0.1346 | −0.8153 | 1.0000 | 1.0000 |
| $N = 256$ | 0.4078 | 0.0952 | 0.3994 | 0.4071 | 0.0957 | −0.5627 | 1.0000 | 1.0000 |
| $p = 2, q = 1, V(\mathbf{\Sigma}) = 0.6$ | | | | | | | | |
| $N = 4$ | 1.2667 | 2.3068 | 0.4407 | 1.2057 | 2.1608 | −1.9958 | 0.1300 | 0.0474 |
| $N = 8$ | 0.8857 | 1.0743 | 0.5312 | 0.8717 | 1.0467 | −0.9497 | 0.2988 | 0.1446 |
| $N = 16$ | 0.7333 | 0.6131 | 0.5716 | 0.7383 | 0.6161 | 0.5690 | 0.6422 | 0.4196 |
| $N = 32$ | 0.6645 | 0.3882 | 0.5839 | 0.6640 | 0.3847 | −0.1014 | 0.9538 | 0.8224 |
| $N = 64$ | 0.6317 | 0.2594 | 0.5896 | 0.6302 | 0.2601 | −0.4312 | 0.9998 | 0.9982 |
| $N = 128$ | 0.6157 | 0.1783 | 0.5994 | 0.6151 | 0.1753 | −0.2601 | 1.0000 | 1.0000 |
| $N = 256$ | 0.6078 | 0.1243 | 0.5978 | 0.6063 | 0.1218 | −0.9039 | 1.0000 | 1.0000 |
| $p = 2, q = 1, V(\mathbf{\Sigma}) = 0.8$ | | | | | | | | |
| $N = 4$ | 1.4667 | 2.6522 | 0.5269 | 1.4956 | 2.9055 | 0.7042 | 0.1656 | 0.0716 |
| $N = 8$ | 1.0857 | 1.2651 | 0.6587 | 1.0792 | 1.3030 | −0.3520 | 0.3674 | 0.1988 |
| $N = 16$ | 0.9333 | 0.7343 | 0.7440 | 0.9364 | 0.7250 | 0.3003 | 0.7620 | 0.5484 |
| $N = 32$ | 0.8645 | 0.4698 | 0.7732 | 0.8675 | 0.4675 | 0.4571 | 0.9878 | 0.9220 |
| $N = 64$ | 0.8317 | 0.3157 | 0.7840 | 0.8333 | 0.3172 | 0.3574 | 1.0000 | 0.9998 |
| $N = 128$ | 0.8157 | 0.2176 | 0.7915 | 0.8154 | 0.2182 | −0.1085 | 1.0000 | 1.0000 |
| $N = 256$ | 0.8078 | 0.1519 | 0.7970 | 0.8111 | 0.1515 | 1.5198 | 1.0000 | 1.0000 |

(*continued*)

**Table S1.** (*continued*)

| | E[$V$(**S**)] | SD[$V$(**S**)] | Median | Mean | ESD | $T$ | Pow. 5% | Pow. 1% |
|---|---|---|---|---|---|---|---|---|
| $p = 4, q = 1, V(\boldsymbol{\Sigma}) = 0.1$ | | | | | | | | |
| $N = 4$ | 0.6167 | 0.7999 | 0.3641 | 0.6246 | 0.8476 | 0.6598 | 0.1002 | 0.0352 |
| $N = 8$ | 0.3214 | 0.3225 | 0.2245 | 0.3334 | 0.3544 | 2.3979 | 0.1866 | 0.0862 |
| $N = 16$ | 0.2033 | 0.1646 | 0.1606 | 0.2052 | 0.1617 | 0.8205 | 0.3528 | 0.1910 |
| $N = 32$ | 0.1500 | 0.0968 | 0.1288 | 0.1513 | 0.0974 | 0.9326 | 0.6670 | 0.4546 |
| $N = 64$ | 0.1246 | 0.0619 | 0.1130 | 0.1253 | 0.0633 | 0.7650 | 0.9492 | 0.8778 |
| $N = 128$ | 0.1122 | 0.0415 | 0.1069 | 0.1129 | 0.0420 | 1.2435 | 0.9994 | 0.9976 |
| $N = 256$ | 0.1061 | 0.0286 | 0.1030 | 0.1056 | 0.0284 | −1.1421 | 1.0000 | 1.0000 |
| $p = 4, q = 1, V(\boldsymbol{\Sigma}) = 0.2$ | | | | | | | | |
| $N = 4$ | 0.7333 | 1.0876 | 0.3700 | 0.7096 | 1.0191 | −1.6486 | 0.1308 | 0.0552 |
| $N = 8$ | 0.4286 | 0.4692 | 0.2755 | 0.4334 | 0.4851 | 0.7103 | 0.2700 | 0.1490 |
| $N = 16$ | 0.3067 | 0.2532 | 0.2379 | 0.3087 | 0.2541 | 0.5529 | 0.5504 | 0.3758 |
| $N = 32$ | 0.2516 | 0.1546 | 0.2207 | 0.2512 | 0.1517 | −0.1925 | 0.8916 | 0.7652 |
| $N = 64$ | 0.2254 | 0.1012 | 0.2085 | 0.2257 | 0.1016 | 0.1918 | 0.9976 | 0.9908 |
| $N = 128$ | 0.2126 | 0.0688 | 0.2031 | 0.2112 | 0.0683 | −1.4846 | 1.0000 | 1.0000 |
| $N = 256$ | 0.2063 | 0.0477 | 0.2029 | 0.2069 | 0.0478 | 0.9460 | 1.0000 | 1.0000 |
| $p = 4, q = 1, V(\boldsymbol{\Sigma}) = 0.4$ | | | | | | | | |
| $N = 4$ | 0.9667 | 1.6064 | 0.4096 | 0.9742 | 1.5788 | 0.3372 | 0.1936 | 0.1026 |
| $N = 8$ | 0.6429 | 0.7343 | 0.3949 | 0.6372 | 0.7364 | −0.5426 | 0.4134 | 0.2690 |
| $N = 16$ | 0.5133 | 0.4137 | 0.3968 | 0.5132 | 0.4183 | −0.0225 | 0.7684 | 0.6234 |
| $N = 32$ | 0.4548 | 0.2598 | 0.3937 | 0.4544 | 0.2661 | −0.1087 | 0.9888 | 0.9584 |
| $N = 64$ | 0.4270 | 0.1728 | 0.4025 | 0.4292 | 0.1756 | 0.8791 | 1.0000 | 0.9998 |
| $N = 128$ | 0.4134 | 0.1185 | 0.4011 | 0.4122 | 0.1168 | −0.7078 | 1.0000 | 1.0000 |
| $N = 256$ | 0.4067 | 0.0825 | 0.3970 | 0.4038 | 0.0806 | −2.5084 | 1.0000 | 1.0000 |
| $p = 4, q = 1, V(\boldsymbol{\Sigma}) = 0.6$ | | | | | | | | |
| $N = 4$ | 1.2000 | 2.0850 | 0.4763 | 1.1770 | 2.0466 | −0.7938 | 0.2426 | 0.1370 |
| $N = 8$ | 0.8571 | 0.9812 | 0.5485 | 0.8444 | 0.9432 | −0.9582 | 0.5276 | 0.3806 |
| $N = 16$ | 0.7200 | 0.5643 | 0.5634 | 0.7104 | 0.5573 | −1.2209 | 0.8732 | 0.7710 |
| $N = 32$ | 0.6581 | 0.3589 | 0.5820 | 0.6539 | 0.3516 | −0.8294 | 0.9970 | 0.9882 |
| $N = 64$ | 0.6286 | 0.2405 | 0.5917 | 0.6291 | 0.2451 | 0.1460 | 1.0000 | 1.0000 |
| $N = 128$ | 0.6142 | 0.1655 | 0.5923 | 0.6084 | 0.1634 | −2.5079 | 1.0000 | 1.0000 |
| $N = 256$ | 0.6071 | 0.1154 | 0.5971 | 0.6067 | 0.1175 | −0.2168 | 1.0000 | 1.0000 |
| $p = 4, q = 1, V(\boldsymbol{\Sigma}) = 0.8$ | | | | | | | | |
| $N = 4$ | 1.4333 | 2.5407 | 0.5520 | 1.3960 | 2.4201 | −1.0902 | 0.2744 | 0.1672 |
| $N = 8$ | 1.0714 | 1.2182 | 0.6903 | 1.0730 | 1.2081 | 0.0917 | 0.6150 | 0.4626 |
| $N = 16$ | 0.9267 | 0.7096 | 0.7454 | 0.9328 | 0.7139 | 0.6117 | 0.9306 | 0.8646 |
| $N = 32$ | 0.8613 | 0.4549 | 0.7682 | 0.8585 | 0.4565 | −0.4368 | 0.9994 | 0.9978 |
| $N = 64$ | 0.8302 | 0.3061 | 0.7903 | 0.8350 | 0.3069 | 1.1137 | 1.0000 | 1.0000 |
| $N = 128$ | 0.8150 | 0.2111 | 0.7929 | 0.8139 | 0.2120 | −0.3642 | 1.0000 | 1.0000 |
| $N = 256$ | 0.8075 | 0.1474 | 0.7948 | 0.8079 | 0.1470 | 0.2033 | 1.0000 | 1.0000 |

(*continued*)

**Table S1.** (*continued*)

| | E[$V$(**S**)] | SD[$V$(**S**)] | Median | Mean | ESD | $T$ | Pow. 5% | Pow. 1% |
|---|---|---|---|---|---|---|---|---|
| $p = 4, q = 2, V(\mathbf{\Sigma}) = 0.1$ | | | | | | | | |
| $N = 4$ | 0.6167 | 0.7295 | 0.3864 | 0.6220 | 0.7469 | 0.5028 | 0.0982 | 0.0316 |
| $N = 8$ | 0.3214 | 0.2772 | 0.2483 | 0.3227 | 0.2693 | 0.3344 | 0.1798 | 0.0652 |
| $N = 16$ | 0.2033 | 0.1318 | 0.1731 | 0.2031 | 0.1311 | −0.1489 | 0.3632 | 0.1676 |
| $N = 32$ | 0.1500 | 0.0725 | 0.1380 | 0.1510 | 0.0723 | 1.0047 | 0.7642 | 0.4980 |
| $N = 64$ | 0.1246 | 0.0441 | 0.1186 | 0.1251 | 0.0455 | 0.7961 | 0.9920 | 0.9602 |
| $N = 128$ | 0.1122 | 0.0286 | 0.1099 | 0.1127 | 0.0288 | 1.1330 | 1.0000 | 1.0000 |
| $N = 256$ | 0.1061 | 0.0193 | 0.1041 | 0.1054 | 0.0193 | −2.5193 | 1.0000 | 1.0000 |
| $p = 4, q = 2, V(\mathbf{\Sigma}) = 0.2$ | | | | | | | | |
| $N = 4$ | 0.7333 | 0.9288 | 0.4256 | 0.7380 | 0.9311 | 0.3541 | 0.1418 | 0.0488 |
| $N = 8$ | 0.4286 | 0.3717 | 0.3198 | 0.4179 | 0.3578 | −2.1073 | 0.2826 | 0.1270 |
| $N = 16$ | 0.3067 | 0.1855 | 0.2651 | 0.3085 | 0.1873 | 0.7019 | 0.6378 | 0.3982 |
| $N = 32$ | 0.2516 | 0.1061 | 0.2384 | 0.2556 | 0.1084 | 2.6090 | 0.9726 | 0.8884 |
| $N = 64$ | 0.2254 | 0.0664 | 0.2173 | 0.2260 | 0.0666 | 0.6088 | 1.0000 | 1.0000 |
| $N = 128$ | 0.2126 | 0.0439 | 0.2088 | 0.2128 | 0.0453 | 0.3361 | 1.0000 | 1.0000 |
| $N = 256$ | 0.2063 | 0.0300 | 0.2050 | 0.2064 | 0.0300 | 0.3695 | 1.0000 | 1.0000 |
| $p = 4, q = 2, V(\mathbf{\Sigma}) = 0.4$ | | | | | | | | |
| (This comformation is impossible) | | | | | | | | |

(*continued*)

**Table S1.** (*continued*)

| | E[$V$($\mathbf{S}$)] | SD[$V$($\mathbf{S}$)] | Median | Mean | ESD | $T$ | Pow. 5% | Pow. 1% |
|---|---|---|---|---|---|---|---|---|
| $p = 8, q = 1, V(\mathbf{\Sigma}) = 0.1$ | | | | | | | | |
| $N = 4$ | 0.5417 | 0.5946 | 0.3545 | 0.5388 | 0.6115 | −0.3270 | 0.1390 | 0.0752 |
| $N = 8$ | 0.2893 | 0.2455 | 0.2127 | 0.2849 | 0.2393 | −1.3039 | 0.2810 | 0.1660 |
| $N = 16$ | 0.1883 | 0.1281 | 0.1522 | 0.1860 | 0.1272 | −1.2985 | 0.5616 | 0.4194 |
| $N = 32$ | 0.1427 | 0.0766 | 0.1271 | 0.1425 | 0.0758 | −0.2525 | 0.9044 | 0.8270 |
| $N = 64$ | 0.1210 | 0.0495 | 0.1122 | 0.1213 | 0.0498 | 0.3606 | 0.9984 | 0.9950 |
| $N = 128$ | 0.1104 | 0.0334 | 0.1061 | 0.1098 | 0.0333 | −1.3971 | 1.0000 | 1.0000 |
| $N = 256$ | 0.1052 | 0.0231 | 0.1030 | 0.1054 | 0.0236 | 0.5513 | 1.0000 | 1.0000 |
| $p = 8, q = 1, V(\mathbf{\Sigma}) = 0.2$ | | | | | | | | |
| $N = 4$ | 0.6667 | 0.8974 | 0.3775 | 0.6835 | 0.9034 | 1.3184 | 0.2118 | 0.1336 |
| $N = 8$ | 0.4000 | 0.3948 | 0.2819 | 0.4013 | 0.3782 | 0.2518 | 0.4278 | 0.3112 |
| $N = 16$ | 0.2933 | 0.2163 | 0.2323 | 0.2891 | 0.2124 | −1.4124 | 0.7642 | 0.6644 |
| $N = 32$ | 0.2452 | 0.1335 | 0.2173 | 0.2469 | 0.1372 | 0.8894 | 0.9836 | 0.9652 |
| $N = 64$ | 0.2222 | 0.0880 | 0.2090 | 0.2224 | 0.0903 | 0.1113 | 1.0000 | 0.9998 |
| $N = 128$ | 0.2110 | 0.0600 | 0.2055 | 0.2122 | 0.0592 | 1.3676 | 1.0000 | 1.0000 |
| $N = 256$ | 0.2055 | 0.0416 | 0.2026 | 0.2054 | 0.0417 | −0.1800 | 1.0000 | 1.0000 |
| $p = 8, q = 1, V(\mathbf{\Sigma}) = 0.4$ | | | | | | | | |
| $N = 4$ | 0.9167 | 1.4545 | 0.4073 | 0.8964 | 1.3932 | −1.0283 | 0.2758 | 0.2010 |
| $N = 8$ | 0.6214 | 0.6726 | 0.3993 | 0.6180 | 0.6724 | −0.3572 | 0.5748 | 0.4674 |
| $N = 16$ | 0.5033 | 0.3821 | 0.3975 | 0.5043 | 0.3798 | 0.1820 | 0.9124 | 0.8712 |
| $N = 32$ | 0.4500 | 0.2412 | 0.4025 | 0.4510 | 0.2414 | 0.2901 | 0.9990 | 0.9964 |
| $N = 64$ | 0.4246 | 0.1610 | 0.4031 | 0.4243 | 0.1586 | −0.1199 | 1.0000 | 1.0000 |
| $N = 128$ | 0.4122 | 0.1105 | 0.3995 | 0.4130 | 0.1108 | 0.5315 | 1.0000 | 1.0000 |
| $N = 256$ | 0.4061 | 0.0770 | 0.3993 | 0.4053 | 0.0762 | −0.7337 | 1.0000 | 1.0000 |
| $p = 8, q = 1, V(\mathbf{\Sigma}) = 0.6$ | | | | | | | | |
| $N = 4$ | 1.1667 | 1.9793 | 0.4641 | 1.1773 | 2.0289 | 0.3698 | 0.3310 | 0.2544 |
| $N = 8$ | 0.8429 | 0.9374 | 0.5468 | 0.8327 | 0.9047 | −0.7964 | 0.6726 | 0.5776 |
| $N = 16$ | 0.7133 | 0.5415 | 0.5891 | 0.7236 | 0.5308 | 1.3632 | 0.9568 | 0.9324 |
| $N = 32$ | 0.6548 | 0.3454 | 0.5764 | 0.6494 | 0.3566 | −1.0725 | 0.9998 | 0.9988 |
| $N = 64$ | 0.6270 | 0.2318 | 0.5906 | 0.6254 | 0.2324 | −0.4722 | 1.0000 | 1.0000 |
| $N = 128$ | 0.6134 | 0.1596 | 0.5988 | 0.6135 | 0.1565 | 0.0522 | 1.0000 | 1.0000 |
| $N = 256$ | 0.6067 | 0.1114 | 0.5957 | 0.6051 | 0.1106 | −0.9916 | 1.0000 | 1.0000 |
| $p = 8, q = 1, V(\mathbf{\Sigma}) = 0.8$ | | | | | | | | |
| $N = 4$ | 1.4167 | 2.4862 | 0.5572 | 1.4707 | 2.7285 | 1.3993 | 0.3852 | 0.3018 |
| $N = 8$ | 1.0643 | 1.1954 | 0.6913 | 1.0524 | 1.1663 | −0.7186 | 0.7356 | 0.6420 |
| $N = 16$ | 0.9233 | 0.6977 | 0.7348 | 0.9074 | 0.6883 | −1.6414 | 0.9750 | 0.9562 |
| $N = 32$ | 0.8597 | 0.4477 | 0.7714 | 0.8582 | 0.4380 | −0.2392 | 1.0000 | 1.0000 |
| $N = 64$ | 0.8294 | 0.3015 | 0.7840 | 0.8222 | 0.2990 | −1.6841 | 1.0000 | 1.0000 |
| $N = 128$ | 0.8146 | 0.2080 | 0.7942 | 0.8167 | 0.2067 | 0.7183 | 1.0000 | 1.0000 |
| $N = 256$ | 0.8073 | 0.1453 | 0.7949 | 0.8054 | 0.1443 | −0.9026 | 1.0000 | 1.0000 |

(*continued*)

**Table S1.** (*continued*)

| | E[$V$(**S**)] | SD[$V$(**S**)] | Median | Mean | ESD | $T$ | Pow. 5% | Pow. 1% |
|---|---|---|---|---|---|---|---|---|
| $p = 8, q = 2, V(\boldsymbol{\Sigma}) = 0.1$ | | | | | | | | |
| $N = 4$ | 0.5417 | 0.5297 | 0.3831 | 0.5358 | 0.5142 | −0.8102 | 0.1444 | 0.0754 |
| $N = 8$ | 0.2893 | 0.2070 | 0.2363 | 0.2909 | 0.2077 | 0.5498 | 0.3160 | 0.1700 |
| $N = 16$ | 0.1883 | 0.1021 | 0.1667 | 0.1870 | 0.0997 | −0.9397 | 0.6386 | 0.4742 |
| $N = 32$ | 0.1427 | 0.0583 | 0.1334 | 0.1435 | 0.0597 | 0.9048 | 0.9594 | 0.9088 |
| $N = 64$ | 0.1210 | 0.0365 | 0.1158 | 0.1195 | 0.0358 | −2.9783 | 0.9998 | 0.9990 |
| $N = 128$ | 0.1104 | 0.0242 | 0.1089 | 0.1106 | 0.0242 | 0.4686 | 1.0000 | 1.0000 |
| $N = 256$ | 0.1052 | 0.0165 | 0.1044 | 0.1052 | 0.0164 | 0.0586 | 1.0000 | 1.0000 |
| $p = 8, q = 2, V(\boldsymbol{\Sigma}) = 0.2$ | | | | | | | | |
| $N = 4$ | 0.6667 | 0.7567 | 0.4338 | 0.6849 | 0.7664 | 1.6786 | 0.2248 | 0.1390 |
| $N = 8$ | 0.4000 | 0.3138 | 0.3126 | 0.3976 | 0.3228 | −0.5188 | 0.4742 | 0.3170 |
| $N = 16$ | 0.2933 | 0.1629 | 0.2554 | 0.2901 | 0.1616 | −1.4358 | 0.8592 | 0.7680 |
| $N = 32$ | 0.2452 | 0.0966 | 0.2299 | 0.2446 | 0.0962 | −0.3991 | 0.9990 | 0.9944 |
| $N = 64$ | 0.2222 | 0.0620 | 0.2144 | 0.2215 | 0.0629 | −0.8630 | 1.0000 | 1.0000 |
| $N = 128$ | 0.2110 | 0.0417 | 0.2077 | 0.2108 | 0.0413 | −0.3914 | 1.0000 | 1.0000 |
| $N = 256$ | 0.2055 | 0.0287 | 0.2044 | 0.2057 | 0.0284 | 0.5480 | 1.0000 | 1.0000 |
| $p = 8, q = 2, V(\boldsymbol{\Sigma}) = 0.4$ | | | | | | | | |
| $N = 4$ | 0.9167 | 1.1622 | 0.5207 | 0.9034 | 1.1518 | −0.8149 | 0.3152 | 0.2154 |
| $N = 8$ | 0.6214 | 0.5071 | 0.4870 | 0.6244 | 0.5230 | 0.4001 | 0.6742 | 0.5432 |
| $N = 16$ | 0.5033 | 0.2744 | 0.4443 | 0.5015 | 0.2685 | −0.4783 | 0.9794 | 0.9514 |
| $N = 32$ | 0.4500 | 0.1675 | 0.4209 | 0.4491 | 0.1695 | −0.3959 | 0.9998 | 0.9998 |
| $N = 64$ | 0.4246 | 0.1095 | 0.4105 | 0.4212 | 0.1079 | −2.2123 | 1.0000 | 1.0000 |
| $N = 128$ | 0.4122 | 0.0744 | 0.4086 | 0.4131 | 0.0736 | 0.8143 | 1.0000 | 1.0000 |
| $N = 256$ | 0.4061 | 0.0515 | 0.4028 | 0.4054 | 0.0514 | −0.9666 | 1.0000 | 1.0000 |
| $p = 8, q = 4, V(\boldsymbol{\Sigma}) = 0.1$ | | | | | | | | |
| $N = 4$ | 0.5417 | 0.4697 | 0.4193 | 0.5603 | 0.4833 | 2.7253 | 0.1542 | 0.0784 |
| $N = 8$ | 0.2893 | 0.1721 | 0.2525 | 0.2901 | 0.1715 | 0.3491 | 0.3180 | 0.1606 |
| $N = 16$ | 0.1883 | 0.0787 | 0.1754 | 0.1887 | 0.0785 | 0.3747 | 0.7164 | 0.5158 |
| $N = 32$ | 0.1427 | 0.0417 | 0.1381 | 0.1435 | 0.0423 | 1.2407 | 0.9950 | 0.9750 |
| $N = 64$ | 0.1210 | 0.0245 | 0.1193 | 0.1215 | 0.0246 | 1.2929 | 1.0000 | 1.0000 |
| $N = 128$ | 0.1104 | 0.0156 | 0.1098 | 0.1107 | 0.0155 | 1.2881 | 1.0000 | 1.0000 |
| $N = 256$ | 0.1052 | 0.0104 | 0.1046 | 0.1049 | 0.0103 | −1.9227 | 1.0000 | 1.0000 |
| $p = 8, q = 4, V(\boldsymbol{\Sigma}) = 0.2$ | | | | | | | | |
| (This comformation is impossible) | | | | | | | | |

(*continued*)

**Table S1.** (*continued*)

| | E[$V$(**S**)] | SD[$V$(**S**)] | Median | Mean | ESD | $T$ | Pow. 5% | Pow. 1% |
|---|---|---|---|---|---|---|---|---|
| $p = 16, q = 1, V(\mathbf{\Sigma}) = 0.1$ | | | | | | | | |
| $N = 4$ | 0.5042 | 0.4994 | 0.3516 | 0.5085 | 0.5040 | 0.6104 | 0.2026 | 0.1218 |
| $N = 8$ | 0.2732 | 0.2102 | 0.2165 | 0.2773 | 0.2092 | 1.3950 | 0.4158 | 0.3142 |
| $N = 16$ | 0.1808 | 0.1114 | 0.1518 | 0.1799 | 0.1115 | −0.6051 | 0.7332 | 0.6342 |
| $N = 32$ | 0.1391 | 0.0673 | 0.1257 | 0.1393 | 0.0685 | 0.1796 | 0.9782 | 0.9564 |
| $N = 64$ | 0.1192 | 0.0438 | 0.1121 | 0.1196 | 0.0443 | 0.5738 | 0.9998 | 0.9998 |
| $N = 128$ | 0.1095 | 0.0297 | 0.1063 | 0.1097 | 0.0302 | 0.3760 | 1.0000 | 1.0000 |
| $N = 256$ | 0.1048 | 0.0205 | 0.1034 | 0.1048 | 0.0205 | 0.3061 | 1.0000 | 1.0000 |
| $p = 16, q = 1, V(\mathbf{\Sigma}) = 0.2$ | | | | | | | | |
| $N = 4$ | 0.6333 | 0.8077 | 0.3713 | 0.6342 | 0.8024 | 0.0729 | 0.2688 | 0.1880 |
| $N = 8$ | 0.3857 | 0.3598 | 0.2756 | 0.3878 | 0.3671 | 0.3950 | 0.5466 | 0.4586 |
| $N = 16$ | 0.2867 | 0.1990 | 0.2375 | 0.2847 | 0.1933 | −0.7042 | 0.8770 | 0.8272 |
| $N = 32$ | 0.2419 | 0.1235 | 0.2168 | 0.2395 | 0.1226 | −1.3960 | 0.9970 | 0.9930 |
| $N = 64$ | 0.2206 | 0.0817 | 0.2092 | 0.2206 | 0.0817 | −0.0454 | 1.0000 | 1.0000 |
| $N = 128$ | 0.2102 | 0.0558 | 0.2030 | 0.2087 | 0.0554 | −1.9179 | 1.0000 | 1.0000 |
| $N = 256$ | 0.2051 | 0.0388 | 0.2015 | 0.2050 | 0.0385 | −0.1718 | 1.0000 | 1.0000 |
| $p = 16, q = 1, V(\mathbf{\Sigma}) = 0.4$ | | | | | | | | |
| $N = 4$ | 0.8917 | 1.3811 | 0.4263 | 0.8759 | 1.2652 | −0.8801 | 0.3518 | 0.2774 |
| $N = 8$ | 0.6107 | 0.6427 | 0.4130 | 0.6165 | 0.6509 | 0.6282 | 0.6822 | 0.6172 |
| $N = 16$ | 0.4983 | 0.3668 | 0.4060 | 0.5036 | 0.3748 | 0.9922 | 0.9542 | 0.9330 |
| $N = 32$ | 0.4476 | 0.2322 | 0.4057 | 0.4492 | 0.2314 | 0.5001 | 0.9998 | 0.9994 |
| $N = 64$ | 0.4234 | 0.1552 | 0.4005 | 0.4227 | 0.1546 | −0.3205 | 1.0000 | 1.0000 |
| $N = 128$ | 0.4116 | 0.1067 | 0.4036 | 0.4144 | 0.1082 | 1.8203 | 1.0000 | 1.0000 |
| $N = 256$ | 0.4058 | 0.0743 | 0.4000 | 0.4056 | 0.0753 | −0.2074 | 1.0000 | 1.0000 |
| $p = 16, q = 1, V(\mathbf{\Sigma}) = 0.6$ | | | | | | | | |
| $N = 4$ | 1.1500 | 1.9274 | 0.4797 | 1.1959 | 2.0814 | 1.5607 | 0.4140 | 0.3374 |
| $N = 8$ | 0.8357 | 0.9159 | 0.5480 | 0.8416 | 0.9194 | 0.4545 | 0.7496 | 0.7020 |
| $N = 16$ | 0.7100 | 0.5304 | 0.5720 | 0.7164 | 0.5430 | 0.8347 | 0.9762 | 0.9668 |
| $N = 32$ | 0.6532 | 0.3388 | 0.5914 | 0.6559 | 0.3432 | 0.5582 | 1.0000 | 1.0000 |
| $N = 64$ | 0.6262 | 0.2275 | 0.5929 | 0.6268 | 0.2284 | 0.1861 | 1.0000 | 1.0000 |
| $N = 128$ | 0.6130 | 0.1568 | 0.6006 | 0.6158 | 0.1603 | 1.2290 | 1.0000 | 1.0000 |
| $N = 256$ | 0.6065 | 0.1094 | 0.5974 | 0.6048 | 0.1087 | −1.0764 | 1.0000 | 1.0000 |
| $p = 16, q = 1, V(\mathbf{\Sigma}) = 0.8$ | | | | | | | | |
| $N = 4$ | 1.4083 | 2.4591 | 0.5328 | 1.3669 | 2.3491 | −1.2462 | 0.4438 | 0.3742 |
| $N = 8$ | 1.0607 | 1.1840 | 0.6778 | 1.0687 | 1.1804 | 0.4765 | 0.8080 | 0.7666 |
| $N = 16$ | 0.9217 | 0.6917 | 0.7517 | 0.9147 | 0.6842 | −0.7221 | 0.9876 | 0.9814 |
| $N = 32$ | 0.8589 | 0.4442 | 0.7847 | 0.8683 | 0.4387 | 1.5248 | 1.0000 | 1.0000 |
| $N = 64$ | 0.8290 | 0.2992 | 0.7925 | 0.8358 | 0.3067 | 1.5855 | 1.0000 | 1.0000 |
| $N = 128$ | 0.8144 | 0.2065 | 0.7840 | 0.8075 | 0.2054 | −2.3764 | 1.0000 | 1.0000 |
| $N = 256$ | 0.8072 | 0.1442 | 0.7955 | 0.8072 | 0.1452 | 0.0030 | 1.0000 | 1.0000 |

(*continued*)

**Table S1.** (*continued*)

| | E[$V$(**S**)] | SD[$V$(**S**)] | Median | Mean | ESD | $T$ | Pow. 5% | Pow. 1% |
|---|---|---|---|---|---|---|---|---|
| $p = 16, q = 2, V(\mathbf{\Sigma}) = 0.1$ | | | | | | | | |
| $N = 4$ | 0.5042 | 0.4346 | 0.3721 | 0.4893 | 0.4066 | −2.5905 | 0.2082 | 0.1128 |
| $N = 8$ | 0.2732 | 0.1733 | 0.2255 | 0.2702 | 0.1699 | −1.2467 | 0.4370 | 0.3182 |
| $N = 16$ | 0.1808 | 0.0874 | 0.1631 | 0.1813 | 0.0871 | 0.3576 | 0.8220 | 0.7294 |
| $N = 32$ | 0.1391 | 0.0508 | 0.1323 | 0.1398 | 0.0516 | 0.9766 | 0.9976 | 0.9928 |
| $N = 64$ | 0.1192 | 0.0322 | 0.1162 | 0.1195 | 0.0325 | 0.5145 | 1.0000 | 1.0000 |
| $N = 128$ | 0.1095 | 0.0215 | 0.1078 | 0.1091 | 0.0214 | −1.3660 | 1.0000 | 1.0000 |
| $N = 256$ | 0.1048 | 0.0148 | 0.1036 | 0.1045 | 0.0148 | −1.3745 | 1.0000 | 1.0000 |
| $p = 16, q = 2, V(\mathbf{\Sigma}) = 0.2$ | | | | | | | | |
| $N = 4$ | 0.6333 | 0.6717 | 0.4132 | 0.6307 | 0.6606 | −0.2827 | 0.3020 | 0.2034 |
| $N = 8$ | 0.3857 | 0.2838 | 0.3111 | 0.3877 | 0.2868 | 0.4978 | 0.6306 | 0.5186 |
| $N = 16$ | 0.2867 | 0.1500 | 0.2536 | 0.2841 | 0.1491 | −1.2343 | 0.9478 | 0.9116 |
| $N = 32$ | 0.2419 | 0.0902 | 0.2291 | 0.2414 | 0.0899 | −0.4062 | 0.9998 | 0.9994 |
| $N = 64$ | 0.2206 | 0.0585 | 0.2120 | 0.2185 | 0.0582 | −2.6399 | 1.0000 | 1.0000 |
| $N = 128$ | 0.2102 | 0.0395 | 0.2068 | 0.2095 | 0.0396 | −1.3000 | 1.0000 | 1.0000 |
| $N = 256$ | 0.2051 | 0.0273 | 0.2042 | 0.2057 | 0.0265 | 1.5602 | 1.0000 | 1.0000 |
| $p = 16, q = 2, V(\mathbf{\Sigma}) = 0.4$ | | | | | | | | |
| $N = 4$ | 0.8917 | 1.1035 | 0.5242 | 0.8854 | 1.0836 | −0.4089 | 0.4082 | 0.3040 |
| $N = 8$ | 0.6107 | 0.4889 | 0.4676 | 0.6002 | 0.4853 | −1.5280 | 0.7828 | 0.7114 |
| $N = 16$ | 0.4983 | 0.2682 | 0.4488 | 0.5020 | 0.2713 | 0.9563 | 0.9924 | 0.9860 |
| $N = 32$ | 0.4476 | 0.1654 | 0.4249 | 0.4473 | 0.1616 | −0.1307 | 1.0000 | 1.0000 |
| $N = 64$ | 0.4234 | 0.1089 | 0.4093 | 0.4237 | 0.1108 | 0.1666 | 1.0000 | 1.0000 |
| $N = 128$ | 0.4116 | 0.0742 | 0.4045 | 0.4114 | 0.0739 | −0.2290 | 1.0000 | 1.0000 |
| $N = 256$ | 0.4058 | 0.0515 | 0.4017 | 0.4057 | 0.0523 | −0.1365 | 1.0000 | 1.0000 |
| $p = 16, q = 4, V(\mathbf{\Sigma}) = 0.1$ | | | | | | | | |
| $N = 4$ | 0.5042 | 0.3844 | 0.4028 | 0.5101 | 0.3902 | 1.0802 | 0.2238 | 0.1162 |
| $N = 8$ | 0.2732 | 0.1449 | 0.2417 | 0.2711 | 0.1416 | −1.0800 | 0.4776 | 0.3316 |
| $N = 16$ | 0.1808 | 0.0688 | 0.1713 | 0.1819 | 0.0702 | 1.1022 | 0.8972 | 0.7998 |
| $N = 32$ | 0.1391 | 0.0379 | 0.1335 | 0.1376 | 0.0372 | −2.8589 | 0.9998 | 0.9986 |
| $N = 64$ | 0.1192 | 0.0232 | 0.1172 | 0.1190 | 0.0229 | −0.8179 | 1.0000 | 1.0000 |
| $N = 128$ | 0.1095 | 0.0151 | 0.1089 | 0.1097 | 0.0151 | 0.7823 | 1.0000 | 1.0000 |
| $N = 256$ | 0.1048 | 0.0102 | 0.1041 | 0.1047 | 0.0101 | −0.5139 | 1.0000 | 1.0000 |
| $p = 16, q = 4, V(\mathbf{\Sigma}) = 0.2$ | | | | | | | | |
| $N = 4$ | 0.6333 | 0.5660 | 0.4525 | 0.6217 | 0.5592 | −1.4728 | 0.3164 | 0.2102 |
| $N = 8$ | 0.3857 | 0.2250 | 0.3395 | 0.3867 | 0.2277 | 0.3213 | 0.6996 | 0.5818 |
| $N = 16$ | 0.2867 | 0.1122 | 0.2670 | 0.2853 | 0.1118 | −0.8900 | 0.9872 | 0.9702 |
| $N = 32$ | 0.2419 | 0.0644 | 0.2345 | 0.2422 | 0.0650 | 0.2472 | 1.0000 | 1.0000 |
| $N = 64$ | 0.2206 | 0.0404 | 0.2167 | 0.2201 | 0.0406 | −1.0141 | 1.0000 | 1.0000 |
| $N = 128$ | 0.2102 | 0.0268 | 0.2086 | 0.2103 | 0.0266 | 0.0662 | 1.0000 | 1.0000 |
| $N = 256$ | 0.2051 | 0.0183 | 0.2042 | 0.2045 | 0.0179 | −2.2267 | 1.0000 | 1.0000 |

(*continued*)

**Table S1.** (*continued*)

| | E[$V$(**S**)] | SD[$V$(**S**)] | Median | Mean | ESD | $T$ | Pow. 5% | Pow. 1% |
|---|---|---|---|---|---|---|---|---|
| $p = 32, q = 1, V(\mathbf{\Sigma}) = 0.1$ | | | | | | | | |
| $N = 4$ | 0.4854 | 0.4534 | 0.3382 | 0.4789 | 0.4380 | −1.0575 | 0.2604 | 0.1900 |
| $N = 8$ | 0.2652 | 0.1932 | 0.2038 | 0.2590 | 0.1893 | −2.3033 | 0.4940 | 0.4178 |
| $N = 16$ | 0.1771 | 0.1033 | 0.1546 | 0.1794 | 0.1038 | 1.5962 | 0.8460 | 0.8034 |
| $N = 32$ | 0.1373 | 0.0628 | 0.1262 | 0.1378 | 0.0615 | 0.6141 | 0.9940 | 0.9912 |
| $N = 64$ | 0.1184 | 0.0410 | 0.1127 | 0.1186 | 0.0411 | 0.4012 | 1.0000 | 1.0000 |
| $N = 128$ | 0.1091 | 0.0278 | 0.1060 | 0.1082 | 0.0278 | −2.1710 | 1.0000 | 1.0000 |
| $N = 256$ | 0.1045 | 0.0193 | 0.1025 | 0.1040 | 0.0189 | −2.1023 | 1.0000 | 1.0000 |
| $p = 32, q = 1, V(\mathbf{\Sigma}) = 0.2$ | | | | | | | | |
| $N = 4$ | 0.6167 | 0.7641 | 0.3490 | 0.5908 | 0.7234 | −2.5279 | 0.3270 | 0.2640 |
| $N = 8$ | 0.3786 | 0.3427 | 0.2744 | 0.3799 | 0.3437 | 0.2828 | 0.6380 | 0.5822 |
| $N = 16$ | 0.2833 | 0.1905 | 0.2312 | 0.2778 | 0.1862 | −2.1149 | 0.9294 | 0.9080 |
| $N = 32$ | 0.2403 | 0.1186 | 0.2181 | 0.2423 | 0.1215 | 1.1513 | 0.9994 | 0.9980 |
| $N = 64$ | 0.2198 | 0.0786 | 0.2093 | 0.2206 | 0.0778 | 0.6652 | 1.0000 | 1.0000 |
| $N = 128$ | 0.2098 | 0.0537 | 0.2050 | 0.2107 | 0.0533 | 1.1501 | 1.0000 | 1.0000 |
| $N = 256$ | 0.2049 | 0.0374 | 0.2027 | 0.2053 | 0.0369 | 0.6690 | 1.0000 | 1.0000 |
| $p = 32, q = 1, V(\mathbf{\Sigma}) = 0.4$ | | | | | | | | |
| $N = 4$ | 0.8792 | 1.3449 | 0.4175 | 0.8790 | 1.3001 | −0.0111 | 0.4194 | 0.3640 |
| $N = 8$ | 0.6054 | 0.6280 | 0.4080 | 0.5999 | 0.6307 | −0.6167 | 0.7424 | 0.7020 |
| $N = 16$ | 0.4958 | 0.3592 | 0.4030 | 0.4931 | 0.3599 | −0.5428 | 0.9756 | 0.9694 |
| $N = 32$ | 0.4464 | 0.2278 | 0.4061 | 0.4517 | 0.2342 | 1.6000 | 0.9998 | 0.9996 |
| $N = 64$ | 0.4228 | 0.1523 | 0.4014 | 0.4228 | 0.1509 | −0.0001 | 1.0000 | 1.0000 |
| $N = 128$ | 0.4113 | 0.1047 | 0.4027 | 0.4142 | 0.1061 | 1.9386 | 1.0000 | 1.0000 |
| $N = 256$ | 0.4056 | 0.0730 | 0.3990 | 0.4049 | 0.0720 | −0.7666 | 1.0000 | 1.0000 |
| $p = 32, q = 1, V(\mathbf{\Sigma}) = 0.6$ | | | | | | | | |
| $N = 4$ | 1.1417 | 1.9016 | 0.4839 | 1.1495 | 1.8884 | 0.2917 | 0.4702 | 0.4220 |
| $N = 8$ | 0.8321 | 0.9053 | 0.5361 | 0.8252 | 0.8771 | −0.5637 | 0.8144 | 0.7848 |
| $N = 16$ | 0.7083 | 0.5248 | 0.5752 | 0.7099 | 0.5382 | 0.2084 | 0.9886 | 0.9850 |
| $N = 32$ | 0.6524 | 0.3355 | 0.5899 | 0.6548 | 0.3395 | 0.4951 | 1.0000 | 1.0000 |
| $N = 64$ | 0.6258 | 0.2254 | 0.5960 | 0.6286 | 0.2259 | 0.8767 | 1.0000 | 1.0000 |
| $N = 128$ | 0.6128 | 0.1553 | 0.5927 | 0.6096 | 0.1548 | −1.4599 | 1.0000 | 1.0000 |
| $N = 256$ | 0.6064 | 0.1084 | 0.5992 | 0.6061 | 0.1086 | −0.1815 | 1.0000 | 1.0000 |
| $p = 32, q = 1, V(\mathbf{\Sigma}) = 0.8$ | | | | | | | | |
| $N = 4$ | 1.4042 | 2.4456 | 0.5315 | 1.3706 | 2.3694 | −1.0015 | 0.4940 | 0.4506 |
| $N = 8$ | 1.0589 | 1.1784 | 0.6869 | 1.0663 | 1.1907 | 0.4355 | 0.8458 | 0.8238 |
| $N = 16$ | 0.9208 | 0.6888 | 0.7275 | 0.9046 | 0.6664 | −1.7216 | 0.9912 | 0.9894 |
| $N = 32$ | 0.8585 | 0.4424 | 0.7738 | 0.8625 | 0.4476 | 0.6339 | 1.0000 | 1.0000 |
| $N = 64$ | 0.8288 | 0.2981 | 0.7822 | 0.8223 | 0.2880 | −1.5782 | 1.0000 | 1.0000 |
| $N = 128$ | 0.8143 | 0.2057 | 0.7978 | 0.8180 | 0.2054 | 1.2902 | 1.0000 | 1.0000 |
| $N = 256$ | 0.8071 | 0.1437 | 0.7946 | 0.8046 | 0.1438 | −1.2523 | 1.0000 | 1.0000 |

(*continued*)

**Table S1.** (*continued*)

| | E[$V$(**S**)] | SD[$V$(**S**)] | Median | Mean | ESD | $T$ | Pow. 5% | Pow. 1% |
|---|---|---|---|---|---|---|---|---|
| $p = 32, q = 2, V(\boldsymbol{\Sigma}) = 0.1$ | | | | | | | | |
| $N = 4$ | 0.4854 | 0.3879 | 0.3699 | 0.4765 | 0.3649 | −1.7243 | 0.2934 | 0.2140 |
| $N = 8$ | 0.2652 | 0.1567 | 0.2254 | 0.2620 | 0.1560 | −1.4249 | 0.5638 | 0.4812 |
| $N = 16$ | 0.1771 | 0.0800 | 0.1596 | 0.1752 | 0.0797 | −1.6907 | 0.9106 | 0.8708 |
| $N = 32$ | 0.1373 | 0.0469 | 0.1300 | 0.1365 | 0.0469 | −1.1689 | 0.9994 | 0.9986 |
| $N = 64$ | 0.1184 | 0.0300 | 0.1151 | 0.1183 | 0.0298 | −0.1368 | 1.0000 | 1.0000 |
| $N = 128$ | 0.1091 | 0.0201 | 0.1071 | 0.1089 | 0.0200 | −0.6488 | 1.0000 | 1.0000 |
| $N = 256$ | 0.1045 | 0.0138 | 0.1043 | 0.1048 | 0.0138 | 1.3050 | 1.0000 | 1.0000 |
| $p = 32, q = 2, V(\boldsymbol{\Sigma}) = 0.2$ | | | | | | | | |
| $N = 4$ | 0.6167 | 0.6296 | 0.4145 | 0.6196 | 0.6367 | 0.3310 | 0.3856 | 0.3140 |
| $N = 8$ | 0.3786 | 0.2686 | 0.3172 | 0.3827 | 0.2633 | 1.1072 | 0.7300 | 0.6708 |
| $N = 16$ | 0.2833 | 0.1433 | 0.2560 | 0.2838 | 0.1414 | 0.2470 | 0.9806 | 0.9714 |
| $N = 32$ | 0.2403 | 0.0867 | 0.2278 | 0.2410 | 0.0876 | 0.5138 | 1.0000 | 1.0000 |
| $N = 64$ | 0.2198 | 0.0565 | 0.2149 | 0.2211 | 0.0577 | 1.5370 | 1.0000 | 1.0000 |
| $N = 128$ | 0.2098 | 0.0382 | 0.2064 | 0.2095 | 0.0381 | −0.5589 | 1.0000 | 1.0000 |
| $N = 256$ | 0.2049 | 0.0265 | 0.2032 | 0.2047 | 0.0261 | −0.4954 | 1.0000 | 1.0000 |
| $p = 32, q = 2, V(\boldsymbol{\Sigma}) = 0.4$ | | | | | | | | |
| $N = 4$ | 0.8792 | 1.0734 | 0.5075 | 0.8708 | 1.0767 | −0.5475 | 0.4758 | 0.4150 |
| $N = 8$ | 0.6054 | 0.4790 | 0.4807 | 0.6065 | 0.4766 | 0.1723 | 0.8488 | 0.8144 |
| $N = 16$ | 0.4958 | 0.2645 | 0.4407 | 0.4946 | 0.2639 | −0.3404 | 0.9966 | 0.9944 |
| $N = 32$ | 0.4464 | 0.1638 | 0.4192 | 0.4468 | 0.1675 | 0.1846 | 1.0000 | 1.0000 |
| $N = 64$ | 0.4228 | 0.1081 | 0.4109 | 0.4226 | 0.1077 | −0.1262 | 1.0000 | 1.0000 |
| $N = 128$ | 0.4113 | 0.0738 | 0.4060 | 0.4117 | 0.0747 | 0.3552 | 1.0000 | 1.0000 |
| $N = 256$ | 0.4056 | 0.0512 | 0.4035 | 0.4066 | 0.0512 | 1.3117 | 1.0000 | 1.0000 |
| $p = 32, q = 4, V(\boldsymbol{\Sigma}) = 0.1$ | | | | | | | | |
| $N = 4$ | 0.4854 | 0.3402 | 0.4019 | 0.4886 | 0.3388 | 0.6713 | 0.3194 | 0.2274 |
| $N = 8$ | 0.2652 | 0.1302 | 0.2360 | 0.2642 | 0.1296 | −0.5586 | 0.6138 | 0.5124 |
| $N = 16$ | 0.1771 | 0.0629 | 0.1674 | 0.1779 | 0.0631 | 0.9277 | 0.9636 | 0.9400 |
| $N = 32$ | 0.1373 | 0.0353 | 0.1328 | 0.1367 | 0.0352 | −1.1192 | 1.0000 | 1.0000 |
| $N = 64$ | 0.1184 | 0.0218 | 0.1165 | 0.1184 | 0.0220 | 0.1788 | 1.0000 | 1.0000 |
| $N = 128$ | 0.1091 | 0.0144 | 0.1083 | 0.1090 | 0.0145 | −0.2811 | 1.0000 | 1.0000 |
| $N = 256$ | 0.1045 | 0.0098 | 0.1040 | 0.1045 | 0.0097 | −0.4697 | 1.0000 | 1.0000 |
| $p = 32, q = 4, V(\boldsymbol{\Sigma}) = 0.2$ | | | | | | | | |
| $N = 4$ | 0.6167 | 0.5314 | 0.4570 | 0.6124 | 0.5398 | −0.5531 | 0.4180 | 0.3282 |
| $N = 8$ | 0.3786 | 0.2145 | 0.3322 | 0.3764 | 0.2149 | −0.6987 | 0.7890 | 0.7306 |
| $N = 16$ | 0.2833 | 0.1087 | 0.2658 | 0.2811 | 0.1065 | −1.4855 | 0.9964 | 0.9934 |
| $N = 32$ | 0.2403 | 0.0633 | 0.2333 | 0.2405 | 0.0644 | 0.2077 | 1.0000 | 1.0000 |
| $N = 64$ | 0.2198 | 0.0402 | 0.2162 | 0.2190 | 0.0398 | −1.5583 | 1.0000 | 1.0000 |
| $N = 128$ | 0.2098 | 0.0268 | 0.2084 | 0.2094 | 0.0268 | −1.0612 | 1.0000 | 1.0000 |
| $N = 256$ | 0.2049 | 0.0184 | 0.2043 | 0.2051 | 0.0184 | 0.6105 | 1.0000 | 1.0000 |

(*continued*)

**Table S1.** (*continued*)

| | E[$V$(**S**)] | SD[$V$(**S**)] | Median | Mean | ESD | $T$ | Pow. 5% | Pow. 1% |
|---|---|---|---|---|---|---|---|---|
| $p$ = 64, $q$ = 1, $V(\boldsymbol{\Sigma})$ = 0.1 | | | | | | | | |
| $N$ = 4 | 0.4760 | 0.4308 | 0.3333 | 0.4758 | 0.4454 | −0.0458 | 0.3174 | 0.2566 |
| $N$ = 8 | 0.2612 | 0.1848 | 0.2043 | 0.2593 | 0.1808 | −0.7361 | 0.5748 | 0.5076 |
| $N$ = 16 | 0.1752 | 0.0994 | 0.1502 | 0.1743 | 0.0977 | −0.6293 | 0.8884 | 0.8608 |
| $N$ = 32 | 0.1364 | 0.0606 | 0.1243 | 0.1368 | 0.0603 | 0.4427 | 0.9986 | 0.9970 |
| $N$ = 64 | 0.1179 | 0.0396 | 0.1123 | 0.1179 | 0.0407 | −0.0656 | 1.0000 | 1.0000 |
| $N$ = 128 | 0.1089 | 0.0269 | 0.1062 | 0.1088 | 0.0268 | −0.2292 | 1.0000 | 1.0000 |
| $N$ = 256 | 0.1044 | 0.0187 | 0.1031 | 0.1046 | 0.0188 | 0.7075 | 1.0000 | 1.0000 |
| $p$ = 64, $q$ = 1, $V(\boldsymbol{\Sigma})$ = 0.2 | | | | | | | | |
| $N$ = 4 | 0.6083 | 0.7426 | 0.3491 | 0.6187 | 0.7564 | 0.9699 | 0.3862 | 0.3406 |
| $N$ = 8 | 0.3750 | 0.3342 | 0.2724 | 0.3693 | 0.3084 | −1.2970 | 0.6930 | 0.6404 |
| $N$ = 16 | 0.2817 | 0.1862 | 0.2395 | 0.2846 | 0.1881 | 1.0946 | 0.9514 | 0.9416 |
| $N$ = 32 | 0.2395 | 0.1162 | 0.2189 | 0.2410 | 0.1161 | 0.8880 | 0.9998 | 0.9998 |
| $N$ = 64 | 0.2194 | 0.0770 | 0.2075 | 0.2185 | 0.0768 | −0.8362 | 1.0000 | 1.0000 |
| $N$ = 128 | 0.2096 | 0.0527 | 0.2032 | 0.2090 | 0.0534 | −0.8819 | 1.0000 | 1.0000 |
| $N$ = 256 | 0.2048 | 0.0366 | 0.2024 | 0.2051 | 0.0368 | 0.5577 | 1.0000 | 1.0000 |
| $p$ = 64, $q$ = 1, $V(\boldsymbol{\Sigma})$ = 0.4 | | | | | | | | |
| $N$ = 4 | 0.8729 | 1.3270 | 0.4193 | 0.8856 | 1.3267 | 0.6763 | 0.4664 | 0.4264 |
| $N$ = 8 | 0.6027 | 0.6206 | 0.4227 | 0.6052 | 0.5988 | 0.2997 | 0.7806 | 0.7568 |
| $N$ = 16 | 0.4946 | 0.3554 | 0.4061 | 0.5021 | 0.3631 | 1.4613 | 0.9818 | 0.9778 |
| $N$ = 32 | 0.4458 | 0.2255 | 0.4003 | 0.4445 | 0.2244 | −0.4124 | 1.0000 | 1.0000 |
| $N$ = 64 | 0.4225 | 0.1509 | 0.3986 | 0.4237 | 0.1546 | 0.5322 | 1.0000 | 1.0000 |
| $N$ = 128 | 0.4112 | 0.1038 | 0.4015 | 0.4125 | 0.1038 | 0.9057 | 1.0000 | 1.0000 |
| $N$ = 256 | 0.4056 | 0.0724 | 0.4003 | 0.4058 | 0.0718 | 0.1941 | 1.0000 | 1.0000 |
| $p$ = 64, $q$ = 1, $V(\boldsymbol{\Sigma})$ = 0.6 | | | | | | | | |
| $N$ = 4 | 1.1375 | 1.8888 | 0.4781 | 1.1311 | 1.8346 | −0.2450 | 0.5040 | 0.4696 |
| $N$ = 8 | 0.8304 | 0.8999 | 0.5503 | 0.8546 | 0.9505 | 1.8043 | 0.8352 | 0.8128 |
| $N$ = 16 | 0.7075 | 0.5220 | 0.5693 | 0.7136 | 0.5361 | 0.8002 | 0.9914 | 0.9892 |
| $N$ = 32 | 0.6520 | 0.3338 | 0.5899 | 0.6555 | 0.3387 | 0.7324 | 1.0000 | 1.0000 |
| $N$ = 64 | 0.6256 | 0.2243 | 0.5963 | 0.6275 | 0.2229 | 0.6155 | 1.0000 | 1.0000 |
| $N$ = 128 | 0.6127 | 0.1546 | 0.5961 | 0.6119 | 0.1532 | −0.3828 | 1.0000 | 1.0000 |
| $N$ = 256 | 0.6063 | 0.1079 | 0.5985 | 0.6067 | 0.1078 | 0.2554 | 1.0000 | 1.0000 |
| $p$ = 64, $q$ = 1, $V(\boldsymbol{\Sigma})$ = 0.8 | | | | | | | | |
| $N$ = 4 | 1.4021 | 2.4389 | 0.5397 | 1.3796 | 2.3986 | −0.6620 | 0.5362 | 0.5010 |
| $N$ = 8 | 1.0580 | 1.1756 | 0.6979 | 1.0755 | 1.2081 | 1.0231 | 0.8678 | 0.8528 |
| $N$ = 16 | 0.9204 | 0.6873 | 0.7463 | 0.9304 | 0.7024 | 1.0083 | 0.9940 | 0.9934 |
| $N$ = 32 | 0.8583 | 0.4415 | 0.7643 | 0.8495 | 0.4352 | −1.4315 | 1.0000 | 1.0000 |
| $N$ = 64 | 0.8287 | 0.2975 | 0.7873 | 0.8292 | 0.2977 | 0.1296 | 1.0000 | 1.0000 |
| $N$ = 128 | 0.8142 | 0.2053 | 0.7973 | 0.8192 | 0.2071 | 1.6973 | 1.0000 | 1.0000 |
| $N$ = 256 | 0.8071 | 0.1434 | 0.7976 | 0.8060 | 0.1420 | −0.5233 | 1.0000 | 1.0000 |

(*continued*)

**Table S1.** (*continued*)

| | E[$V(\mathbf{S})$] | SD[$V(\mathbf{S})$] | Median | Mean | ESD | $T$ | Pow. 5% | Pow. 1% |
|---|---|---|---|---|---|---|---|---|
| $p = 64, q = 2, V(\mathbf{\Sigma}) = 0.1$ | | | | | | | | |
| $N = 4$ | 0.4760 | 0.3647 | 0.3655 | 0.4774 | 0.3706 | 0.2548 | 0.3576 | 0.2884 |
| $N = 8$ | 0.2612 | 0.1484 | 0.2234 | 0.2611 | 0.1483 | −0.0339 | 0.6472 | 0.5736 |
| $N = 16$ | 0.1752 | 0.0763 | 0.1602 | 0.1761 | 0.0777 | 0.8173 | 0.9538 | 0.9342 |
| $N = 32$ | 0.1364 | 0.0450 | 0.1300 | 0.1364 | 0.0459 | −0.0466 | 1.0000 | 0.9998 |
| $N = 64$ | 0.1179 | 0.0288 | 0.1151 | 0.1178 | 0.0284 | −0.3278 | 1.0000 | 1.0000 |
| $N = 128$ | 0.1089 | 0.0194 | 0.1076 | 0.1090 | 0.0193 | 0.4459 | 1.0000 | 1.0000 |
| $N = 256$ | 0.1044 | 0.0133 | 0.1037 | 0.1045 | 0.0133 | 0.1905 | 1.0000 | 1.0000 |
| $p = 64, q = 2, V(\mathbf{\Sigma}) = 0.2$ | | | | | | | | |
| $N = 4$ | 0.6083 | 0.6086 | 0.4144 | 0.6043 | 0.5838 | −0.4854 | 0.4464 | 0.3862 |
| $N = 8$ | 0.3750 | 0.2610 | 0.3152 | 0.3799 | 0.2633 | 1.3251 | 0.7896 | 0.7500 |
| $N = 16$ | 0.2817 | 0.1398 | 0.2526 | 0.2804 | 0.1409 | −0.6185 | 0.9872 | 0.9804 |
| $N = 32$ | 0.2395 | 0.0849 | 0.2266 | 0.2399 | 0.0865 | 0.3471 | 1.0000 | 1.0000 |
| $N = 64$ | 0.2194 | 0.0554 | 0.2136 | 0.2187 | 0.0554 | −0.9000 | 1.0000 | 1.0000 |
| $N = 128$ | 0.2096 | 0.0376 | 0.2069 | 0.2100 | 0.0379 | 0.7484 | 1.0000 | 1.0000 |
| $N = 256$ | 0.2048 | 0.0260 | 0.2037 | 0.2047 | 0.0260 | −0.3568 | 1.0000 | 1.0000 |
| $p = 64, q = 2, V(\mathbf{\Sigma}) = 0.4$ | | | | | | | | |
| $N = 4$ | 0.8729 | 1.0582 | 0.5340 | 0.8914 | 1.0620 | 1.2328 | 0.5470 | 0.4978 |
| $N = 8$ | 0.6027 | 0.4740 | 0.4713 | 0.5962 | 0.4669 | −0.9845 | 0.8740 | 0.8518 |
| $N = 16$ | 0.4946 | 0.2625 | 0.4487 | 0.4998 | 0.2602 | 1.4063 | 0.9984 | 0.9982 |
| $N = 32$ | 0.4458 | 0.1629 | 0.4222 | 0.4444 | 0.1605 | −0.6045 | 1.0000 | 1.0000 |
| $N = 64$ | 0.4225 | 0.1076 | 0.4081 | 0.4211 | 0.1085 | −0.8938 | 1.0000 | 1.0000 |
| $N = 128$ | 0.4112 | 0.0735 | 0.4061 | 0.4104 | 0.0715 | −0.7768 | 1.0000 | 1.0000 |
| $N = 256$ | 0.4056 | 0.0511 | 0.4024 | 0.4048 | 0.0512 | −1.0341 | 1.0000 | 1.0000 |
| $p = 64, q = 4, V(\mathbf{\Sigma}) = 0.1$ | | | | | | | | |
| $N = 4$ | 0.4760 | 0.3178 | 0.3945 | 0.4743 | 0.3195 | −0.3833 | 0.3938 | 0.3076 |
| $N = 8$ | 0.2612 | 0.1226 | 0.2379 | 0.2603 | 0.1197 | −0.5099 | 0.7108 | 0.6280 |
| $N = 16$ | 0.1752 | 0.0598 | 0.1658 | 0.1756 | 0.0600 | 0.4656 | 0.9836 | 0.9720 |
| $N = 32$ | 0.1364 | 0.0338 | 0.1325 | 0.1363 | 0.0341 | −0.0890 | 1.0000 | 1.0000 |
| $N = 64$ | 0.1179 | 0.0211 | 0.1167 | 0.1182 | 0.0214 | 0.8725 | 1.0000 | 1.0000 |
| $N = 128$ | 0.1089 | 0.0139 | 0.1082 | 0.1087 | 0.0139 | −1.0247 | 1.0000 | 1.0000 |
| $N = 256$ | 0.1044 | 0.0095 | 0.1041 | 0.1044 | 0.0094 | 0.1697 | 1.0000 | 1.0000 |
| $p = 64, q = 4, V(\mathbf{\Sigma}) = 0.2$ | | | | | | | | |
| $N = 4$ | 0.6083 | 0.5135 | 0.4635 | 0.6089 | 0.5082 | 0.0738 | 0.4944 | 0.4314 |
| $N = 8$ | 0.3750 | 0.2089 | 0.3325 | 0.3748 | 0.2074 | −0.0607 | 0.8514 | 0.8052 |
| $N = 16$ | 0.2817 | 0.1067 | 0.2634 | 0.2821 | 0.1090 | 0.2833 | 0.9986 | 0.9978 |
| $N = 32$ | 0.2395 | 0.0625 | 0.2332 | 0.2399 | 0.0631 | 0.4617 | 1.0000 | 1.0000 |
| $N = 64$ | 0.2194 | 0.0398 | 0.2163 | 0.2193 | 0.0391 | −0.1922 | 1.0000 | 1.0000 |
| $N = 128$ | 0.2096 | 0.0266 | 0.2082 | 0.2097 | 0.0269 | 0.2535 | 1.0000 | 1.0000 |
| $N = 256$ | 0.2048 | 0.0183 | 0.2041 | 0.2048 | 0.0180 | −0.0932 | 1.0000 | 1.0000 |

(*continued*)

**Table S1.** (*continued*)

| | E[$V$(**S**)] | SD[$V$(**S**)] | Median | Mean | ESD | $T$ | Pow. 5% | Pow. 1% |
|---|---|---|---|---|---|---|---|---|
| $p = 128, q = 1, V(\mathbf{\Sigma}) = 0.1$ | | | | | | | | |
| $N = 4$ | 0.4714 | 0.4196 | 0.3291 | 0.4669 | 0.4247 | −0.7376 | 0.3562 | 0.3112 |
| $N = 8$ | 0.2592 | 0.1806 | 0.2074 | 0.2641 | 0.1886 | 1.8401 | 0.6340 | 0.5912 |
| $N = 16$ | 0.1743 | 0.0974 | 0.1518 | 0.1752 | 0.0980 | 0.7020 | 0.9224 | 0.9012 |
| $N = 32$ | 0.1359 | 0.0595 | 0.1265 | 0.1368 | 0.0594 | 0.9989 | 0.9992 | 0.9986 |
| $N = 64$ | 0.1177 | 0.0389 | 0.1114 | 0.1171 | 0.0390 | −0.9930 | 1.0000 | 1.0000 |
| $N = 128$ | 0.1088 | 0.0265 | 0.1056 | 0.1084 | 0.0266 | −1.1044 | 1.0000 | 1.0000 |
| $N = 256$ | 0.1044 | 0.0183 | 0.1033 | 0.1043 | 0.0182 | −0.1878 | 1.0000 | 1.0000 |
| $p = 128, q = 1, V(\mathbf{\Sigma}) = 0.2$ | | | | | | | | |
| $N = 4$ | 0.6042 | 0.7319 | 0.3521 | 0.5941 | 0.7030 | −1.0089 | 0.4288 | 0.3906 |
| $N = 8$ | 0.3732 | 0.3300 | 0.2737 | 0.3806 | 0.3437 | 1.5247 | 0.7364 | 0.7076 |
| $N = 16$ | 0.2808 | 0.1841 | 0.2350 | 0.2831 | 0.1925 | 0.8332 | 0.9602 | 0.9538 |
| $N = 32$ | 0.2391 | 0.1150 | 0.2179 | 0.2394 | 0.1158 | 0.1938 | 1.0000 | 1.0000 |
| $N = 64$ | 0.2192 | 0.0763 | 0.2096 | 0.2198 | 0.0762 | 0.5014 | 1.0000 | 1.0000 |
| $N = 128$ | 0.2095 | 0.0522 | 0.2045 | 0.2108 | 0.0529 | 1.7078 | 1.0000 | 1.0000 |
| $N = 256$ | 0.2048 | 0.0363 | 0.2012 | 0.2045 | 0.0363 | −0.5260 | 1.0000 | 1.0000 |
| $p = 128, q = 1, V(\mathbf{\Sigma}) = 0.4$ | | | | | | | | |
| $N = 4$ | 0.8698 | 1.3180 | 0.4240 | 0.8793 | 1.3211 | 0.5077 | 0.5012 | 0.4752 |
| $N = 8$ | 0.6013 | 0.6170 | 0.4109 | 0.6010 | 0.6166 | −0.0393 | 0.8032 | 0.7886 |
| $N = 16$ | 0.4940 | 0.3536 | 0.3964 | 0.4838 | 0.3447 | −2.0799 | 0.9856 | 0.9832 |
| $N = 32$ | 0.4455 | 0.2244 | 0.4018 | 0.4478 | 0.2249 | 0.7386 | 1.0000 | 1.0000 |
| $N = 64$ | 0.4224 | 0.1502 | 0.3989 | 0.4221 | 0.1513 | −0.1182 | 1.0000 | 1.0000 |
| $N = 128$ | 0.4111 | 0.1033 | 0.4000 | 0.4103 | 0.1041 | −0.5203 | 1.0000 | 1.0000 |
| $N = 256$ | 0.4055 | 0.0720 | 0.4015 | 0.4075 | 0.0718 | 1.8953 | 1.0000 | 1.0000 |
| $p = 128, q = 1, V(\mathbf{\Sigma}) = 0.6$ | | | | | | | | |
| $N = 4$ | 1.1354 | 1.8824 | 0.4797 | 1.1748 | 1.9845 | 1.4033 | 0.5288 | 0.5084 |
| $N = 8$ | 0.8295 | 0.8973 | 0.5462 | 0.8334 | 0.8762 | 0.3179 | 0.8454 | 0.8346 |
| $N = 16$ | 0.7071 | 0.5206 | 0.5797 | 0.7083 | 0.5151 | 0.1716 | 0.9932 | 0.9926 |
| $N = 32$ | 0.6518 | 0.3330 | 0.5881 | 0.6544 | 0.3402 | 0.5470 | 1.0000 | 1.0000 |
| $N = 64$ | 0.6255 | 0.2238 | 0.5970 | 0.6273 | 0.2257 | 0.5778 | 1.0000 | 1.0000 |
| $N = 128$ | 0.6126 | 0.1543 | 0.5996 | 0.6139 | 0.1541 | 0.5810 | 1.0000 | 1.0000 |
| $N = 256$ | 0.6063 | 0.1077 | 0.5977 | 0.6059 | 0.1071 | −0.2419 | 1.0000 | 1.0000 |
| $p = 128, q = 1, V(\mathbf{\Sigma}) = 0.8$ | | | | | | | | |
| $N = 4$ | 1.4010 | 2.4355 | 0.5423 | 1.3818 | 2.6633 | −0.5106 | 0.5578 | 0.5376 |
| $N = 8$ | 1.0576 | 1.1742 | 0.6488 | 1.0314 | 1.1393 | −1.6229 | 0.8726 | 0.8598 |
| $N = 16$ | 0.9202 | 0.6865 | 0.7373 | 0.9009 | 0.6625 | −2.0655 | 0.9960 | 0.9958 |
| $N = 32$ | 0.8582 | 0.4411 | 0.7650 | 0.8551 | 0.4435 | −0.4819 | 1.0000 | 1.0000 |
| $N = 64$ | 0.8286 | 0.2972 | 0.7932 | 0.8344 | 0.2987 | 1.3577 | 1.0000 | 1.0000 |
| $N = 128$ | 0.8142 | 0.2051 | 0.7948 | 0.8175 | 0.2073 | 1.1298 | 1.0000 | 1.0000 |
| $N = 256$ | 0.8071 | 0.1433 | 0.7941 | 0.8035 | 0.1427 | −1.7684 | 1.0000 | 1.0000 |

(*continued*)

**Table S1.** (*continued*)

| | E[$V$(**S**)] | SD[$V$(**S**)] | Median | Mean | ESD | $T$ | Pow. 5% | Pow. 1% |
|---|---|---|---|---|---|---|---|---|
| $p = 128, q = 2, V(\mathbf{\Sigma}) = 0.1$ | | | | | | | | |
| $N = 4$ | 0.4714 | 0.3532 | 0.3619 | 0.4658 | 0.3508 | −1.1137 | 0.4058 | 0.3562 |
| $N = 8$ | 0.2592 | 0.1443 | 0.2255 | 0.2602 | 0.1458 | 0.4873 | 0.7038 | 0.6668 |
| $N = 16$ | 0.1743 | 0.0744 | 0.1585 | 0.1736 | 0.0734 | −0.6676 | 0.9692 | 0.9588 |
| $N = 32$ | 0.1359 | 0.0440 | 0.1292 | 0.1347 | 0.0428 | −2.0088 | 1.0000 | 1.0000 |
| $N = 64$ | 0.1177 | 0.0283 | 0.1151 | 0.1181 | 0.0284 | 1.0985 | 1.0000 | 1.0000 |
| $N = 128$ | 0.1088 | 0.0190 | 0.1073 | 0.1089 | 0.0190 | 0.6346 | 1.0000 | 1.0000 |
| $N = 256$ | 0.1044 | 0.0131 | 0.1033 | 0.1043 | 0.0130 | −0.1708 | 1.0000 | 1.0000 |
| $p = 128, q = 2, V(\mathbf{\Sigma}) = 0.2$ | | | | | | | | |
| $N = 4$ | 0.6042 | 0.5982 | 0.4138 | 0.6028 | 0.6034 | −0.1628 | 0.4912 | 0.4532 |
| $N = 8$ | 0.3732 | 0.2572 | 0.3105 | 0.3757 | 0.2607 | 0.6690 | 0.8210 | 0.7964 |
| $N = 16$ | 0.2808 | 0.1381 | 0.2512 | 0.2803 | 0.1416 | −0.2466 | 0.9918 | 0.9896 |
| $N = 32$ | 0.2391 | 0.0840 | 0.2269 | 0.2387 | 0.0826 | −0.3229 | 1.0000 | 1.0000 |
| $N = 64$ | 0.2192 | 0.0549 | 0.2138 | 0.2191 | 0.0556 | −0.2106 | 1.0000 | 1.0000 |
| $N = 128$ | 0.2095 | 0.0372 | 0.2063 | 0.2095 | 0.0378 | −0.0391 | 1.0000 | 1.0000 |
| $N = 256$ | 0.2048 | 0.0258 | 0.2036 | 0.2047 | 0.0259 | −0.1840 | 1.0000 | 1.0000 |
| $p = 128, q = 2, V(\mathbf{\Sigma}) = 0.4$ | | | | | | | | |
| $N = 4$ | 0.8698 | 1.0506 | 0.5457 | 0.8847 | 1.0750 | 0.9798 | 0.5790 | 0.5540 |
| $N = 8$ | 0.6013 | 0.4714 | 0.4772 | 0.6031 | 0.4732 | 0.2588 | 0.9008 | 0.8864 |
| $N = 16$ | 0.4940 | 0.2615 | 0.4404 | 0.4875 | 0.2552 | −1.7810 | 0.9990 | 0.9988 |
| $N = 32$ | 0.4455 | 0.1625 | 0.4232 | 0.4457 | 0.1604 | 0.0840 | 1.0000 | 1.0000 |
| $N = 64$ | 0.4224 | 0.1074 | 0.4136 | 0.4238 | 0.1074 | 0.9228 | 1.0000 | 1.0000 |
| $N = 128$ | 0.4111 | 0.0734 | 0.4042 | 0.4099 | 0.0734 | −1.1445 | 1.0000 | 1.0000 |
| $N = 256$ | 0.4055 | 0.0510 | 0.4026 | 0.4058 | 0.0508 | 0.3393 | 1.0000 | 1.0000 |
| $p = 128, q = 4, V(\mathbf{\Sigma}) = 0.1$ | | | | | | | | |
| $N = 4$ | 0.4714 | 0.3065 | 0.3956 | 0.4754 | 0.3052 | 0.9466 | 0.4594 | 0.4060 |
| $N = 8$ | 0.2592 | 0.1187 | 0.2303 | 0.2571 | 0.1193 | −1.2327 | 0.7618 | 0.7164 |
| $N = 16$ | 0.1743 | 0.0582 | 0.1659 | 0.1748 | 0.0586 | 0.6611 | 0.9918 | 0.9884 |
| $N = 32$ | 0.1359 | 0.0331 | 0.1314 | 0.1355 | 0.0331 | −0.9953 | 1.0000 | 1.0000 |
| $N = 64$ | 0.1177 | 0.0206 | 0.1160 | 0.1174 | 0.0208 | −1.0099 | 1.0000 | 1.0000 |
| $N = 128$ | 0.1088 | 0.0137 | 0.1078 | 0.1087 | 0.0136 | −0.3031 | 1.0000 | 1.0000 |
| $N = 256$ | 0.1044 | 0.0093 | 0.1040 | 0.1041 | 0.0093 | −1.8770 | 1.0000 | 1.0000 |
| $p = 128, q = 4, V(\mathbf{\Sigma}) = 0.2$ | | | | | | | | |
| $N = 4$ | 0.6042 | 0.5044 | 0.4654 | 0.5986 | 0.4922 | −0.8044 | 0.5486 | 0.5040 |
| $N = 8$ | 0.3732 | 0.2059 | 0.3269 | 0.3715 | 0.2029 | −0.5889 | 0.8868 | 0.8642 |
| $N = 16$ | 0.2808 | 0.1056 | 0.2666 | 0.2834 | 0.1064 | 1.7057 | 0.9996 | 0.9988 |
| $N = 32$ | 0.2391 | 0.0621 | 0.2328 | 0.2400 | 0.0629 | 1.0371 | 1.0000 | 1.0000 |
| $N = 64$ | 0.2192 | 0.0396 | 0.2166 | 0.2192 | 0.0400 | −0.0044 | 1.0000 | 1.0000 |
| $N = 128$ | 0.2095 | 0.0265 | 0.2077 | 0.2093 | 0.0261 | −0.7459 | 1.0000 | 1.0000 |
| $N = 256$ | 0.2048 | 0.0183 | 0.2044 | 0.2049 | 0.0183 | 0.5881 | 1.0000 | 1.0000 |

(*continued*)

**Table S1.** (*continued*)

| | E[$V$(**S**)] | SD[$V$(**S**)] | Median | Mean | ESD | $T$ | Pow. 5% | Pow. 1% |
|---|---|---|---|---|---|---|---|---|
| $p$ = 256, $q$ = 1, $V(\mathbf{\Sigma})$ = 0.1 | | | | | | | | |
| $N$ = 4 | 0.4690 | 0.4140 | 0.3288 | 0.4640 | 0.3993 | −0.8845 | 0.3936 | 0.3548 |
| $N$ = 8 | 0.2581 | 0.1786 | 0.1992 | 0.2538 | 0.1758 | −1.7463 | 0.6480 | 0.6122 |
| $N$ = 16 | 0.1738 | 0.0964 | 0.1501 | 0.1722 | 0.0929 | −1.2156 | 0.9370 | 0.9300 |
| $N$ = 32 | 0.1357 | 0.0589 | 0.1226 | 0.1338 | 0.0571 | −2.3990 | 0.9986 | 0.9984 |
| $N$ = 64 | 0.1176 | 0.0386 | 0.1122 | 0.1177 | 0.0384 | 0.1675 | 1.0000 | 1.0000 |
| $N$ = 128 | 0.1087 | 0.0262 | 0.1063 | 0.1086 | 0.0261 | −0.1824 | 1.0000 | 1.0000 |
| $N$ = 256 | 0.1043 | 0.0182 | 0.1027 | 0.1043 | 0.0184 | −0.2755 | 1.0000 | 1.0000 |
| $p$ = 256, $q$ = 1, $V(\mathbf{\Sigma})$ = 0.2 | | | | | | | | |
| $N$ = 4 | 0.6021 | 0.7265 | 0.3493 | 0.5926 | 0.7138 | −0.9406 | 0.4574 | 0.4288 |
| $N$ = 8 | 0.3723 | 0.3279 | 0.2676 | 0.3679 | 0.3236 | −0.9750 | 0.7410 | 0.7150 |
| $N$ = 16 | 0.2804 | 0.1831 | 0.2332 | 0.2806 | 0.1813 | 0.0870 | 0.9712 | 0.9664 |
| $N$ = 32 | 0.2389 | 0.1144 | 0.2166 | 0.2377 | 0.1138 | −0.7437 | 1.0000 | 1.0000 |
| $N$ = 64 | 0.2191 | 0.0759 | 0.2104 | 0.2196 | 0.0772 | 0.4276 | 1.0000 | 1.0000 |
| $N$ = 128 | 0.2095 | 0.0519 | 0.2059 | 0.2104 | 0.0518 | 1.2576 | 1.0000 | 1.0000 |
| $N$ = 256 | 0.2047 | 0.0361 | 0.2017 | 0.2050 | 0.0361 | 0.5110 | 1.0000 | 1.0000 |
| $p$ = 256, $q$ = 1, $V(\mathbf{\Sigma})$ = 0.4 | | | | | | | | |
| $N$ = 4 | 0.8682 | 1.3136 | 0.4188 | 0.8545 | 1.2792 | −0.7569 | 0.5160 | 0.4972 |
| $N$ = 8 | 0.6007 | 0.6151 | 0.4140 | 0.6076 | 0.6137 | 0.7995 | 0.8272 | 0.8158 |
| $N$ = 16 | 0.4936 | 0.3526 | 0.4000 | 0.4873 | 0.3492 | −1.2751 | 0.9882 | 0.9872 |
| $N$ = 32 | 0.4453 | 0.2239 | 0.4068 | 0.4465 | 0.2213 | 0.3813 | 1.0000 | 1.0000 |
| $N$ = 64 | 0.4223 | 0.1499 | 0.4003 | 0.4234 | 0.1505 | 0.5054 | 1.0000 | 1.0000 |
| $N$ = 128 | 0.4111 | 0.1031 | 0.3965 | 0.4095 | 0.1027 | −1.0603 | 1.0000 | 1.0000 |
| $N$ = 256 | 0.4055 | 0.0719 | 0.4012 | 0.4060 | 0.0719 | 0.4926 | 1.0000 | 1.0000 |
| $p$ = 256, $q$ = 1, $V(\mathbf{\Sigma})$ = 0.6 | | | | | | | | |
| $N$ = 4 | 1.1344 | 1.8792 | 0.4711 | 1.0961 | 1.7882 | −1.5150 | 0.5510 | 0.5306 |
| $N$ = 8 | 0.8290 | 0.8960 | 0.5433 | 0.8354 | 0.8985 | 0.5049 | 0.8694 | 0.8588 |
| $N$ = 16 | 0.7069 | 0.5200 | 0.5747 | 0.7078 | 0.5192 | 0.1261 | 0.9946 | 0.9936 |
| $N$ = 32 | 0.6517 | 0.3326 | 0.5949 | 0.6581 | 0.3348 | 1.3577 | 1.0000 | 1.0000 |
| $N$ = 64 | 0.6254 | 0.2235 | 0.5884 | 0.6207 | 0.2239 | −1.4937 | 1.0000 | 1.0000 |
| $N$ = 128 | 0.6126 | 0.1541 | 0.5974 | 0.6123 | 0.1544 | −0.1619 | 1.0000 | 1.0000 |
| $N$ = 256 | 0.6063 | 0.1076 | 0.5994 | 0.6061 | 0.1066 | −0.1095 | 1.0000 | 1.0000 |
| $p$ = 256, $q$ = 1, $V(\mathbf{\Sigma})$ = 0.8 | | | | | | | | |
| $N$ = 4 | 1.4005 | 2.4338 | 0.5288 | 1.4172 | 2.5212 | 0.4676 | 0.5650 | 0.5524 |
| $N$ = 8 | 1.0574 | 1.1735 | 0.6982 | 1.0818 | 1.1935 | 1.4492 | 0.8902 | 0.8828 |
| $N$ = 16 | 0.9201 | 0.6862 | 0.7373 | 0.9122 | 0.6746 | −0.8321 | 0.9964 | 0.9964 |
| $N$ = 32 | 0.8581 | 0.4409 | 0.7749 | 0.8663 | 0.4540 | 1.2689 | 1.0000 | 1.0000 |
| $N$ = 64 | 0.8286 | 0.2971 | 0.7824 | 0.8219 | 0.2873 | −1.6582 | 1.0000 | 1.0000 |
| $N$ = 128 | 0.8142 | 0.2050 | 0.7977 | 0.8154 | 0.2033 | 0.4365 | 1.0000 | 1.0000 |
| $N$ = 256 | 0.8071 | 0.1432 | 0.7974 | 0.8086 | 0.1460 | 0.7277 | 1.0000 | 1.0000 |

(*continued*)

**Table S1.** (*continued*)

| | E[$V$(**S**)] | SD[$V$(**S**)] | Median | Mean | ESD | $T$ | Pow. 5% | Pow. 1% |
|---|---|---|---|---|---|---|---|---|
| $p$ = 256, $q$ = 2, $V(\boldsymbol{\Sigma})$ = 0.1 | | | | | | | | |
| $N$ = 4 | 0.4690 | 0.3474 | 0.3654 | 0.4634 | 0.3480 | −1.1409 | 0.4556 | 0.4024 |
| $N$ = 8 | 0.2581 | 0.1423 | 0.2253 | 0.2595 | 0.1427 | 0.6890 | 0.7438 | 0.7116 |
| $N$ = 16 | 0.1738 | 0.0735 | 0.1578 | 0.1718 | 0.0736 | −1.9007 | 0.9722 | 0.9690 |
| $N$ = 32 | 0.1357 | 0.0435 | 0.1283 | 0.1348 | 0.0426 | −1.5449 | 1.0000 | 1.0000 |
| $N$ = 64 | 0.1176 | 0.0280 | 0.1144 | 0.1177 | 0.0281 | 0.2848 | 1.0000 | 1.0000 |
| $N$ = 128 | 0.1087 | 0.0188 | 0.1069 | 0.1084 | 0.0189 | −1.0010 | 1.0000 | 1.0000 |
| $N$ = 256 | 0.1043 | 0.0130 | 0.1040 | 0.1046 | 0.0131 | 1.4744 | 1.0000 | 1.0000 |
| $p$ = 256, $q$ = 2, $V(\boldsymbol{\Sigma})$ = 0.2 | | | | | | | | |
| $N$ = 4 | 0.6021 | 0.5930 | 0.4144 | 0.5971 | 0.6015 | −0.5910 | 0.5248 | 0.4916 |
| $N$ = 8 | 0.3723 | 0.2553 | 0.3056 | 0.3701 | 0.2520 | −0.6132 | 0.8336 | 0.8142 |
| $N$ = 16 | 0.2804 | 0.1372 | 0.2539 | 0.2790 | 0.1348 | −0.7403 | 0.9926 | 0.9912 |
| $N$ = 32 | 0.2389 | 0.0836 | 0.2263 | 0.2379 | 0.0827 | −0.8512 | 1.0000 | 1.0000 |
| $N$ = 64 | 0.2191 | 0.0546 | 0.2131 | 0.2187 | 0.0547 | −0.6298 | 1.0000 | 1.0000 |
| $N$ = 128 | 0.2095 | 0.0370 | 0.2066 | 0.2100 | 0.0367 | 1.0076 | 1.0000 | 1.0000 |
| $N$ = 256 | 0.2047 | 0.0256 | 0.2033 | 0.2046 | 0.0255 | −0.4003 | 1.0000 | 1.0000 |
| $p$ = 256, $q$ = 2, $V(\boldsymbol{\Sigma})$ = 0.4 | | | | | | | | |
| $N$ = 4 | 0.8682 | 1.0468 | 0.5365 | 0.8927 | 1.0995 | 1.5729 | 0.6016 | 0.5770 |
| $N$ = 8 | 0.6007 | 0.4701 | 0.4639 | 0.5909 | 0.4642 | −1.4846 | 0.9042 | 0.8960 |
| $N$ = 16 | 0.4936 | 0.2609 | 0.4474 | 0.5025 | 0.2686 | 2.3407 | 0.9996 | 0.9996 |
| $N$ = 32 | 0.4453 | 0.1622 | 0.4253 | 0.4481 | 0.1594 | 1.2589 | 1.0000 | 1.0000 |
| $N$ = 64 | 0.4223 | 0.1073 | 0.4107 | 0.4221 | 0.1070 | −0.1076 | 1.0000 | 1.0000 |
| $N$ = 128 | 0.4111 | 0.0733 | 0.4038 | 0.4095 | 0.0729 | −1.5460 | 1.0000 | 1.0000 |
| $N$ = 256 | 0.4055 | 0.0509 | 0.4043 | 0.4063 | 0.0516 | 1.1106 | 1.0000 | 1.0000 |
| $p$ = 256, $q$ = 4, $V(\boldsymbol{\Sigma})$ = 0.1 | | | | | | | | |
| $N$ = 4 | 0.4690 | 0.3008 | 0.3970 | 0.4698 | 0.2910 | 0.1839 | 0.5062 | 0.4560 |
| $N$ = 8 | 0.2581 | 0.1168 | 0.2346 | 0.2563 | 0.1142 | −1.1614 | 0.8074 | 0.7734 |
| $N$ = 16 | 0.1738 | 0.0574 | 0.1645 | 0.1723 | 0.0557 | −1.9201 | 0.9940 | 0.9924 |
| $N$ = 32 | 0.1357 | 0.0327 | 0.1315 | 0.1352 | 0.0326 | −1.1484 | 1.0000 | 1.0000 |
| $N$ = 64 | 0.1176 | 0.0204 | 0.1162 | 0.1178 | 0.0206 | 0.8992 | 1.0000 | 1.0000 |
| $N$ = 128 | 0.1087 | 0.0135 | 0.1082 | 0.1087 | 0.0134 | −0.1494 | 1.0000 | 1.0000 |
| $N$ = 256 | 0.1043 | 0.0092 | 0.1041 | 0.1045 | 0.0092 | 1.2177 | 1.0000 | 1.0000 |
| $p$ = 256, $q$ = 4, $V(\boldsymbol{\Sigma})$ = 0.2 | | | | | | | | |
| $N$ = 4 | 0.6021 | 0.4998 | 0.4605 | 0.5996 | 0.4947 | −0.3569 | 0.5792 | 0.5418 |
| $N$ = 8 | 0.3723 | 0.2044 | 0.3280 | 0.3694 | 0.2072 | −0.9800 | 0.8912 | 0.8774 |
| $N$ = 16 | 0.2804 | 0.1050 | 0.2652 | 0.2818 | 0.1068 | 0.9360 | 0.9994 | 0.9994 |
| $N$ = 32 | 0.2389 | 0.0618 | 0.2323 | 0.2377 | 0.0614 | −1.4261 | 1.0000 | 1.0000 |
| $N$ = 64 | 0.2191 | 0.0395 | 0.2166 | 0.2189 | 0.0395 | −0.4363 | 1.0000 | 1.0000 |
| $N$ = 128 | 0.2095 | 0.0265 | 0.2082 | 0.2099 | 0.0270 | 1.1301 | 1.0000 | 1.0000 |
| $N$ = 256 | 0.2047 | 0.0182 | 0.2040 | 0.2048 | 0.0180 | 0.1575 | 1.0000 | 1.0000 |

(*continued*)

**Table S1.** (*continued*)

| | E[$V$(**S**)] | SD[$V$(**S**)] | Median | Mean | ESD | $T$ | Pow. 5% | Pow. 1% |
|---|---|---|---|---|---|---|---|---|
| $p$ = 1024, $q$ = 1, $V(\boldsymbol{\Sigma})$ = 0.1 | | | | | | | | |
| $N$ = 4 | 0.4673 | 0.4098 | 0.3340 | 0.4614 | 0.3938 | −1.0472 | 0.4516 | 0.4324 |
| $N$ = 8 | 0.2574 | 0.1770 | 0.2071 | 0.2598 | 0.1802 | 0.9277 | 0.7062 | 0.6892 |
| $N$ = 16 | 0.1735 | 0.0957 | 0.1506 | 0.1731 | 0.0955 | −0.2362 | 0.9464 | 0.9400 |
| $N$ = 32 | 0.1355 | 0.0585 | 0.1229 | 0.1345 | 0.0579 | −1.2252 | 1.0000 | 1.0000 |
| $N$ = 64 | 0.1175 | 0.0383 | 0.1119 | 0.1176 | 0.0385 | 0.1237 | 1.0000 | 1.0000 |
| $N$ = 128 | 0.1087 | 0.0261 | 0.1056 | 0.1081 | 0.0258 | −1.5172 | 1.0000 | 1.0000 |
| $N$ = 256 | 0.1043 | 0.0181 | 0.1028 | 0.1041 | 0.0181 | −1.0113 | 1.0000 | 1.0000 |
| $p$ = 1024, $q$ = 1, $V(\boldsymbol{\Sigma})$ = 0.2 | | | | | | | | |
| $N$ = 4 | 0.6005 | 0.7225 | 0.3486 | 0.6004 | 0.7467 | −0.0149 | 0.4836 | 0.4722 |
| $N$ = 8 | 0.3717 | 0.3263 | 0.2718 | 0.3697 | 0.3275 | −0.4127 | 0.7766 | 0.7646 |
| $N$ = 16 | 0.2801 | 0.1823 | 0.2393 | 0.2820 | 0.1793 | 0.7472 | 0.9752 | 0.9740 |
| $N$ = 32 | 0.2388 | 0.1139 | 0.2163 | 0.2375 | 0.1138 | −0.7599 | 1.0000 | 1.0000 |
| $N$ = 64 | 0.2191 | 0.0756 | 0.2098 | 0.2201 | 0.0761 | 0.9238 | 1.0000 | 1.0000 |
| $N$ = 128 | 0.2095 | 0.0517 | 0.2029 | 0.2079 | 0.0514 | −2.1703 | 1.0000 | 1.0000 |
| $N$ = 256 | 0.2047 | 0.0360 | 0.2010 | 0.2036 | 0.0356 | −2.2748 | 1.0000 | 1.0000 |
| $p$ = 1024, $q$ = 1, $V(\boldsymbol{\Sigma})$ = 0.4 | | | | | | | | |
| $N$ = 4 | 0.8671 | 1.3102 | 0.4015 | 0.8405 | 1.3164 | −1.4242 | 0.5364 | 0.5234 |
| $N$ = 8 | 0.6002 | 0.6138 | 0.3961 | 0.5940 | 0.6361 | −0.6814 | 0.8298 | 0.8208 |
| $N$ = 16 | 0.4934 | 0.3519 | 0.4015 | 0.4977 | 0.3638 | 0.8432 | 0.9866 | 0.9860 |
| $N$ = 32 | 0.4452 | 0.2235 | 0.4037 | 0.4489 | 0.2304 | 1.1498 | 1.0000 | 1.0000 |
| $N$ = 64 | 0.4222 | 0.1496 | 0.4020 | 0.4240 | 0.1526 | 0.8140 | 1.0000 | 1.0000 |
| $N$ = 128 | 0.4110 | 0.1029 | 0.3979 | 0.4104 | 0.1038 | −0.4592 | 1.0000 | 1.0000 |
| $N$ = 256 | 0.4055 | 0.0717 | 0.4020 | 0.4063 | 0.0728 | 0.7559 | 1.0000 | 1.0000 |
| $p$ = 1024, $q$ = 1, $V(\boldsymbol{\Sigma})$ = 0.6 | | | | | | | | |
| $N$ = 4 | 1.1336 | 1.8768 | 0.4611 | 1.1261 | 1.9068 | −0.2763 | 0.5542 | 0.5472 |
| $N$ = 8 | 0.8287 | 0.8950 | 0.5476 | 0.8168 | 0.8639 | −0.9740 | 0.8732 | 0.8690 |
| $N$ = 16 | 0.7067 | 0.5194 | 0.5792 | 0.7003 | 0.4944 | −0.9163 | 0.9940 | 0.9938 |
| $N$ = 32 | 0.6516 | 0.3323 | 0.5888 | 0.6511 | 0.3351 | −0.1071 | 1.0000 | 1.0000 |
| $N$ = 64 | 0.6254 | 0.2233 | 0.5952 | 0.6228 | 0.2197 | −0.8267 | 1.0000 | 1.0000 |
| $N$ = 128 | 0.6126 | 0.1539 | 0.5969 | 0.6127 | 0.1505 | 0.0278 | 1.0000 | 1.0000 |
| $N$ = 256 | 0.6063 | 0.1075 | 0.5971 | 0.6056 | 0.1075 | −0.4205 | 1.0000 | 1.0000 |
| $p$ = 1024, $q$ = 1, $V(\boldsymbol{\Sigma})$ = 0.8 | | | | | | | | |
| $N$ = 4 | 1.4001 | 2.4326 | 0.5395 | 1.3638 | 2.3850 | −1.0764 | 0.5938 | 0.5872 |
| $N$ = 8 | 1.0572 | 1.1729 | 0.6881 | 1.0573 | 1.1723 | 0.0072 | 0.8892 | 0.8864 |
| $N$ = 16 | 0.9200 | 0.6859 | 0.7655 | 0.9344 | 0.6943 | 1.4624 | 0.9964 | 0.9962 |
| $N$ = 32 | 0.8581 | 0.4407 | 0.7709 | 0.8531 | 0.4414 | −0.7945 | 1.0000 | 1.0000 |
| $N$ = 64 | 0.8286 | 0.2969 | 0.7840 | 0.8292 | 0.2987 | 0.1364 | 1.0000 | 1.0000 |
| $N$ = 128 | 0.8142 | 0.2049 | 0.7958 | 0.8135 | 0.2091 | −0.2323 | 1.0000 | 1.0000 |
| $N$ = 256 | 0.8071 | 0.1432 | 0.7897 | 0.8021 | 0.1457 | −2.4090 | 1.0000 | 1.0000 |

(*continued*)

**Table S1.** (*continued*)

| | E[$V$(**S**)] | SD[$V$(**S**)] | Median | Mean | ESD | $T$ | Pow. 5% | Pow. 1% |
|---|---|---|---|---|---|---|---|---|
| $p = 1024$, $q = 2$, $V(\mathbf{\Sigma}) = 0.1$ | | | | | | | | |
| $N = 4$ | 0.4673 | 0.3431 | 0.3680 | 0.4677 | 0.3375 | 0.0847 | 0.5096 | 0.4900 |
| $N = 8$ | 0.2574 | 0.1407 | 0.2234 | 0.2551 | 0.1380 | −1.1672 | 0.7816 | 0.7650 |
| $N = 16$ | 0.1735 | 0.0728 | 0.1589 | 0.1725 | 0.0728 | −0.8882 | 0.9808 | 0.9794 |
| $N = 32$ | 0.1355 | 0.0432 | 0.1290 | 0.1351 | 0.0430 | −0.7093 | 1.0000 | 1.0000 |
| $N = 64$ | 0.1175 | 0.0277 | 0.1154 | 0.1181 | 0.0280 | 1.5804 | 1.0000 | 1.0000 |
| $N = 128$ | 0.1087 | 0.0187 | 0.1075 | 0.1088 | 0.0185 | 0.4350 | 1.0000 | 1.0000 |
| $N = 256$ | 0.1043 | 0.0129 | 0.1037 | 0.1043 | 0.0129 | 0.1578 | 1.0000 | 1.0000 |
| $p = 1024$, $q = 2$, $V(\mathbf{\Sigma}) = 0.2$ | | | | | | | | |
| $N = 4$ | 0.6005 | 0.5891 | 0.4114 | 0.5972 | 0.6001 | −0.3908 | 0.5562 | 0.5428 |
| $N = 8$ | 0.3717 | 0.2538 | 0.3100 | 0.3740 | 0.2543 | 0.6557 | 0.8622 | 0.8536 |
| $N = 16$ | 0.2801 | 0.1366 | 0.2497 | 0.2786 | 0.1375 | −0.7571 | 0.9954 | 0.9954 |
| $N = 32$ | 0.2388 | 0.0832 | 0.2278 | 0.2396 | 0.0843 | 0.7147 | 1.0000 | 1.0000 |
| $N = 64$ | 0.2191 | 0.0544 | 0.2124 | 0.2181 | 0.0540 | −1.2282 | 1.0000 | 1.0000 |
| $N = 128$ | 0.2095 | 0.0369 | 0.2071 | 0.2092 | 0.0363 | −0.4385 | 1.0000 | 1.0000 |
| $N = 256$ | 0.2047 | 0.0256 | 0.2042 | 0.2050 | 0.0255 | 0.7656 | 1.0000 | 1.0000 |
| $p = 1024$, $q = 2$, $V(\mathbf{\Sigma}) = 0.4$ | | | | | | | | |
| $N = 4$ | 0.8671 | 1.0439 | 0.5247 | 0.8745 | 1.0769 | 0.4858 | 0.6186 | 0.6108 |
| $N = 8$ | 0.6002 | 0.4691 | 0.4631 | 0.5869 | 0.4547 | −2.0640 | 0.9268 | 0.9202 |
| $N = 16$ | 0.4934 | 0.2605 | 0.4428 | 0.4976 | 0.2683 | 1.0984 | 0.9992 | 0.9990 |
| $N = 32$ | 0.4452 | 0.1620 | 0.4194 | 0.4427 | 0.1615 | −1.0754 | 1.0000 | 1.0000 |
| $N = 64$ | 0.4222 | 0.1072 | 0.4127 | 0.4236 | 0.1082 | 0.8702 | 1.0000 | 1.0000 |
| $N = 128$ | 0.4110 | 0.0732 | 0.4088 | 0.4135 | 0.0733 | 2.4063 | 1.0000 | 1.0000 |
| $N = 256$ | 0.4055 | 0.0509 | 0.4024 | 0.4058 | 0.0506 | 0.3761 | 1.0000 | 1.0000 |
| $p = 1024$, $q = 4$, $V(\mathbf{\Sigma}) = 0.1$ | | | | | | | | |
| $N = 4$ | 0.4673 | 0.2966 | 0.3913 | 0.4672 | 0.2960 | −0.0035 | 0.5506 | 0.5286 |
| $N = 8$ | 0.2574 | 0.1153 | 0.2361 | 0.2596 | 0.1170 | 1.3608 | 0.8494 | 0.8328 |
| $N = 16$ | 0.1735 | 0.0567 | 0.1645 | 0.1741 | 0.0584 | 0.8447 | 0.9966 | 0.9960 |
| $N = 32$ | 0.1355 | 0.0324 | 0.1313 | 0.1349 | 0.0329 | −1.2922 | 1.0000 | 1.0000 |
| $N = 64$ | 0.1175 | 0.0203 | 0.1157 | 0.1168 | 0.0200 | −2.4767 | 1.0000 | 1.0000 |
| $N = 128$ | 0.1087 | 0.0134 | 0.1082 | 0.1088 | 0.0136 | 0.8393 | 1.0000 | 1.0000 |
| $N = 256$ | 0.1043 | 0.0092 | 0.1040 | 0.1043 | 0.0090 | 0.1824 | 1.0000 | 1.0000 |
| $p = 1024$, $q = 4$, $V(\mathbf{\Sigma}) = 0.2$ | | | | | | | | |
| $N = 4$ | 0.6005 | 0.4964 | 0.4682 | 0.5980 | 0.4856 | −0.3698 | 0.6212 | 0.6076 |
| $N = 8$ | 0.3717 | 0.2033 | 0.3306 | 0.3706 | 0.2005 | −0.3631 | 0.9174 | 0.9120 |
| $N = 16$ | 0.2801 | 0.1046 | 0.2681 | 0.2846 | 0.1044 | 3.0183 | 1.0000 | 1.0000 |
| $N = 32$ | 0.2388 | 0.0616 | 0.2315 | 0.2376 | 0.0602 | −1.3527 | 1.0000 | 1.0000 |
| $N = 64$ | 0.2191 | 0.0394 | 0.2156 | 0.2182 | 0.0389 | −1.6629 | 1.0000 | 1.0000 |
| $N = 128$ | 0.2095 | 0.0264 | 0.2076 | 0.2089 | 0.0261 | −1.5712 | 1.0000 | 1.0000 |
| $N = 256$ | 0.2047 | 0.0182 | 0.2038 | 0.2045 | 0.0181 | −0.9541 | 1.0000 | 1.0000 |

(*continued*)

**Table S1.** (*continued*)

| | E[$V$(**S**)] | SD[$V$(**S**)] | Median | Mean | ESD | $T$ | Pow. 5% | Pow. 1% |
|---|---|---|---|---|---|---|---|---|
| $p$ = 2, linearly decreasing λ ($V(\boldsymbol{\Sigma})$ = 0.1111) | | | | | | | | |
| $N$ = 4 | 0.7778 | 1.3168 | 0.3244 | 0.7933 | 1.3342 | 0.8238 | 0.0708 | 0.0190 |
| $N$ = 8 | 0.3968 | 0.5318 | 0.2164 | 0.3997 | 0.5313 | 0.3812 | 0.1042 | 0.0328 |
| $N$ = 16 | 0.2444 | 0.2693 | 0.1575 | 0.2433 | 0.2687 | −0.2967 | 0.1820 | 0.0676 |
| $N$ = 32 | 0.1756 | 0.1565 | 0.1352 | 0.1795 | 0.1599 | 1.7054 | 0.3436 | 0.1446 |
| $N$ = 64 | 0.1429 | 0.0992 | 0.1232 | 0.1434 | 0.1000 | 0.4129 | 0.6262 | 0.4204 |
| $N$ = 128 | 0.1269 | 0.0661 | 0.1169 | 0.1281 | 0.0676 | 1.3481 | 0.9248 | 0.8128 |
| $N$ = 256 | 0.1190 | 0.0453 | 0.1140 | 0.1192 | 0.0455 | 0.4317 | 0.9980 | 0.9892 |
| $p$ = 4, linearly decreasing λ ($V(\boldsymbol{\Sigma})$ = 0.0667) | | | | | | | | |
| $N$ = 4 | 0.5778 | 0.6569 | 0.3679 | 0.5751 | 0.6531 | −0.2908 | 0.0824 | 0.0210 |
| $N$ = 8 | 0.2857 | 0.2434 | 0.2159 | 0.2877 | 0.2528 | 0.5523 | 0.1362 | 0.0496 |
| $N$ = 16 | 0.1689 | 0.1133 | 0.1381 | 0.1670 | 0.1128 | −1.1736 | 0.2488 | 0.0998 |
| $N$ = 32 | 0.1161 | 0.0613 | 0.1008 | 0.1140 | 0.0603 | −2.5412 | 0.5314 | 0.2710 |
| $N$ = 64 | 0.0910 | 0.0369 | 0.0855 | 0.0917 | 0.0374 | 1.2880 | 0.9250 | 0.8034 |
| $N$ = 128 | 0.0787 | 0.0238 | 0.0754 | 0.0786 | 0.0239 | −0.5522 | 1.0000 | 0.9984 |
| $N$ = 256 | 0.0727 | 0.0160 | 0.0717 | 0.0729 | 0.0160 | 0.7927 | 1.0000 | 1.0000 |
| $p$ = 8, linearly decreasing λ ($V(\boldsymbol{\Sigma})$ = 0.0370) | | | | | | | | |
| $N$ = 4 | 0.4630 | 0.3521 | 0.3683 | 0.4628 | 0.3535 | −0.0264 | 0.0898 | 0.0322 |
| $N$ = 8 | 0.2196 | 0.1211 | 0.1903 | 0.2180 | 0.1228 | −0.8922 | 0.1508 | 0.0504 |
| $N$ = 16 | 0.1222 | 0.0520 | 0.1125 | 0.1220 | 0.0520 | −0.2921 | 0.3076 | 0.1458 |
| $N$ = 32 | 0.0783 | 0.0261 | 0.0748 | 0.0785 | 0.0264 | 0.6822 | 0.6982 | 0.4684 |
| $N$ = 64 | 0.0573 | 0.0148 | 0.0558 | 0.0574 | 0.0148 | 0.4841 | 0.9930 | 0.9622 |
| $N$ = 128 | 0.0471 | 0.0091 | 0.0461 | 0.0469 | 0.0090 | −1.5428 | 1.0000 | 1.0000 |
| $N$ = 256 | 0.0420 | 0.0060 | 0.0416 | 0.0421 | 0.0060 | 1.1109 | 1.0000 | 1.0000 |
| $p$ = 16, linearly decreasing λ ($V(\boldsymbol{\Sigma})$ = 0.0196) | | | | | | | | |
| $N$ = 4 | 0.4003 | 0.2049 | 0.3565 | 0.4022 | 0.2110 | 0.6379 | 0.0916 | 0.0268 |
| $N$ = 8 | 0.1828 | 0.0658 | 0.1722 | 0.1821 | 0.0656 | −0.6784 | 0.1430 | 0.0518 |
| $N$ = 16 | 0.0958 | 0.0260 | 0.0936 | 0.0964 | 0.0263 | 1.8197 | 0.3258 | 0.1462 |
| $N$ = 32 | 0.0565 | 0.0120 | 0.0552 | 0.0564 | 0.0121 | −0.0814 | 0.7550 | 0.5230 |
| $N$ = 64 | 0.0377 | 0.0063 | 0.0373 | 0.0378 | 0.0064 | 0.4353 | 0.9976 | 0.9880 |
| $N$ = 128 | 0.0286 | 0.0036 | 0.0284 | 0.0286 | 0.0037 | −0.5174 | 1.0000 | 1.0000 |
| $N$ = 256 | 0.0241 | 0.0023 | 0.0240 | 0.0241 | 0.0022 | 0.9463 | 1.0000 | 1.0000 |
| $p$ = 32, linearly decreasing λ ($V(\boldsymbol{\Sigma})$ = 0.0101) | | | | | | | | |
| $N$ = 4 | 0.3674 | 0.1284 | 0.3498 | 0.3676 | 0.1275 | 0.0762 | 0.0890 | 0.0322 |
| $N$ = 8 | 0.1632 | 0.0391 | 0.1574 | 0.1611 | 0.0386 | −3.8961 | 0.1182 | 0.0470 |
| $N$ = 16 | 0.0816 | 0.0143 | 0.0803 | 0.0814 | 0.0141 | −0.7872 | 0.2772 | 0.1328 |
| $N$ = 32 | 0.0447 | 0.0060 | 0.0443 | 0.0447 | 0.0061 | 0.2237 | 0.7446 | 0.5416 |
| $N$ = 64 | 0.0271 | 0.0029 | 0.0270 | 0.0271 | 0.0029 | 0.3384 | 0.9986 | 0.9954 |
| $N$ = 128 | 0.0185 | 0.0015 | 0.0185 | 0.0185 | 0.0015 | 0.2761 | 1.0000 | 1.0000 |
| $N$ = 256 | 0.0143 | 0.0009 | 0.0143 | 0.0143 | 0.0009 | 0.0449 | 1.0000 | 1.0000 |

(*continued*)

**Table S1.** (*continued*)

| | E[$V$(**S**)] | SD[$V$(**S**)] | Median | Mean | ESD | $T$ | Pow. 5% | Pow. 1% |
|---|---|---|---|---|---|---|---|---|
| $p$ = 64, linearly decreasing λ ($V(\mathbf{\Sigma})$ = 0.0051) | | | | | | | | |
| $N$ = 4 | 0.3505 | 0.08474 | 0.3397 | 0.3483 | 0.08321 | −1.8765 | 0.0926 | 0.0218 |
| $N$ = 8 | 0.1532 | 0.02489 | 0.1512 | 0.1531 | 0.02482 | −0.1999 | 0.1152 | 0.0330 |
| $N$ = 16 | 0.0742 | 0.00863 | 0.0738 | 0.0743 | 0.00855 | 0.5331 | 0.2424 | 0.0954 |
| $N$ = 32 | 0.0386 | 0.00335 | 0.0384 | 0.0386 | 0.00337 | 0.1913 | 0.6256 | 0.4168 |
| $N$ = 64 | 0.0216 | 0.00145 | 0.0215 | 0.0216 | 0.00144 | 0.5035 | 0.9970 | 0.9890 |
| $N$ = 128 | 0.0133 | 0.00070 | 0.0133 | 0.0133 | 0.00071 | −0.1560 | 1.0000 | 1.0000 |
| $N$ = 256 | 0.0092 | 0.00038 | 0.0092 | 0.0092 | 0.00037 | −2.0535 | 1.0000 | 1.0000 |
| $p$ = 128, linearly decreasing λ ($V(\mathbf{\Sigma})$ = 0.0026) | | | | | | | | |
| $N$ = 4 | 0.3420 | 0.05775 | 0.3379 | 0.3416 | 0.05769 | −0.4209 | 0.0894 | 0.0274 |
| $N$ = 8 | 0.1480 | 0.01660 | 0.1471 | 0.1478 | 0.01665 | −0.8164 | 0.1096 | 0.0346 |
| $N$ = 16 | 0.0705 | 0.00554 | 0.0704 | 0.0705 | 0.00558 | −0.0138 | 0.1858 | 0.0718 |
| $N$ = 32 | 0.0354 | 0.00203 | 0.0354 | 0.0354 | 0.00203 | 0.4273 | 0.4964 | 0.2800 |
| $N$ = 64 | 0.0187 | 0.00081 | 0.0187 | 0.0188 | 0.00081 | 0.6380 | 0.9844 | 0.9502 |
| $N$ = 128 | 0.0106 | 0.00036 | 0.0106 | 0.0106 | 0.00035 | 0.2818 | 1.0000 | 1.0000 |
| $N$ = 256 | 0.0066 | 0.00017 | 0.0066 | 0.0066 | 0.00017 | 1.8720 | 1.0000 | 1.0000 |
| $p$ = 256, linearly decreasing λ ($V(\mathbf{\Sigma})$ = 0.0013) | | | | | | | | |
| $N$ = 4 | 0.3377 | 0.04006 | 0.3359 | 0.3382 | 0.04042 | 0.8696 | 0.0924 | 0.0236 |
| $N$ = 8 | 0.1455 | 0.01138 | 0.1452 | 0.1455 | 0.01124 | 0.4174 | 0.0908 | 0.0208 |
| $N$ = 16 | 0.0686 | 0.00372 | 0.0685 | 0.0686 | 0.00372 | 0.2241 | 0.1694 | 0.0666 |
| $N$ = 32 | 0.0338 | 0.00131 | 0.0338 | 0.0338 | 0.00131 | −0.6163 | 0.3578 | 0.1798 |
| $N$ = 64 | 0.0173 | 0.00049 | 0.0173 | 0.0173 | 0.00050 | 1.3107 | 0.9134 | 0.7978 |
| $N$ = 128 | 0.0092 | 0.00020 | 0.0092 | 0.0092 | 0.00020 | −0.1534 | 1.0000 | 1.0000 |
| $N$ = 256 | 0.0053 | 0.00009 | 0.0053 | 0.0053 | 0.00009 | −0.1959 | 1.0000 | 1.0000 |
| $p$ = 1024, linearly decreasing λ ($V(\mathbf{\Sigma})$ = 0.0003) | | | | | | | | |
| $N$ = 4 | 0.3344 | 0.01974 | 0.3339 | 0.3345 | 0.01985 | 0.2764 | 0.0848 | 0.0270 |
| $N$ = 8 | 0.1435 | 0.00556 | 0.1434 | 0.1435 | 0.00552 | −0.5480 | 0.0880 | 0.0190 |
| $N$ = 16 | 0.0671 | 0.00178 | 0.0671 | 0.0671 | 0.00178 | −0.0478 | 0.1052 | 0.0338 |
| $N$ = 32 | 0.0327 | 0.00061 | 0.0327 | 0.0327 | 0.00062 | 0.5148 | 0.2036 | 0.0790 |
| $N$ = 64 | 0.0162 | 0.00021 | 0.0162 | 0.0162 | 0.00021 | 0.3321 | 0.5512 | 0.3070 |
| $N$ = 128 | 0.0082 | 0.00008 | 0.0082 | 0.0082 | 0.00008 | 1.8861 | 0.9988 | 0.9906 |
| $N$ = 256 | 0.0043 | 0.00003 | 0.0043 | 0.0043 | 0.00003 | 0.8650 | 1.0000 | 1.0000 |

(*continued*)

**Table S1.** (*continued*)

| | E[$V$(**S**)] | SD[$V$(**S**)] | Median | Mean | ESD | $T$ | Pow. 5% | Pow. 1% |
|---|---|---|---|---|---|---|---|---|
| $p$ = 2, quadratically decreasing λ ($V$(**Σ**) = 0.3600) | | | | | | | | |
| $N$ = 4 | 1.0267 | 1.8585 | 0.3865 | 0.9886 | 1.7577 | −1.5298 | 0.1000 | 0.0320 |
| $N$ = 8 | 0.6457 | 0.8290 | 0.3649 | 0.6528 | 0.8393 | 0.5941 | 0.2108 | 0.0958 |
| $N$ = 16 | 0.4933 | 0.4583 | 0.3551 | 0.4902 | 0.4570 | −0.4788 | 0.4410 | 0.2418 |
| $N$ = 32 | 0.4245 | 0.2842 | 0.3563 | 0.4239 | 0.2886 | −0.1479 | 0.7944 | 0.5390 |
| $N$ = 64 | 0.3917 | 0.1877 | 0.3601 | 0.3932 | 0.1889 | 0.5256 | 0.9884 | 0.9542 |
| $N$ = 128 | 0.3757 | 0.1282 | 0.3611 | 0.3769 | 0.1279 | 0.6613 | 1.0000 | 0.9998 |
| $N$ = 256 | 0.3678 | 0.0890 | 0.3567 | 0.3669 | 0.0896 | −0.7454 | 1.0000 | 1.0000 |
| $p$ = 4, quadratically decreasing λ ($V$(**Σ**) = 0.1911) | | | | | | | | |
| $N$ = 4 | 0.7230 | 0.9730 | 0.4085 | 0.7294 | 0.9809 | 0.4633 | 0.1306 | 0.0548 |
| $N$ = 8 | 0.4190 | 0.4031 | 0.3059 | 0.4178 | 0.3900 | −0.2228 | 0.2756 | 0.1264 |
| $N$ = 16 | 0.2975 | 0.2094 | 0.2416 | 0.2960 | 0.2056 | −0.5254 | 0.5702 | 0.3578 |
| $N$ = 32 | 0.2426 | 0.1242 | 0.2203 | 0.2436 | 0.1233 | 0.5682 | 0.9398 | 0.8156 |
| $N$ = 64 | 0.2164 | 0.0798 | 0.2046 | 0.2155 | 0.0785 | −0.8868 | 1.0000 | 0.9996 |
| $N$ = 128 | 0.2037 | 0.0537 | 0.1980 | 0.2042 | 0.0534 | 0.6874 | 1.0000 | 1.0000 |
| $N$ = 256 | 0.1974 | 0.0370 | 0.1945 | 0.1974 | 0.0368 | 0.1120 | 1.0000 | 1.0000 |
| $p$ = 8, quadratically decreasing λ ($V$(**Σ**) = 0.0980) | | | | | | | | |
| $N$ = 4 | 0.5392 | 0.4970 | 0.3844 | 0.5303 | 0.4841 | −1.3072 | 0.1458 | 0.0692 |
| $N$ = 8 | 0.2871 | 0.1893 | 0.2402 | 0.2887 | 0.1957 | 0.5796 | 0.3016 | 0.1590 |
| $N$ = 16 | 0.1863 | 0.0910 | 0.1663 | 0.1856 | 0.0920 | −0.5266 | 0.6552 | 0.4694 |
| $N$ = 32 | 0.1407 | 0.0508 | 0.1312 | 0.1399 | 0.0506 | −1.1144 | 0.9838 | 0.9450 |
| $N$ = 64 | 0.1190 | 0.0313 | 0.1150 | 0.1190 | 0.0317 | −0.1024 | 1.0000 | 1.0000 |
| $N$ = 128 | 0.1085 | 0.0205 | 0.1064 | 0.1085 | 0.0206 | −0.0111 | 1.0000 | 1.0000 |
| $N$ = 256 | 0.1032 | 0.0139 | 0.1020 | 0.1030 | 0.0140 | −0.9112 | 1.0000 | 1.0000 |
| $p$ = 16, quadratically decreasing λ ($V$(**Σ**) = 0.0496) | | | | | | | | |
| $N$ = 4 | 0.4390 | 0.2711 | 0.3719 | 0.4354 | 0.2697 | −0.9575 | 0.1392 | 0.0572 |
| $N$ = 8 | 0.2165 | 0.0946 | 0.1975 | 0.2151 | 0.0922 | −1.0611 | 0.2892 | 0.1544 |
| $N$ = 16 | 0.1274 | 0.0415 | 0.1209 | 0.1270 | 0.0412 | −0.8072 | 0.6622 | 0.4782 |
| $N$ = 32 | 0.0872 | 0.0213 | 0.0847 | 0.0874 | 0.0217 | 0.5232 | 0.9916 | 0.9652 |
| $N$ = 64 | 0.0681 | 0.0123 | 0.0670 | 0.0682 | 0.0124 | 0.6871 | 1.0000 | 1.0000 |
| $N$ = 128 | 0.0588 | 0.0078 | 0.0580 | 0.0586 | 0.0077 | −1.3435 | 1.0000 | 1.0000 |
| $N$ = 256 | 0.0541 | 0.0052 | 0.0540 | 0.0542 | 0.0052 | 1.1584 | 1.0000 | 1.0000 |
| $p$ = 32, quadratically decreasing λ ($V$(**Σ**) = 0.0249) | | | | | | | | |
| $N$ = 4 | 0.3868 | 0.1611 | 0.3537 | 0.3840 | 0.1566 | −1.2819 | 0.1400 | 0.0626 |
| $N$ = 8 | 0.1800 | 0.0520 | 0.1740 | 0.1803 | 0.0518 | 0.3846 | 0.2702 | 0.1556 |
| $N$ = 16 | 0.0973 | 0.0207 | 0.0950 | 0.0971 | 0.0207 | −0.7694 | 0.6166 | 0.4606 |
| $N$ = 32 | 0.0599 | 0.0097 | 0.0594 | 0.0601 | 0.0099 | 0.9872 | 0.9894 | 0.9688 |
| $N$ = 64 | 0.0421 | 0.0051 | 0.0418 | 0.0421 | 0.0051 | 0.0932 | 1.0000 | 1.0000 |
| $N$ = 128 | 0.0334 | 0.0030 | 0.0333 | 0.0334 | 0.0031 | −0.4273 | 1.0000 | 1.0000 |
| $N$ = 256 | 0.0292 | 0.0019 | 0.0291 | 0.0292 | 0.0019 | 1.2425 | 1.0000 | 1.0000 |

(*continued*)

**Table S1.** (*continued*)

| | E[$V$(**S**)] | SD[$V$(**S**)] | Median | Mean | ESD | $T$ | Pow. 5% | Pow. 1% |
|---|---|---|---|---|---|---|---|---|
| $p$ = 64, quadratically decreasing λ ($V$(**Σ**) = 0.0125) | | | | | | | | |
| $N$ = 4 | 0.3603 | 0.10274 | 0.3469 | 0.3599 | 0.10569 | −0.2366 | 0.1464 | 0.0638 |
| $N$ = 8 | 0.1615 | 0.03129 | 0.1581 | 0.1616 | 0.03125 | 0.2612 | 0.2324 | 0.1032 |
| $N$ = 16 | 0.0820 | 0.01150 | 0.0815 | 0.0826 | 0.01166 | 3.4198 | 0.5504 | 0.3684 |
| $N$ = 32 | 0.0461 | 0.00486 | 0.0458 | 0.0460 | 0.00484 | −1.3132 | 0.9774 | 0.9342 |
| $N$ = 64 | 0.0290 | 0.00233 | 0.0290 | 0.0291 | 0.00236 | 1.9871 | 1.0000 | 1.0000 |
| $N$ = 128 | 0.0207 | 0.00126 | 0.0206 | 0.0207 | 0.00126 | −2.1347 | 1.0000 | 1.0000 |
| $N$ = 256 | 0.0166 | 0.00075 | 0.0165 | 0.0166 | 0.00075 | 0.4776 | 1.0000 | 1.0000 |
| $p$ = 128, quadratically decreasing λ ($V$(**Σ**) = 0.0062) | | | | | | | | |
| $N$ = 4 | 0.3468 | 0.06862 | 0.3414 | 0.3476 | 0.06800 | 0.8011 | 0.1350 | 0.0620 |
| $N$ = 8 | 0.1522 | 0.02014 | 0.1508 | 0.1520 | 0.02010 | −0.6510 | 0.2016 | 0.0944 |
| $N$ = 16 | 0.0744 | 0.00697 | 0.0738 | 0.0742 | 0.00692 | −1.6078 | 0.4236 | 0.2468 |
| $N$ = 32 | 0.0392 | 0.00271 | 0.0392 | 0.0392 | 0.00272 | 0.1739 | 0.9186 | 0.8312 |
| $N$ = 64 | 0.0225 | 0.00118 | 0.0224 | 0.0224 | 0.00118 | −1.0233 | 1.0000 | 1.0000 |
| $N$ = 128 | 0.0143 | 0.00057 | 0.0143 | 0.0143 | 0.00058 | 0.4957 | 1.0000 | 1.0000 |
| $N$ = 256 | 0.0103 | 0.00031 | 0.0102 | 0.0102 | 0.00032 | −0.7049 | 1.0000 | 1.0000 |
| $p$ = 256, quadratically decreasing λ ($V$(**Σ**) = 0.0031) | | | | | | | | |
| $N$ = 4 | 0.3401 | 0.04709 | 0.3378 | 0.3400 | 0.04713 | −0.1581 | 0.1338 | 0.0460 |
| $N$ = 8 | 0.1475 | 0.01353 | 0.1469 | 0.1474 | 0.01375 | −0.7367 | 0.1654 | 0.0648 |
| $N$ = 16 | 0.0705 | 0.00451 | 0.0703 | 0.0705 | 0.00454 | −0.5367 | 0.3418 | 0.1896 |
| $N$ = 32 | 0.0357 | 0.00165 | 0.0357 | 0.0357 | 0.00164 | −0.2453 | 0.8128 | 0.6602 |
| $N$ = 64 | 0.0192 | 0.00066 | 0.0191 | 0.0191 | 0.00066 | −2.5120 | 1.0000 | 1.0000 |
| $N$ = 128 | 0.0111 | 0.00029 | 0.0111 | 0.0111 | 0.00029 | −3.0020 | 1.0000 | 1.0000 |
| $N$ = 256 | 0.0071 | 0.00014 | 0.0071 | 0.0071 | 0.00014 | −1.4241 | 1.0000 | 1.0000 |
| $p$ = 1024, quadratically decreasing λ ($V$(**Σ**) = 0.0008) | | | | | | | | |
| $N$ = 4 | 0.3350 | 0.02300 | 0.3346 | 0.3352 | 0.02282 | 0.4279 | 0.1146 | 0.0486 |
| $N$ = 8 | 0.1440 | 0.00649 | 0.1439 | 0.1440 | 0.00650 | −0.0425 | 0.1416 | 0.0490 |
| $N$ = 16 | 0.0676 | 0.00210 | 0.0676 | 0.0677 | 0.00210 | 0.8367 | 0.2058 | 0.0988 |
| $N$ = 32 | 0.0331 | 0.00072 | 0.0331 | 0.0331 | 0.00071 | 0.7721 | 0.4686 | 0.2736 |
| $N$ = 64 | 0.0167 | 0.00026 | 0.0167 | 0.0167 | 0.00027 | −1.3843 | 0.9656 | 0.9110 |
| $N$ = 128 | 0.0087 | 0.00010 | 0.0087 | 0.0087 | 0.00010 | −1.8267 | 1.0000 | 1.0000 |
| $N$ = 256 | 0.0047 | 0.00004 | 0.0047 | 0.0047 | 0.00004 | 0.4628 | 1.0000 | 1.0000 |

**Table S2.** Summary of simulation results for $V_{rel}(\mathbf{S})$. Theoretical expectation ($E[V_{rel}(\mathbf{S})]$) and standard deviation ($SD[V_{rel}(\mathbf{S})]$; approximate results are shown for these in non-null conditions), as well as empirical median, mean, standard deviation (ESD), bias in standard error unit ($T$), and critical points or power (for null and non-null conditions, respectively) at α = 0.05 and 0.01 (CP/Pow. 5% and 1%) from 5000 simulation runs are shown. See Table S1 for further information.

| | $E[V_{rel}(\mathbf{S})]$ | $SD[V_{rel}(\mathbf{S})]$ | Median | Mean | ESD | $T$ | CP 5% | CP 1% |
|---|---|---|---|---|---|---|---|---|
| $p = 2$, $V_{rel}(\mathbf{\Sigma}) = 0$ | | | | | | | | |
| $N = 4$ | 0.5000 | 0.2887 | 0.5073 | 0.5039 | 0.2885 | 0.9603 | 0.9500 | 0.9897 |
| $N = 8$ | 0.2500 | 0.1936 | 0.2079 | 0.2514 | 0.1924 | 0.4961 | 0.6335 | 0.7686 |
| $N = 16$ | 0.1250 | 0.1102 | 0.0953 | 0.1259 | 0.1098 | 0.6000 | 0.3454 | 0.4862 |
| $N = 32$ | 0.0625 | 0.0587 | 0.0448 | 0.0628 | 0.0597 | 0.3234 | 0.1830 | 0.2744 |
| $N = 64$ | 0.0313 | 0.0303 | 0.0224 | 0.0309 | 0.0289 | −0.8301 | 0.0893 | 0.1266 |
| $N = 128$ | 0.0156 | 0.0154 | 0.0109 | 0.0154 | 0.0149 | −1.1474 | 0.0455 | 0.0646 |
| $N = 256$ | 0.0078 | 0.0078 | 0.0055 | 0.0079 | 0.0078 | 0.5516 | 0.0232 | 0.0351 |
| $p = 4$, $V_{rel}(\mathbf{\Sigma}) = 0$ | | | | | | | | |
| $N = 4$ | 0.4286 | 0.1506 | 0.4028 | 0.4293 | 0.1509 | 0.3344 | 0.7168 | 0.8275 |
| $N = 8$ | 0.2000 | 0.0840 | 0.1856 | 0.2002 | 0.0858 | 0.1644 | 0.3641 | 0.4610 |
| $N = 16$ | 0.0968 | 0.0433 | 0.0907 | 0.0968 | 0.0434 | 0.0620 | 0.1774 | 0.2307 |
| $N = 32$ | 0.0476 | 0.0219 | 0.0438 | 0.0474 | 0.0221 | −0.6146 | 0.0885 | 0.1159 |
| $N = 64$ | 0.0236 | 0.0110 | 0.0218 | 0.0237 | 0.0110 | 0.6544 | 0.0442 | 0.0558 |
| $N = 128$ | 0.0118 | 0.0055 | 0.0109 | 0.0118 | 0.0056 | 0.1782 | 0.0225 | 0.0287 |
| $N = 256$ | 0.0059 | 0.0028 | 0.0054 | 0.0058 | 0.0027 | −1.1927 | 0.0107 | 0.0138 |
| $p = 8$, $V_{rel}(\mathbf{\Sigma}) = 0$ | | | | | | | | |
| $N = 4$ | 0.3846 | 0.0803 | 0.3732 | 0.3863 | 0.0811 | 1.5112 | 0.5406 | 0.6279 |
| $N = 8$ | 0.1724 | 0.0397 | 0.1685 | 0.1725 | 0.0390 | 0.2005 | 0.2434 | 0.2820 |
| $N = 16$ | 0.0820 | 0.0193 | 0.0802 | 0.0821 | 0.0190 | 0.5489 | 0.1163 | 0.1340 |
| $N = 32$ | 0.0400 | 0.0095 | 0.0390 | 0.0401 | 0.0096 | 0.7661 | 0.0574 | 0.0668 |
| $N = 64$ | 0.0198 | 0.0047 | 0.0193 | 0.0198 | 0.0047 | 0.2662 | 0.0281 | 0.0324 |
| $N = 128$ | 0.0098 | 0.0023 | 0.0096 | 0.0098 | 0.0024 | −0.4846 | 0.0140 | 0.0163 |
| $N = 256$ | 0.0049 | 0.0012 | 0.0048 | 0.0049 | 0.0012 | 1.1240 | 0.0070 | 0.0082 |
| $p = 16$, $V_{rel}(\mathbf{\Sigma}) = 0$ | | | | | | | | |
| $N = 4$ | 0.3600 | 0.0418 | 0.3532 | 0.3601 | 0.0412 | 0.2428 | 0.4385 | 0.4926 |
| $N = 8$ | 0.1579 | 0.0193 | 0.1562 | 0.1580 | 0.0197 | 0.3764 | 0.1934 | 0.2135 |
| $N = 16$ | 0.0744 | 0.0091 | 0.0738 | 0.0746 | 0.0094 | 1.4158 | 0.0911 | 0.0991 |
| $N = 32$ | 0.0361 | 0.0044 | 0.0357 | 0.0361 | 0.0044 | −1.2604 | 0.0439 | 0.0477 |
| $N = 64$ | 0.0178 | 0.0022 | 0.0177 | 0.0178 | 0.0022 | 0.4338 | 0.0215 | 0.0234 |
| $N = 128$ | 0.0088 | 0.0011 | 0.0088 | 0.0088 | 0.0011 | −1.2039 | 0.0107 | 0.0116 |
| $N = 256$ | 0.0044 | 0.0005 | 0.0044 | 0.0044 | 0.0005 | 1.8858 | 0.0053 | 0.0058 |
| $p = 32$, $V_{rel}(\mathbf{\Sigma}) = 0$ | | | | | | | | |
| $N = 4$ | 0.3469 | 0.0214 | 0.3429 | 0.3468 | 0.0212 | −0.4007 | 0.3872 | 0.4132 |
| $N = 8$ | 0.1504 | 0.0095 | 0.1492 | 0.1503 | 0.0096 | −0.8935 | 0.1680 | 0.1777 |
| $N = 16$ | 0.0705 | 0.0044 | 0.0701 | 0.0704 | 0.0044 | −1.6332 | 0.0780 | 0.0823 |
| $N = 32$ | 0.0342 | 0.0021 | 0.0341 | 0.0342 | 0.0021 | −0.7810 | 0.0378 | 0.0397 |
| $N = 64$ | 0.0168 | 0.0010 | 0.0168 | 0.0168 | 0.0010 | −0.7681 | 0.0186 | 0.0195 |
| $N = 128$ | 0.0084 | 0.0005 | 0.0083 | 0.0084 | 0.0005 | 0.0999 | 0.0093 | 0.0096 |
| $N = 256$ | 0.0042 | 0.0003 | 0.0042 | 0.0042 | 0.0003 | 0.1845 | 0.0046 | 0.0048 |

(*continued*)

**Table S2.** (*continued*)

| | $E[V_{rel}(\mathbf{S})]$ | $SD[V_{rel}(\mathbf{S})]$ | Median | Mean | ESD | $T$ | CP 5% | CP 1% |
|---|---|---|---|---|---|---|---|---|
| $p = 64$, $V_{rel}(\mathbf{\Sigma}) = 0$ | | | | | | | | |
| $N = 4$ | 0.3402 | 0.01084 | 0.3381 | 0.3403 | 0.01099 | 0.3923 | 0.3616 | 0.3763 |
| $N = 8$ | 0.1467 | 0.00472 | 0.1462 | 0.1466 | 0.00464 | −0.6909 | 0.1549 | 0.1594 |
| $N = 16$ | 0.0686 | 0.00218 | 0.0685 | 0.0686 | 0.00216 | 0.5618 | 0.0723 | 0.0741 |
| $N = 32$ | 0.0332 | 0.00104 | 0.0332 | 0.0332 | 0.00106 | 0.0446 | 0.0350 | 0.0359 |
| $N = 64$ | 0.0164 | 0.00051 | 0.0163 | 0.0164 | 0.00050 | −0.5869 | 0.0172 | 0.0176 |
| $N = 128$ | 0.0081 | 0.00025 | 0.0081 | 0.0081 | 0.00025 | 0.6710 | 0.0085 | 0.0087 |
| $N = 256$ | 0.0040 | 0.00013 | 0.0040 | 0.0040 | 0.00013 | 1.8656 | 0.0043 | 0.0044 |
| $p = 128$, $V_{rel}(\mathbf{\Sigma}) = 0$ | | | | | | | | |
| $N = 4$ | 0.3368 | 0.00545 | 0.3358 | 0.3368 | 0.00545 | 0.1266 | 0.3471 | 0.3539 |
| $N = 8$ | 0.1448 | 0.00235 | 0.1446 | 0.1447 | 0.00230 | −0.0173 | 0.1488 | 0.1509 |
| $N = 16$ | 0.0676 | 0.00108 | 0.0676 | 0.0677 | 0.00107 | 1.4545 | 0.0695 | 0.0704 |
| $N = 32$ | 0.0327 | 0.00052 | 0.0327 | 0.0327 | 0.00052 | −0.7481 | 0.0336 | 0.0340 |
| $N = 64$ | 0.0161 | 0.00025 | 0.0161 | 0.0161 | 0.00025 | 0.3608 | 0.0165 | 0.0167 |
| $N = 128$ | 0.0080 | 0.00012 | 0.0080 | 0.0080 | 0.00012 | 0.8117 | 0.0082 | 0.0083 |
| $N = 256$ | 0.0040 | 0.00006 | 0.0040 | 0.0040 | 0.00006 | 0.5283 | 0.0041 | 0.0041 |
| $p = 256$, $V_{rel}(\mathbf{\Sigma}) = 0$ | | | | | | | | |
| $N = 4$ | 0.3351 | 0.00274 | 0.3345 | 0.3351 | 0.00275 | 0.9603 | 0.3403 | 0.3437 |
| $N = 8$ | 0.1438 | 0.00117 | 0.1437 | 0.1438 | 0.00118 | 0.4961 | 0.1459 | 0.1472 |
| $N = 16$ | 0.0672 | 0.00054 | 0.0671 | 0.0672 | 0.00053 | 0.6000 | 0.0681 | 0.0685 |
| $N = 32$ | 0.0325 | 0.00026 | 0.0325 | 0.0325 | 0.00025 | 0.3234 | 0.0329 | 0.0331 |
| $N = 64$ | 0.0160 | 0.00013 | 0.0160 | 0.0160 | 0.00013 | −0.8301 | 0.0162 | 0.0163 |
| $N = 128$ | 0.0079 | 0.00006 | 0.0079 | 0.0079 | 0.00006 | −1.1474 | 0.0080 | 0.0081 |
| $N = 256$ | 0.0040 | 0.00003 | 0.0040 | 0.0040 | 0.00003 | 0.5516 | 0.0040 | 0.0040 |
| $p = 1024$, $V_{rel}(\mathbf{\Sigma}) = 0$ | | | | | | | | |
| $N = 4$ | 0.3338 | 0.00069 | 0.3336 | 0.3338 | 0.00070 | 1.2938 | 0.3351 | 0.3360 |
| $N = 8$ | 0.1431 | 0.00029 | 0.1431 | 0.1431 | 0.00029 | −0.5105 | 0.1436 | 0.1439 |
| $N = 16$ | 0.0668 | 0.00013 | 0.0668 | 0.0668 | 0.00013 | 0.0231 | 0.0670 | 0.0671 |
| $N = 32$ | 0.0323 | 0.00006 | 0.0323 | 0.0323 | 0.00006 | −1.3055 | 0.0324 | 0.0325 |
| $N = 64$ | 0.0159 | 0.00003 | 0.0159 | 0.0159 | 0.00003 | −0.0022 | 0.0160 | 0.0160 |
| $N = 128$ | 0.0079 | 0.00002 | 0.0079 | 0.0079 | 0.00002 | 0.3824 | 0.0079 | 0.0079 |
| $N = 256$ | 0.0039 | 0.00001 | 0.0039 | 0.0039 | 0.00001 | −0.3392 | 0.0039 | 0.0039 |

(*continued*)

**Table S2.** (*continued*)

| | $\approx E[V_{rel}(\mathbf{S})]$ | $\approx SD[V_{rel}(\mathbf{S})]$ | Median | Mean | ESD | $T$ | Pow. 5% | Pow. 1% |
|---|---|---|---|---|---|---|---|---|
| $p = 2, q = 1, V_{rel}(\mathbf{\Sigma}) = 0.1$ | | | | | | | | |
| $N = 4$ | 0.4469 | 0.4633 | 0.5363 | 0.5248 | 0.2904 | 18.9688 | 0.0586 | 0.0122 |
| $N = 8$ | 0.2782 | 0.2935 | 0.2711 | 0.3061 | 0.2126 | 9.2675 | 0.0882 | 0.0254 |
| $N = 16$ | 0.1920 | 0.1811 | 0.1640 | 0.1932 | 0.1428 | 0.5873 | 0.1592 | 0.0406 |
| $N = 32$ | 0.1469 | 0.1157 | 0.1328 | 0.1505 | 0.1028 | 2.4955 | 0.3320 | 0.1290 |
| $N = 64$ | 0.1237 | 0.0768 | 0.1132 | 0.1238 | 0.0710 | 0.0973 | 0.6406 | 0.4366 |
| $N = 128$ | 0.1119 | 0.0523 | 0.1063 | 0.1112 | 0.0507 | −1.0195 | 0.9184 | 0.8118 |
| $N = 256$ | 0.1060 | 0.0363 | 0.1051 | 0.1070 | 0.0355 | 2.1216 | 0.9982 | 0.9914 |
| $p = 2, q = 1, V_{rel}(\mathbf{\Sigma}) = 0.2$ | | | | | | | | |
| $N = 4$ | 0.4290 | 0.4462 | 0.5869 | 0.5583 | 0.2857 | 31.9985 | 0.0660 | 0.0140 |
| $N = 8$ | 0.3197 | 0.3082 | 0.3399 | 0.3586 | 0.2254 | 12.1908 | 0.1356 | 0.0464 |
| $N = 16$ | 0.2642 | 0.2030 | 0.2591 | 0.2743 | 0.1661 | 4.3048 | 0.3274 | 0.1210 |
| $N = 32$ | 0.2335 | 0.1359 | 0.2264 | 0.2368 | 0.1214 | 1.9558 | 0.6386 | 0.3574 |
| $N = 64$ | 0.2171 | 0.0930 | 0.2153 | 0.2182 | 0.0883 | 0.9017 | 0.9382 | 0.8402 |
| $N = 128$ | 0.2087 | 0.0645 | 0.2060 | 0.2088 | 0.0622 | 0.2154 | 0.9994 | 0.9958 |
| $N = 256$ | 0.2044 | 0.0452 | 0.2041 | 0.2051 | 0.0442 | 1.1662 | 1.0000 | 1.0000 |
| $p = 2, q = 1, V_{rel}(\mathbf{\Sigma}) = 0.4$ | | | | | | | | |
| $N = 4$ | 0.4756 | 0.3691 | 0.6779 | 0.6252 | 0.2745 | 38.5387 | 0.0958 | 0.0184 |
| $N = 8$ | 0.4377 | 0.2825 | 0.4939 | 0.4804 | 0.2334 | 12.9356 | 0.2916 | 0.1148 |
| $N = 16$ | 0.4232 | 0.1982 | 0.4457 | 0.4384 | 0.1764 | 6.0670 | 0.6988 | 0.4114 |
| $N = 32$ | 0.4132 | 0.1378 | 0.4169 | 0.4152 | 0.1297 | 1.1224 | 0.9592 | 0.8522 |
| $N = 64$ | 0.4070 | 0.0963 | 0.4108 | 0.4083 | 0.0945 | 0.9214 | 1.0000 | 0.9990 |
| $N = 128$ | 0.4036 | 0.0676 | 0.4051 | 0.4042 | 0.0661 | 0.6227 | 1.0000 | 1.0000 |
| $N = 256$ | 0.4018 | 0.0476 | 0.4019 | 0.4014 | 0.0479 | −0.6817 | 1.0000 | 1.0000 |
| $p = 2, q = 1, V_{rel}(\mathbf{\Sigma}) = 0.6$ | | | | | | | | |
| $N = 4$ | 0.6025 | 0.2596 | 0.7743 | 0.7012 | 0.2530 | 27.5684 | 0.1468 | 0.0342 |
| $N = 8$ | 0.5952 | 0.2126 | 0.6625 | 0.6262 | 0.2094 | 10.4774 | 0.5502 | 0.2920 |
| $N = 16$ | 0.6000 | 0.1549 | 0.6271 | 0.6099 | 0.1560 | 4.4717 | 0.9312 | 0.8046 |
| $N = 32$ | 0.6011 | 0.1100 | 0.6136 | 0.6029 | 0.1090 | 1.1799 | 0.9996 | 0.9940 |
| $N = 64$ | 0.6009 | 0.0777 | 0.6038 | 0.6004 | 0.0780 | −0.4144 | 1.0000 | 1.0000 |
| $N = 128$ | 0.6005 | 0.0549 | 0.6020 | 0.6004 | 0.0553 | −0.2033 | 1.0000 | 1.0000 |
| $N = 256$ | 0.6003 | 0.0388 | 0.6015 | 0.6002 | 0.0384 | −0.1776 | 1.0000 | 1.0000 |
| $p = 2, q = 1, V_{rel}(\mathbf{\Sigma}) = 0.8$ | | | | | | | | |
| $N = 4$ | 0.7836 | 0.1336 | 0.8863 | 0.8078 | 0.2061 | 8.3207 | 0.2682 | 0.0708 |
| $N = 8$ | 0.7847 | 0.1153 | 0.8314 | 0.7947 | 0.1473 | 4.8016 | 0.8766 | 0.6756 |
| $N = 16$ | 0.7929 | 0.0866 | 0.8164 | 0.7963 | 0.1004 | 2.4327 | 0.9976 | 0.9890 |
| $N = 32$ | 0.7969 | 0.0625 | 0.8090 | 0.7990 | 0.0672 | 2.2207 | 1.0000 | 1.0000 |
| $N = 64$ | 0.7986 | 0.0445 | 0.8030 | 0.7983 | 0.0470 | −0.3966 | 1.0000 | 1.0000 |
| $N = 128$ | 0.7993 | 0.0315 | 0.8008 | 0.7989 | 0.0318 | −0.9023 | 1.0000 | 1.0000 |
| $N = 256$ | 0.7997 | 0.0223 | 0.8009 | 0.8003 | 0.0223 | 1.8873 | 1.0000 | 1.0000 |

(*continued*)

**Table S2.** (*continued*)

| | $\approx E[V_{rel}(\mathbf{S})]$ | $\approx SD[V_{rel}(\mathbf{S})]$ | Median | Mean | ESD | $T$ | Pow. 5% | Pow. 1% |
|---|---|---|---|---|---|---|---|---|
| $p = 4, q = 1, V_{rel}(\mathbf{\Sigma}) = 0.1$ | | | | | | | | |
| $N = 4$ | 0.4100 | 0.2621 | 0.4457 | 0.4695 | 0.1653 | 25.4117 | 0.0964 | 0.0254 |
| $N = 8$ | 0.2514 | 0.1644 | 0.2472 | 0.2694 | 0.1191 | 10.6867 | 0.1984 | 0.0760 |
| $N = 16$ | 0.1758 | 0.1029 | 0.1661 | 0.1809 | 0.0855 | 4.2193 | 0.4512 | 0.2578 |
| $N = 32$ | 0.1380 | 0.0671 | 0.1316 | 0.1392 | 0.0602 | 1.3758 | 0.7884 | 0.6014 |
| $N = 64$ | 0.1190 | 0.0453 | 0.1150 | 0.1195 | 0.0433 | 0.7437 | 0.9820 | 0.9506 |
| $N = 128$ | 0.1095 | 0.0312 | 0.1084 | 0.1104 | 0.0305 | 2.1101 | 0.9998 | 0.9994 |
| $N = 256$ | 0.1048 | 0.0218 | 0.1034 | 0.1045 | 0.0214 | −0.9490 | 1.0000 | 1.0000 |
| $p = 4, q = 1, V_{rel}(\mathbf{\Sigma}) = 0.2$ | | | | | | | | |
| $N = 4$ | 0.4037 | 0.2878 | 0.4878 | 0.5064 | 0.1746 | 41.5546 | 0.1402 | 0.0424 |
| $N = 8$ | 0.3041 | 0.1990 | 0.3186 | 0.3363 | 0.1442 | 15.8368 | 0.3792 | 0.2024 |
| $N = 16$ | 0.2545 | 0.1316 | 0.2556 | 0.2624 | 0.1086 | 5.0868 | 0.7618 | 0.5808 |
| $N = 32$ | 0.2280 | 0.0887 | 0.2250 | 0.2292 | 0.0792 | 1.0147 | 0.9722 | 0.9310 |
| $N = 64$ | 0.2142 | 0.0610 | 0.2112 | 0.2144 | 0.0581 | 0.2411 | 1.0000 | 0.9998 |
| $N = 128$ | 0.2072 | 0.0425 | 0.2049 | 0.2064 | 0.0411 | −1.3741 | 1.0000 | 1.0000 |
| $N = 256$ | 0.2036 | 0.0298 | 0.2033 | 0.2040 | 0.0296 | 0.8971 | 1.0000 | 1.0000 |
| $p = 4, q = 1, V_{rel}(\mathbf{\Sigma}) = 0.4$ | | | | | | | | |
| $N = 4$ | 0.4602 | 0.2677 | 0.5962 | 0.5896 | 0.1887 | 48.4850 | 0.2948 | 0.1162 |
| $N = 8$ | 0.4319 | 0.2065 | 0.4730 | 0.4675 | 0.1658 | 15.1734 | 0.7126 | 0.5234 |
| $N = 16$ | 0.4200 | 0.1444 | 0.4316 | 0.4288 | 0.1272 | 4.8902 | 0.9750 | 0.9318 |
| $N = 32$ | 0.4113 | 0.1001 | 0.4147 | 0.4125 | 0.0944 | 0.9058 | 1.0000 | 0.9994 |
| $N = 64$ | 0.4060 | 0.0699 | 0.4073 | 0.4067 | 0.0682 | 0.7503 | 1.0000 | 1.0000 |
| $N = 128$ | 0.4031 | 0.0490 | 0.4038 | 0.4026 | 0.0480 | −0.7644 | 1.0000 | 1.0000 |
| $N = 256$ | 0.4016 | 0.0345 | 0.4012 | 0.4008 | 0.0335 | −1.7085 | 1.0000 | 1.0000 |
| $p = 4, q = 1, V_{rel}(\mathbf{\Sigma}) = 0.6$ | | | | | | | | |
| $N = 4$ | 0.5944 | 0.1973 | 0.7186 | 0.6836 | 0.1866 | 33.8195 | 0.5030 | 0.2728 |
| $N = 8$ | 0.5930 | 0.1644 | 0.6442 | 0.6214 | 0.1590 | 12.5992 | 0.9218 | 0.8380 |
| $N = 16$ | 0.5992 | 0.1195 | 0.6198 | 0.6055 | 0.1174 | 3.8086 | 0.9988 | 0.9962 |
| $N = 32$ | 0.6006 | 0.0846 | 0.6086 | 0.6022 | 0.0830 | 1.3653 | 1.0000 | 1.0000 |
| $N = 64$ | 0.6006 | 0.0596 | 0.6043 | 0.6007 | 0.0601 | 0.1178 | 1.0000 | 1.0000 |
| $N = 128$ | 0.6004 | 0.0420 | 0.6017 | 0.5991 | 0.0421 | −2.1326 | 1.0000 | 1.0000 |
| $N = 256$ | 0.6002 | 0.0297 | 0.6011 | 0.6001 | 0.0303 | −0.2881 | 1.0000 | 1.0000 |
| $p = 4, q = 1, V_{rel}(\mathbf{\Sigma}) = 0.8$ | | | | | | | | |
| $N = 4$ | 0.7812 | 0.1037 | 0.8548 | 0.8020 | 0.1610 | 9.1171 | 0.7666 | 0.5686 |
| $N = 8$ | 0.7841 | 0.0920 | 0.8207 | 0.7927 | 0.1170 | 5.1526 | 0.9928 | 0.9804 |
| $N = 16$ | 0.7927 | 0.0692 | 0.8086 | 0.7945 | 0.0793 | 1.6175 | 0.9998 | 0.9998 |
| $N = 32$ | 0.7968 | 0.0498 | 0.8039 | 0.7967 | 0.0537 | −0.1966 | 1.0000 | 1.0000 |
| $N = 64$ | 0.7985 | 0.0354 | 0.8026 | 0.7993 | 0.0368 | 1.5352 | 1.0000 | 1.0000 |
| $N = 128$ | 0.7993 | 0.0251 | 0.8012 | 0.7994 | 0.0255 | 0.2206 | 1.0000 | 1.0000 |
| $N = 256$ | 0.7997 | 0.0177 | 0.8003 | 0.7997 | 0.0176 | 0.0875 | 1.0000 | 1.0000 |

(*continued*)

**Table S2.** (*continued*)

| | ≈ E[$V_{rel}$(**S**)] | ≈ SD[$V_{rel}$(**S**)] | Median | Mean | ESD | $T$ | Pow. 5% | Pow. 1% |
|---|---|---|---|---|---|---|---|---|
| $p = 4$, $q = 2$, $V_{rel}(\mathbf{\Sigma}) = 0.1$ | | | | | | | | |
| $N = 4$ | 0.4605 | 0.2422 | 0.4572 | 0.4810 | 0.1640 | 8.8203 | 0.0980 | 0.0270 |
| $N = 8$ | 0.2728 | 0.1327 | 0.2646 | 0.2784 | 0.1053 | 3.7235 | 0.1932 | 0.0616 |
| $N = 16$ | 0.1851 | 0.0733 | 0.1780 | 0.1855 | 0.0664 | 0.3458 | 0.5050 | 0.2212 |
| $N = 32$ | 0.1423 | 0.0429 | 0.1409 | 0.1433 | 0.0408 | 1.7920 | 0.9230 | 0.7422 |
| $N = 64$ | 0.1211 | 0.0267 | 0.1204 | 0.1215 | 0.0263 | 1.0538 | 0.9998 | 0.9976 |
| $N = 128$ | 0.1105 | 0.0175 | 0.1102 | 0.1107 | 0.0173 | 0.6470 | 1.0000 | 1.0000 |
| $N = 256$ | 0.1053 | 0.0118 | 0.1048 | 0.1049 | 0.0118 | −2.0594 | 1.0000 | 1.0000 |
| $p = 4$, $q = 2$, $V_{rel}(\mathbf{\Sigma}) = 0.2$ | | | | | | | | |
| $N = 4$ | 0.5313 | 0.2669 | 0.5324 | 0.5467 | 0.1745 | 6.2103 | 0.1934 | 0.0684 |
| $N = 8$ | 0.3615 | 0.1449 | 0.3441 | 0.3667 | 0.1203 | 3.1014 | 0.4370 | 0.2064 |
| $N = 16$ | 0.2803 | 0.0771 | 0.2710 | 0.2804 | 0.0687 | 0.1260 | 0.9646 | 0.7720 |
| $N = 32$ | 0.2401 | 0.0426 | 0.2379 | 0.2413 | 0.0412 | 2.1470 | 1.0000 | 0.9994 |
| $N = 64$ | 0.2200 | 0.0249 | 0.2186 | 0.2199 | 0.0246 | −0.4333 | 1.0000 | 1.0000 |
| $N = 128$ | 0.2100 | 0.0154 | 0.2097 | 0.2099 | 0.0158 | −0.5985 | 1.0000 | 1.0000 |
| $N = 256$ | 0.2050 | 0.0101 | 0.2049 | 0.2051 | 0.0100 | 0.3652 | 1.0000 | 1.0000 |
| $p = 4$, $q = 2$, $V_{rel}(\mathbf{\Sigma}) = 0.4$ | | | | | | | | |
| (This comformation is impossible) | | | | | | | | |

(*continued*)

**Table S2.** (*continued*)

| | $\approx$ E[$V_{rel}$(**S**)] | $\approx$ SD[$V_{rel}$(**S**)] | Median | Mean | ESD | $T$ | Pow. 5% | Pow. 1% |
|---|---|---|---|---|---|---|---|---|
| $p = 8, q = 1, V_{rel}(\boldsymbol{\Sigma}) = 0.1$ | | | | | | | | |
| $N = 4$ | 0.3892 | 0.1871 | 0.4158 | 0.4355 | 0.1090 | 30.0381 | 0.1688 | 0.0620 |
| $N = 8$ | 0.2368 | 0.1191 | 0.2305 | 0.2475 | 0.0838 | 9.0124 | 0.4426 | 0.2874 |
| $N = 16$ | 0.1673 | 0.0758 | 0.1604 | 0.1698 | 0.0621 | 2.8253 | 0.7950 | 0.6780 |
| $N = 32$ | 0.1334 | 0.0502 | 0.1299 | 0.1343 | 0.0452 | 1.4051 | 0.9828 | 0.9576 |
| $N = 64$ | 0.1167 | 0.0342 | 0.1140 | 0.1171 | 0.0326 | 0.8569 | 0.9998 | 0.9994 |
| $N = 128$ | 0.1083 | 0.0237 | 0.1069 | 0.1080 | 0.0230 | −1.0270 | 1.0000 | 1.0000 |
| $N = 256$ | 0.1042 | 0.0166 | 0.1034 | 0.1043 | 0.0166 | 0.7600 | 1.0000 | 1.0000 |
| $p = 8, q = 1, V_{rel}(\boldsymbol{\Sigma}) = 0.2$ | | | | | | | | |
| $N = 4$ | 0.3906 | 0.2315 | 0.4683 | 0.4859 | 0.1313 | 51.3142 | 0.3222 | 0.1598 |
| $N = 8$ | 0.2957 | 0.1608 | 0.3077 | 0.3226 | 0.1115 | 17.0187 | 0.7298 | 0.5902 |
| $N = 16$ | 0.2495 | 0.1066 | 0.2474 | 0.2544 | 0.0874 | 3.9350 | 0.9640 | 0.9300 |
| $N = 32$ | 0.2252 | 0.0721 | 0.2229 | 0.2269 | 0.0661 | 1.7453 | 1.0000 | 0.9996 |
| $N = 64$ | 0.2127 | 0.0496 | 0.2102 | 0.2125 | 0.0480 | −0.2832 | 1.0000 | 1.0000 |
| $N = 128$ | 0.2064 | 0.0346 | 0.2067 | 0.2073 | 0.0333 | 1.8083 | 1.0000 | 1.0000 |
| $N = 256$ | 0.2032 | 0.0243 | 0.2029 | 0.2030 | 0.0241 | −0.6017 | 1.0000 | 1.0000 |
| $p = 8, q = 1, V_{rel}(\boldsymbol{\Sigma}) = 0.4$ | | | | | | | | |
| $N = 4$ | 0.4525 | 0.2334 | 0.5701 | 0.5731 | 0.1549 | 55.1063 | 0.5656 | 0.3908 |
| $N = 8$ | 0.4290 | 0.1803 | 0.4659 | 0.4613 | 0.1429 | 15.9923 | 0.9252 | 0.8754 |
| $N = 16$ | 0.4183 | 0.1256 | 0.4316 | 0.4290 | 0.1092 | 6.9108 | 0.9986 | 0.9980 |
| $N = 32$ | 0.4103 | 0.0870 | 0.4153 | 0.4128 | 0.0823 | 2.1081 | 1.0000 | 1.0000 |
| $N = 64$ | 0.4055 | 0.0606 | 0.4078 | 0.4063 | 0.0585 | 0.9585 | 1.0000 | 1.0000 |
| $N = 128$ | 0.4028 | 0.0425 | 0.4038 | 0.4034 | 0.0417 | 1.0354 | 1.0000 | 1.0000 |
| $N = 256$ | 0.4014 | 0.0299 | 0.4013 | 0.4011 | 0.0292 | −0.8338 | 1.0000 | 1.0000 |
| $p = 8, q = 1, V_{rel}(\boldsymbol{\Sigma}) = 0.6$ | | | | | | | | |
| $N = 4$ | 0.5903 | 0.1765 | 0.6998 | 0.6731 | 0.1648 | 35.5306 | 0.7656 | 0.6362 |
| $N = 8$ | 0.5920 | 0.1482 | 0.6382 | 0.6170 | 0.1435 | 12.3162 | 0.9902 | 0.9768 |
| $N = 16$ | 0.5987 | 0.1074 | 0.6205 | 0.6077 | 0.1052 | 6.0548 | 0.9998 | 0.9998 |
| $N = 32$ | 0.6004 | 0.0758 | 0.6060 | 0.5997 | 0.0766 | −0.6096 | 1.0000 | 1.0000 |
| $N = 64$ | 0.6005 | 0.0534 | 0.6031 | 0.6005 | 0.0536 | 0.0160 | 1.0000 | 1.0000 |
| $N = 128$ | 0.6003 | 0.0376 | 0.6023 | 0.6003 | 0.0370 | −0.1299 | 1.0000 | 1.0000 |
| $N = 256$ | 0.6002 | 0.0265 | 0.6012 | 0.6001 | 0.0266 | −0.3410 | 1.0000 | 1.0000 |
| $p = 8, q = 1, V_{rel}(\boldsymbol{\Sigma}) = 0.8$ | | | | | | | | |
| $N = 4$ | 0.7800 | 0.0936 | 0.8464 | 0.8015 | 0.1448 | 10.4973 | 0.9262 | 0.8738 |
| $N = 8$ | 0.7839 | 0.0843 | 0.8159 | 0.7910 | 0.1083 | 4.6211 | 0.9996 | 0.9990 |
| $N = 16$ | 0.7926 | 0.0634 | 0.8056 | 0.7930 | 0.0728 | 0.3423 | 1.0000 | 1.0000 |
| $N = 32$ | 0.7968 | 0.0455 | 0.8030 | 0.7978 | 0.0473 | 1.5850 | 1.0000 | 1.0000 |
| $N = 64$ | 0.7985 | 0.0323 | 0.8008 | 0.7982 | 0.0331 | −0.7649 | 1.0000 | 1.0000 |
| $N = 128$ | 0.7993 | 0.0229 | 0.8011 | 0.7997 | 0.0231 | 1.0882 | 1.0000 | 1.0000 |
| $N = 256$ | 0.7997 | 0.0162 | 0.8001 | 0.7994 | 0.0162 | −1.1600 | 1.0000 | 1.0000 |

(*continued*)

**Table S2.** (*continued*)

| | ≈ E[$V_{rel}$(**S**)] | ≈ SD[$V_{rel}$(**S**)] | Median | Mean | ESD | $T$ | Pow. 5% | Pow. 1% |
|---|---|---|---|---|---|---|---|---|
| $p = 8, q = 2, V_{rel}(\mathbf{\Sigma}) = 0.1$ | | | | | | | | |
| $N = 4$ | 0.4290 | 0.1604 | 0.4268 | 0.4460 | 0.1079 | 11.1170 | 0.1910 | 0.0696 |
| $N = 8$ | 0.2529 | 0.0895 | 0.2481 | 0.2581 | 0.0725 | 5.1536 | 0.5284 | 0.3212 |
| $N = 16$ | 0.1742 | 0.0517 | 0.1705 | 0.1746 | 0.0460 | 0.5660 | 0.9114 | 0.8146 |
| $N = 32$ | 0.1366 | 0.0319 | 0.1351 | 0.1370 | 0.0302 | 0.8562 | 0.9996 | 0.9974 |
| $N = 64$ | 0.1182 | 0.0208 | 0.1167 | 0.1176 | 0.0202 | −2.1649 | 1.0000 | 1.0000 |
| $N = 128$ | 0.1091 | 0.0141 | 0.1090 | 0.1091 | 0.0139 | 0.4297 | 1.0000 | 1.0000 |
| $N = 256$ | 0.1045 | 0.0097 | 0.1045 | 0.1045 | 0.0096 | −0.2327 | 1.0000 | 1.0000 |
| $p = 8, q = 2, V_{rel}(\mathbf{\Sigma}) = 0.2$ | | | | | | | | |
| $N = 4$ | 0.4926 | 0.1956 | 0.4995 | 0.5161 | 0.1295 | 12.8805 | 0.3960 | 0.2078 |
| $N = 8$ | 0.3393 | 0.1091 | 0.3322 | 0.3433 | 0.0889 | 3.1755 | 0.8854 | 0.7434 |
| $N = 16$ | 0.2687 | 0.0615 | 0.2643 | 0.2696 | 0.0573 | 1.1307 | 0.9990 | 0.9956 |
| $N = 32$ | 0.2341 | 0.0368 | 0.2330 | 0.2346 | 0.0361 | 0.9088 | 1.0000 | 1.0000 |
| $N = 64$ | 0.2170 | 0.0234 | 0.2166 | 0.2168 | 0.0231 | −0.5343 | 1.0000 | 1.0000 |
| $N = 128$ | 0.2085 | 0.0155 | 0.2083 | 0.2083 | 0.0153 | −0.7404 | 1.0000 | 1.0000 |
| $N = 256$ | 0.2042 | 0.0106 | 0.2043 | 0.2042 | 0.0103 | −0.0092 | 1.0000 | 1.0000 |
| $p = 8, q = 2, V_{rel}(\mathbf{\Sigma}) = 0.4$ | | | | | | | | |
| $N = 4$ | 0.6820 | 0.2513 | 0.6817 | 0.6803 | 0.1556 | −0.7726 | 0.7586 | 0.5986 |
| $N = 8$ | 0.5399 | 0.1346 | 0.5145 | 0.5385 | 0.1050 | −0.9238 | 1.0000 | 1.0000 |
| $N = 16$ | 0.4698 | 0.0683 | 0.4529 | 0.4693 | 0.0602 | −0.6598 | 1.0000 | 1.0000 |
| $N = 32$ | 0.4349 | 0.0344 | 0.4258 | 0.4348 | 0.0316 | −0.3472 | 1.0000 | 1.0000 |
| $N = 64$ | 0.4175 | 0.0174 | 0.4131 | 0.4176 | 0.0169 | 0.4181 | 1.0000 | 1.0000 |
| $N = 128$ | 0.4087 | 0.0089 | 0.4066 | 0.4088 | 0.0088 | 0.2251 | 1.0000 | 1.0000 |
| $N = 256$ | 0.4044 | 0.0047 | 0.4034 | 0.4044 | 0.0048 | 0.2576 | 1.0000 | 1.0000 |
| $p = 8, q = 4, V_{rel}(\mathbf{\Sigma}) = 0.1$ | | | | | | | | |
| $N = 4$ | 0.4646 | 0.1481 | 0.4458 | 0.4666 | 0.1129 | 1.2361 | 0.2366 | 0.0988 |
| $N = 8$ | 0.2682 | 0.0717 | 0.2611 | 0.2699 | 0.0636 | 1.8384 | 0.6264 | 0.3590 |
| $N = 16$ | 0.1811 | 0.0351 | 0.1770 | 0.1817 | 0.0335 | 1.3536 | 0.9980 | 0.9606 |
| $N = 32$ | 0.1398 | 0.0177 | 0.1379 | 0.1397 | 0.0170 | −0.3943 | 1.0000 | 1.0000 |
| $N = 64$ | 0.1197 | 0.0093 | 0.1189 | 0.1198 | 0.0092 | 0.7866 | 1.0000 | 1.0000 |
| $N = 128$ | 0.1098 | 0.0052 | 0.1095 | 0.1098 | 0.0051 | 0.1577 | 1.0000 | 1.0000 |
| $N = 256$ | 0.1049 | 0.0030 | 0.1048 | 0.1049 | 0.0030 | −0.9935 | 1.0000 | 1.0000 |
| $p = 8, q = 4, V_{rel}(\mathbf{\Sigma}) = 0.2$ | | | | | | | | |
| (This comformation is impossible) | | | | | | | | |

(*continued*)

**Table S2.** (*continued*)

| | ≈ E[$V_{rel}$(**S**)] | ≈ SD[$V_{rel}$(**S**)] | Median | Mean | ESD | $T$ | Pow. 5% | Pow. 1% |
|---|---|---|---|---|---|---|---|---|
| $p = 16, q = 1, V_{rel}(\mathbf{\Sigma}) = 0.1$ | | | | | | | | |
| $N = 4$ | 0.3781 | 0.1547 | 0.3993 | 0.4200 | 0.0844 | 35.1233 | 0.3406 | 0.1980 |
| $N = 8$ | 0.2291 | 0.1000 | 0.2267 | 0.2407 | 0.0704 | 11.6244 | 0.7040 | 0.5780 |
| $N = 16$ | 0.1629 | 0.0643 | 0.1566 | 0.1645 | 0.0534 | 2.1175 | 0.9522 | 0.9166 |
| $N = 32$ | 0.1311 | 0.0429 | 0.1278 | 0.1315 | 0.0392 | 0.7208 | 0.9994 | 0.9984 |
| $N = 64$ | 0.1155 | 0.0293 | 0.1132 | 0.1157 | 0.0283 | 0.6514 | 1.0000 | 1.0000 |
| $N = 128$ | 0.1077 | 0.0204 | 0.1069 | 0.1079 | 0.0201 | 0.5728 | 1.0000 | 1.0000 |
| $N = 256$ | 0.1039 | 0.0143 | 0.1033 | 0.1039 | 0.0142 | 0.0771 | 1.0000 | 1.0000 |
| $p = 16, q = 1, V_{rel}(\mathbf{\Sigma}) = 0.2$ | | | | | | | | |
| $N = 4$ | 0.3839 | 0.2075 | 0.4479 | 0.4692 | 0.1109 | 54.4282 | 0.5346 | 0.3688 |
| $N = 8$ | 0.2914 | 0.1443 | 0.3019 | 0.3139 | 0.1010 | 15.7447 | 0.8904 | 0.8238 |
| $N = 16$ | 0.2469 | 0.0958 | 0.2456 | 0.2524 | 0.0791 | 4.9064 | 0.9968 | 0.9936 |
| $N = 32$ | 0.2238 | 0.0648 | 0.2218 | 0.2241 | 0.0585 | 0.3753 | 1.0000 | 1.0000 |
| $N = 64$ | 0.2120 | 0.0447 | 0.2112 | 0.2122 | 0.0426 | 0.3554 | 1.0000 | 1.0000 |
| $N = 128$ | 0.2060 | 0.0312 | 0.2048 | 0.2055 | 0.0303 | −1.2781 | 1.0000 | 1.0000 |
| $N = 256$ | 0.2030 | 0.0219 | 0.2026 | 0.2030 | 0.0216 | −0.1191 | 1.0000 | 1.0000 |
| $p = 16, q = 1, V_{rel}(\mathbf{\Sigma}) = 0.4$ | | | | | | | | |
| $N = 4$ | 0.4486 | 0.2189 | 0.5668 | 0.5678 | 0.1418 | 59.4299 | 0.7750 | 0.6596 |
| $N = 8$ | 0.4275 | 0.1691 | 0.4657 | 0.4626 | 0.1346 | 18.4219 | 0.9828 | 0.9698 |
| $N = 16$ | 0.4174 | 0.1175 | 0.4302 | 0.4270 | 0.1049 | 6.4710 | 1.0000 | 1.0000 |
| $N = 32$ | 0.4098 | 0.0813 | 0.4161 | 0.4129 | 0.0762 | 2.8682 | 1.0000 | 1.0000 |
| $N = 64$ | 0.4052 | 0.0566 | 0.4070 | 0.4058 | 0.0548 | 0.7118 | 1.0000 | 1.0000 |
| $N = 128$ | 0.4027 | 0.0397 | 0.4042 | 0.4038 | 0.0394 | 1.9337 | 1.0000 | 1.0000 |
| $N = 256$ | 0.4014 | 0.0279 | 0.4013 | 0.4012 | 0.0280 | −0.3594 | 1.0000 | 1.0000 |
| $p = 16, q = 1, V_{rel}(\mathbf{\Sigma}) = 0.6$ | | | | | | | | |
| $N = 4$ | 0.5882 | 0.1678 | 0.6919 | 0.6700 | 0.1577 | 36.6795 | 0.8948 | 0.8326 |
| $N = 8$ | 0.5915 | 0.1412 | 0.6344 | 0.6165 | 0.1383 | 12.8135 | 0.9982 | 0.9954 |
| $N = 16$ | 0.5985 | 0.1023 | 0.6172 | 0.6057 | 0.1014 | 5.0298 | 1.0000 | 1.0000 |
| $N = 32$ | 0.6003 | 0.0721 | 0.6085 | 0.6022 | 0.0714 | 1.9527 | 1.0000 | 1.0000 |
| $N = 64$ | 0.6004 | 0.0507 | 0.6032 | 0.6007 | 0.0506 | 0.4295 | 1.0000 | 1.0000 |
| $N = 128$ | 0.6003 | 0.0357 | 0.6020 | 0.6007 | 0.0361 | 0.8161 | 1.0000 | 1.0000 |
| $N = 256$ | 0.6002 | 0.0252 | 0.6006 | 0.5999 | 0.0253 | −0.8000 | 1.0000 | 1.0000 |
| $p = 16, q = 1, V_{rel}(\mathbf{\Sigma}) = 0.8$ | | | | | | | | |
| $N = 4$ | 0.7795 | 0.0893 | 0.8397 | 0.7967 | 0.1413 | 8.6401 | 0.9688 | 0.9514 |
| $N = 8$ | 0.7838 | 0.0810 | 0.8153 | 0.7913 | 0.1049 | 5.0688 | 0.9998 | 0.9998 |
| $N = 16$ | 0.7926 | 0.0609 | 0.8067 | 0.7943 | 0.0696 | 1.7415 | 1.0000 | 1.0000 |
| $N = 32$ | 0.7968 | 0.0437 | 0.8044 | 0.7989 | 0.0452 | 3.3474 | 1.0000 | 1.0000 |
| $N = 64$ | 0.7985 | 0.0310 | 0.8019 | 0.7990 | 0.0323 | 1.0675 | 1.0000 | 1.0000 |
| $N = 128$ | 0.7993 | 0.0220 | 0.8000 | 0.7988 | 0.0223 | −1.7432 | 1.0000 | 1.0000 |
| $N = 256$ | 0.7997 | 0.0155 | 0.8000 | 0.7996 | 0.0157 | −0.2993 | 1.0000 | 1.0000 |

(*continued*)

**Table S2.** (*continued*)

| | ≈ $E[V_{rel}(\mathbf{S})]$ | ≈ $SD[V_{rel}(\mathbf{S})]$ | Median | Mean | ESD | $T$ | Pow. 5% | Pow. 1% |
|---|---|---|---|---|---|---|---|---|
| $p = 16, q = 2, V_{rel}(\mathbf{\Sigma}) = 0.1$ | | | | | | | | |
| $N = 4$ | 0.4143 | 0.1238 | 0.4134 | 0.4286 | 0.0815 | 12.3917 | 0.3806 | 0.1976 |
| $N = 8$ | 0.2434 | 0.0706 | 0.2383 | 0.2455 | 0.0557 | 2.5532 | 0.8294 | 0.6884 |
| $N = 16$ | 0.1691 | 0.0419 | 0.1666 | 0.1696 | 0.0369 | 1.0663 | 0.9958 | 0.9908 |
| $N = 32$ | 0.1339 | 0.0266 | 0.1333 | 0.1345 | 0.0255 | 1.5713 | 1.0000 | 1.0000 |
| $N = 64$ | 0.1168 | 0.0176 | 0.1164 | 0.1170 | 0.0173 | 0.8504 | 1.0000 | 1.0000 |
| $N = 128$ | 0.1084 | 0.0120 | 0.1079 | 0.1082 | 0.0119 | −1.0759 | 1.0000 | 1.0000 |
| $N = 256$ | 0.1042 | 0.0084 | 0.1037 | 0.1040 | 0.0084 | −1.3172 | 1.0000 | 1.0000 |
| $p = 16, q = 2, V_{rel}(\mathbf{\Sigma}) = 0.2$ | | | | | | | | |
| $N = 4$ | 0.4775 | 0.1649 | 0.4801 | 0.4984 | 0.1080 | 13.7304 | 0.6566 | 0.4588 |
| $N = 8$ | 0.3305 | 0.0936 | 0.3271 | 0.3362 | 0.0744 | 5.3878 | 0.9894 | 0.9730 |
| $N = 16$ | 0.2640 | 0.0543 | 0.2616 | 0.2646 | 0.0499 | 0.8966 | 1.0000 | 1.0000 |
| $N = 32$ | 0.2317 | 0.0334 | 0.2306 | 0.2316 | 0.0320 | −0.2412 | 1.0000 | 1.0000 |
| $N = 64$ | 0.2158 | 0.0217 | 0.2152 | 0.2151 | 0.0213 | −2.2294 | 1.0000 | 1.0000 |
| $N = 128$ | 0.2079 | 0.0147 | 0.2077 | 0.2077 | 0.0146 | −0.9653 | 1.0000 | 1.0000 |
| $N = 256$ | 0.2039 | 0.0101 | 0.2043 | 0.2042 | 0.0098 | 1.6488 | 1.0000 | 1.0000 |
| $p = 16, q = 2, V_{rel}(\mathbf{\Sigma}) = 0.4$ | | | | | | | | |
| $N = 4$ | 0.6595 | 0.2219 | 0.6539 | 0.6600 | 0.1378 | 0.2882 | 0.9756 | 0.8632 |
| $N = 8$ | 0.5273 | 0.1187 | 0.5057 | 0.5269 | 0.0936 | −0.3636 | 1.0000 | 1.0000 |
| $N = 16$ | 0.4634 | 0.0608 | 0.4512 | 0.4639 | 0.0540 | 0.5775 | 1.0000 | 1.0000 |
| $N = 32$ | 0.4317 | 0.0314 | 0.4254 | 0.4310 | 0.0285 | −1.7871 | 1.0000 | 1.0000 |
| $N = 64$ | 0.4158 | 0.0167 | 0.4131 | 0.4155 | 0.0159 | −1.6153 | 1.0000 | 1.0000 |
| $N = 128$ | 0.4079 | 0.0093 | 0.4068 | 0.4079 | 0.0093 | −0.0692 | 1.0000 | 1.0000 |
| $N = 256$ | 0.4040 | 0.0054 | 0.4035 | 0.4039 | 0.0055 | −1.0354 | 1.0000 | 1.0000 |
| $p = 16, q = 4, V_{rel}(\mathbf{\Sigma}) = 0.1$ | | | | | | | | |
| $N = 4$ | 0.4407 | 0.1087 | 0.4285 | 0.4431 | 0.0808 | 2.0856 | 0.4498 | 0.2388 |
| $N = 8$ | 0.2545 | 0.0534 | 0.2481 | 0.2548 | 0.0460 | 0.4484 | 0.9444 | 0.8240 |
| $N = 16$ | 0.1740 | 0.0273 | 0.1716 | 0.1746 | 0.0259 | 1.7728 | 1.0000 | 1.0000 |
| $N = 32$ | 0.1362 | 0.0149 | 0.1349 | 0.1358 | 0.0144 | −2.2596 | 1.0000 | 1.0000 |
| $N = 64$ | 0.1179 | 0.0087 | 0.1176 | 0.1178 | 0.0085 | −1.1815 | 1.0000 | 1.0000 |
| $N = 128$ | 0.1089 | 0.0055 | 0.1089 | 0.1091 | 0.0054 | 2.0563 | 1.0000 | 1.0000 |
| $N = 256$ | 0.1044 | 0.0036 | 0.1044 | 0.1044 | 0.0036 | −0.7657 | 1.0000 | 1.0000 |
| $p = 16, q = 4, V_{rel}(\mathbf{\Sigma}) = 0.2$ | | | | | | | | |
| $N = 4$ | 0.5429 | 0.1577 | 0.5188 | 0.5399 | 0.1191 | −1.7637 | 0.8000 | 0.5890 |
| $N = 8$ | 0.3600 | 0.0765 | 0.3492 | 0.3588 | 0.0665 | −1.2879 | 1.0000 | 1.0000 |
| $N = 16$ | 0.2774 | 0.0369 | 0.2721 | 0.2773 | 0.0347 | −0.1483 | 1.0000 | 1.0000 |
| $N = 32$ | 0.2381 | 0.0181 | 0.2359 | 0.2382 | 0.0176 | 0.5801 | 1.0000 | 1.0000 |
| $N = 64$ | 0.2189 | 0.0089 | 0.2174 | 0.2188 | 0.0088 | −0.4406 | 1.0000 | 1.0000 |
| $N = 128$ | 0.2094 | 0.0044 | 0.2087 | 0.2095 | 0.0045 | 1.3265 | 1.0000 | 1.0000 |
| $N = 256$ | 0.2047 | 0.0022 | 0.2043 | 0.2047 | 0.0022 | −1.4616 | 1.0000 | 1.0000 |

(*continued*)

**Table S2.** (*continued*)

| | ≈ E[$V_{rel}$(**S**)] | ≈ SD[$V_{rel}$(**S**)] | Median | Mean | ESD | $T$ | Pow. 5% | Pow. 1% |
|---|---|---|---|---|---|---|---|---|
| $p = 32, q = 1, V_{rel}(\mathbf{\Sigma}) = 0.1$ | | | | | | | | |
| $N = 4$ | 0.3722 | 0.1401 | 0.3890 | 0.4085 | 0.0722 | 35.5187 | 0.5116 | 0.3740 |
| $N = 8$ | 0.2252 | 0.0913 | 0.2179 | 0.2316 | 0.0629 | 7.1557 | 0.8658 | 0.7946 |
| $N = 16$ | 0.1607 | 0.0590 | 0.1578 | 0.1642 | 0.0492 | 4.9649 | 0.9948 | 0.9884 |
| $N = 32$ | 0.1300 | 0.0395 | 0.1274 | 0.1307 | 0.0355 | 1.5884 | 1.0000 | 1.0000 |
| $N = 64$ | 0.1149 | 0.0271 | 0.1135 | 0.1151 | 0.0258 | 0.5694 | 1.0000 | 1.0000 |
| $N = 128$ | 0.1074 | 0.0188 | 0.1063 | 0.1069 | 0.0185 | −2.1220 | 1.0000 | 1.0000 |
| $N = 256$ | 0.1037 | 0.0132 | 0.1028 | 0.1033 | 0.0129 | −2.0896 | 1.0000 | 1.0000 |
| $p = 32, q = 1, V_{rel}(\mathbf{\Sigma}) = 0.2$ | | | | | | | | |
| $N = 4$ | 0.3805 | 0.1965 | 0.4370 | 0.4585 | 0.1007 | 54.7782 | 0.6940 | 0.5928 |
| $N = 8$ | 0.2893 | 0.1367 | 0.2993 | 0.3106 | 0.0957 | 15.8065 | 0.9646 | 0.9436 |
| $N = 16$ | 0.2456 | 0.0907 | 0.2423 | 0.2484 | 0.0741 | 2.6792 | 1.0000 | 0.9996 |
| $N = 32$ | 0.2231 | 0.0614 | 0.2216 | 0.2249 | 0.0564 | 2.2321 | 1.0000 | 1.0000 |
| $N = 64$ | 0.2116 | 0.0424 | 0.2110 | 0.2126 | 0.0401 | 1.6316 | 1.0000 | 1.0000 |
| $N = 128$ | 0.2058 | 0.0296 | 0.2061 | 0.2065 | 0.0284 | 1.6193 | 1.0000 | 1.0000 |
| $N = 256$ | 0.2029 | 0.0208 | 0.2028 | 0.2031 | 0.0202 | 0.5471 | 1.0000 | 1.0000 |
| $p = 32, q = 1, V_{rel}(\mathbf{\Sigma}) = 0.4$ | | | | | | | | |
| $N = 4$ | 0.4467 | 0.2122 | 0.5598 | 0.5635 | 0.1376 | 60.0007 | 0.8810 | 0.8234 |
| $N = 8$ | 0.4268 | 0.1638 | 0.4631 | 0.4583 | 0.1291 | 17.2526 | 0.9966 | 0.9944 |
| $N = 16$ | 0.4170 | 0.1137 | 0.4271 | 0.4250 | 0.0995 | 5.6662 | 1.0000 | 1.0000 |
| $N = 32$ | 0.4096 | 0.0786 | 0.4154 | 0.4128 | 0.0756 | 2.9659 | 1.0000 | 1.0000 |
| $N = 64$ | 0.4051 | 0.0547 | 0.4072 | 0.4057 | 0.0529 | 0.7484 | 1.0000 | 1.0000 |
| $N = 128$ | 0.4026 | 0.0384 | 0.4038 | 0.4036 | 0.0382 | 1.8820 | 1.0000 | 1.0000 |
| $N = 256$ | 0.4013 | 0.0270 | 0.4010 | 0.4011 | 0.0265 | −0.5358 | 1.0000 | 1.0000 |
| $p = 32, q = 1, V_{rel}(\mathbf{\Sigma}) = 0.6$ | | | | | | | | |
| $N = 4$ | 0.5872 | 0.1637 | 0.6947 | 0.6706 | 0.1518 | 38.8361 | 0.9460 | 0.9222 |
| $N = 8$ | 0.5912 | 0.1380 | 0.6345 | 0.6170 | 0.1313 | 13.8710 | 1.0000 | 0.9998 |
| $N = 16$ | 0.5984 | 0.0998 | 0.6160 | 0.6041 | 0.0995 | 4.0499 | 1.0000 | 1.0000 |
| $N = 32$ | 0.6002 | 0.0703 | 0.6081 | 0.6018 | 0.0706 | 1.6224 | 1.0000 | 1.0000 |
| $N = 64$ | 0.6004 | 0.0494 | 0.6041 | 0.6014 | 0.0492 | 1.4951 | 1.0000 | 1.0000 |
| $N = 128$ | 0.6003 | 0.0348 | 0.6004 | 0.5996 | 0.0348 | −1.3790 | 1.0000 | 1.0000 |
| $N = 256$ | 0.6002 | 0.0246 | 0.6011 | 0.6002 | 0.0245 | 0.0139 | 1.0000 | 1.0000 |
| $p = 32, q = 1, V_{rel}(\mathbf{\Sigma}) = 0.8$ | | | | | | | | |
| $N = 4$ | 0.7792 | 0.0873 | 0.8365 | 0.7949 | 0.1393 | 7.9763 | 0.9844 | 0.9772 |
| $N = 8$ | 0.7837 | 0.0794 | 0.8157 | 0.7922 | 0.1009 | 5.9763 | 1.0000 | 1.0000 |
| $N = 16$ | 0.7926 | 0.0597 | 0.8050 | 0.7935 | 0.0684 | 0.9693 | 1.0000 | 1.0000 |
| $N = 32$ | 0.7967 | 0.0429 | 0.8033 | 0.7977 | 0.0452 | 1.5534 | 1.0000 | 1.0000 |
| $N = 64$ | 0.7985 | 0.0304 | 0.8008 | 0.7983 | 0.0308 | −0.6108 | 1.0000 | 1.0000 |
| $N = 128$ | 0.7993 | 0.0215 | 0.8011 | 0.7998 | 0.0215 | 1.6288 | 1.0000 | 1.0000 |
| $N = 256$ | 0.7997 | 0.0152 | 0.8001 | 0.7994 | 0.0154 | −1.2286 | 1.0000 | 1.0000 |

(*continued*)

**Table S2.** (*continued*)

| | ≈ E[$V_{rel}(\mathbf{S})$] | ≈ SD[$V_{rel}(\mathbf{S})$] | Median | Mean | ESD | $T$ | Pow. 5% | Pow. 1% |
|---|---|---|---|---|---|---|---|---|
| $p = 32, q = 2, V_{rel}(\mathbf{\Sigma}) = 0.1$ | | | | | | | | |
| $N = 4$ | 0.4069 | 0.1066 | 0.4070 | 0.4201 | 0.0673 | 13.8253 | 0.6254 | 0.4612 |
| $N = 8$ | 0.2388 | 0.0619 | 0.2344 | 0.2412 | 0.0491 | 3.4320 | 0.9654 | 0.9284 |
| $N = 16$ | 0.1665 | 0.0374 | 0.1634 | 0.1660 | 0.0336 | −1.0262 | 0.9998 | 0.9996 |
| $N = 32$ | 0.1326 | 0.0240 | 0.1319 | 0.1326 | 0.0229 | −0.0886 | 1.0000 | 1.0000 |
| $N = 64$ | 0.1162 | 0.0161 | 0.1160 | 0.1163 | 0.0156 | 0.7616 | 1.0000 | 1.0000 |
| $N = 128$ | 0.1080 | 0.0110 | 0.1077 | 0.1080 | 0.0108 | −0.3648 | 1.0000 | 1.0000 |
| $N = 256$ | 0.1040 | 0.0077 | 0.1041 | 0.1041 | 0.0076 | 1.0367 | 1.0000 | 1.0000 |
| $p = 32, q = 2, V_{rel}(\mathbf{\Sigma}) = 0.2$ | | | | | | | | |
| $N = 4$ | 0.4706 | 0.1506 | 0.4771 | 0.4921 | 0.0964 | 15.8001 | 0.8662 | 0.7700 |
| $N = 8$ | 0.3264 | 0.0864 | 0.3271 | 0.3329 | 0.0703 | 6.4974 | 0.9990 | 0.9980 |
| $N = 16$ | 0.2618 | 0.0508 | 0.2609 | 0.2630 | 0.0454 | 1.8913 | 1.0000 | 1.0000 |
| $N = 32$ | 0.2306 | 0.0317 | 0.2300 | 0.2310 | 0.0305 | 0.8537 | 1.0000 | 1.0000 |
| $N = 64$ | 0.2152 | 0.0208 | 0.2153 | 0.2158 | 0.0207 | 1.9430 | 1.0000 | 1.0000 |
| $N = 128$ | 0.2076 | 0.0141 | 0.2075 | 0.2076 | 0.0141 | −0.0879 | 1.0000 | 1.0000 |
| $N = 256$ | 0.2038 | 0.0098 | 0.2038 | 0.2038 | 0.0096 | −0.1538 | 1.0000 | 1.0000 |
| $p = 32, q = 2, V_{rel}(\mathbf{\Sigma}) = 0.4$ | | | | | | | | |
| $N = 4$ | 0.6497 | 0.2088 | 0.6458 | 0.6505 | 0.1291 | 0.4379 | 0.9976 | 0.9916 |
| $N = 8$ | 0.5219 | 0.1118 | 0.5050 | 0.5229 | 0.0885 | 0.7836 | 1.0000 | 1.0000 |
| $N = 16$ | 0.4607 | 0.0578 | 0.4507 | 0.4615 | 0.0518 | 1.0509 | 1.0000 | 1.0000 |
| $N = 32$ | 0.4303 | 0.0303 | 0.4253 | 0.4303 | 0.0291 | −0.0698 | 1.0000 | 1.0000 |
| $N = 64$ | 0.4152 | 0.0166 | 0.4135 | 0.4153 | 0.0162 | 0.5373 | 1.0000 | 1.0000 |
| $N = 128$ | 0.4076 | 0.0095 | 0.4071 | 0.4076 | 0.0095 | 0.2889 | 1.0000 | 1.0000 |
| $N = 256$ | 0.4038 | 0.0058 | 0.4037 | 0.4039 | 0.0058 | 1.8489 | 1.0000 | 1.0000 |
| $p = 32, q = 4, V_{rel}(\mathbf{\Sigma}) = 0.1$ | | | | | | | | |
| $N = 4$ | 0.4304 | 0.0901 | 0.4220 | 0.4353 | 0.0680 | 5.1163 | 0.7372 | 0.5570 |
| $N = 8$ | 0.2485 | 0.0448 | 0.2434 | 0.2492 | 0.0396 | 1.2297 | 0.9980 | 0.9914 |
| $N = 16$ | 0.1708 | 0.0236 | 0.1696 | 0.1716 | 0.0223 | 2.5933 | 1.0000 | 1.0000 |
| $N = 32$ | 0.1346 | 0.0134 | 0.1341 | 0.1346 | 0.0132 | 0.2360 | 1.0000 | 1.0000 |
| $N = 64$ | 0.1171 | 0.0082 | 0.1171 | 0.1173 | 0.0081 | 1.2790 | 1.0000 | 1.0000 |
| $N = 128$ | 0.1085 | 0.0053 | 0.1085 | 0.1085 | 0.0053 | −0.2026 | 1.0000 | 1.0000 |
| $N = 256$ | 0.1042 | 0.0036 | 0.1042 | 0.1042 | 0.0035 | −0.3077 | 1.0000 | 1.0000 |
| $p = 32, q = 4, V_{rel}(\mathbf{\Sigma}) = 0.2$ | | | | | | | | |
| $N = 4$ | 0.5291 | 0.1397 | 0.5095 | 0.5278 | 0.1049 | −0.8747 | 0.9464 | 0.8860 |
| $N = 8$ | 0.3524 | 0.0674 | 0.3431 | 0.3521 | 0.0579 | −0.3826 | 1.0000 | 1.0000 |
| $N = 16$ | 0.2735 | 0.0326 | 0.2679 | 0.2727 | 0.0304 | −1.9324 | 1.0000 | 1.0000 |
| $N = 32$ | 0.2362 | 0.0161 | 0.2335 | 0.2357 | 0.0156 | −1.9882 | 1.0000 | 1.0000 |
| $N = 64$ | 0.2179 | 0.0081 | 0.2169 | 0.2179 | 0.0079 | −0.6360 | 1.0000 | 1.0000 |
| $N = 128$ | 0.2089 | 0.0042 | 0.2084 | 0.2090 | 0.0042 | 0.5130 | 1.0000 | 1.0000 |
| $N = 256$ | 0.2045 | 0.0022 | 0.2043 | 0.2045 | 0.0022 | 0.5881 | 1.0000 | 1.0000 |

(*continued*)

**Table S2.** (*continued*)

| | ≈ E[$V_{rel}$(**S**)] | ≈ SD[$V_{rel}$(**S**)] | Median | Mean | ESD | $T$ | Pow. 5% | Pow. 1% |
|---|---|---|---|---|---|---|---|---|
| $p = 64$, $q = 1$, $V_{rel}(\boldsymbol{\Sigma}) = 0.1$ | | | | | | | | |
| $N = 4$ | 0.3693 | 0.1331 | 0.3833 | 0.4049 | 0.0688 | 36.6235 | 0.6648 | 0.5412 |
| $N = 8$ | 0.2232 | 0.0871 | 0.2177 | 0.2312 | 0.0609 | 9.2822 | 0.9624 | 0.9330 |
| $N = 16$ | 0.1596 | 0.0565 | 0.1555 | 0.1614 | 0.0468 | 2.6219 | 1.0000 | 0.9996 |
| $N = 32$ | 0.1294 | 0.0378 | 0.1264 | 0.1301 | 0.0344 | 1.4771 | 1.0000 | 1.0000 |
| $N = 64$ | 0.1146 | 0.0260 | 0.1130 | 0.1146 | 0.0253 | −0.0493 | 1.0000 | 1.0000 |
| $N = 128$ | 0.1073 | 0.0181 | 0.1065 | 0.1073 | 0.0175 | −0.0405 | 1.0000 | 1.0000 |
| $N = 256$ | 0.1036 | 0.0127 | 0.1032 | 0.1037 | 0.0126 | 0.6022 | 1.0000 | 1.0000 |
| $p = 64$, $q = 1$, $V_{rel}(\boldsymbol{\Sigma}) = 0.2$ | | | | | | | | |
| $N = 4$ | 0.3788 | 0.1912 | 0.4356 | 0.4583 | 0.1014 | 55.4640 | 0.8090 | 0.7328 |
| $N = 8$ | 0.2881 | 0.1330 | 0.2971 | 0.3077 | 0.0917 | 15.0979 | 0.9922 | 0.9844 |
| $N = 16$ | 0.2450 | 0.0883 | 0.2472 | 0.2512 | 0.0732 | 6.0300 | 0.9998 | 0.9998 |
| $N = 32$ | 0.2227 | 0.0598 | 0.2228 | 0.2247 | 0.0546 | 2.5121 | 1.0000 | 1.0000 |
| $N = 64$ | 0.2114 | 0.0412 | 0.2098 | 0.2112 | 0.0392 | −0.3645 | 1.0000 | 1.0000 |
| $N = 128$ | 0.2057 | 0.0288 | 0.2046 | 0.2053 | 0.0285 | −1.0540 | 1.0000 | 1.0000 |
| $N = 256$ | 0.2029 | 0.0202 | 0.2025 | 0.2030 | 0.0200 | 0.2987 | 1.0000 | 1.0000 |
| $p = 64$, $q = 1$, $V_{rel}(\boldsymbol{\Sigma}) = 0.4$ | | | | | | | | |
| $N = 4$ | 0.4458 | 0.2090 | 0.5557 | 0.5594 | 0.1376 | 58.3687 | 0.9216 | 0.8926 |
| $N = 8$ | 0.4264 | 0.1613 | 0.4670 | 0.4599 | 0.1273 | 18.6017 | 0.9980 | 0.9974 |
| $N = 16$ | 0.4168 | 0.1119 | 0.4286 | 0.4266 | 0.0996 | 6.9651 | 1.0000 | 1.0000 |
| $N = 32$ | 0.4095 | 0.0773 | 0.4125 | 0.4109 | 0.0726 | 1.3497 | 1.0000 | 1.0000 |
| $N = 64$ | 0.4050 | 0.0538 | 0.4063 | 0.4056 | 0.0532 | 0.7287 | 1.0000 | 1.0000 |
| $N = 128$ | 0.4026 | 0.0377 | 0.4041 | 0.4032 | 0.0371 | 1.1293 | 1.0000 | 1.0000 |
| $N = 256$ | 0.4013 | 0.0266 | 0.4018 | 0.4015 | 0.0261 | 0.4964 | 1.0000 | 1.0000 |
| $p = 64$, $q = 1$, $V_{rel}(\boldsymbol{\Sigma}) = 0.6$ | | | | | | | | |
| $N = 4$ | 0.5867 | 0.1617 | 0.6883 | 0.6668 | 0.1510 | 37.5484 | 0.9718 | 0.9596 |
| $N = 8$ | 0.5911 | 0.1364 | 0.6354 | 0.6177 | 0.1314 | 14.3427 | 0.9998 | 0.9998 |
| $N = 16$ | 0.5983 | 0.0987 | 0.6151 | 0.6052 | 0.0974 | 5.0151 | 1.0000 | 1.0000 |
| $N = 32$ | 0.6002 | 0.0695 | 0.6085 | 0.6022 | 0.0693 | 2.0323 | 1.0000 | 1.0000 |
| $N = 64$ | 0.6004 | 0.0488 | 0.6039 | 0.6012 | 0.0484 | 1.1728 | 1.0000 | 1.0000 |
| $N = 128$ | 0.6003 | 0.0344 | 0.6015 | 0.6002 | 0.0342 | −0.0756 | 1.0000 | 1.0000 |
| $N = 256$ | 0.6001 | 0.0243 | 0.6009 | 0.6002 | 0.0241 | 0.2174 | 1.0000 | 1.0000 |
| $p = 64$, $q = 1$, $V_{rel}(\boldsymbol{\Sigma}) = 0.8$ | | | | | | | | |
| $N = 4$ | 0.7790 | 0.0864 | 0.8377 | 0.7928 | 0.1416 | 6.9078 | 0.9904 | 0.9854 |
| $N = 8$ | 0.7837 | 0.0787 | 0.8157 | 0.7915 | 0.1022 | 5.4258 | 1.0000 | 1.0000 |
| $N = 16$ | 0.7926 | 0.0592 | 0.8072 | 0.7956 | 0.0669 | 3.2349 | 1.0000 | 1.0000 |
| $N = 32$ | 0.7967 | 0.0425 | 0.8021 | 0.7967 | 0.0447 | −0.0106 | 1.0000 | 1.0000 |
| $N = 64$ | 0.7985 | 0.0301 | 0.8018 | 0.7989 | 0.0308 | 0.8231 | 1.0000 | 1.0000 |
| $N = 128$ | 0.7993 | 0.0213 | 0.8012 | 0.7999 | 0.0216 | 1.8961 | 1.0000 | 1.0000 |
| $N = 256$ | 0.7997 | 0.0151 | 0.8006 | 0.7996 | 0.0152 | −0.2806 | 1.0000 | 1.0000 |

(*continued*)

**Table S2.** (*continued*)

| | ≈ E[$V_{rel}$(**S**)] | ≈ SD[$V_{rel}$(**S**)] | Median | Mean | ESD | $T$ | Pow. 5% | Pow. 1% |
|---|---|---|---|---|---|---|---|---|
| $p = 64, q = 2, V_{rel}(\mathbf{\Sigma}) = 0.1$ | | | | | | | | |
| $N = 4$ | 0.4033 | 0.0983 | 0.4018 | 0.4170 | 0.0625 | 15.5134 | 0.8144 | 0.7052 |
| $N = 8$ | 0.2365 | 0.0577 | 0.2331 | 0.2394 | 0.0473 | 4.4814 | 0.9960 | 0.9914 |
| $N = 16$ | 0.1653 | 0.0352 | 0.1636 | 0.1662 | 0.0321 | 2.1126 | 1.0000 | 1.0000 |
| $N = 32$ | 0.1320 | 0.0227 | 0.1311 | 0.1319 | 0.0219 | −0.0435 | 1.0000 | 1.0000 |
| $N = 64$ | 0.1158 | 0.0153 | 0.1156 | 0.1159 | 0.0148 | 0.1625 | 1.0000 | 1.0000 |
| $N = 128$ | 0.1079 | 0.0105 | 0.1079 | 0.1080 | 0.0103 | 0.9248 | 1.0000 | 1.0000 |
| $N = 256$ | 0.1039 | 0.0073 | 0.1039 | 0.1040 | 0.0073 | 0.3798 | 1.0000 | 1.0000 |
| $p = 64, q = 2, V_{rel}(\mathbf{\Sigma}) = 0.2$ | | | | | | | | |
| $N = 4$ | 0.4672 | 0.1436 | 0.4719 | 0.4877 | 0.0920 | 15.6853 | 0.9534 | 0.9086 |
| $N = 8$ | 0.3244 | 0.0829 | 0.3238 | 0.3306 | 0.0666 | 6.5016 | 1.0000 | 1.0000 |
| $N = 16$ | 0.2607 | 0.0490 | 0.2589 | 0.2610 | 0.0449 | 0.4644 | 1.0000 | 1.0000 |
| $N = 32$ | 0.2300 | 0.0308 | 0.2295 | 0.2304 | 0.0297 | 0.7439 | 1.0000 | 1.0000 |
| $N = 64$ | 0.2149 | 0.0203 | 0.2148 | 0.2146 | 0.0197 | −1.1127 | 1.0000 | 1.0000 |
| $N = 128$ | 0.2075 | 0.0138 | 0.2075 | 0.2076 | 0.0138 | 0.9941 | 1.0000 | 1.0000 |
| $N = 256$ | 0.2037 | 0.0096 | 0.2038 | 0.2037 | 0.0095 | −0.4661 | 1.0000 | 1.0000 |
| $p = 64, q = 2, V_{rel}(\mathbf{\Sigma}) = 0.4$ | | | | | | | | |
| $N = 4$ | 0.6451 | 0.2025 | 0.6453 | 0.6520 | 0.1274 | 3.8135 | 0.9992 | 0.9986 |
| $N = 8$ | 0.5194 | 0.1086 | 0.4985 | 0.5178 | 0.0858 | −1.2882 | 1.0000 | 1.0000 |
| $N = 16$ | 0.4594 | 0.0563 | 0.4507 | 0.4604 | 0.0496 | 1.3591 | 1.0000 | 1.0000 |
| $N = 32$ | 0.4297 | 0.0299 | 0.4253 | 0.4303 | 0.0290 | 1.4675 | 1.0000 | 1.0000 |
| $N = 64$ | 0.4148 | 0.0165 | 0.4132 | 0.4149 | 0.0164 | 0.2356 | 1.0000 | 1.0000 |
| $N = 128$ | 0.4074 | 0.0097 | 0.4067 | 0.4072 | 0.0094 | −1.3898 | 1.0000 | 1.0000 |
| $N = 256$ | 0.4037 | 0.0060 | 0.4033 | 0.4036 | 0.0061 | −0.7394 | 1.0000 | 1.0000 |
| $p = 64, q = 4, V_{rel}(\mathbf{\Sigma}) = 0.1$ | | | | | | | | |
| $N = 4$ | 0.4254 | 0.0810 | 0.4164 | 0.4276 | 0.0588 | 2.6768 | 0.9004 | 0.8048 |
| $N = 8$ | 0.2456 | 0.0407 | 0.2411 | 0.2458 | 0.0350 | 0.4676 | 1.0000 | 1.0000 |
| $N = 16$ | 0.1693 | 0.0218 | 0.1676 | 0.1694 | 0.0203 | 0.4186 | 1.0000 | 1.0000 |
| $N = 32$ | 0.1338 | 0.0126 | 0.1333 | 0.1339 | 0.0123 | 0.2041 | 1.0000 | 1.0000 |
| $N = 64$ | 0.1167 | 0.0078 | 0.1167 | 0.1168 | 0.0079 | 0.6494 | 1.0000 | 1.0000 |
| $N = 128$ | 0.1083 | 0.0051 | 0.1083 | 0.1082 | 0.0051 | −1.1718 | 1.0000 | 1.0000 |
| $N = 256$ | 0.1041 | 0.0035 | 0.1042 | 0.1042 | 0.0035 | 0.2485 | 1.0000 | 1.0000 |
| $p = 64, q = 4, V_{rel}(\mathbf{\Sigma}) = 0.2$ | | | | | | | | |
| $N = 4$ | 0.5229 | 0.1312 | 0.5065 | 0.5241 | 0.0977 | 0.8973 | 0.9866 | 0.9726 |
| $N = 8$ | 0.3489 | 0.0633 | 0.3406 | 0.3487 | 0.0551 | −0.2548 | 1.0000 | 1.0000 |
| $N = 16$ | 0.2718 | 0.0307 | 0.2678 | 0.2719 | 0.0292 | 0.1980 | 1.0000 | 1.0000 |
| $N = 32$ | 0.2353 | 0.0153 | 0.2336 | 0.2353 | 0.0148 | 0.4010 | 1.0000 | 1.0000 |
| $N = 64$ | 0.2175 | 0.0079 | 0.2164 | 0.2175 | 0.0079 | 0.5288 | 1.0000 | 1.0000 |
| $N = 128$ | 0.2087 | 0.0042 | 0.2084 | 0.2088 | 0.0042 | 1.3044 | 1.0000 | 1.0000 |
| $N = 256$ | 0.2043 | 0.0024 | 0.2042 | 0.2043 | 0.0023 | 0.0456 | 1.0000 | 1.0000 |

(*continued*)

**Table S2.** (*continued*)

| | ≈ $E[V_{rel}(\mathbf{S})]$ | ≈ $SD[V_{rel}(\mathbf{S})]$ | Median | Mean | ESD | $T$ | Pow. 5% | Pow. 1% |
|---|---|---|---|---|---|---|---|---|
| $p = 128, q = 1, V_{rel}(\mathbf{\Sigma}) = 0.1$ | | | | | | | | |
| $N = 4$ | 0.3678 | 0.1298 | 0.3805 | 0.4016 | 0.0662 | 36.1631 | 0.7914 | 0.7176 |
| $N = 8$ | 0.2222 | 0.0851 | 0.2202 | 0.2319 | 0.0608 | 11.2526 | 0.9884 | 0.9816 |
| $N = 16$ | 0.1591 | 0.0552 | 0.1554 | 0.1613 | 0.0462 | 3.3949 | 1.0000 | 1.0000 |
| $N = 32$ | 0.1291 | 0.0370 | 0.1275 | 0.1301 | 0.0336 | 2.0940 | 1.0000 | 1.0000 |
| $N = 64$ | 0.1144 | 0.0254 | 0.1125 | 0.1141 | 0.0244 | −0.8564 | 1.0000 | 1.0000 |
| $N = 128$ | 0.1072 | 0.0177 | 0.1061 | 0.1070 | 0.0174 | −0.9694 | 1.0000 | 1.0000 |
| $N = 256$ | 0.1036 | 0.0124 | 0.1035 | 0.1036 | 0.0122 | −0.0474 | 1.0000 | 1.0000 |
| $p = 128, q = 1, V_{rel}(\mathbf{\Sigma}) = 0.2$ | | | | | | | | |
| $N = 4$ | 0.3779 | 0.1886 | 0.4357 | 0.4560 | 0.0968 | 56.9972 | 0.8926 | 0.8560 |
| $N = 8$ | 0.2876 | 0.1312 | 0.2974 | 0.3095 | 0.0922 | 16.8239 | 0.9976 | 0.9956 |
| $N = 16$ | 0.2446 | 0.0871 | 0.2447 | 0.2497 | 0.0738 | 4.8935 | 1.0000 | 1.0000 |
| $N = 32$ | 0.2226 | 0.0590 | 0.2220 | 0.2240 | 0.0534 | 1.8409 | 1.0000 | 1.0000 |
| $N = 64$ | 0.2114 | 0.0407 | 0.2111 | 0.2119 | 0.0385 | 1.0824 | 1.0000 | 1.0000 |
| $N = 128$ | 0.2057 | 0.0284 | 0.2055 | 0.2065 | 0.0279 | 1.9585 | 1.0000 | 1.0000 |
| $N = 256$ | 0.2028 | 0.0199 | 0.2022 | 0.2027 | 0.0197 | −0.3729 | 1.0000 | 1.0000 |
| $p = 128, q = 1, V_{rel}(\mathbf{\Sigma}) = 0.4$ | | | | | | | | |
| $N = 4$ | 0.4453 | 0.2074 | 0.5559 | 0.5594 | 0.1346 | 59.9265 | 0.9670 | 0.9496 |
| $N = 8$ | 0.4262 | 0.1601 | 0.4626 | 0.4578 | 0.1262 | 17.7102 | 0.9998 | 0.9996 |
| $N = 16$ | 0.4167 | 0.1110 | 0.4252 | 0.4220 | 0.0981 | 3.8079 | 1.0000 | 1.0000 |
| $N = 32$ | 0.4094 | 0.0766 | 0.4134 | 0.4124 | 0.0715 | 2.9484 | 1.0000 | 1.0000 |
| $N = 64$ | 0.4050 | 0.0534 | 0.4051 | 0.4053 | 0.0515 | 0.4527 | 1.0000 | 1.0000 |
| $N = 128$ | 0.4026 | 0.0374 | 0.4035 | 0.4024 | 0.0372 | −0.3393 | 1.0000 | 1.0000 |
| $N = 256$ | 0.4013 | 0.0263 | 0.4022 | 0.4021 | 0.0258 | 2.1191 | 1.0000 | 1.0000 |
| $p = 128, q = 1, V_{rel}(\mathbf{\Sigma}) = 0.6$ | | | | | | | | |
| $N = 4$ | 0.5864 | 0.1608 | 0.6876 | 0.6652 | 0.1523 | 36.6050 | 0.9842 | 0.9788 |
| $N = 8$ | 0.5910 | 0.1357 | 0.6346 | 0.6155 | 0.1327 | 13.0148 | 1.0000 | 1.0000 |
| $N = 16$ | 0.5983 | 0.0981 | 0.6162 | 0.6051 | 0.0963 | 5.0064 | 1.0000 | 1.0000 |
| $N = 32$ | 0.6001 | 0.0691 | 0.6069 | 0.6016 | 0.0696 | 1.4855 | 1.0000 | 1.0000 |
| $N = 64$ | 0.6004 | 0.0485 | 0.6044 | 0.6011 | 0.0485 | 1.0441 | 1.0000 | 1.0000 |
| $N = 128$ | 0.6003 | 0.0342 | 0.6023 | 0.6007 | 0.0339 | 0.8364 | 1.0000 | 1.0000 |
| $N = 256$ | 0.6001 | 0.0241 | 0.6010 | 0.6001 | 0.0241 | −0.1653 | 1.0000 | 1.0000 |
| $p = 128, q = 1, V_{rel}(\mathbf{\Sigma}) = 0.8$ | | | | | | | | |
| $N = 4$ | 0.7789 | 0.0859 | 0.8372 | 0.7960 | 0.1372 | 8.7820 | 0.9956 | 0.9936 |
| $N = 8$ | 0.7836 | 0.0783 | 0.8095 | 0.7880 | 0.1014 | 3.0677 | 1.0000 | 1.0000 |
| $N = 16$ | 0.7926 | 0.0589 | 0.8052 | 0.7934 | 0.0665 | 0.8762 | 1.0000 | 1.0000 |
| $N = 32$ | 0.7967 | 0.0423 | 0.8021 | 0.7970 | 0.0447 | 0.3646 | 1.0000 | 1.0000 |
| $N = 64$ | 0.7985 | 0.0300 | 0.8018 | 0.7991 | 0.0311 | 1.3807 | 1.0000 | 1.0000 |
| $N = 128$ | 0.7993 | 0.0212 | 0.8009 | 0.7996 | 0.0214 | 1.1186 | 1.0000 | 1.0000 |
| $N = 256$ | 0.7997 | 0.0150 | 0.8000 | 0.7993 | 0.0151 | −1.7751 | 1.0000 | 1.0000 |

(*continued*)

**Table S2.** (*continued*)

| | ≈ E[$V_{rel}$(**S**)] | ≈ SD[$V_{rel}$(**S**)] | Median | Mean | ESD | $T$ | Pow. 5% | Pow. 1% |
|---|---|---|---|---|---|---|---|---|
| $p$ = 128, $q$ = 2, $V_{rel}$(**Σ**) = 0.1 | | | | | | | | |
| $N$ = 4 | 0.4014 | 0.0943 | 0.3998 | 0.4133 | 0.0587 | 14.3539 | 0.9222 | 0.8732 |
| $N$ = 8 | 0.2353 | 0.0557 | 0.2325 | 0.2385 | 0.0452 | 4.9532 | 0.9994 | 0.9988 |
| $N$ = 16 | 0.1646 | 0.0341 | 0.1624 | 0.1650 | 0.0306 | 0.8469 | 1.0000 | 1.0000 |
| $N$ = 32 | 0.1316 | 0.0221 | 0.1302 | 0.1313 | 0.0207 | −1.2201 | 1.0000 | 1.0000 |
| $N$ = 64 | 0.1157 | 0.0149 | 0.1156 | 0.1160 | 0.0145 | 1.5574 | 1.0000 | 1.0000 |
| $N$ = 128 | 0.1078 | 0.0103 | 0.1076 | 0.1079 | 0.0102 | 0.9428 | 1.0000 | 1.0000 |
| $N$ = 256 | 0.1039 | 0.0072 | 0.1038 | 0.1039 | 0.0071 | 0.1376 | 1.0000 | 1.0000 |
| $p$ = 128, $q$ = 2, $V_{rel}$(**Σ**) = 0.2 | | | | | | | | |
| $N$ = 4 | 0.4656 | 0.1402 | 0.4709 | 0.4855 | 0.0889 | 15.7910 | 0.9894 | 0.9774 |
| $N$ = 8 | 0.3235 | 0.0812 | 0.3232 | 0.3291 | 0.0663 | 5.9794 | 1.0000 | 1.0000 |
| $N$ = 16 | 0.2602 | 0.0482 | 0.2586 | 0.2608 | 0.0442 | 1.0265 | 1.0000 | 1.0000 |
| $N$ = 32 | 0.2298 | 0.0303 | 0.2290 | 0.2301 | 0.0288 | 0.8073 | 1.0000 | 1.0000 |
| $N$ = 64 | 0.2148 | 0.0200 | 0.2149 | 0.2147 | 0.0199 | −0.2697 | 1.0000 | 1.0000 |
| $N$ = 128 | 0.2074 | 0.0137 | 0.2074 | 0.2074 | 0.0137 | 0.2450 | 1.0000 | 1.0000 |
| $N$ = 256 | 0.2037 | 0.0095 | 0.2039 | 0.2037 | 0.0095 | −0.0761 | 1.0000 | 1.0000 |
| $p$ = 128, $q$ = 2, $V_{rel}$(**Σ**) = 0.4 | | | | | | | | |
| $N$ = 4 | 0.6429 | 0.1994 | 0.6451 | 0.6482 | 0.1253 | 2.9897 | 0.9998 | 0.9990 |
| $N$ = 8 | 0.5182 | 0.1070 | 0.5029 | 0.5197 | 0.0852 | 1.2226 | 1.0000 | 1.0000 |
| $N$ = 16 | 0.4588 | 0.0557 | 0.4479 | 0.4586 | 0.0500 | −0.2879 | 1.0000 | 1.0000 |
| $N$ = 32 | 0.4294 | 0.0296 | 0.4251 | 0.4294 | 0.0286 | 0.0105 | 1.0000 | 1.0000 |
| $N$ = 64 | 0.4147 | 0.0165 | 0.4133 | 0.4148 | 0.0161 | 0.5386 | 1.0000 | 1.0000 |
| $N$ = 128 | 0.4073 | 0.0097 | 0.4069 | 0.4073 | 0.0098 | −0.1664 | 1.0000 | 1.0000 |
| $N$ = 256 | 0.4037 | 0.0061 | 0.4036 | 0.4037 | 0.0061 | 0.9004 | 1.0000 | 1.0000 |
| $p$ = 128, $q$ = 4, $V_{rel}$(**Σ**) = 0.1 | | | | | | | | |
| $N$ = 4 | 0.4230 | 0.0766 | 0.4169 | 0.4273 | 0.0555 | 5.4744 | 0.9766 | 0.9560 |
| $N$ = 8 | 0.2441 | 0.0387 | 0.2396 | 0.2444 | 0.0342 | 0.5399 | 1.0000 | 1.0000 |
| $N$ = 16 | 0.1685 | 0.0209 | 0.1674 | 0.1690 | 0.0196 | 1.6338 | 1.0000 | 1.0000 |
| $N$ = 32 | 0.1334 | 0.0122 | 0.1328 | 0.1334 | 0.0119 | −0.2881 | 1.0000 | 1.0000 |
| $N$ = 64 | 0.1165 | 0.0076 | 0.1164 | 0.1165 | 0.0076 | −0.6600 | 1.0000 | 1.0000 |
| $N$ = 128 | 0.1082 | 0.0050 | 0.1081 | 0.1082 | 0.0050 | −0.6723 | 1.0000 | 1.0000 |
| $N$ = 256 | 0.1041 | 0.0034 | 0.1040 | 0.1040 | 0.0034 | −1.7921 | 1.0000 | 1.0000 |
| $p$ = 128, $q$ = 4, $V_{rel}$(**Σ**) = 0.2 | | | | | | | | |
| $N$ = 4 | 0.5199 | 0.1270 | 0.5004 | 0.5186 | 0.0928 | −0.9804 | 0.9976 | 0.9954 |
| $N$ = 8 | 0.3473 | 0.0613 | 0.3401 | 0.3480 | 0.0527 | 0.9132 | 1.0000 | 1.0000 |
| $N$ = 16 | 0.2709 | 0.0298 | 0.2678 | 0.2714 | 0.0275 | 1.0597 | 1.0000 | 1.0000 |
| $N$ = 32 | 0.2348 | 0.0150 | 0.2329 | 0.2348 | 0.0145 | −0.2822 | 1.0000 | 1.0000 |
| $N$ = 64 | 0.2173 | 0.0078 | 0.2164 | 0.2172 | 0.0076 | −0.5165 | 1.0000 | 1.0000 |
| $N$ = 128 | 0.2086 | 0.0042 | 0.2084 | 0.2087 | 0.0042 | 0.9662 | 1.0000 | 1.0000 |
| $N$ = 256 | 0.2043 | 0.0024 | 0.2042 | 0.2043 | 0.0024 | −0.3874 | 1.0000 | 1.0000 |

(*continued*)

**Table S2.** (*continued*)

| | ≈ E[$V_{rel}$(**S**)] | ≈ SD[$V_{rel}$(**S**)] | Median | Mean | ESD | $T$ | Pow. 5% | Pow. 1% |
|---|---|---|---|---|---|---|---|---|
| $p = 256, q = 1, V_{rel}(\mathbf{\Sigma}) = 0.1$ | | | | | | | | |
| $N = 4$ | 0.3670 | 0.1282 | 0.3802 | 0.4010 | 0.0649 | 36.9664 | 0.8680 | 0.8214 |
| $N = 8$ | 0.2217 | 0.0841 | 0.2161 | 0.2282 | 0.0586 | 7.8635 | 0.9948 | 0.9910 |
| $N = 16$ | 0.1588 | 0.0546 | 0.1547 | 0.1603 | 0.0443 | 2.3327 | 1.0000 | 1.0000 |
| $N = 32$ | 0.1289 | 0.0366 | 0.1252 | 0.1283 | 0.0328 | −1.3113 | 1.0000 | 1.0000 |
| $N = 64$ | 0.1144 | 0.0252 | 0.1131 | 0.1145 | 0.0238 | 0.5640 | 1.0000 | 1.0000 |
| $N = 128$ | 0.1072 | 0.0175 | 0.1067 | 0.1072 | 0.0171 | 0.0656 | 1.0000 | 1.0000 |
| $N = 256$ | 0.1036 | 0.0123 | 0.1029 | 0.1035 | 0.0123 | −0.2719 | 1.0000 | 1.0000 |
| $p = 256, q = 1, V_{rel}(\mathbf{\Sigma}) = 0.2$ | | | | | | | | |
| $N = 4$ | 0.3775 | 0.1873 | 0.4345 | 0.4545 | 0.0960 | 56.7566 | 0.9408 | 0.9150 |
| $N = 8$ | 0.2873 | 0.1303 | 0.2947 | 0.3059 | 0.0906 | 14.5168 | 0.9998 | 0.9996 |
| $N = 16$ | 0.2445 | 0.0865 | 0.2439 | 0.2494 | 0.0717 | 4.8522 | 1.0000 | 1.0000 |
| $N = 32$ | 0.2225 | 0.0586 | 0.2207 | 0.2230 | 0.0532 | 0.7110 | 1.0000 | 1.0000 |
| $N = 64$ | 0.2113 | 0.0404 | 0.2113 | 0.2117 | 0.0393 | 0.7681 | 1.0000 | 1.0000 |
| $N = 128$ | 0.2057 | 0.0282 | 0.2060 | 0.2063 | 0.0273 | 1.6400 | 1.0000 | 1.0000 |
| $N = 256$ | 0.2028 | 0.0198 | 0.2025 | 0.2030 | 0.0195 | 0.6016 | 1.0000 | 1.0000 |
| $p = 256, q = 1, V_{rel}(\mathbf{\Sigma}) = 0.4$ | | | | | | | | |
| $N = 4$ | 0.4451 | 0.2066 | 0.5553 | 0.5569 | 0.1336 | 59.1992 | 0.9778 | 0.9692 |
| $N = 8$ | 0.4261 | 0.1594 | 0.4637 | 0.4606 | 0.1249 | 19.5222 | 1.0000 | 1.0000 |
| $N = 16$ | 0.4166 | 0.1105 | 0.4262 | 0.4228 | 0.0977 | 4.4222 | 1.0000 | 1.0000 |
| $N = 32$ | 0.4094 | 0.0763 | 0.4148 | 0.4120 | 0.0718 | 2.5791 | 1.0000 | 1.0000 |
| $N = 64$ | 0.4050 | 0.0531 | 0.4066 | 0.4058 | 0.0520 | 1.1405 | 1.0000 | 1.0000 |
| $N = 128$ | 0.4026 | 0.0373 | 0.4017 | 0.4021 | 0.0365 | −0.8094 | 1.0000 | 1.0000 |
| $N = 256$ | 0.4013 | 0.0262 | 0.4020 | 0.4016 | 0.0260 | 0.7194 | 1.0000 | 1.0000 |
| $p = 256, q = 1, V_{rel}(\mathbf{\Sigma}) = 0.6$ | | | | | | | | |
| $N = 4$ | 0.5863 | 0.1603 | 0.6891 | 0.6644 | 0.1489 | 37.1120 | 0.9924 | 0.9878 |
| $N = 8$ | 0.5910 | 0.1353 | 0.6340 | 0.6178 | 0.1277 | 14.8192 | 0.9998 | 0.9998 |
| $N = 16$ | 0.5983 | 0.0978 | 0.6147 | 0.6052 | 0.0952 | 5.1681 | 1.0000 | 1.0000 |
| $N = 32$ | 0.6001 | 0.0689 | 0.6085 | 0.6029 | 0.0685 | 2.8470 | 1.0000 | 1.0000 |
| $N = 64$ | 0.6004 | 0.0484 | 0.6027 | 0.5997 | 0.0483 | −0.9421 | 1.0000 | 1.0000 |
| $N = 128$ | 0.6003 | 0.0341 | 0.6019 | 0.6002 | 0.0342 | −0.0204 | 1.0000 | 1.0000 |
| $N = 256$ | 0.6001 | 0.0240 | 0.6008 | 0.6002 | 0.0240 | 0.0870 | 1.0000 | 1.0000 |
| $p = 256, q = 1, V_{rel}(\mathbf{\Sigma}) = 0.8$ | | | | | | | | |
| $N = 4$ | 0.7789 | 0.0857 | 0.8347 | 0.7932 | 0.1395 | 7.2244 | 0.9968 | 0.9958 |
| $N = 8$ | 0.7836 | 0.0781 | 0.8150 | 0.7930 | 0.0994 | 6.6508 | 1.0000 | 1.0000 |
| $N = 16$ | 0.7926 | 0.0588 | 0.8058 | 0.7944 | 0.0663 | 1.9392 | 1.0000 | 1.0000 |
| $N = 32$ | 0.7967 | 0.0422 | 0.8033 | 0.7976 | 0.0455 | 1.3138 | 1.0000 | 1.0000 |
| $N = 64$ | 0.7985 | 0.0299 | 0.8010 | 0.7983 | 0.0303 | −0.5863 | 1.0000 | 1.0000 |
| $N = 128$ | 0.7993 | 0.0212 | 0.8010 | 0.7995 | 0.0212 | 0.7960 | 1.0000 | 1.0000 |
| $N = 256$ | 0.7997 | 0.0150 | 0.8004 | 0.7998 | 0.0153 | 0.5080 | 1.0000 | 1.0000 |

(*continued*)

**Table S2.** (*continued*)

| | ≈ E[$V_{rel}$(**S**)] | ≈ SD[$V_{rel}$(**S**)] | Median | Mean | ESD | $T$ | Pow. 5% | Pow. 1% |
|---|---|---|---|---|---|---|---|---|
| $p = 256, q = 2, V_{rel}(\mathbf{\Sigma}) = 0.1$ | | | | | | | | |
| $N = 4$ | 0.4005 | 0.0923 | 0.3999 | 0.4120 | 0.0570 | 14.2491 | 0.9688 | 0.9502 |
| $N = 8$ | 0.2347 | 0.0547 | 0.2321 | 0.2380 | 0.0440 | 5.3249 | 1.0000 | 1.0000 |
| $N = 16$ | 0.1643 | 0.0336 | 0.1618 | 0.1640 | 0.0302 | −0.6560 | 1.0000 | 1.0000 |
| $N = 32$ | 0.1315 | 0.0218 | 0.1301 | 0.1312 | 0.0204 | −0.8040 | 1.0000 | 1.0000 |
| $N = 64$ | 0.1156 | 0.0147 | 0.1149 | 0.1157 | 0.0144 | 0.4099 | 1.0000 | 1.0000 |
| $N = 128$ | 0.1077 | 0.0101 | 0.1074 | 0.1076 | 0.0101 | −0.9266 | 1.0000 | 1.0000 |
| $N = 256$ | 0.1039 | 0.0071 | 0.1039 | 0.1040 | 0.0071 | 1.4076 | 1.0000 | 1.0000 |
| $p = 256, q = 2, V_{rel}(\mathbf{\Sigma}) = 0.2$ | | | | | | | | |
| $N = 4$ | 0.4648 | 0.1385 | 0.4678 | 0.4835 | 0.0865 | 15.2564 | 0.9948 | 0.9912 |
| $N = 8$ | 0.3230 | 0.0803 | 0.3213 | 0.3272 | 0.0645 | 4.5921 | 1.0000 | 1.0000 |
| $N = 16$ | 0.2599 | 0.0478 | 0.2587 | 0.2606 | 0.0430 | 1.0712 | 1.0000 | 1.0000 |
| $N = 32$ | 0.2296 | 0.0301 | 0.2296 | 0.2295 | 0.0289 | −0.2494 | 1.0000 | 1.0000 |
| $N = 64$ | 0.2147 | 0.0199 | 0.2145 | 0.2146 | 0.0197 | −0.4946 | 1.0000 | 1.0000 |
| $N = 128$ | 0.2074 | 0.0136 | 0.2074 | 0.2076 | 0.0133 | 1.2036 | 1.0000 | 1.0000 |
| $N = 256$ | 0.2037 | 0.0094 | 0.2037 | 0.2036 | 0.0094 | −0.3174 | 1.0000 | 1.0000 |
| $p = 256, q = 2, V_{rel}(\mathbf{\Sigma}) = 0.4$ | | | | | | | | |
| $N = 4$ | 0.6418 | 0.1979 | 0.6414 | 0.6484 | 0.1237 | 3.7632 | 0.9998 | 0.9996 |
| $N = 8$ | 0.5176 | 0.1062 | 0.4989 | 0.5166 | 0.0846 | −0.8435 | 1.0000 | 1.0000 |
| $N = 16$ | 0.4585 | 0.0553 | 0.4502 | 0.4599 | 0.0496 | 2.0152 | 1.0000 | 1.0000 |
| $N = 32$ | 0.4292 | 0.0295 | 0.4255 | 0.4294 | 0.0276 | 0.4661 | 1.0000 | 1.0000 |
| $N = 64$ | 0.4146 | 0.0165 | 0.4129 | 0.4145 | 0.0159 | −0.5531 | 1.0000 | 1.0000 |
| $N = 128$ | 0.4073 | 0.0097 | 0.4068 | 0.4072 | 0.0096 | −0.8852 | 1.0000 | 1.0000 |
| $N = 256$ | 0.4036 | 0.0061 | 0.4037 | 0.4037 | 0.0062 | 1.0291 | 1.0000 | 1.0000 |
| $p = 256, q = 4, V_{rel}(\mathbf{\Sigma}) = 0.1$ | | | | | | | | |
| $N = 4$ | 0.4218 | 0.0744 | 0.4150 | 0.4253 | 0.0544 | 4.6252 | 0.9958 | 0.9918 |
| $N = 8$ | 0.2434 | 0.0377 | 0.2390 | 0.2431 | 0.0321 | −0.6534 | 1.0000 | 1.0000 |
| $N = 16$ | 0.1682 | 0.0204 | 0.1663 | 0.1679 | 0.0187 | −1.0115 | 1.0000 | 1.0000 |
| $N = 32$ | 0.1333 | 0.0120 | 0.1324 | 0.1331 | 0.0116 | −1.1964 | 1.0000 | 1.0000 |
| $N = 64$ | 0.1164 | 0.0075 | 0.1164 | 0.1165 | 0.0075 | 1.0008 | 1.0000 | 1.0000 |
| $N = 128$ | 0.1082 | 0.0050 | 0.1082 | 0.1081 | 0.0049 | −0.3841 | 1.0000 | 1.0000 |
| $N = 256$ | 0.1041 | 0.0034 | 0.1041 | 0.1041 | 0.0034 | 1.4474 | 1.0000 | 1.0000 |
| $p = 256, q = 4, V_{rel}(\mathbf{\Sigma}) = 0.2$ | | | | | | | | |
| $N = 4$ | 0.5184 | 0.1250 | 0.5027 | 0.5188 | 0.0921 | 0.2504 | 0.9990 | 0.9986 |
| $N = 8$ | 0.3465 | 0.0603 | 0.3376 | 0.3459 | 0.0517 | −0.8305 | 1.0000 | 1.0000 |
| $N = 16$ | 0.2705 | 0.0294 | 0.2673 | 0.2711 | 0.0278 | 1.3770 | 1.0000 | 1.0000 |
| $N = 32$ | 0.2346 | 0.0148 | 0.2332 | 0.2350 | 0.0146 | 2.0066 | 1.0000 | 1.0000 |
| $N = 64$ | 0.2172 | 0.0078 | 0.2164 | 0.2171 | 0.0076 | −0.2420 | 1.0000 | 1.0000 |
| $N = 128$ | 0.2085 | 0.0043 | 0.2082 | 0.2086 | 0.0043 | 0.4033 | 1.0000 | 1.0000 |
| $N = 256$ | 0.2043 | 0.0025 | 0.2042 | 0.2043 | 0.0025 | 0.7756 | 1.0000 | 1.0000 |

(*continued*)

**Table S2.** (*continued*)

| | ≈ E[$V_{rel}$(**S**)] | ≈ SD[$V_{rel}$(**S**)] | Median | Mean | ESD | $T$ | Pow. 5% | Pow. 1% |
|---|---|---|---|---|---|---|---|---|
| $p = 1024$, $q = 1$, $V_{rel}(\boldsymbol{\Sigma}) = 0.1$ | | | | | | | | |
| $N = 4$ | 0.3664 | 0.1269 | 0.3824 | 0.4005 | 0.0628 | 38.3853 | 0.9530 | 0.9368 |
| $N = 8$ | 0.2214 | 0.0834 | 0.2187 | 0.2300 | 0.0594 | 10.3495 | 0.9996 | 0.9992 |
| $N = 16$ | 0.1586 | 0.0541 | 0.1546 | 0.1604 | 0.0450 | 2.7974 | 1.0000 | 1.0000 |
| $N = 32$ | 0.1288 | 0.0363 | 0.1252 | 0.1288 | 0.0327 | −0.1243 | 1.0000 | 1.0000 |
| $N = 64$ | 0.1143 | 0.0250 | 0.1129 | 0.1144 | 0.0239 | 0.3962 | 1.0000 | 1.0000 |
| $N = 128$ | 0.1071 | 0.0174 | 0.1061 | 0.1068 | 0.0168 | −1.3612 | 1.0000 | 1.0000 |
| $N = 256$ | 0.1036 | 0.0122 | 0.1030 | 0.1034 | 0.0121 | −0.9759 | 1.0000 | 1.0000 |
| $p = 1024$, $q = 1$, $V_{rel}(\boldsymbol{\Sigma}) = 0.2$ | | | | | | | | |
| $N = 4$ | 0.3772 | 0.1863 | 0.4316 | 0.4538 | 0.0966 | 56.1171 | 0.9792 | 0.9712 |
| $N = 8$ | 0.2871 | 0.1296 | 0.2968 | 0.3061 | 0.0902 | 14.9086 | 1.0000 | 1.0000 |
| $N = 16$ | 0.2443 | 0.0860 | 0.2460 | 0.2502 | 0.0711 | 5.8614 | 1.0000 | 1.0000 |
| $N = 32$ | 0.2224 | 0.0583 | 0.2206 | 0.2229 | 0.0530 | 0.6929 | 1.0000 | 1.0000 |
| $N = 64$ | 0.2113 | 0.0402 | 0.2110 | 0.2121 | 0.0385 | 1.4655 | 1.0000 | 1.0000 |
| $N = 128$ | 0.2057 | 0.0281 | 0.2044 | 0.2049 | 0.0273 | −2.0171 | 1.0000 | 1.0000 |
| $N = 256$ | 0.2028 | 0.0197 | 0.2019 | 0.2022 | 0.0194 | −2.1194 | 1.0000 | 1.0000 |
| $p = 1024$, $q = 1$, $V_{rel}(\boldsymbol{\Sigma}) = 0.4$ | | | | | | | | |
| $N = 4$ | 0.4449 | 0.2060 | 0.5508 | 0.5536 | 0.1326 | 57.9415 | 0.9934 | 0.9894 |
| $N = 8$ | 0.4261 | 0.1590 | 0.4566 | 0.4548 | 0.1261 | 16.1041 | 1.0000 | 1.0000 |
| $N = 16$ | 0.4166 | 0.1102 | 0.4272 | 0.4246 | 0.0990 | 5.7314 | 1.0000 | 1.0000 |
| $N = 32$ | 0.4094 | 0.0761 | 0.4140 | 0.4123 | 0.0718 | 2.8690 | 1.0000 | 1.0000 |
| $N = 64$ | 0.4050 | 0.0530 | 0.4068 | 0.4058 | 0.0521 | 1.1494 | 1.0000 | 1.0000 |
| $N = 128$ | 0.4026 | 0.0371 | 0.4024 | 0.4024 | 0.0366 | −0.2967 | 1.0000 | 1.0000 |
| $N = 256$ | 0.4013 | 0.0261 | 0.4024 | 0.4016 | 0.0263 | 0.7143 | 1.0000 | 1.0000 |
| $p = 1024$, $q = 1$, $V_{rel}(\boldsymbol{\Sigma}) = 0.6$ | | | | | | | | |
| $N = 4$ | 0.5862 | 0.1600 | 0.6832 | 0.6637 | 0.1492 | 36.7630 | 0.9966 | 0.9950 |
| $N = 8$ | 0.5910 | 0.1350 | 0.6344 | 0.6160 | 0.1281 | 13.8435 | 1.0000 | 1.0000 |
| $N = 16$ | 0.5983 | 0.0976 | 0.6170 | 0.6051 | 0.0949 | 5.0647 | 1.0000 | 1.0000 |
| $N = 32$ | 0.6001 | 0.0687 | 0.6076 | 0.6012 | 0.0692 | 1.0874 | 1.0000 | 1.0000 |
| $N = 64$ | 0.6004 | 0.0483 | 0.6040 | 0.6005 | 0.0476 | 0.1575 | 1.0000 | 1.0000 |
| $N = 128$ | 0.6003 | 0.0340 | 0.6016 | 0.6005 | 0.0334 | 0.5738 | 1.0000 | 1.0000 |
| $N = 256$ | 0.6001 | 0.0240 | 0.6006 | 0.6000 | 0.0240 | −0.4044 | 1.0000 | 1.0000 |
| $p = 1024$, $q = 1$, $V_{rel}(\boldsymbol{\Sigma}) = 0.8$ | | | | | | | | |
| $N = 4$ | 0.7789 | 0.0855 | 0.8366 | 0.7933 | 0.1392 | 7.3423 | 0.9990 | 0.9986 |
| $N = 8$ | 0.7836 | 0.0780 | 0.8140 | 0.7904 | 0.1011 | 4.7652 | 1.0000 | 1.0000 |
| $N = 16$ | 0.7926 | 0.0587 | 0.8081 | 0.7959 | 0.0665 | 3.5530 | 1.0000 | 1.0000 |
| $N = 32$ | 0.7967 | 0.0421 | 0.8026 | 0.7967 | 0.0448 | −0.0218 | 1.0000 | 1.0000 |
| $N = 64$ | 0.7985 | 0.0299 | 0.8012 | 0.7987 | 0.0307 | 0.4338 | 1.0000 | 1.0000 |
| $N = 128$ | 0.7993 | 0.0211 | 0.8009 | 0.7991 | 0.0220 | −0.6352 | 1.0000 | 1.0000 |
| $N = 256$ | 0.7997 | 0.0149 | 0.7996 | 0.7991 | 0.0154 | −2.7295 | 1.0000 | 1.0000 |

(*continued*)

**Table S2.** (*continued*)

| | ≈ $E[V_{rel}(\mathbf{S})]$ | ≈ $SD[V_{rel}(\mathbf{S})]$ | Median | Mean | ESD | $T$ | Pow. 5% | Pow. 1% |
|---|---|---|---|---|---|---|---|---|
| $p = 1024, q = 2, V_{rel}(\mathbf{\Sigma}) = 0.1$ | | | | | | | | |
| $N = 4$ | 0.3998 | 0.0908 | 0.4004 | 0.4135 | 0.0570 | 16.9530 | 0.9966 | 0.9932 |
| $N = 8$ | 0.2343 | 0.0540 | 0.2310 | 0.2367 | 0.0428 | 3.9784 | 1.0000 | 1.0000 |
| $N = 16$ | 0.1641 | 0.0332 | 0.1618 | 0.1645 | 0.0300 | 0.8667 | 1.0000 | 1.0000 |
| $N = 32$ | 0.1314 | 0.0216 | 0.1304 | 0.1312 | 0.0205 | −0.3665 | 1.0000 | 1.0000 |
| $N = 64$ | 0.1155 | 0.0146 | 0.1157 | 0.1158 | 0.0143 | 1.6711 | 1.0000 | 1.0000 |
| $N = 128$ | 0.1077 | 0.0101 | 0.1076 | 0.1078 | 0.0098 | 0.6646 | 1.0000 | 1.0000 |
| $N = 256$ | 0.1038 | 0.0070 | 0.1038 | 0.1039 | 0.0070 | 0.2842 | 1.0000 | 1.0000 |
| $p = 1024, q = 2, V_{rel}(\mathbf{\Sigma}) = 0.2$ | | | | | | | | |
| $N = 4$ | 0.4642 | 0.1372 | 0.4699 | 0.4843 | 0.0868 | 16.3768 | 0.9992 | 0.9988 |
| $N = 8$ | 0.3226 | 0.0797 | 0.3230 | 0.3279 | 0.0648 | 5.7031 | 1.0000 | 1.0000 |
| $N = 16$ | 0.2597 | 0.0474 | 0.2583 | 0.2605 | 0.0433 | 1.1566 | 1.0000 | 1.0000 |
| $N = 32$ | 0.2295 | 0.0300 | 0.2301 | 0.2302 | 0.0291 | 1.5109 | 1.0000 | 1.0000 |
| $N = 64$ | 0.2147 | 0.0198 | 0.2143 | 0.2143 | 0.0195 | −1.2399 | 1.0000 | 1.0000 |
| $N = 128$ | 0.2073 | 0.0135 | 0.2074 | 0.2073 | 0.0132 | −0.3207 | 1.0000 | 1.0000 |
| $N = 256$ | 0.2037 | 0.0094 | 0.2039 | 0.2038 | 0.0093 | 0.8660 | 1.0000 | 1.0000 |
| $p = 1024, q = 2, V_{rel}(\mathbf{\Sigma}) = 0.4$ | | | | | | | | |
| $N = 4$ | 0.6410 | 0.1968 | 0.6318 | 0.6431 | 0.1222 | 1.2092 | 1.0000 | 1.0000 |
| $N = 8$ | 0.5171 | 0.1057 | 0.5005 | 0.5176 | 0.0835 | 0.3728 | 1.0000 | 1.0000 |
| $N = 16$ | 0.4583 | 0.0551 | 0.4494 | 0.4586 | 0.0495 | 0.5014 | 1.0000 | 1.0000 |
| $N = 32$ | 0.4291 | 0.0294 | 0.4251 | 0.4290 | 0.0284 | −0.1381 | 1.0000 | 1.0000 |
| $N = 64$ | 0.4145 | 0.0165 | 0.4133 | 0.4146 | 0.0159 | 0.1152 | 1.0000 | 1.0000 |
| $N = 128$ | 0.4073 | 0.0098 | 0.4072 | 0.4075 | 0.0096 | 1.4718 | 1.0000 | 1.0000 |
| $N = 256$ | 0.4036 | 0.0061 | 0.4035 | 0.4037 | 0.0061 | 0.6990 | 1.0000 | 1.0000 |
| $p = 1024, q = 4, V_{rel}(\mathbf{\Sigma}) = 0.1$ | | | | | | | | |
| $N = 4$ | 0.4209 | 0.0727 | 0.4147 | 0.4244 | 0.0526 | 4.6637 | 0.9998 | 0.9998 |
| $N = 8$ | 0.2429 | 0.0370 | 0.2400 | 0.2444 | 0.0329 | 3.3100 | 1.0000 | 1.0000 |
| $N = 16$ | 0.1679 | 0.0201 | 0.1664 | 0.1681 | 0.0194 | 1.0058 | 1.0000 | 1.0000 |
| $N = 32$ | 0.1331 | 0.0118 | 0.1324 | 0.1329 | 0.0117 | −1.3092 | 1.0000 | 1.0000 |
| $N = 64$ | 0.1164 | 0.0075 | 0.1161 | 0.1162 | 0.0074 | −1.8472 | 1.0000 | 1.0000 |
| $N = 128$ | 0.1081 | 0.0049 | 0.1082 | 0.1082 | 0.0050 | 0.8669 | 1.0000 | 1.0000 |
| $N = 256$ | 0.1041 | 0.0034 | 0.1041 | 0.1041 | 0.0033 | 0.3270 | 1.0000 | 1.0000 |
| $p = 1024, q = 4, V_{rel}(\mathbf{\Sigma}) = 0.2$ | | | | | | | | |
| $N = 4$ | 0.5174 | 0.1234 | 0.5036 | 0.5197 | 0.0916 | 1.7658 | 1.0000 | 1.0000 |
| $N = 8$ | 0.3459 | 0.0596 | 0.3379 | 0.3465 | 0.0516 | 0.8457 | 1.0000 | 1.0000 |
| $N = 16$ | 0.2702 | 0.0291 | 0.2672 | 0.2711 | 0.0272 | 2.2078 | 1.0000 | 1.0000 |
| $N = 32$ | 0.2345 | 0.0147 | 0.2329 | 0.2345 | 0.0142 | −0.0558 | 1.0000 | 1.0000 |
| $N = 64$ | 0.2171 | 0.0077 | 0.2164 | 0.2171 | 0.0076 | −0.3159 | 1.0000 | 1.0000 |
| $N = 128$ | 0.2085 | 0.0043 | 0.2082 | 0.2085 | 0.0042 | −0.4645 | 1.0000 | 1.0000 |
| $N = 256$ | 0.2042 | 0.0025 | 0.2041 | 0.2042 | 0.0025 | −0.3507 | 1.0000 | 1.0000 |

(*continued*)

**Table S2.** (*continued*)

| | $\approx E[V_{rel}(\mathbf{S})]$ | $\approx SD[V_{rel}(\mathbf{S})]$ | Median | Mean | ESD | $T$ | Pow. 5% | Pow. 1% |
|---|---|---|---|---|---|---|---|---|
| $p = 2$, linearly decreasing λ ($V_{rel}(\mathbf{\Sigma}) = 0.1111$) | | | | | | | | |
| $N = 4$ | 0.4433 | 0.4624 | 0.5454 | 0.5308 | 0.2864 | 21.5974 | 0.0550 | 0.0116 |
| $N = 8$ | 0.2822 | 0.2964 | 0.2773 | 0.3101 | 0.2141 | 9.2064 | 0.0964 | 0.0266 |
| $N = 16$ | 0.1998 | 0.1848 | 0.1783 | 0.2080 | 0.1505 | 3.8473 | 0.1900 | 0.0548 |
| $N = 32$ | 0.1564 | 0.1190 | 0.1422 | 0.1606 | 0.1065 | 2.8070 | 0.3726 | 0.1508 |
| $N = 64$ | 0.1340 | 0.0794 | 0.1260 | 0.1346 | 0.0747 | 0.6060 | 0.6880 | 0.4968 |
| $N = 128$ | 0.1226 | 0.0543 | 0.1191 | 0.1235 | 0.0534 | 1.1103 | 0.9492 | 0.8676 |
| $N = 256$ | 0.1169 | 0.0377 | 0.1145 | 0.1167 | 0.0369 | −0.3463 | 0.9990 | 0.9940 |
| $p = 4$, linearly decreasing λ ($V_{rel}(\mathbf{\Sigma}) = 0.0667$) | | | | | | | | |
| $N = 4$ | 0.4440 | 0.2334 | 0.4352 | 0.4602 | 0.1600 | 7.1593 | 0.0810 | 0.0180 |
| $N = 8$ | 0.2464 | 0.1278 | 0.2350 | 0.2515 | 0.1022 | 3.5335 | 0.1358 | 0.0408 |
| $N = 16$ | 0.1549 | 0.0713 | 0.1466 | 0.1551 | 0.0626 | 0.3162 | 0.3068 | 0.1148 |
| $N = 32$ | 0.1104 | 0.0423 | 0.1044 | 0.1098 | 0.0396 | −1.0912 | 0.6722 | 0.3788 |
| $N = 64$ | 0.0884 | 0.0267 | 0.0863 | 0.0890 | 0.0258 | 1.4457 | 0.9832 | 0.9172 |
| $N = 128$ | 0.0775 | 0.0176 | 0.0760 | 0.0775 | 0.0174 | −0.2476 | 1.0000 | 1.0000 |
| $N = 256$ | 0.0721 | 0.0120 | 0.0718 | 0.0724 | 0.0117 | 1.6817 | 1.0000 | 1.0000 |
| $p = 8$, linearly decreasing λ ($V_{rel}(\mathbf{\Sigma}) = 0.0370$) | | | | | | | | |
| $N = 4$ | 0.4089 | 0.1184 | 0.3958 | 0.4129 | 0.0932 | 2.9996 | 0.1068 | 0.0292 |
| $N = 8$ | 0.2058 | 0.0584 | 0.1989 | 0.2060 | 0.0510 | 0.2640 | 0.1982 | 0.0808 |
| $N = 16$ | 0.1178 | 0.0302 | 0.1139 | 0.1176 | 0.0282 | −0.4178 | 0.4668 | 0.2476 |
| $N = 32$ | 0.0766 | 0.0168 | 0.0754 | 0.0771 | 0.0165 | 2.2504 | 0.9060 | 0.7168 |
| $N = 64$ | 0.0566 | 0.0100 | 0.0559 | 0.0567 | 0.0098 | 0.6109 | 1.0000 | 1.0000 |
| $N = 128$ | 0.0468 | 0.0063 | 0.0463 | 0.0467 | 0.0062 | −1.1788 | 1.0000 | 1.0000 |
| $N = 256$ | 0.0419 | 0.0042 | 0.0416 | 0.0420 | 0.0042 | 1.1985 | 1.0000 | 1.0000 |
| $p = 16$, linearly decreasing λ ($V_{rel}(\mathbf{\Sigma}) = 0.0196$) | | | | | | | | |
| $N = 4$ | 0.3773 | 0.0593 | 0.3693 | 0.3780 | 0.0526 | 1.0046 | 0.1246 | 0.0366 |
| $N = 8$ | 0.1776 | 0.0274 | 0.1747 | 0.1776 | 0.0251 | 0.0584 | 0.2290 | 0.0860 |
| $N = 16$ | 0.0942 | 0.0134 | 0.0932 | 0.0945 | 0.0132 | 1.1528 | 0.5654 | 0.3212 |
| $N = 32$ | 0.0559 | 0.0071 | 0.0555 | 0.0560 | 0.0071 | 0.9064 | 0.9740 | 0.8924 |
| $N = 64$ | 0.0375 | 0.0040 | 0.0372 | 0.0375 | 0.0040 | 0.0096 | 1.0000 | 1.0000 |
| $N = 128$ | 0.0285 | 0.0024 | 0.0284 | 0.0285 | 0.0024 | −0.6234 | 1.0000 | 1.0000 |
| $N = 256$ | 0.0240 | 0.0015 | 0.0240 | 0.0240 | 0.0015 | 0.0265 | 1.0000 | 1.0000 |
| $p = 32$, linearly decreasing λ ($V_{rel}(\mathbf{\Sigma}) = 0.0101$) | | | | | | | | |
| $N = 4$ | 0.3570 | 0.0295 | 0.3524 | 0.3575 | 0.0271 | 1.2012 | 0.1346 | 0.0410 |
| $N = 8$ | 0.1611 | 0.0131 | 0.1595 | 0.1608 | 0.0125 | −1.6115 | 0.2570 | 0.0966 |
| $N = 16$ | 0.0810 | 0.0063 | 0.0804 | 0.0809 | 0.0061 | −1.1890 | 0.6594 | 0.3810 |
| $N = 32$ | 0.0445 | 0.0032 | 0.0443 | 0.0445 | 0.0031 | −0.6711 | 0.9894 | 0.9498 |
| $N = 64$ | 0.0271 | 0.0017 | 0.0270 | 0.0271 | 0.0017 | 0.5277 | 1.0000 | 1.0000 |
| $N = 128$ | 0.0185 | 0.0010 | 0.0185 | 0.0185 | 0.0010 | 0.8847 | 1.0000 | 1.0000 |
| $N = 256$ | 0.0143 | 0.0006 | 0.0143 | 0.0143 | 0.0006 | −0.1711 | 1.0000 | 1.0000 |

(*continued*)

**Table S2.** (*continued*)

| | ≈ E[$V_{rel}$(**S**)] | ≈ SD[$V_{rel}$(**S**)] | Median | Mean | ESD | $T$ | Pow. 5% | Pow. 1% |
|---|---|---|---|---|---|---|---|---|
| $p$ = 64, linearly decreasing λ ($V_{rel}$(**Σ**) = 0.0051) | | | | | | | | |
| $N$ = 4 | 0.3456 | 0.01471 | 0.3427 | 0.3456 | 0.01414 | 0.1623 | 0.1296 | 0.0366 |
| $N$ = 8 | 0.1522 | 0.00642 | 0.1514 | 0.1522 | 0.00633 | 0.0749 | 0.2962 | 0.1338 |
| $N$ = 16 | 0.0740 | 0.00300 | 0.0737 | 0.0739 | 0.00296 | −1.7247 | 0.6844 | 0.4484 |
| $N$ = 32 | 0.0385 | 0.00148 | 0.0384 | 0.0385 | 0.00148 | 0.6798 | 0.9936 | 0.9696 |
| $N$ = 64 | 0.0216 | 0.00076 | 0.0215 | 0.0216 | 0.00076 | 0.7268 | 1.0000 | 1.0000 |
| $N$ = 128 | 0.0133 | 0.00041 | 0.0133 | 0.0133 | 0.00042 | −0.5858 | 1.0000 | 1.0000 |
| $N$ = 256 | 0.0092 | 0.00024 | 0.0092 | 0.0092 | 0.00023 | −2.1676 | 1.0000 | 1.0000 |
| $p$ = 128, linearly decreasing λ ($V_{rel}$(**Σ**) = 0.0026) | | | | | | | | |
| $N$ = 4 | 0.3396 | 0.00734 | 0.3381 | 0.3395 | 0.00715 | −1.1276 | 0.1388 | 0.0446 |
| $N$ = 8 | 0.1476 | 0.00317 | 0.1473 | 0.1476 | 0.00319 | 1.3934 | 0.3240 | 0.1514 |
| $N$ = 16 | 0.0704 | 0.00146 | 0.0702 | 0.0703 | 0.00146 | −1.6467 | 0.7072 | 0.4642 |
| $N$ = 32 | 0.0354 | 0.00071 | 0.0354 | 0.0354 | 0.00071 | −0.1477 | 0.9956 | 0.9772 |
| $N$ = 64 | 0.0187 | 0.00036 | 0.0187 | 0.0187 | 0.00036 | −0.4842 | 1.0000 | 1.0000 |
| $N$ = 128 | 0.0106 | 0.00019 | 0.0106 | 0.0106 | 0.00019 | 0.7478 | 1.0000 | 1.0000 |
| $N$ = 256 | 0.0066 | 0.00010 | 0.0066 | 0.0066 | 0.00010 | 1.7065 | 1.0000 | 1.0000 |
| $p$ = 256, linearly decreasing λ ($V_{rel}$(**Σ**) = 0.0013) | | | | | | | | |
| $N$ = 4 | 0.3365 | 0.00366 | 0.3357 | 0.3365 | 0.00365 | 0.5477 | 0.1388 | 0.0488 |
| $N$ = 8 | 0.1452 | 0.00157 | 0.1451 | 0.1453 | 0.00155 | 0.9592 | 0.3200 | 0.1046 |
| $N$ = 16 | 0.0685 | 0.00072 | 0.0685 | 0.0685 | 0.00071 | 2.6723 | 0.7506 | 0.4980 |
| $N$ = 32 | 0.0338 | 0.00035 | 0.0338 | 0.0338 | 0.00035 | −0.3700 | 0.9968 | 0.9796 |
| $N$ = 64 | 0.0173 | 0.00017 | 0.0173 | 0.0173 | 0.00017 | 0.7181 | 1.0000 | 1.0000 |
| $N$ = 128 | 0.0092 | 0.00009 | 0.0092 | 0.0092 | 0.00009 | −0.0474 | 1.0000 | 1.0000 |
| $N$ = 256 | 0.0053 | 0.00005 | 0.0053 | 0.0053 | 0.00005 | −0.2356 | 1.0000 | 1.0000 |
| $p$ = 1024, linearly decreasing λ ($V_{rel}$(**Σ**) = 0.0003) | | | | | | | | |
| $N$ = 4 | 0.3341 | 0.00092 | 0.3340 | 0.3341 | 0.00093 | 0.5238 | 0.1328 | 0.0428 |
| $N$ = 8 | 0.1435 | 0.00039 | 0.1434 | 0.1435 | 0.00039 | −0.0304 | 0.3188 | 0.1342 |
| $N$ = 16 | 0.0671 | 0.00018 | 0.0671 | 0.0671 | 0.00018 | 0.0071 | 0.7104 | 0.5334 |
| $N$ = 32 | 0.0327 | 0.00009 | 0.0327 | 0.0327 | 0.00009 | 1.9692 | 0.9984 | 0.9884 |
| $N$ = 64 | 0.0162 | 0.00004 | 0.0162 | 0.0162 | 0.00004 | −1.4251 | 1.0000 | 1.0000 |
| $N$ = 128 | 0.0082 | 0.00002 | 0.0082 | 0.0082 | 0.00002 | −0.7182 | 1.0000 | 1.0000 |
| $N$ = 256 | 0.0043 | 0.00001 | 0.0043 | 0.0043 | 0.00001 | 0.4714 | 1.0000 | 1.0000 |

(*continued*)

**Table S2.** (*continued*)

| | $\approx E[V_{rel}(\mathbf{S})]$ | $\approx SD[V_{rel}(\mathbf{S})]$ | Median | Mean | ESD | $T$ | Pow. 5% | Pow. 1% |
|---|---|---|---|---|---|---|---|---|
| $p = 2$, quadratically decreasing $\lambda$ ($V_{rel}(\mathbf{\Sigma}) = 0.3600$) | | | | | | | | |
| $N = 4$ | 0.4588 | 0.3879 | 0.6400 | 0.6006 | 0.2781 | 36.0479 | 0.0902 | 0.0184 |
| $N = 8$ | 0.4107 | 0.2919 | 0.4696 | 0.4570 | 0.2354 | 13.8894 | 0.2646 | 0.0982 |
| $N = 16$ | 0.3899 | 0.2029 | 0.4064 | 0.4019 | 0.1785 | 4.7316 | 0.6200 | 0.3362 |
| $N = 32$ | 0.3766 | 0.1403 | 0.3794 | 0.3790 | 0.1297 | 1.3433 | 0.9294 | 0.7882 |
| $N = 64$ | 0.3687 | 0.0977 | 0.3710 | 0.3700 | 0.0946 | 0.9496 | 0.9992 | 0.9960 |
| $N = 128$ | 0.3645 | 0.0685 | 0.3644 | 0.3653 | 0.0671 | 0.8978 | 1.0000 | 1.0000 |
| $N = 256$ | 0.3623 | 0.0482 | 0.3629 | 0.3620 | 0.0480 | −0.4354 | 1.0000 | 1.0000 |
| $p = 4$, quadratically decreasing $\lambda$ ($V_{rel}(\mathbf{\Sigma}) = 0.1911$) | | | | | | | | |
| $N = 4$ | 0.4786 | 0.2770 | 0.5126 | 0.5300 | 0.1744 | 20.8377 | 0.1746 | 0.0572 |
| $N = 8$ | 0.3332 | 0.1705 | 0.3275 | 0.3461 | 0.1250 | 7.3018 | 0.3820 | 0.1802 |
| $N = 16$ | 0.2628 | 0.1044 | 0.2523 | 0.2658 | 0.0869 | 2.4188 | 0.8598 | 0.6038 |
| $N = 32$ | 0.2272 | 0.0667 | 0.2212 | 0.2286 | 0.0606 | 1.6646 | 0.9998 | 0.9914 |
| $N = 64$ | 0.2092 | 0.0444 | 0.2043 | 0.2090 | 0.0418 | −0.3674 | 1.0000 | 1.0000 |
| $N = 128$ | 0.2002 | 0.0303 | 0.1977 | 0.2004 | 0.0293 | 0.5275 | 1.0000 | 1.0000 |
| $N = 256$ | 0.1957 | 0.0211 | 0.1948 | 0.1956 | 0.0205 | −0.2530 | 1.0000 | 1.0000 |
| $p = 8$, quadratically decreasing $\lambda$ ($V_{rel}(\mathbf{\Sigma}) = 0.0980$) | | | | | | | | |
| $N = 4$ | 0.4455 | 0.1557 | 0.4359 | 0.4568 | 0.1104 | 7.2433 | 0.2120 | 0.0808 |
| $N = 8$ | 0.2589 | 0.0830 | 0.2499 | 0.2608 | 0.0683 | 1.8730 | 0.5416 | 0.3214 |
| $N = 16$ | 0.1759 | 0.0461 | 0.1704 | 0.1762 | 0.0410 | 0.6287 | 0.9666 | 0.8714 |
| $N = 32$ | 0.1363 | 0.0274 | 0.1333 | 0.1365 | 0.0257 | 0.3997 | 1.0000 | 1.0000 |
| $N = 64$ | 0.1170 | 0.0174 | 0.1152 | 0.1169 | 0.0170 | −0.7032 | 1.0000 | 1.0000 |
| $N = 128$ | 0.1075 | 0.0116 | 0.1067 | 0.1075 | 0.0114 | 0.0192 | 1.0000 | 1.0000 |
| $N = 256$ | 0.1028 | 0.0079 | 0.1023 | 0.1028 | 0.0078 | 0.2060 | 1.0000 | 1.0000 |
| $p = 16$, quadratically decreasing $\lambda$ ($V_{rel}(\mathbf{\Sigma}) = 0.0496$) | | | | | | | | |
| $N = 4$ | 0.4022 | 0.0801 | 0.3929 | 0.4044 | 0.0655 | 2.4555 | 0.2552 | 0.1008 |
| $N = 8$ | 0.2069 | 0.0388 | 0.2025 | 0.2075 | 0.0346 | 1.1343 | 0.6284 | 0.3734 |
| $N = 16$ | 0.1243 | 0.0201 | 0.1219 | 0.1242 | 0.0189 | −0.2887 | 0.9884 | 0.9362 |
| $N = 32$ | 0.0860 | 0.0112 | 0.0851 | 0.0861 | 0.0108 | 0.3780 | 1.0000 | 1.0000 |
| $N = 64$ | 0.0676 | 0.0068 | 0.0670 | 0.0676 | 0.0066 | 0.5759 | 1.0000 | 1.0000 |
| $N = 128$ | 0.0585 | 0.0043 | 0.0582 | 0.0584 | 0.0042 | −1.0260 | 1.0000 | 1.0000 |
| $N = 256$ | 0.0540 | 0.0029 | 0.0540 | 0.0541 | 0.0029 | 2.0991 | 1.0000 | 1.0000 |
| $p = 32$, quadratically decreasing $\lambda$ ($V_{rel}(\mathbf{\Sigma}) = 0.0249$) | | | | | | | | |
| $N = 4$ | 0.3713 | 0.0401 | 0.3642 | 0.3715 | 0.0363 | 0.3888 | 0.2768 | 0.1296 |
| $N = 8$ | 0.1765 | 0.0183 | 0.1741 | 0.1765 | 0.0177 | 0.2092 | 0.6570 | 0.4126 |
| $N = 16$ | 0.0962 | 0.0090 | 0.0953 | 0.0960 | 0.0087 | −1.6594 | 0.9944 | 0.9622 |
| $N = 32$ | 0.0595 | 0.0048 | 0.0593 | 0.0596 | 0.0047 | 0.8578 | 1.0000 | 1.0000 |
| $N = 64$ | 0.0420 | 0.0027 | 0.0418 | 0.0420 | 0.0027 | −0.3644 | 1.0000 | 1.0000 |
| $N = 128$ | 0.0334 | 0.0017 | 0.0333 | 0.0334 | 0.0017 | 0.6213 | 1.0000 | 1.0000 |
| $N = 256$ | 0.0291 | 0.0011 | 0.0291 | 0.0291 | 0.0011 | 0.8151 | 1.0000 | 1.0000 |

(*continued*)

**Table S2.** (*continued*)

| | ≈ E[$V_{rel}$(**S**)] | ≈ SD[$V_{rel}$(**S**)] | Median | Mean | ESD | $T$ | Pow. 5% | Pow. 1% |
|---|---|---|---|---|---|---|---|---|
| $p$ = 64, quadratically decreasing λ ($V_{rel}$(**Σ**) = 0.0125) | | | | | | | | |
| $N$ = 4 | 0.3532 | 0.01994 | 0.3501 | 0.3537 | 0.01885 | 1.7900 | 0.2828 | 0.1174 |
| $N$ = 8 | 0.1601 | 0.00883 | 0.1590 | 0.1600 | 0.00873 | −0.6981 | 0.6952 | 0.4790 |
| $N$ = 16 | 0.0816 | 0.00420 | 0.0814 | 0.0818 | 0.00424 | 2.1275 | 0.9958 | 0.9770 |
| $N$ = 32 | 0.0460 | 0.00213 | 0.0459 | 0.0460 | 0.00209 | 0.5724 | 1.0000 | 1.0000 |
| $N$ = 64 | 0.0290 | 0.00115 | 0.0290 | 0.0290 | 0.00114 | 2.1029 | 1.0000 | 1.0000 |
| $N$ = 128 | 0.0207 | 0.00066 | 0.0206 | 0.0207 | 0.00066 | −1.9228 | 1.0000 | 1.0000 |
| $N$ = 256 | 0.0166 | 0.00041 | 0.0166 | 0.0166 | 0.00041 | 0.8326 | 1.0000 | 1.0000 |
| $p$ = 128, quadratically decreasing λ ($V_{rel}$(**Σ**) = 0.0062) | | | | | | | | |
| $N$ = 4 | 0.3435 | 0.00993 | 0.3416 | 0.3436 | 0.00964 | 0.2942 | 0.2888 | 0.1320 |
| $N$ = 8 | 0.1516 | 0.00432 | 0.1511 | 0.1515 | 0.00424 | −0.6585 | 0.7264 | 0.5188 |
| $N$ = 16 | 0.0742 | 0.00202 | 0.0741 | 0.0742 | 0.00200 | −0.8615 | 0.9962 | 0.9824 |
| $N$ = 32 | 0.0392 | 0.00100 | 0.0391 | 0.0391 | 0.00100 | −1.0978 | 1.0000 | 1.0000 |
| $N$ = 64 | 0.0224 | 0.00052 | 0.0224 | 0.0224 | 0.00051 | −1.0415 | 1.0000 | 1.0000 |
| $N$ = 128 | 0.0143 | 0.00028 | 0.0143 | 0.0143 | 0.00028 | −0.0377 | 1.0000 | 1.0000 |
| $N$ = 256 | 0.0102 | 0.00016 | 0.0102 | 0.0102 | 0.00016 | −0.4172 | 1.0000 | 1.0000 |
| $p$ = 256, quadratically decreasing λ ($V_{rel}$(**Σ**) = 0.0031) | | | | | | | | |
| $N$ = 4 | 0.3385 | 0.00495 | 0.3375 | 0.3385 | 0.00483 | −0.3735 | 0.2886 | 0.1312 |
| $N$ = 8 | 0.1472 | 0.00214 | 0.1470 | 0.1472 | 0.00212 | 0.1107 | 0.7280 | 0.4636 |
| $N$ = 16 | 0.0704 | 0.00099 | 0.0704 | 0.0704 | 0.00098 | −1.6367 | 0.9976 | 0.9846 |
| $N$ = 32 | 0.0357 | 0.00048 | 0.0357 | 0.0357 | 0.00048 | 0.6626 | 1.0000 | 1.0000 |
| $N$ = 64 | 0.0192 | 0.00024 | 0.0192 | 0.0192 | 0.00024 | −1.5591 | 1.0000 | 1.0000 |
| $N$ = 128 | 0.0111 | 0.00013 | 0.0111 | 0.0111 | 0.00013 | −2.4955 | 1.0000 | 1.0000 |
| $N$ = 256 | 0.0071 | 0.00007 | 0.0071 | 0.0071 | 0.00007 | 0.0791 | 1.0000 | 1.0000 |
| $p$ = 1024, quadratically decreasing λ ($V_{rel}$(**Σ**) = 0.0008) | | | | | | | | |
| $N$ = 4 | 0.3346 | 0.00124 | 0.3344 | 0.3346 | 0.00121 | 0.6326 | 0.2838 | 0.1306 |
| $N$ = 8 | 0.1440 | 0.00053 | 0.1439 | 0.1440 | 0.00054 | 1.2686 | 0.7300 | 0.5134 |
| $N$ = 16 | 0.0676 | 0.00024 | 0.0676 | 0.0676 | 0.00024 | −1.5337 | 0.9968 | 0.9880 |
| $N$ = 32 | 0.0331 | 0.00012 | 0.0331 | 0.0331 | 0.00011 | 0.5647 | 1.0000 | 1.0000 |
| $N$ = 64 | 0.0167 | 0.00006 | 0.0167 | 0.0167 | 0.00006 | −0.7195 | 1.0000 | 1.0000 |
| $N$ = 128 | 0.0087 | 0.00003 | 0.0087 | 0.0087 | 0.00003 | −1.0076 | 1.0000 | 1.0000 |
| $N$ = 256 | 0.0047 | 0.00001 | 0.0047 | 0.0047 | 0.00002 | −0.1773 | 1.0000 | 1.0000 |

**Table S3.** Summary of simulation results for $V_{rel}(\mathbf{R})$. Theoretical expectation ($E[V_{rel}(\mathbf{R})]$) and standard deviation ($SD[V_{rel}(\mathbf{R})]$; asymptotic results are shown for the latter in non-null conditions with $p > 2$), and empirical median, mean, standard deviation (ESD), bias in standard error unit ($T$), and critical points or power (for null and non-null conditions, respectively) at $\alpha = 0.05$ and 0.01 (CP/Pow. 5% and 1%) from 5000 simulation runs are shown. See Table S1 for further information.

| | $E[V_{rel}(\mathbf{R})]$ | $SD[V_{rel}(\mathbf{R})]$ | Median | Mean | ESD | $T$ | CP 5% | CP 1% |
|---|---|---|---|---|---|---|---|---|
| $p = 2$, $V_{rel}(\mathbf{P}) = 0$ | | | | | | | | |
| $N = 4$ | 0.3333 | 0.2981 | 0.2517 | 0.3325 | 0.2989 | −0.1990 | 0.9021 | 0.9807 |
| $N = 8$ | 0.1429 | 0.1650 | 0.0792 | 0.1429 | 0.1635 | 0.0363 | 0.4979 | 0.6857 |
| $N = 16$ | 0.0667 | 0.0856 | 0.0336 | 0.0679 | 0.0862 | 1.0235 | 0.2456 | 0.3876 |
| $N = 32$ | 0.0323 | 0.0435 | 0.0163 | 0.0326 | 0.0427 | 0.6113 | 0.1194 | 0.1934 |
| $N = 64$ | 0.0159 | 0.0219 | 0.0075 | 0.0156 | 0.0205 | −1.0380 | 0.0583 | 0.0938 |
| $N = 128$ | 0.0079 | 0.0110 | 0.0035 | 0.0076 | 0.0104 | −1.5798 | 0.0295 | 0.0467 |
| $N = 256$ | 0.0039 | 0.0055 | 0.0018 | 0.0039 | 0.0054 | −0.3981 | 0.0148 | 0.0252 |
| $p = 4$, $V_{rel}(\mathbf{P}) = 0$ | | | | | | | | |
| $N = 4$ | 0.3333 | 0.1217 | 0.3152 | 0.3327 | 0.1214 | −0.3906 | 0.5646 | 0.7079 |
| $N = 8$ | 0.1429 | 0.0673 | 0.1332 | 0.1432 | 0.0673 | 0.3148 | 0.2728 | 0.3435 |
| $N = 16$ | 0.0667 | 0.0349 | 0.0608 | 0.0661 | 0.0344 | −1.0696 | 0.1289 | 0.1728 |
| $N = 32$ | 0.0323 | 0.0178 | 0.0292 | 0.0321 | 0.0175 | −0.6906 | 0.0642 | 0.0866 |
| $N = 64$ | 0.0159 | 0.0090 | 0.0142 | 0.0157 | 0.0087 | −1.0749 | 0.0323 | 0.0426 |
| $N = 128$ | 0.0079 | 0.0045 | 0.0070 | 0.0079 | 0.0045 | 0.0033 | 0.0165 | 0.0215 |
| $N = 256$ | 0.0039 | 0.0023 | 0.0035 | 0.0039 | 0.0022 | −0.9161 | 0.0081 | 0.0107 |
| $p = 8$, $V_{rel}(\mathbf{P}) = 0$ | | | | | | | | |
| $N = 4$ | 0.3333 | 0.0563 | 0.3228 | 0.3336 | 0.0569 | 0.3780 | 0.4418 | 0.5149 |
| $N = 8$ | 0.1429 | 0.0312 | 0.1402 | 0.1428 | 0.0312 | −0.1003 | 0.1975 | 0.2302 |
| $N = 16$ | 0.0667 | 0.0162 | 0.0654 | 0.0668 | 0.0159 | 0.4573 | 0.0948 | 0.1102 |
| $N = 32$ | 0.0323 | 0.0082 | 0.0315 | 0.0322 | 0.0082 | −0.1190 | 0.0465 | 0.0544 |
| $N = 64$ | 0.0159 | 0.0041 | 0.0156 | 0.0159 | 0.0041 | 0.2305 | 0.0232 | 0.0270 |
| $N = 128$ | 0.0079 | 0.0021 | 0.0077 | 0.0079 | 0.0021 | 0.0628 | 0.0117 | 0.0136 |
| $N = 256$ | 0.0039 | 0.0010 | 0.0039 | 0.0039 | 0.0010 | 1.2483 | 0.0058 | 0.0067 |
| $p = 16$, $V_{rel}(\mathbf{P}) = 0$ | | | | | | | | |
| $N = 4$ | 0.3333 | 0.0272 | 0.3281 | 0.3332 | 0.0265 | −0.4751 | 0.3850 | 0.4135 |
| $N = 8$ | 0.1429 | 0.0151 | 0.1411 | 0.1428 | 0.0151 | −0.0917 | 0.1698 | 0.1840 |
| $N = 16$ | 0.0667 | 0.0078 | 0.0663 | 0.0669 | 0.0080 | 1.7743 | 0.0810 | 0.0879 |
| $N = 32$ | 0.0323 | 0.0040 | 0.0319 | 0.0322 | 0.0039 | −1.7517 | 0.0389 | 0.0428 |
| $N = 64$ | 0.0159 | 0.0020 | 0.0158 | 0.0159 | 0.0020 | 0.7127 | 0.0192 | 0.0209 |
| $N = 128$ | 0.0079 | 0.0010 | 0.0078 | 0.0079 | 0.0010 | −0.7405 | 0.0096 | 0.0104 |
| $N = 256$ | 0.0039 | 0.0005 | 0.0039 | 0.0039 | 0.0005 | 1.3884 | 0.0048 | 0.0052 |
| $p = 32$, $V_{rel}(\mathbf{P}) = 0$ | | | | | | | | |
| $N = 4$ | 0.3333 | 0.0134 | 0.3306 | 0.3332 | 0.0133 | −0.7714 | 0.3586 | 0.3770 |
| $N = 8$ | 0.1429 | 0.0074 | 0.1418 | 0.1426 | 0.0074 | −2.0764 | 0.1562 | 0.1627 |
| $N = 16$ | 0.0667 | 0.0038 | 0.0665 | 0.0666 | 0.0038 | −1.7329 | 0.0732 | 0.0766 |
| $N = 32$ | 0.0323 | 0.0020 | 0.0322 | 0.0322 | 0.0019 | −1.3105 | 0.0355 | 0.0370 |
| $N = 64$ | 0.0159 | 0.0010 | 0.0158 | 0.0159 | 0.0010 | −0.5099 | 0.0175 | 0.0183 |
| $N = 128$ | 0.0079 | 0.0005 | 0.0079 | 0.0079 | 0.0005 | −0.0426 | 0.0087 | 0.0091 |
| $N = 256$ | 0.0039 | 0.0002 | 0.0039 | 0.0039 | 0.0002 | −0.0743 | 0.0043 | 0.0045 |

(*continued*)

**Table S3.** (*continued*)

| | $E[V_{rel}(\mathbf{R})]$ | $SD[V_{rel}(\mathbf{R})]$ | Median | Mean | ESD | $T$ | CP 5% | CP 1% |
|---|---|---|---|---|---|---|---|---|
| $p = 64$, $V_{rel}(\mathbf{P}) = 0$ | | | | | | | | |
| $N = 4$ | 0.3333 | 0.00664 | 0.3320 | 0.3333 | 0.00677 | −0.0624 | 0.3463 | 0.3543 |
| $N = 8$ | 0.1429 | 0.00367 | 0.1425 | 0.1428 | 0.00363 | −0.4617 | 0.1492 | 0.1530 |
| $N = 16$ | 0.0667 | 0.00191 | 0.0666 | 0.0667 | 0.00188 | 0.4407 | 0.0699 | 0.0716 |
| $N = 32$ | 0.0323 | 0.00097 | 0.0322 | 0.0323 | 0.00099 | −0.0725 | 0.0339 | 0.0347 |
| $N = 64$ | 0.0159 | 0.00049 | 0.0159 | 0.0159 | 0.00048 | −0.6525 | 0.0167 | 0.0170 |
| $N = 128$ | 0.0079 | 0.00025 | 0.0079 | 0.0079 | 0.00025 | 0.8165 | 0.0083 | 0.0085 |
| $N = 256$ | 0.0039 | 0.00012 | 0.0039 | 0.0039 | 0.00012 | 1.7441 | 0.0041 | 0.0042 |
| $p = 128$, $V_{rel}(\mathbf{P}) = 0$ | | | | | | | | |
| $N = 4$ | 0.3333 | 0.00331 | 0.3327 | 0.3333 | 0.00327 | 0.1648 | 0.3397 | 0.3433 |
| $N = 8$ | 0.1429 | 0.00183 | 0.1427 | 0.1428 | 0.00180 | −1.3211 | 0.1460 | 0.1478 |
| $N = 16$ | 0.0667 | 0.00095 | 0.0666 | 0.0667 | 0.00094 | 0.7843 | 0.0683 | 0.0692 |
| $N = 32$ | 0.0323 | 0.00048 | 0.0322 | 0.0323 | 0.00049 | 0.3062 | 0.0331 | 0.0334 |
| $N = 64$ | 0.0159 | 0.00024 | 0.0159 | 0.0159 | 0.00024 | −0.7890 | 0.0163 | 0.0165 |
| $N = 128$ | 0.0079 | 0.00012 | 0.0079 | 0.0079 | 0.00012 | −0.8957 | 0.0081 | 0.0082 |
| $N = 256$ | 0.0039 | 0.00006 | 0.0039 | 0.0039 | 0.00006 | 0.3419 | 0.0040 | 0.0041 |
| $p = 256$, $V_{rel}(\mathbf{P}) = 0$ | | | | | | | | |
| $N = 4$ | 0.3333 | 0.00165 | 0.3330 | 0.3334 | 0.00170 | 1.8964 | 0.3366 | 0.3388 |
| $N = 8$ | 0.1429 | 0.00091 | 0.1428 | 0.1429 | 0.00091 | 0.0105 | 0.1445 | 0.1453 |
| $N = 16$ | 0.0667 | 0.00047 | 0.0667 | 0.0667 | 0.00047 | 1.8676 | 0.0675 | 0.0679 |
| $N = 32$ | 0.0323 | 0.00024 | 0.0323 | 0.0323 | 0.00024 | −0.9701 | 0.0327 | 0.0328 |
| $N = 64$ | 0.0159 | 0.00012 | 0.0159 | 0.0159 | 0.00012 | 0.3244 | 0.0161 | 0.0162 |
| $N = 128$ | 0.0079 | 0.00006 | 0.0079 | 0.0079 | 0.00006 | 0.8789 | 0.0080 | 0.0080 |
| $N = 256$ | 0.0039 | 0.00003 | 0.0039 | 0.0039 | 0.00003 | 0.5874 | 0.0040 | 0.0040 |
| $p = 1024$, $V_{rel}(\mathbf{P}) = 0$ | | | | | | | | |
| $N = 4$ | 0.3333 | 0.00041 | 0.3333 | 0.3333 | 0.00041 | 1.4469 | 0.3342 | 0.3346 |
| $N = 8$ | 0.1429 | 0.00023 | 0.1428 | 0.1429 | 0.00023 | −0.9639 | 0.1433 | 0.1435 |
| $N = 16$ | 0.0667 | 0.00012 | 0.0667 | 0.0667 | 0.00012 | 0.5220 | 0.0669 | 0.0670 |
| $N = 32$ | 0.0323 | 0.00006 | 0.0323 | 0.0323 | 0.00006 | −1.8393 | 0.0324 | 0.0324 |
| $N = 64$ | 0.0159 | 0.00003 | 0.0159 | 0.0159 | 0.00003 | −0.0357 | 0.0159 | 0.0159 |
| $N = 128$ | 0.0079 | 0.00002 | 0.0079 | 0.0079 | 0.00002 | 0.1288 | 0.0079 | 0.0079 |
| $N = 256$ | 0.0039 | 0.00001 | 0.0039 | 0.0039 | 0.00001 | −0.3305 | 0.0039 | 0.0039 |

(*continued*)

**Table S3.** (*continued*)

| | $E[V_{rel}(\mathbf{R})]$ | $SD[V_{rel}(\mathbf{R})]$ | Median | Mean | ESD | $T$ | Pow. 5% | Pow. 1% |
|---|---|---|---|---|---|---|---|---|
| $p = 2, q = 1, V_{rel}(\mathbf{P}) = 0.1$ | | | | | | | | |
| $N = 4$ | 0.3745 | 0.3092 | 0.3229 | 0.3769 | 0.3071 | 0.5423 | 0.0606 | 0.0126 |
| $N = 8$ | 0.2108 | 0.2061 | 0.1513 | 0.2161 | 0.2092 | 1.7989 | 0.1222 | 0.0372 |
| $N = 16$ | 0.1499 | 0.1420 | 0.1031 | 0.1437 | 0.1389 | −3.1691 | 0.2126 | 0.0712 |
| $N = 32$ | 0.1237 | 0.1001 | 0.1050 | 0.1259 | 0.1012 | 1.5152 | 0.4484 | 0.2328 |
| $N = 64$ | 0.1115 | 0.0709 | 0.1011 | 0.1117 | 0.0702 | 0.2070 | 0.7482 | 0.5448 |
| $N = 128$ | 0.1057 | 0.0502 | 0.0996 | 0.1047 | 0.0503 | −1.4136 | 0.9612 | 0.8832 |
| $N = 256$ | 0.1028 | 0.0355 | 0.1018 | 0.1038 | 0.0355 | 1.9926 | 0.9998 | 0.9968 |
| $p = 2, q = 1, V_{rel}(\mathbf{P}) = 0.2$ | | | | | | | | |
| $N = 4$ | 0.4184 | 0.3169 | 0.3856 | 0.4190 | 0.3181 | 0.1492 | 0.0826 | 0.0178 |
| $N = 8$ | 0.2814 | 0.2314 | 0.2368 | 0.2799 | 0.2312 | −0.4442 | 0.1958 | 0.0660 |
| $N = 16$ | 0.2350 | 0.1696 | 0.2114 | 0.2330 | 0.1682 | −0.8121 | 0.4312 | 0.1894 |
| $N = 32$ | 0.2162 | 0.1229 | 0.2045 | 0.2162 | 0.1232 | 0.0219 | 0.7550 | 0.5344 |
| $N = 64$ | 0.2078 | 0.0881 | 0.2054 | 0.2083 | 0.0890 | 0.3926 | 0.9686 | 0.9028 |
| $N = 128$ | 0.2038 | 0.0628 | 0.2018 | 0.2040 | 0.0623 | 0.1506 | 1.0000 | 0.9990 |
| $N = 256$ | 0.2019 | 0.0446 | 0.2014 | 0.2025 | 0.0442 | 1.0388 | 1.0000 | 1.0000 |
| $p = 2, q = 1, V_{rel}(\mathbf{P}) = 0.4$ | | | | | | | | |
| $N = 4$ | 0.5158 | 0.3203 | 0.5639 | 0.5254 | 0.3179 | 2.1361 | 0.1410 | 0.0274 |
| $N = 8$ | 0.4318 | 0.2495 | 0.4362 | 0.4294 | 0.2516 | −0.6571 | 0.4256 | 0.1854 |
| $N = 16$ | 0.4111 | 0.1844 | 0.4230 | 0.4147 | 0.1832 | 1.3640 | 0.8050 | 0.5666 |
| $N = 32$ | 0.4046 | 0.1326 | 0.4056 | 0.4041 | 0.1322 | −0.2806 | 0.9858 | 0.9388 |
| $N = 64$ | 0.4021 | 0.0944 | 0.4058 | 0.4025 | 0.0954 | 0.3294 | 1.0000 | 1.0000 |
| $N = 128$ | 0.4010 | 0.0669 | 0.4020 | 0.4013 | 0.0665 | 0.3770 | 1.0000 | 1.0000 |
| $N = 256$ | 0.4005 | 0.0474 | 0.4006 | 0.4000 | 0.0480 | −0.7435 | 1.0000 | 1.0000 |
| $p = 2, q = 1, V_{rel}(\mathbf{P}) = 0.6$ | | | | | | | | |
| $N = 4$ | 0.6313 | 0.3022 | 0.7219 | 0.6328 | 0.3010 | 0.3570 | 0.2224 | 0.0522 |
| $N = 8$ | 0.5973 | 0.2285 | 0.6376 | 0.5970 | 0.2281 | −0.0946 | 0.7126 | 0.4182 |
| $N = 16$ | 0.5963 | 0.1608 | 0.6175 | 0.5980 | 0.1623 | 0.7784 | 0.9678 | 0.8914 |
| $N = 32$ | 0.5978 | 0.1120 | 0.6072 | 0.5973 | 0.1108 | −0.3007 | 0.9998 | 0.9990 |
| $N = 64$ | 0.5988 | 0.0784 | 0.6017 | 0.5978 | 0.0787 | −0.8984 | 1.0000 | 1.0000 |
| $N = 128$ | 0.5994 | 0.0551 | 0.6010 | 0.5991 | 0.0556 | −0.3529 | 1.0000 | 1.0000 |
| $N = 256$ | 0.5997 | 0.0389 | 0.6008 | 0.5996 | 0.0385 | −0.2220 | 1.0000 | 1.0000 |
| $p = 2, q = 1, V_{rel}(\mathbf{P}) = 0.8$ | | | | | | | | |
| $N = 4$ | 0.7768 | 0.2440 | 0.8733 | 0.7748 | 0.2470 | −0.5570 | 0.4220 | 0.1180 |
| $N = 8$ | 0.7831 | 0.1596 | 0.8250 | 0.7853 | 0.1571 | 0.9790 | 0.9352 | 0.8012 |
| $N = 16$ | 0.7918 | 0.1015 | 0.8141 | 0.7928 | 0.1033 | 0.6253 | 0.9996 | 0.9956 |
| $N = 32$ | 0.7961 | 0.0674 | 0.8076 | 0.7976 | 0.0680 | 1.5536 | 1.0000 | 1.0000 |
| $N = 64$ | 0.7981 | 0.0462 | 0.8024 | 0.7976 | 0.0473 | −0.6617 | 1.0000 | 1.0000 |
| $N = 128$ | 0.7991 | 0.0321 | 0.8006 | 0.7986 | 0.0319 | −1.0007 | 1.0000 | 1.0000 |
| $N = 256$ | 0.7995 | 0.0225 | 0.8008 | 0.8001 | 0.0223 | 1.8536 | 1.0000 | 1.0000 |

(*continued*)

**Table S3.** (*continued*)

| | $E[V_{rel}(\mathbf{R})]$ | $ASD_K$ | $ASD_{PF}$ | Median | Mean | ESD | $T$ | Pow. 5% | Pow. 1% |
|---|---|---|---|---|---|---|---|---|---|
| $p = 4, q = 1, V_{rel}(\mathbf{P}) = 0.1$ | | | | | | | | | |
| $N = 4$ | 0.3745 | 0.1986 | 0.1977 | 0.3467 | 0.3755 | 0.1470 | 0.4579 | 0.1144 | 0.0368 |
| $N = 8$ | 0.2108 | 0.1300 | 0.1285 | 0.1917 | 0.2134 | 0.1115 | 1.6596 | 0.2484 | 0.1262 |
| $N = 16$ | 0.1499 | 0.0888 | 0.0877 | 0.1373 | 0.1511 | 0.0815 | 1.0182 | 0.5380 | 0.3360 |
| $N = 32$ | 0.1237 | 0.0618 | 0.0613 | 0.1155 | 0.1240 | 0.0594 | 0.3959 | 0.8520 | 0.7024 |
| $N = 64$ | 0.1115 | 0.0433 | 0.0431 | 0.1070 | 0.1116 | 0.0429 | 0.0896 | 0.9884 | 0.9720 |
| $N = 128$ | 0.1057 | 0.0305 | 0.0304 | 0.1045 | 0.1065 | 0.0303 | 1.9867 | 1.0000 | 0.9998 |
| $N = 256$ | 0.1028 | 0.0215 | 0.0215 | 0.1016 | 0.1025 | 0.0213 | −1.0710 | 1.0000 | 1.0000 |
| $p = 4, q = 1, V_{rel}(\mathbf{P}) = 0.2$ | | | | | | | | | |
| $N = 4$ | 0.4184 | 0.2729 | 0.2441 | 0.3831 | 0.4167 | 0.1663 | −0.7193 | 0.1916 | 0.0690 |
| $N = 8$ | 0.2814 | 0.1786 | 0.1672 | 0.2562 | 0.2822 | 0.1433 | 0.4227 | 0.4578 | 0.2962 |
| $N = 16$ | 0.2350 | 0.1220 | 0.1176 | 0.2242 | 0.2352 | 0.1090 | 0.1625 | 0.8206 | 0.6788 |
| $N = 32$ | 0.2162 | 0.0849 | 0.0833 | 0.2107 | 0.2156 | 0.0792 | −0.5585 | 0.9862 | 0.9592 |
| $N = 64$ | 0.2078 | 0.0595 | 0.0590 | 0.2040 | 0.2076 | 0.0584 | −0.1842 | 1.0000 | 1.0000 |
| $N = 128$ | 0.2038 | 0.0419 | 0.0417 | 0.2011 | 0.2028 | 0.0411 | −1.6816 | 1.0000 | 1.0000 |
| $N = 256$ | 0.2019 | 0.0296 | 0.0295 | 0.2016 | 0.2023 | 0.0297 | 0.8644 | 1.0000 | 1.0000 |
| $p = 4, q = 1, V_{rel}(\mathbf{P}) = 0.4$ | | | | | | | | | |
| $N = 4$ | 0.5158 | 0.3175 | 0.2755 | 0.4927 | 0.5144 | 0.2023 | −0.4921 | 0.3980 | 0.2132 |
| $N = 8$ | 0.4318 | 0.2078 | 0.1950 | 0.4314 | 0.4287 | 0.1750 | −1.2271 | 0.7814 | 0.6600 |
| $N = 16$ | 0.4111 | 0.1420 | 0.1380 | 0.4141 | 0.4093 | 0.1317 | −0.9769 | 0.9866 | 0.9596 |
| $N = 32$ | 0.4046 | 0.0988 | 0.0975 | 0.4054 | 0.4034 | 0.0960 | −0.8743 | 1.0000 | 1.0000 |
| $N = 64$ | 0.4021 | 0.0693 | 0.0688 | 0.4033 | 0.4022 | 0.0687 | 0.1270 | 1.0000 | 1.0000 |
| $N = 128$ | 0.4010 | 0.0488 | 0.0486 | 0.4016 | 0.4003 | 0.0484 | −0.9561 | 1.0000 | 1.0000 |
| $N = 256$ | 0.4005 | 0.0344 | 0.0344 | 0.3998 | 0.3997 | 0.0336 | −1.7358 | 1.0000 | 1.0000 |
| $p = 4, q = 1, V_{rel}(\mathbf{P}) = 0.6$ | | | | | | | | | |
| $N = 4$ | 0.6313 | 0.2736 | 0.2472 | 0.6644 | 0.6329 | 0.2101 | 0.5400 | 0.6206 | 0.4366 |
| $N = 8$ | 0.5973 | 0.1791 | 0.1743 | 0.6219 | 0.5978 | 0.1714 | 0.2242 | 0.9456 | 0.9046 |
| $N = 16$ | 0.5963 | 0.1223 | 0.1215 | 0.6106 | 0.5949 | 0.1216 | −0.7641 | 0.9994 | 0.9986 |
| $N = 32$ | 0.5978 | 0.0851 | 0.0849 | 0.6047 | 0.5975 | 0.0846 | −0.1828 | 1.0000 | 1.0000 |
| $N = 64$ | 0.5988 | 0.0597 | 0.0597 | 0.6019 | 0.5985 | 0.0606 | −0.3532 | 1.0000 | 1.0000 |
| $N = 128$ | 0.5994 | 0.0420 | 0.0420 | 0.6005 | 0.5980 | 0.0422 | −2.2753 | 1.0000 | 1.0000 |
| $N = 256$ | 0.5997 | 0.0297 | 0.0297 | 0.6006 | 0.5996 | 0.0304 | −0.3144 | 1.0000 | 1.0000 |
| $p = 4, q = 1, V_{rel}(\mathbf{P}) = 0.8$ | | | | | | | | | |
| $N = 4$ | 0.7768 | 0.1640 | 0.1666 | 0.8388 | 0.7754 | 0.1873 | −0.5322 | 0.8496 | 0.7278 |
| $N = 8$ | 0.7831 | 0.1073 | 0.1115 | 0.8149 | 0.7840 | 0.1249 | 0.4605 | 0.9970 | 0.9896 |
| $N = 16$ | 0.7918 | 0.0733 | 0.0749 | 0.8059 | 0.7913 | 0.0815 | −0.4682 | 1.0000 | 0.9998 |
| $N = 32$ | 0.7961 | 0.0510 | 0.0516 | 0.8027 | 0.7953 | 0.0543 | −1.0173 | 1.0000 | 1.0000 |
| $N = 64$ | 0.7981 | 0.0358 | 0.0360 | 0.8019 | 0.7987 | 0.0370 | 1.2077 | 1.0000 | 1.0000 |
| $N = 128$ | 0.7991 | 0.0252 | 0.0253 | 0.8008 | 0.7991 | 0.0255 | 0.1156 | 1.0000 | 1.0000 |
| $N = 256$ | 0.7995 | 0.0178 | 0.0178 | 0.8003 | 0.7995 | 0.0176 | 0.0407 | 1.0000 | 1.0000 |

$ASD_K$: Asymptotic standard deviation of $V_{rel}(\mathbf{R})$ based on Konishi's (1979) theory (eq. 39).
$ASD_{PF}$: Approximate standard deviation of $V_{rel}(\mathbf{R})$ based on Pan & Frank's (2004) approach (eqs. 36–38).

(*continued*)

**Table S3.** (*continued*)

| | E[$V_{rel}$(**R**)] | $ASD_K$ | $ASD_{PF}$ | Median | Mean | ESD | $T$ | Pow. 5% | Pow. 1% |
|---|---|---|---|---|---|---|---|---|---|
| $p = 4$, $q = 2$, $V_{rel}$(**P**) = 0.1 | | | | | | | | | |
| $N = 4$ | 0.3773 | 0.1043 | 0.1806 | 0.3512 | 0.3774 | 0.1431 | 0.0581 | 0.1082 | 0.0342 |
| $N = 8$ | 0.2135 | 0.0683 | 0.0975 | 0.2034 | 0.2142 | 0.0884 | 0.5188 | 0.2168 | 0.0820 |
| $N = 16$ | 0.1518 | 0.0467 | 0.0578 | 0.1465 | 0.1514 | 0.0570 | −0.4810 | 0.6370 | 0.3098 |
| $N = 32$ | 0.1248 | 0.0325 | 0.0366 | 0.1233 | 0.1255 | 0.0361 | 1.3664 | 0.9658 | 0.8672 |
| $N = 64$ | 0.1121 | 0.0228 | 0.0243 | 0.1115 | 0.1125 | 0.0245 | 1.0436 | 1.0000 | 0.9992 |
| $N = 128$ | 0.1060 | 0.0160 | 0.0166 | 0.1057 | 0.1061 | 0.0166 | 0.3845 | 1.0000 | 1.0000 |
| $N = 256$ | 0.1030 | 0.0113 | 0.0115 | 0.1025 | 0.1027 | 0.0116 | −1.9263 | 1.0000 | 1.0000 |
| $p = 4$, $q = 2$, $V_{rel}$(**P**) = 0.2 | | | | | | | | | |
| $N = 4$ | 0.4327 | 0.0843 | 0.2312 | 0.3983 | 0.4377 | 0.1622 | 2.1772 | 0.2110 | 0.0806 |
| $N = 8$ | 0.2943 | 0.0552 | 0.1140 | 0.2784 | 0.2948 | 0.1011 | 0.3354 | 0.5266 | 0.2416 |
| $N = 16$ | 0.2432 | 0.0377 | 0.0616 | 0.2373 | 0.2428 | 0.0578 | −0.4744 | 0.9880 | 0.9142 |
| $N = 32$ | 0.2208 | 0.0262 | 0.0356 | 0.2202 | 0.2212 | 0.0357 | 0.8814 | 1.0000 | 1.0000 |
| $N = 64$ | 0.2102 | 0.0184 | 0.0220 | 0.2097 | 0.2100 | 0.0220 | −0.4393 | 1.0000 | 1.0000 |
| $N = 128$ | 0.2050 | 0.0130 | 0.0143 | 0.2052 | 0.2049 | 0.0146 | −0.5257 | 1.0000 | 1.0000 |
| $N = 256$ | 0.2025 | 0.0091 | 0.0096 | 0.2025 | 0.2025 | 0.0096 | 0.2252 | 1.0000 | 1.0000 |
| $p = 4$, $q = 2$, $V_{rel}$(**P**) = 0.4 | | | | | | | | | |
| (This comformation is impossible) | | | | | | | | | |

$ASD_K$: Asymptotic standard deviation of $V_{rel}$(**R**) based on Konishi's (1979) theory (eq. 39).
$ASD_{PF}$: Approximate standard deviation of $V_{rel}$(**R**) based on Pan & Frank's (2004) approach (eqs. 36–38).

(*continued*)

**Table S3.** (*continued*)

| | $E[V_{rel}(\mathbf{R})]$ | $ASD_K$ | $ASD_{PF}$ | Median | Mean | ESD | $T$ | Pow. 5% | Pow. 1% |
|---|---|---|---|---|---|---|---|---|---|
| $p = 8, q = 1, V_{rel}(\mathbf{P}) = 0.1$ | | | | | | | | | |
| $N = 4$ | 0.3745 | 0.1516 | 0.1518 | 0.3546 | 0.3735 | 0.0876 | −0.7927 | 0.1848 | 0.0746 |
| $N = 8$ | 0.2108 | 0.0993 | 0.0987 | 0.1930 | 0.2101 | 0.0762 | −0.6583 | 0.4750 | 0.3218 |
| $N = 16$ | 0.1499 | 0.0678 | 0.0674 | 0.1394 | 0.1496 | 0.0590 | −0.3952 | 0.8288 | 0.7152 |
| $N = 32$ | 0.1237 | 0.0472 | 0.0470 | 0.1193 | 0.1238 | 0.0440 | 0.2041 | 0.9884 | 0.9706 |
| $N = 64$ | 0.1115 | 0.0331 | 0.0330 | 0.1091 | 0.1119 | 0.0323 | 0.7713 | 0.9998 | 0.9996 |
| $N = 128$ | 0.1057 | 0.0233 | 0.0233 | 0.1045 | 0.1053 | 0.0228 | −1.1017 | 1.0000 | 1.0000 |
| $N = 256$ | 0.1028 | 0.0164 | 0.0164 | 0.1020 | 0.1030 | 0.0165 | 0.6914 | 1.0000 | 1.0000 |
| $p = 8, q = 1, V_{rel}(\mathbf{P}) = 0.2$ | | | | | | | | | |
| $N = 4$ | 0.4184 | 0.2228 | 0.2075 | 0.3921 | 0.4189 | 0.1187 | 0.3123 | 0.3430 | 0.1946 |
| $N = 8$ | 0.2814 | 0.1459 | 0.1404 | 0.2643 | 0.2826 | 0.1088 | 0.7886 | 0.7554 | 0.6256 |
| $N = 16$ | 0.2350 | 0.0997 | 0.0977 | 0.2253 | 0.2336 | 0.0864 | −1.1163 | 0.9706 | 0.9458 |
| $N = 32$ | 0.2162 | 0.0693 | 0.0686 | 0.2124 | 0.2164 | 0.0658 | 0.1980 | 1.0000 | 0.9998 |
| $N = 64$ | 0.2078 | 0.0486 | 0.0484 | 0.2052 | 0.2072 | 0.0480 | −0.9238 | 1.0000 | 1.0000 |
| $N = 128$ | 0.2038 | 0.0342 | 0.0342 | 0.2040 | 0.2046 | 0.0333 | 1.6580 | 1.0000 | 1.0000 |
| $N = 256$ | 0.2019 | 0.0242 | 0.0241 | 0.2015 | 0.2017 | 0.0241 | −0.6367 | 1.0000 | 1.0000 |
| $p = 8, q = 1, V_{rel}(\mathbf{P}) = 0.4$ | | | | | | | | | |
| $N = 4$ | 0.5158 | 0.2753 | 0.2491 | 0.4939 | 0.5144 | 0.1590 | −0.6168 | 0.6076 | 0.4548 |
| $N = 8$ | 0.4318 | 0.1802 | 0.1728 | 0.4270 | 0.4286 | 0.1493 | −1.5260 | 0.9392 | 0.8950 |
| $N = 16$ | 0.4111 | 0.1231 | 0.1208 | 0.4143 | 0.4127 | 0.1120 | 1.0025 | 0.9990 | 0.9986 |
| $N = 32$ | 0.4046 | 0.0856 | 0.0849 | 0.4066 | 0.4046 | 0.0835 | −0.0028 | 1.0000 | 1.0000 |
| $N = 64$ | 0.4021 | 0.0601 | 0.0598 | 0.4032 | 0.4023 | 0.0590 | 0.2713 | 1.0000 | 1.0000 |
| $N = 128$ | 0.4010 | 0.0423 | 0.0422 | 0.4019 | 0.4014 | 0.0419 | 0.7597 | 1.0000 | 1.0000 |
| $N = 256$ | 0.4005 | 0.0299 | 0.0298 | 0.4003 | 0.4001 | 0.0293 | −0.8777 | 1.0000 | 1.0000 |
| $p = 8, q = 1, V_{rel}(\mathbf{P}) = 0.6$ | | | | | | | | | |
| $N = 4$ | 0.6313 | 0.2447 | 0.2249 | 0.6492 | 0.6303 | 0.1808 | −0.3763 | 0.8000 | 0.7026 |
| $N = 8$ | 0.5973 | 0.1602 | 0.1557 | 0.6178 | 0.5958 | 0.1530 | −0.6788 | 0.9932 | 0.9826 |
| $N = 16$ | 0.5963 | 0.1094 | 0.1082 | 0.6130 | 0.5984 | 0.1090 | 1.4179 | 0.9998 | 0.9998 |
| $N = 32$ | 0.5978 | 0.0761 | 0.0758 | 0.6017 | 0.5954 | 0.0778 | −2.0986 | 1.0000 | 1.0000 |
| $N = 64$ | 0.5988 | 0.0534 | 0.0533 | 0.6008 | 0.5984 | 0.0540 | −0.5764 | 1.0000 | 1.0000 |
| $N = 128$ | 0.5994 | 0.0376 | 0.0376 | 0.6012 | 0.5992 | 0.0371 | −0.3409 | 1.0000 | 1.0000 |
| $N = 256$ | 0.5997 | 0.0265 | 0.0265 | 0.6008 | 0.5995 | 0.0266 | −0.4008 | 1.0000 | 1.0000 |
| $p = 8, q = 1, V_{rel}(\mathbf{P}) = 0.8$ | | | | | | | | | |
| $N = 4$ | 0.7768 | 0.1496 | 0.1446 | 0.8337 | 0.7780 | 0.1658 | 0.5054 | 0.9350 | 0.8998 |
| $N = 8$ | 0.7831 | 0.0979 | 0.0977 | 0.8104 | 0.7828 | 0.1154 | −0.2029 | 0.9996 | 0.9992 |
| $N = 16$ | 0.7918 | 0.0669 | 0.0669 | 0.8028 | 0.7899 | 0.0749 | −1.8621 | 1.0000 | 1.0000 |
| $N = 32$ | 0.7961 | 0.0465 | 0.0466 | 0.8017 | 0.7965 | 0.0479 | 0.6755 | 1.0000 | 1.0000 |
| $N = 64$ | 0.7981 | 0.0326 | 0.0327 | 0.8001 | 0.7975 | 0.0332 | −1.1567 | 1.0000 | 1.0000 |
| $N = 128$ | 0.7991 | 0.0230 | 0.0230 | 0.8008 | 0.7994 | 0.0232 | 0.9506 | 1.0000 | 1.0000 |
| $N = 256$ | 0.7995 | 0.0162 | 0.0162 | 0.7999 | 0.7993 | 0.0162 | −1.2035 | 1.0000 | 1.0000 |

$ASD_K$: Asymptotic standard deviation of $V_{rel}(\mathbf{R})$ based on Konishi's (1979) theory (eq. 39).
$ASD_{PF}$: Approximate standard deviation of $V_{rel}(\mathbf{R})$ based on Pan & Frank's (2004) approach (eqs. 36–38).

(*continued*)

**Table S3.** (*continued*)

| | E[$V_{rel}(\mathbf{R})$] | $ASD_K$ | $ASD_{PF}$ | Median | Mean | ESD | $T$ | Pow. 5% | Pow. 1% |
|---|---|---|---|---|---|---|---|---|---|
| $p = 8$, $q = 2$, $V_{rel}(\mathbf{P}) = 0.1$ | | | | | | | | | |
| $N = 4$ | 0.3763 | 0.0874 | 0.1278 | 0.3606 | 0.3752 | 0.0815 | −0.9284 | 0.1786 | 0.0626 |
| $N = 8$ | 0.2126 | 0.0572 | 0.0705 | 0.2057 | 0.2129 | 0.0588 | 0.4217 | 0.5632 | 0.3392 |
| $N = 16$ | 0.1511 | 0.0391 | 0.0437 | 0.1473 | 0.1509 | 0.0402 | −0.4468 | 0.9354 | 0.8492 |
| $N = 32$ | 0.1244 | 0.0272 | 0.0288 | 0.1231 | 0.1244 | 0.0277 | −0.0301 | 1.0000 | 0.9984 |
| $N = 64$ | 0.1119 | 0.0191 | 0.0196 | 0.1107 | 0.1112 | 0.0193 | −2.8395 | 1.0000 | 1.0000 |
| $N = 128$ | 0.1059 | 0.0134 | 0.0136 | 0.1058 | 0.1060 | 0.0135 | 0.7249 | 1.0000 | 1.0000 |
| $N = 256$ | 0.1029 | 0.0095 | 0.0095 | 0.1029 | 0.1029 | 0.0095 | −0.3405 | 1.0000 | 1.0000 |
| $p = 8$, $q = 2$, $V_{rel}(\mathbf{P}) = 0.2$ | | | | | | | | | |
| $N = 4$ | 0.4270 | 0.0943 | 0.1738 | 0.4066 | 0.4284 | 0.1047 | 0.9149 | 0.3598 | 0.1860 |
| $N = 8$ | 0.2895 | 0.0617 | 0.0891 | 0.2813 | 0.2891 | 0.0748 | −0.4132 | 0.9128 | 0.7928 |
| $N = 16$ | 0.2403 | 0.0422 | 0.0521 | 0.2369 | 0.2398 | 0.0502 | −0.7208 | 0.9996 | 0.9974 |
| $N = 32$ | 0.2192 | 0.0293 | 0.0329 | 0.2186 | 0.2191 | 0.0325 | −0.2502 | 1.0000 | 1.0000 |
| $N = 64$ | 0.2094 | 0.0206 | 0.0219 | 0.2092 | 0.2091 | 0.0220 | −0.7467 | 1.0000 | 1.0000 |
| $N = 128$ | 0.2046 | 0.0145 | 0.0150 | 0.2045 | 0.2046 | 0.0149 | −0.1112 | 1.0000 | 1.0000 |
| $N = 256$ | 0.2023 | 0.0102 | 0.0104 | 0.2024 | 0.2023 | 0.0101 | −0.1530 | 1.0000 | 1.0000 |
| $p = 8$, $q = 2$, $V_{rel}(\mathbf{P}) = 0.4$ | | | | | | | | | |
| $N = 4$ | 0.5793 | 0.0182 | 0.2517 | 0.5317 | 0.5781 | 0.1562 | −0.5295 | 0.7550 | 0.5354 |
| $N = 8$ | 0.4772 | 0.0119 | 0.1092 | 0.4436 | 0.4764 | 0.0891 | −0.6747 | 1.0000 | 1.0000 |
| $N = 16$ | 0.4363 | 0.0082 | 0.0516 | 0.4192 | 0.4355 | 0.0469 | −1.0874 | 1.0000 | 1.0000 |
| $N = 32$ | 0.4176 | 0.0057 | 0.0254 | 0.4098 | 0.4178 | 0.0244 | 0.5427 | 1.0000 | 1.0000 |
| $N = 64$ | 0.4087 | 0.0040 | 0.0128 | 0.4053 | 0.4086 | 0.0126 | −0.1456 | 1.0000 | 1.0000 |
| $N = 128$ | 0.4043 | 0.0028 | 0.0067 | 0.4030 | 0.4043 | 0.0066 | 0.2277 | 1.0000 | 1.0000 |
| $N = 256$ | 0.4021 | 0.0020 | 0.0036 | 0.4016 | 0.4022 | 0.0037 | 0.5998 | 1.0000 | 1.0000 |
| $p = 8$, $q = 4$, $V_{rel}(\mathbf{P}) = 0.1$ | | | | | | | | | |
| $N = 4$ | 0.3856 | 0.0207 | 0.1264 | 0.3724 | 0.3860 | 0.0845 | 0.3568 | 0.2128 | 0.0802 |
| $N = 8$ | 0.2206 | 0.0136 | 0.0583 | 0.2147 | 0.2213 | 0.0493 | 0.9737 | 0.6566 | 0.3774 |
| $N = 16$ | 0.1561 | 0.0093 | 0.0289 | 0.1529 | 0.1564 | 0.0271 | 0.8081 | 0.9998 | 0.9846 |
| $N = 32$ | 0.1271 | 0.0064 | 0.0149 | 0.1256 | 0.1271 | 0.0142 | 0.1225 | 1.0000 | 1.0000 |
| $N = 64$ | 0.1133 | 0.0045 | 0.0081 | 0.1127 | 0.1134 | 0.0080 | 0.7534 | 1.0000 | 1.0000 |
| $N = 128$ | 0.1066 | 0.0032 | 0.0046 | 0.1064 | 0.1066 | 0.0046 | −0.6650 | 1.0000 | 1.0000 |
| $N = 256$ | 0.1033 | 0.0022 | 0.0028 | 0.1032 | 0.1033 | 0.0028 | −1.0577 | 1.0000 | 1.0000 |
| $p = 8$, $q = 4$, $V_{rel}(\mathbf{P}) = 0.2$ | | | | | | | | | |
| (This comformation is impossible) | | | | | | | | | |

$ASD_K$: Asymptotic standard deviation of $V_{rel}(\mathbf{R})$ based on Konishi's (1979) theory (eq. 39).
$ASD_{PF}$: Approximate standard deviation of $V_{rel}(\mathbf{R})$ based on Pan & Frank's (2004) approach (eqs. 36–38).

(*continued*)

**Table S3.** (*continued*)

| | $E[V_{rel}(\mathbf{R})]$ | $ASD_K$ | $ASD_{PF}$ | Median | Mean | ESD | $T$ | Pow. 5% | Pow. 1% |
|---|---|---|---|---|---|---|---|---|---|
| $p = 16$, $q = 1$, $V_{rel}(\mathbf{P}) = 0.1$ | | | | | | | | | |
| $N = 4$ | 0.3745 | 0.1309 | 0.1291 | 0.3567 | 0.3753 | 0.0655 | 0.8685 | 0.3218 | 0.2192 |
| $N = 8$ | 0.2108 | 0.0857 | 0.0847 | 0.1975 | 0.2120 | 0.0627 | 1.3379 | 0.7088 | 0.5934 |
| $N = 16$ | 0.1499 | 0.0585 | 0.0582 | 0.1417 | 0.1494 | 0.0505 | −0.6621 | 0.9526 | 0.9194 |
| $N = 32$ | 0.1237 | 0.0407 | 0.0406 | 0.1198 | 0.1236 | 0.0382 | −0.1200 | 0.9994 | 0.9990 |
| $N = 64$ | 0.1115 | 0.0286 | 0.0285 | 0.1090 | 0.1117 | 0.0279 | 0.3795 | 1.0000 | 1.0000 |
| $N = 128$ | 0.1057 | 0.0201 | 0.0201 | 0.1049 | 0.1058 | 0.0199 | 0.4730 | 1.0000 | 1.0000 |
| $N = 256$ | 0.1028 | 0.0142 | 0.0142 | 0.1024 | 0.1028 | 0.0141 | 0.0272 | 1.0000 | 1.0000 |
| $p = 16$, $q = 1$, $V_{rel}(\mathbf{P}) = 0.2$ | | | | | | | | | |
| $N = 4$ | 0.4184 | 0.2009 | 0.1879 | 0.3923 | 0.4189 | 0.0975 | 0.3846 | 0.5308 | 0.4138 |
| $N = 8$ | 0.2814 | 0.1315 | 0.1273 | 0.2670 | 0.2819 | 0.0970 | 0.3873 | 0.8970 | 0.8414 |
| $N = 16$ | 0.2350 | 0.0898 | 0.0884 | 0.2273 | 0.2353 | 0.0780 | 0.2734 | 0.9974 | 0.9944 |
| $N = 32$ | 0.2162 | 0.0625 | 0.0620 | 0.2130 | 0.2152 | 0.0580 | −1.2665 | 1.0000 | 1.0000 |
| $N = 64$ | 0.2078 | 0.0438 | 0.0437 | 0.2068 | 0.2077 | 0.0425 | −0.1913 | 1.0000 | 1.0000 |
| $N = 128$ | 0.2038 | 0.0309 | 0.0308 | 0.2026 | 0.2032 | 0.0303 | −1.4818 | 1.0000 | 1.0000 |
| $N = 256$ | 0.2019 | 0.0218 | 0.0218 | 0.2014 | 0.2018 | 0.0216 | −0.1586 | 1.0000 | 1.0000 |
| $p = 16$, $q = 1$, $V_{rel}(\mathbf{P}) = 0.4$ | | | | | | | | | |
| $N = 4$ | 0.5158 | 0.2570 | 0.2347 | 0.4947 | 0.5167 | 0.1433 | 0.4595 | 0.7744 | 0.6966 |
| $N = 8$ | 0.4318 | 0.1682 | 0.1621 | 0.4321 | 0.4329 | 0.1391 | 0.5780 | 0.9820 | 0.9714 |
| $N = 16$ | 0.4111 | 0.1149 | 0.1130 | 0.4149 | 0.4120 | 0.1073 | 0.5980 | 1.0000 | 1.0000 |
| $N = 32$ | 0.4046 | 0.0799 | 0.0793 | 0.4084 | 0.4055 | 0.0773 | 0.7996 | 1.0000 | 1.0000 |
| $N = 64$ | 0.4021 | 0.0561 | 0.0559 | 0.4030 | 0.4021 | 0.0552 | −0.0283 | 1.0000 | 1.0000 |
| $N = 128$ | 0.4010 | 0.0395 | 0.0394 | 0.4022 | 0.4019 | 0.0395 | 1.7153 | 1.0000 | 1.0000 |
| $N = 256$ | 0.4005 | 0.0279 | 0.0278 | 0.4005 | 0.4003 | 0.0281 | −0.4201 | 1.0000 | 1.0000 |
| $p = 16$, $q = 1$, $V_{rel}(\mathbf{P}) = 0.6$ | | | | | | | | | |
| $N = 4$ | 0.6313 | 0.2322 | 0.2143 | 0.6468 | 0.6306 | 0.1707 | −0.2806 | 0.8956 | 0.8606 |
| $N = 8$ | 0.5973 | 0.1520 | 0.1477 | 0.6162 | 0.5970 | 0.1467 | −0.1359 | 0.9974 | 0.9960 |
| $N = 16$ | 0.5963 | 0.1039 | 0.1026 | 0.6084 | 0.5967 | 0.1047 | 0.3048 | 1.0000 | 1.0000 |
| $N = 32$ | 0.5978 | 0.0722 | 0.0718 | 0.6040 | 0.5979 | 0.0726 | 0.1795 | 1.0000 | 1.0000 |
| $N = 64$ | 0.5988 | 0.0507 | 0.0505 | 0.6013 | 0.5986 | 0.0510 | −0.2188 | 1.0000 | 1.0000 |
| $N = 128$ | 0.5994 | 0.0357 | 0.0356 | 0.6010 | 0.5997 | 0.0362 | 0.6137 | 1.0000 | 1.0000 |
| $N = 256$ | 0.5997 | 0.0252 | 0.0252 | 0.6001 | 0.5994 | 0.0253 | −0.8911 | 1.0000 | 1.0000 |
| $p = 16$, $q = 1$, $V_{rel}(\mathbf{P}) = 0.8$ | | | | | | | | | |
| $N = 4$ | 0.7768 | 0.1435 | 0.1368 | 0.8260 | 0.7751 | 0.1589 | −0.7708 | 0.9692 | 0.9554 |
| $N = 8$ | 0.7831 | 0.0939 | 0.0927 | 0.8102 | 0.7835 | 0.1116 | 0.2370 | 0.9998 | 0.9998 |
| $N = 16$ | 0.7918 | 0.0642 | 0.0638 | 0.8045 | 0.7913 | 0.0716 | −0.5326 | 1.0000 | 1.0000 |
| $N = 32$ | 0.7961 | 0.0446 | 0.0445 | 0.8035 | 0.7976 | 0.0457 | 2.4043 | 1.0000 | 1.0000 |
| $N = 64$ | 0.7981 | 0.0313 | 0.0313 | 0.8013 | 0.7984 | 0.0325 | 0.7108 | 1.0000 | 1.0000 |
| $N = 128$ | 0.7991 | 0.0221 | 0.0220 | 0.7997 | 0.7985 | 0.0224 | −1.8772 | 1.0000 | 1.0000 |
| $N = 256$ | 0.7995 | 0.0156 | 0.0156 | 0.7998 | 0.7995 | 0.0157 | −0.3465 | 1.0000 | 1.0000 |

$ASD_K$: Asymptotic standard deviation of $V_{rel}(\mathbf{R})$ based on Konishi's (1979) theory (eq. 39).
$ASD_{PF}$: Approximate standard deviation of $V_{rel}(\mathbf{R})$ based on Pan & Frank's (2004) approach (eqs. 36–38).

(*continued*)

**Table S3.** (*continued*)

| | E[$V_{rel}$(**R**)] | $ASD_K$ | $ASD_{PF}$ | Median | Mean | ESD | *T* | Pow. 5% | Pow. 1% |
|---|---|---|---|---|---|---|---|---|---|
| $p = 16$, $q = 2$, $V_{rel}(\mathbf{P}) = 0.1$ | | | | | | | | | |
| $N = 4$ | 0.3760 | 0.0759 | 0.1026 | 0.3629 | 0.3754 | 0.0568 | −0.7850 | 0.3492 | 0.2086 |
| $N = 8$ | 0.2123 | 0.0497 | 0.0579 | 0.2064 | 0.2116 | 0.0442 | −1.0713 | 0.8342 | 0.7150 |
| $N = 16$ | 0.1510 | 0.0340 | 0.0367 | 0.1484 | 0.1508 | 0.0321 | −0.2952 | 0.9966 | 0.9900 |
| $N = 32$ | 0.1243 | 0.0236 | 0.0246 | 0.1239 | 0.1246 | 0.0236 | 0.8037 | 1.0000 | 1.0000 |
| $N = 64$ | 0.1119 | 0.0166 | 0.0169 | 0.1114 | 0.1120 | 0.0165 | 0.5617 | 1.0000 | 1.0000 |
| $N = 128$ | 0.1059 | 0.0117 | 0.0118 | 0.1055 | 0.1057 | 0.0116 | −1.1433 | 1.0000 | 1.0000 |
| $N = 256$ | 0.1029 | 0.0082 | 0.0083 | 0.1024 | 0.1027 | 0.0083 | −1.6560 | 1.0000 | 1.0000 |
| $p = 16$, $q = 2$, $V_{rel}(\mathbf{P}) = 0.2$ | | | | | | | | | |
| $N = 4$ | 0.4256 | 0.0909 | 0.1483 | 0.4075 | 0.4256 | 0.0827 | 0.0263 | 0.6414 | 0.4660 |
| $N = 8$ | 0.2882 | 0.0595 | 0.0781 | 0.2814 | 0.2885 | 0.0624 | 0.2671 | 0.9890 | 0.9756 |
| $N = 16$ | 0.2395 | 0.0406 | 0.0471 | 0.2368 | 0.2388 | 0.0439 | −1.0447 | 1.0000 | 1.0000 |
| $N = 32$ | 0.2188 | 0.0283 | 0.0305 | 0.2180 | 0.2186 | 0.0298 | −0.2548 | 1.0000 | 1.0000 |
| $N = 64$ | 0.2091 | 0.0198 | 0.0206 | 0.2087 | 0.2084 | 0.0205 | −2.4858 | 1.0000 | 1.0000 |
| $N = 128$ | 0.2045 | 0.0140 | 0.0143 | 0.2045 | 0.2044 | 0.0142 | −0.7333 | 1.0000 | 1.0000 |
| $N = 256$ | 0.2022 | 0.0099 | 0.0100 | 0.2024 | 0.2025 | 0.0096 | 1.5185 | 1.0000 | 1.0000 |
| $p = 16$, $q = 2$, $V_{rel}(\mathbf{P}) = 0.4$ | | | | | | | | | |
| $N = 4$ | 0.5640 | 0.0370 | 0.2217 | 0.5251 | 0.5645 | 0.1328 | 0.2332 | 0.9850 | 0.9482 |
| $N = 8$ | 0.4687 | 0.0242 | 0.0971 | 0.4450 | 0.4679 | 0.0790 | −0.7910 | 1.0000 | 1.0000 |
| $N = 16$ | 0.4322 | 0.0165 | 0.0470 | 0.4220 | 0.4323 | 0.0439 | 0.2191 | 1.0000 | 1.0000 |
| $N = 32$ | 0.4156 | 0.0115 | 0.0242 | 0.4115 | 0.4153 | 0.0229 | −1.0161 | 1.0000 | 1.0000 |
| $N = 64$ | 0.4077 | 0.0081 | 0.0132 | 0.4064 | 0.4076 | 0.0128 | −0.6659 | 1.0000 | 1.0000 |
| $N = 128$ | 0.4038 | 0.0057 | 0.0077 | 0.4033 | 0.4039 | 0.0078 | 0.2858 | 1.0000 | 1.0000 |
| $N = 256$ | 0.4019 | 0.0040 | 0.0048 | 0.4017 | 0.4018 | 0.0048 | −1.0195 | 1.0000 | 1.0000 |
| $p = 16$, $q = 4$, $V_{rel}(\mathbf{P}) = 0.1$ | | | | | | | | | |
| $N = 4$ | 0.3808 | 0.0305 | 0.0953 | 0.3711 | 0.3802 | 0.0542 | −0.7788 | 0.3896 | 0.2272 |
| $N = 8$ | 0.2168 | 0.0199 | 0.0444 | 0.2115 | 0.2161 | 0.0343 | −1.4581 | 0.9428 | 0.8348 |
| $N = 16$ | 0.1538 | 0.0136 | 0.0232 | 0.1516 | 0.1542 | 0.0213 | 1.0745 | 1.0000 | 1.0000 |
| $N = 32$ | 0.1259 | 0.0095 | 0.0132 | 0.1249 | 0.1255 | 0.0127 | −2.3988 | 1.0000 | 1.0000 |
| $N = 64$ | 0.1127 | 0.0066 | 0.0080 | 0.1127 | 0.1127 | 0.0079 | −0.1133 | 1.0000 | 1.0000 |
| $N = 128$ | 0.1063 | 0.0047 | 0.0052 | 0.1063 | 0.1064 | 0.0052 | 1.6837 | 1.0000 | 1.0000 |
| $N = 256$ | 0.1031 | 0.0033 | 0.0035 | 0.1031 | 0.1031 | 0.0035 | −0.2266 | 1.0000 | 1.0000 |
| $p = 16$, $q = 4$, $V_{rel}(\mathbf{P}) = 0.2$ | | | | | | | | | |
| $N = 4$ | 0.4667 | $0^*$ | 0.1510 | 0.4502 | 0.4641 | 0.0972 | −1.8454 | 0.7790 | 0.6708 |
| $N = 8$ | 0.3143 | $0^*$ | 0.0653 | 0.3049 | 0.3131 | 0.0529 | −1.6415 | 1.0000 | 1.0000 |
| $N = 16$ | 0.2533 | $0^*$ | 0.0306 | 0.2485 | 0.2528 | 0.0275 | −1.2473 | 1.0000 | 1.0000 |
| $N = 32$ | 0.2258 | $0^*$ | 0.0149 | 0.2234 | 0.2258 | 0.0142 | 0.1036 | 1.0000 | 1.0000 |
| $N = 64$ | 0.2127 | $0^*$ | 0.0073 | 0.2113 | 0.2126 | 0.0071 | −1.1940 | 1.0000 | 1.0000 |
| $N = 128$ | 0.2063 | $0^*$ | 0.0036 | 0.2057 | 0.2064 | 0.0036 | 1.7598 | 1.0000 | 1.0000 |
| $N = 256$ | 0.2031 | $0^*$ | 0.0018 | 0.2028 | 0.2031 | 0.0018 | −0.4979 | 1.0000 | 1.0000 |

$ASD_K$: Asymptotic standard deviation of $V_{rel}$(**R**) based on Konishi's (1979) theory (eq. 39).
$ASD_{PF}$: Approximate standard deviation of $V_{rel}$(**R**) based on Pan & Frank's (2004) approach (eqs. 36–38).
*In this case, **P** is singular and the asymptotic expression yields 0 (which is obviously spurious).

(*continued*)

**Table S3.** (*continued*)

| | $E[V_{rel}(\mathbf{R})]$ | $ASD_K$ | $ASD_{PF}$ | Median | Mean | ESD | $T$ | Pow. 5% | Pow. 1% |
|---|---|---|---|---|---|---|---|---|---|
| $p = 32, q = 1, V_{rel}(\mathbf{P}) = 0.1$ | | | | | | | | | |
| $N = 4$ | 0.3745 | 0.1211 | 0.1175 | 0.3560 | 0.3737 | 0.0551 | −1.1234 | 0.4740 | 0.3322 |
| $N = 8$ | 0.2108 | 0.0793 | 0.0779 | 0.1957 | 0.2087 | 0.0550 | −2.6857 | 0.8568 | 0.8010 |
| $N = 16$ | 0.1499 | 0.0542 | 0.0537 | 0.1442 | 0.1511 | 0.0463 | 1.8301 | 0.9952 | 0.9878 |
| $N = 32$ | 0.1237 | 0.0377 | 0.0375 | 0.1203 | 0.1240 | 0.0345 | 0.6352 | 1.0000 | 1.0000 |
| $N = 64$ | 0.1115 | 0.0264 | 0.0264 | 0.1100 | 0.1117 | 0.0254 | 0.3329 | 1.0000 | 1.0000 |
| $N = 128$ | 0.1057 | 0.0186 | 0.0186 | 0.1046 | 0.1051 | 0.0184 | −2.2279 | 1.0000 | 1.0000 |
| $N = 256$ | 0.1028 | 0.0131 | 0.0131 | 0.1019 | 0.1024 | 0.0128 | −2.1683 | 1.0000 | 1.0000 |
| $p = 32, q = 1, V_{rel}(\mathbf{P}) = 0.2$ | | | | | | | | | |
| $N = 4$ | 0.4184 | 0.1905 | 0.1776 | 0.3874 | 0.4152 | 0.0870 | −2.5287 | 0.6732 | 0.5566 |
| $N = 8$ | 0.2814 | 0.1247 | 0.1207 | 0.2661 | 0.2821 | 0.0911 | 0.5770 | 0.9640 | 0.9422 |
| $N = 16$ | 0.2350 | 0.0852 | 0.0839 | 0.2265 | 0.2329 | 0.0728 | −2.0126 | 0.9996 | 0.9996 |
| $N = 32$ | 0.2162 | 0.0593 | 0.0588 | 0.2132 | 0.2168 | 0.0560 | 0.7146 | 1.0000 | 1.0000 |
| $N = 64$ | 0.2078 | 0.0416 | 0.0414 | 0.2069 | 0.2084 | 0.0400 | 1.1078 | 1.0000 | 1.0000 |
| $N = 128$ | 0.2038 | 0.0293 | 0.0292 | 0.2038 | 0.2044 | 0.0283 | 1.3960 | 1.0000 | 1.0000 |
| $N = 256$ | 0.2019 | 0.0207 | 0.0206 | 0.2017 | 0.2020 | 0.0202 | 0.4522 | 1.0000 | 1.0000 |
| $p = 32, q = 1, V_{rel}(\mathbf{P}) = 0.4$ | | | | | | | | | |
| $N = 4$ | 0.5158 | 0.2484 | 0.2271 | 0.4965 | 0.5174 | 0.1383 | 0.8217 | 0.8696 | 0.8084 |
| $N = 8$ | 0.4318 | 0.1626 | 0.1568 | 0.4294 | 0.4304 | 0.1324 | −0.7173 | 0.9966 | 0.9946 |
| $N = 16$ | 0.4111 | 0.1111 | 0.1093 | 0.4124 | 0.4104 | 0.1016 | −0.5116 | 1.0000 | 1.0000 |
| $N = 32$ | 0.4046 | 0.0773 | 0.0767 | 0.4082 | 0.4056 | 0.0764 | 0.9578 | 1.0000 | 1.0000 |
| $N = 64$ | 0.4021 | 0.0542 | 0.0540 | 0.4033 | 0.4021 | 0.0533 | 0.0552 | 1.0000 | 1.0000 |
| $N = 128$ | 0.4010 | 0.0382 | 0.0381 | 0.4021 | 0.4019 | 0.0383 | 1.6504 | 1.0000 | 1.0000 |
| $N = 256$ | 0.4005 | 0.0269 | 0.0269 | 0.4002 | 0.4002 | 0.0265 | −0.6218 | 1.0000 | 1.0000 |
| $p = 32, q = 1, V_{rel}(\mathbf{P}) = 0.6$ | | | | | | | | | |
| $N = 4$ | 0.6313 | 0.2264 | 0.2090 | 0.6504 | 0.6334 | 0.1637 | 0.9116 | 0.9400 | 0.9140 |
| $N = 8$ | 0.5973 | 0.1482 | 0.1439 | 0.6168 | 0.5978 | 0.1394 | 0.2755 | 1.0000 | 0.9998 |
| $N = 16$ | 0.5963 | 0.1013 | 0.1000 | 0.6071 | 0.5952 | 0.1026 | −0.7337 | 1.0000 | 1.0000 |
| $N = 32$ | 0.5978 | 0.0704 | 0.0700 | 0.6041 | 0.5976 | 0.0717 | −0.1040 | 1.0000 | 1.0000 |
| $N = 64$ | 0.5988 | 0.0494 | 0.0493 | 0.6023 | 0.5994 | 0.0496 | 0.8511 | 1.0000 | 1.0000 |
| $N = 128$ | 0.5994 | 0.0348 | 0.0348 | 0.5994 | 0.5986 | 0.0349 | −1.5793 | 1.0000 | 1.0000 |
| $N = 256$ | 0.5997 | 0.0246 | 0.0245 | 0.6006 | 0.5997 | 0.0245 | −0.0753 | 1.0000 | 1.0000 |
| $p = 32, q = 1, V_{rel}(\mathbf{P}) = 0.8$ | | | | | | | | | |
| $N = 4$ | 0.7768 | 0.1406 | 0.1336 | 0.8222 | 0.7737 | 0.1558 | −1.3932 | 0.9822 | 0.9742 |
| $N = 8$ | 0.7831 | 0.0921 | 0.0906 | 0.8099 | 0.7846 | 0.1075 | 0.9636 | 1.0000 | 1.0000 |
| $N = 16$ | 0.7918 | 0.0629 | 0.0625 | 0.8024 | 0.7906 | 0.0703 | −1.2364 | 1.0000 | 1.0000 |
| $N = 32$ | 0.7961 | 0.0438 | 0.0436 | 0.8022 | 0.7965 | 0.0458 | 0.6030 | 1.0000 | 1.0000 |
| $N = 64$ | 0.7981 | 0.0307 | 0.0306 | 0.8001 | 0.7977 | 0.0310 | −0.9765 | 1.0000 | 1.0000 |
| $N = 128$ | 0.7991 | 0.0216 | 0.0216 | 0.8008 | 0.7995 | 0.0215 | 1.4951 | 1.0000 | 1.0000 |
| $N = 256$ | 0.7995 | 0.0153 | 0.0152 | 0.7999 | 0.7992 | 0.0154 | −1.2828 | 1.0000 | 1.0000 |

$ASD_K$: Asymptotic standard deviation of $V_{rel}(\mathbf{R})$ based on Konishi's (1979) theory (eq. 39).
$ASD_{PF}$: Approximate standard deviation of $V_{rel}(\mathbf{R})$ based on Pan & Frank's (2004) approach (eqs. 36–38).

(*continued*)

**Table S3.** (*continued*)

| | E[$V_{rel}$(**R**)] | $ASD_K$ | $ASD_{PF}$ | Median | Mean | ESD | *T* | Pow. 5% | Pow. 1% |
|---|---|---|---|---|---|---|---|---|---|
| $p = 32, q = 2, V_{rel}(\mathbf{P}) = 0.1$ | | | | | | | | | |
| *N* = 4 | 0.3759 | 0.0699 | 0.0895 | 0.3654 | 0.3756 | 0.0446 | −0.5018 | 0.5744 | 0.3864 |
| *N* = 8 | 0.2122 | 0.0458 | 0.0516 | 0.2063 | 0.2115 | 0.0385 | −1.2294 | 0.9618 | 0.9252 |
| *N* = 16 | 0.1509 | 0.0313 | 0.0332 | 0.1477 | 0.1500 | 0.0289 | −2.1212 | 0.9998 | 0.9998 |
| *N* = 32 | 0.1243 | 0.0218 | 0.0224 | 0.1230 | 0.1241 | 0.0211 | −0.6802 | 1.0000 | 1.0000 |
| *N* = 64 | 0.1119 | 0.0153 | 0.0155 | 0.1115 | 0.1119 | 0.0150 | 0.3859 | 1.0000 | 1.0000 |
| *N* = 128 | 0.1059 | 0.0107 | 0.0108 | 0.1056 | 0.1058 | 0.0105 | −0.4723 | 1.0000 | 1.0000 |
| *N* = 256 | 0.1029 | 0.0076 | 0.0076 | 0.1030 | 0.1030 | 0.0075 | 1.0315 | 1.0000 | 1.0000 |
| $p = 32, q = 2, V_{rel}(\mathbf{P}) = 0.2$ | | | | | | | | | |
| *N* = 4 | 0.4250 | 0.0881 | 0.1354 | 0.4112 | 0.4255 | 0.0721 | 0.4438 | 0.8450 | 0.7344 |
| *N* = 8 | 0.2877 | 0.0577 | 0.0727 | 0.2840 | 0.2892 | 0.0578 | 1.7986 | 0.9992 | 0.9984 |
| *N* = 16 | 0.2392 | 0.0394 | 0.0445 | 0.2385 | 0.2394 | 0.0402 | 0.3791 | 1.0000 | 1.0000 |
| *N* = 32 | 0.2186 | 0.0274 | 0.0292 | 0.2183 | 0.2188 | 0.0285 | 0.5549 | 1.0000 | 1.0000 |
| *N* = 64 | 0.2091 | 0.0192 | 0.0199 | 0.2095 | 0.2094 | 0.0199 | 1.3979 | 1.0000 | 1.0000 |
| *N* = 128 | 0.2045 | 0.0135 | 0.0138 | 0.2045 | 0.2044 | 0.0136 | −0.3347 | 1.0000 | 1.0000 |
| *N* = 256 | 0.2022 | 0.0096 | 0.0096 | 0.2022 | 0.2022 | 0.0095 | −0.1785 | 1.0000 | 1.0000 |
| $p = 32, q = 2, V_{rel}(\mathbf{P}) = 0.4$ | | | | | | | | | |
| *N* = 4 | 0.5591 | 0.0436 | 0.2077 | 0.5191 | 0.5568 | 0.1217 | −1.3170 | 0.9976 | 0.9924 |
| *N* = 8 | 0.4656 | 0.0285 | 0.0917 | 0.4458 | 0.4652 | 0.0759 | −0.3893 | 1.0000 | 1.0000 |
| *N* = 16 | 0.4306 | 0.0195 | 0.0451 | 0.4235 | 0.4311 | 0.0424 | 0.7716 | 1.0000 | 1.0000 |
| *N* = 32 | 0.4148 | 0.0136 | 0.0239 | 0.4117 | 0.4149 | 0.0238 | 0.1904 | 1.0000 | 1.0000 |
| *N* = 64 | 0.4073 | 0.0095 | 0.0136 | 0.4065 | 0.4073 | 0.0134 | −0.2999 | 1.0000 | 1.0000 |
| *N* = 128 | 0.4036 | 0.0067 | 0.0082 | 0.4035 | 0.4037 | 0.0083 | 0.3745 | 1.0000 | 1.0000 |
| *N* = 256 | 0.4018 | 0.0047 | 0.0053 | 0.4019 | 0.4019 | 0.0053 | 1.3114 | 1.0000 | 1.0000 |
| $p = 32, q = 4, V_{rel}(\mathbf{P}) = 0.1$ | | | | | | | | | |
| *N* = 4 | 0.3797 | 0.0310 | 0.0801 | 0.3717 | 0.3802 | 0.0418 | 0.8412 | 0.6484 | 0.4402 |
| *N* = 8 | 0.2158 | 0.0203 | 0.0378 | 0.2114 | 0.2156 | 0.0293 | −0.4720 | 0.9966 | 0.9904 |
| *N* = 16 | 0.1532 | 0.0139 | 0.0204 | 0.1517 | 0.1533 | 0.0184 | 0.3585 | 1.0000 | 1.0000 |
| *N* = 32 | 0.1256 | 0.0097 | 0.0121 | 0.1251 | 0.1255 | 0.0117 | −0.3757 | 1.0000 | 1.0000 |
| *N* = 64 | 0.1126 | 0.0068 | 0.0077 | 0.1126 | 0.1127 | 0.0076 | 1.0420 | 1.0000 | 1.0000 |
| *N* = 128 | 0.1062 | 0.0048 | 0.0051 | 0.1063 | 0.1062 | 0.0051 | −0.1750 | 1.0000 | 1.0000 |
| *N* = 256 | 0.1031 | 0.0034 | 0.0035 | 0.1031 | 0.1031 | 0.0034 | −0.5193 | 1.0000 | 1.0000 |
| $p = 32, q = 4, V_{rel}(\mathbf{P}) = 0.2$ | | | | | | | | | |
| *N* = 4 | 0.4513 | 0.0104 | 0.1337 | 0.4391 | 0.4515 | 0.0789 | 0.2035 | 0.9148 | 0.8394 |
| *N* = 8 | 0.3073 | 0.0068 | 0.0577 | 0.3008 | 0.3073 | 0.0460 | −0.0126 | 1.0000 | 1.0000 |
| *N* = 16 | 0.2502 | 0.0046 | 0.0272 | 0.2459 | 0.2497 | 0.0246 | −1.3316 | 1.0000 | 1.0000 |
| *N* = 32 | 0.2243 | 0.0032 | 0.0134 | 0.2220 | 0.2240 | 0.0126 | −1.7985 | 1.0000 | 1.0000 |
| *N* = 64 | 0.2120 | 0.0023 | 0.0068 | 0.2110 | 0.2118 | 0.0065 | −1.2715 | 1.0000 | 1.0000 |
| *N* = 128 | 0.2059 | 0.0016 | 0.0035 | 0.2054 | 0.2060 | 0.0036 | 0.2589 | 1.0000 | 1.0000 |
| *N* = 256 | 0.2030 | 0.0011 | 0.0019 | 0.2028 | 0.2030 | 0.0019 | 0.3123 | 1.0000 | 1.0000 |

$ASD_K$: Asymptotic standard deviation of $V_{rel}$(**R**) based on Konishi's (1979) theory (eq. 39).
$ASD_{PF}$: Approximate standard deviation of $V_{rel}$(**R**) based on Pan & Frank's (2004) approach (eqs. 36–38).

(*continued*)

**Table S3.** (*continued*)

| | $E[V_{rel}(\mathbf{R})]$ | $ASD_K$ | $ASD_{PF}$ | Median | Mean | ESD | $T$ | Pow. 5% | Pow. 1% |
|---|---|---|---|---|---|---|---|---|---|
| $p = 64, q = 1, V_{rel}(\mathbf{P}) = 0.1$ | | | | | | | | | |
| $N = 4$ | 0.3745 | 0.1163 | 0.1115 | 0.3556 | 0.3741 | 0.0519 | −0.5438 | 0.6180 | 0.5168 |
| $N = 8$ | 0.2108 | 0.0762 | 0.0745 | 0.1968 | 0.2103 | 0.0532 | −0.6481 | 0.9514 | 0.9188 |
| $N = 16$ | 0.1499 | 0.0520 | 0.0514 | 0.1437 | 0.1497 | 0.0438 | −0.3383 | 0.9998 | 0.9988 |
| $N = 32$ | 0.1237 | 0.0362 | 0.0360 | 0.1202 | 0.1239 | 0.0334 | 0.4985 | 1.0000 | 1.0000 |
| $N = 64$ | 0.1115 | 0.0254 | 0.0253 | 0.1098 | 0.1114 | 0.0249 | −0.3128 | 1.0000 | 1.0000 |
| $N = 128$ | 0.1057 | 0.0179 | 0.0179 | 0.1050 | 0.1057 | 0.0174 | −0.1546 | 1.0000 | 1.0000 |
| $N = 256$ | 0.1028 | 0.0126 | 0.0126 | 0.1025 | 0.1029 | 0.0126 | 0.5461 | 1.0000 | 1.0000 |
| $p = 64, q = 1, V_{rel}(\mathbf{P}) = 0.2$ | | | | | | | | | |
| $N = 4$ | 0.4184 | 0.1855 | 0.1723 | 0.3898 | 0.4192 | 0.0880 | 0.6633 | 0.7800 | 0.7118 |
| $N = 8$ | 0.2814 | 0.1214 | 0.1174 | 0.2667 | 0.2808 | 0.0868 | −0.4795 | 0.9898 | 0.9808 |
| $N = 16$ | 0.2350 | 0.0829 | 0.0816 | 0.2309 | 0.2361 | 0.0718 | 1.1428 | 0.9998 | 0.9998 |
| $N = 32$ | 0.2162 | 0.0577 | 0.0573 | 0.2145 | 0.2169 | 0.0541 | 0.8648 | 1.0000 | 1.0000 |
| $N = 64$ | 0.2078 | 0.0405 | 0.0403 | 0.2059 | 0.2073 | 0.0391 | −0.9173 | 1.0000 | 1.0000 |
| $N = 128$ | 0.2038 | 0.0285 | 0.0285 | 0.2026 | 0.2033 | 0.0284 | −1.2701 | 1.0000 | 1.0000 |
| $N = 256$ | 0.2019 | 0.0201 | 0.0201 | 0.2015 | 0.2020 | 0.0199 | 0.2640 | 1.0000 | 1.0000 |
| $p = 64, q = 1, V_{rel}(\mathbf{P}) = 0.4$ | | | | | | | | | |
| $N = 4$ | 0.5158 | 0.2442 | 0.2232 | 0.4971 | 0.5163 | 0.1365 | 0.2233 | 0.9130 | 0.8796 |
| $N = 8$ | 0.4318 | 0.1598 | 0.1542 | 0.4366 | 0.4332 | 0.1304 | 0.7802 | 0.9978 | 0.9970 |
| $N = 16$ | 0.4111 | 0.1092 | 0.1074 | 0.4144 | 0.4129 | 0.1015 | 1.2079 | 1.0000 | 1.0000 |
| $N = 32$ | 0.4046 | 0.0760 | 0.0754 | 0.4052 | 0.4039 | 0.0733 | −0.6457 | 1.0000 | 1.0000 |
| $N = 64$ | 0.4021 | 0.0533 | 0.0531 | 0.4029 | 0.4021 | 0.0535 | 0.0493 | 1.0000 | 1.0000 |
| $N = 128$ | 0.4010 | 0.0375 | 0.0375 | 0.4022 | 0.4014 | 0.0373 | 0.8757 | 1.0000 | 1.0000 |
| $N = 256$ | 0.4005 | 0.0265 | 0.0265 | 0.4009 | 0.4006 | 0.0262 | 0.3899 | 1.0000 | 1.0000 |
| $p = 64, q = 1, V_{rel}(\mathbf{P}) = 0.6$ | | | | | | | | | |
| $N = 4$ | 0.6313 | 0.2236 | 0.2064 | 0.6456 | 0.6312 | 0.1616 | −0.0385 | 0.9674 | 0.9538 |
| $N = 8$ | 0.5973 | 0.1464 | 0.1421 | 0.6171 | 0.5990 | 0.1390 | 0.8570 | 0.9998 | 0.9998 |
| $N = 16$ | 0.5963 | 0.1000 | 0.0987 | 0.6063 | 0.5966 | 0.1005 | 0.2119 | 1.0000 | 1.0000 |
| $N = 32$ | 0.5978 | 0.0696 | 0.0692 | 0.6042 | 0.5981 | 0.0705 | 0.2972 | 1.0000 | 1.0000 |
| $N = 64$ | 0.5988 | 0.0488 | 0.0487 | 0.6020 | 0.5992 | 0.0487 | 0.5630 | 1.0000 | 1.0000 |
| $N = 128$ | 0.5994 | 0.0344 | 0.0343 | 0.6006 | 0.5993 | 0.0343 | −0.2839 | 1.0000 | 1.0000 |
| $N = 256$ | 0.5997 | 0.0243 | 0.0242 | 0.6004 | 0.5997 | 0.0242 | 0.1422 | 1.0000 | 1.0000 |
| $p = 64, q = 1, V_{rel}(\mathbf{P}) = 0.8$ | | | | | | | | | |
| $N = 4$ | 0.7768 | 0.1393 | 0.1322 | 0.8244 | 0.7723 | 0.1573 | −2.0189 | 0.9896 | 0.9850 |
| $N = 8$ | 0.7831 | 0.0912 | 0.0896 | 0.8098 | 0.7841 | 0.1083 | 0.6205 | 1.0000 | 1.0000 |
| $N = 16$ | 0.7918 | 0.0623 | 0.0619 | 0.8046 | 0.7928 | 0.0687 | 0.9579 | 1.0000 | 1.0000 |
| $N = 32$ | 0.7961 | 0.0433 | 0.0432 | 0.8009 | 0.7955 | 0.0452 | −0.9741 | 1.0000 | 1.0000 |
| $N = 64$ | 0.7981 | 0.0304 | 0.0303 | 0.8012 | 0.7983 | 0.0309 | 0.4419 | 1.0000 | 1.0000 |
| $N = 128$ | 0.7991 | 0.0214 | 0.0214 | 0.8010 | 0.7996 | 0.0217 | 1.7584 | 1.0000 | 1.0000 |
| $N = 256$ | 0.7995 | 0.0151 | 0.0151 | 0.8004 | 0.7995 | 0.0153 | −0.3298 | 1.0000 | 1.0000 |

$ASD_K$: Asymptotic standard deviation of $V_{rel}(\mathbf{R})$ based on Konishi's (1979) theory (eq. 39).
$ASD_{PF}$: Approximate standard deviation of $V_{rel}(\mathbf{R})$ based on Pan & Frank's (2004) approach (eqs. 36–38).

(*continued*)

**Table S3.** (*continued*)

| | E[$V_{rel}$(**R**)] | $ASD_K$ | $ASD_{PF}$ | Median | Mean | ESD | $T$ | Pow. 5% | Pow. 1% |
|---|---|---|---|---|---|---|---|---|---|
| $p = 64$, $q = 2$, $V_{rel}$(**P**) = 0.1 | | | | | | | | | |
| $N = 4$ | 0.3759 | 0.0669 | 0.0827 | 0.3657 | 0.3763 | 0.0404 | 0.7925 | 0.7720 | 0.6518 |
| $N = 8$ | 0.2122 | 0.0438 | 0.0484 | 0.2069 | 0.2121 | 0.0364 | −0.0292 | 0.9936 | 0.9878 |
| $N = 16$ | 0.1509 | 0.0299 | 0.0314 | 0.1487 | 0.1511 | 0.0280 | 0.6190 | 1.0000 | 1.0000 |
| $N = 32$ | 0.1242 | 0.0208 | 0.0213 | 0.1233 | 0.1241 | 0.0204 | −0.4629 | 1.0000 | 1.0000 |
| $N = 64$ | 0.1118 | 0.0146 | 0.0148 | 0.1114 | 0.1118 | 0.0142 | −0.0792 | 1.0000 | 1.0000 |
| $N = 128$ | 0.1059 | 0.0103 | 0.0103 | 0.1058 | 0.1060 | 0.0102 | 0.8201 | 1.0000 | 1.0000 |
| $N = 256$ | 0.1029 | 0.0073 | 0.0073 | 0.1029 | 0.1029 | 0.0072 | 0.1922 | 1.0000 | 1.0000 |
| $p = 64$, $q = 2$, $V_{rel}$(**P**) = 0.2 | | | | | | | | | |
| $N = 4$ | 0.4248 | 0.0866 | 0.1288 | 0.4108 | 0.4247 | 0.0675 | −0.0638 | 0.9378 | 0.8990 |
| $N = 8$ | 0.2875 | 0.0567 | 0.0699 | 0.2828 | 0.2886 | 0.0558 | 1.4147 | 1.0000 | 1.0000 |
| $N = 16$ | 0.2390 | 0.0387 | 0.0432 | 0.2379 | 0.2389 | 0.0400 | −0.2156 | 1.0000 | 1.0000 |
| $N = 32$ | 0.2185 | 0.0269 | 0.0285 | 0.2184 | 0.2186 | 0.0277 | 0.3109 | 1.0000 | 1.0000 |
| $N = 64$ | 0.2090 | 0.0189 | 0.0194 | 0.2090 | 0.2087 | 0.0189 | −1.1574 | 1.0000 | 1.0000 |
| $N = 128$ | 0.2045 | 0.0133 | 0.0135 | 0.2045 | 0.2046 | 0.0135 | 0.7096 | 1.0000 | 1.0000 |
| $N = 256$ | 0.2022 | 0.0094 | 0.0095 | 0.2024 | 0.2021 | 0.0095 | −0.4674 | 1.0000 | 1.0000 |
| $p = 64$, $q = 2$, $V_{rel}$(**P**) = 0.4 | | | | | | | | | |
| $N = 4$ | 0.5570 | 0.0463 | 0.2009 | 0.5220 | 0.5603 | 0.1204 | 1.9801 | 0.9990 | 0.9984 |
| $N = 8$ | 0.4643 | 0.0303 | 0.0892 | 0.4463 | 0.4627 | 0.0728 | −1.5320 | 1.0000 | 1.0000 |
| $N = 16$ | 0.4299 | 0.0207 | 0.0443 | 0.4241 | 0.4308 | 0.0405 | 1.6245 | 1.0000 | 1.0000 |
| $N = 32$ | 0.4145 | 0.0144 | 0.0238 | 0.4121 | 0.4149 | 0.0237 | 1.1474 | 1.0000 | 1.0000 |
| $N = 64$ | 0.4071 | 0.0101 | 0.0138 | 0.4063 | 0.4070 | 0.0137 | −0.7212 | 1.0000 | 1.0000 |
| $N = 128$ | 0.4035 | 0.0071 | 0.0085 | 0.4032 | 0.4035 | 0.0084 | −0.6864 | 1.0000 | 1.0000 |
| $N = 256$ | 0.4018 | 0.0050 | 0.0055 | 0.4016 | 0.4017 | 0.0056 | −0.8478 | 1.0000 | 1.0000 |
| $p = 64$, $q = 4$, $V_{rel}$(**P**) = 0.1 | | | | | | | | | |
| $N = 4$ | 0.3793 | 0.0307 | 0.0722 | 0.3714 | 0.3788 | 0.0354 | −0.9303 | 0.8442 | 0.7356 |
| $N = 8$ | 0.2154 | 0.0201 | 0.0345 | 0.2121 | 0.2150 | 0.0257 | −0.9970 | 1.0000 | 1.0000 |
| $N = 16$ | 0.1530 | 0.0137 | 0.0190 | 0.1517 | 0.1529 | 0.0170 | −0.3265 | 1.0000 | 1.0000 |
| $N = 32$ | 0.1255 | 0.0095 | 0.0115 | 0.1249 | 0.1255 | 0.0111 | 0.0425 | 1.0000 | 1.0000 |
| $N = 64$ | 0.1125 | 0.0067 | 0.0074 | 0.1125 | 0.1126 | 0.0074 | 0.9617 | 1.0000 | 1.0000 |
| $N = 128$ | 0.1062 | 0.0047 | 0.0050 | 0.1062 | 0.1061 | 0.0050 | −1.1861 | 1.0000 | 1.0000 |
| $N = 256$ | 0.1031 | 0.0033 | 0.0034 | 0.1031 | 0.1031 | 0.0034 | 0.4460 | 1.0000 | 1.0000 |
| $p = 64$, $q = 4$, $V_{rel}$(**P**) = 0.2 | | | | | | | | | |
| $N = 4$ | 0.4473 | 0.0142 | 0.1252 | 0.4362 | 0.4472 | 0.0703 | −0.0188 | 0.9738 | 0.9562 |
| $N = 8$ | 0.3049 | 0.0093 | 0.0542 | 0.2990 | 0.3053 | 0.0440 | 0.6636 | 1.0000 | 1.0000 |
| $N = 16$ | 0.2490 | 0.0063 | 0.0257 | 0.2455 | 0.2488 | 0.0231 | −0.5385 | 1.0000 | 1.0000 |
| $N = 32$ | 0.2237 | 0.0044 | 0.0128 | 0.2222 | 0.2237 | 0.0124 | 0.1394 | 1.0000 | 1.0000 |
| $N = 64$ | 0.2117 | 0.0031 | 0.0067 | 0.2108 | 0.2117 | 0.0067 | 0.4724 | 1.0000 | 1.0000 |
| $N = 128$ | 0.2058 | 0.0022 | 0.0037 | 0.2056 | 0.2058 | 0.0037 | 0.7531 | 1.0000 | 1.0000 |
| $N = 256$ | 0.2029 | 0.0015 | 0.0021 | 0.2028 | 0.2029 | 0.0021 | −0.1420 | 1.0000 | 1.0000 |

$ASD_K$: Asymptotic standard deviation of $V_{rel}$(**R**) based on Konishi's (1979) theory (eq. 39).
$ASD_{PF}$: Approximate standard deviation of $V_{rel}$(**R**) based on Pan & Frank's (2004) approach (eqs. 36–38).

(*continued*)

**Table S3.** (*continued*)

| | $E[V_{rel}(\mathbf{R})]$ | $ASD_K$ | $ASD_{PF}$ | Median | Mean | ESD | $T$ | Pow. 5% | Pow. 1% |
|---|---|---|---|---|---|---|---|---|---|
| $p = 128, q = 1, V_{rel}(\mathbf{P}) = 0.1$ | | | | | | | | | |
| $N = 4$ | 0.3745 | 0.1140 | 0.1085 | 0.3554 | 0.3736 | 0.0496 | −1.3663 | 0.7384 | 0.6688 |
| $N = 8$ | 0.2108 | 0.0746 | 0.0727 | 0.1999 | 0.2121 | 0.0532 | 1.7827 | 0.9838 | 0.9758 |
| $N = 16$ | 0.1499 | 0.0510 | 0.0503 | 0.1440 | 0.1502 | 0.0432 | 0.5427 | 1.0000 | 1.0000 |
| $N = 32$ | 0.1237 | 0.0355 | 0.0352 | 0.1216 | 0.1242 | 0.0325 | 1.1239 | 1.0000 | 1.0000 |
| $N = 64$ | 0.1115 | 0.0249 | 0.0248 | 0.1095 | 0.1111 | 0.0240 | −1.2279 | 1.0000 | 1.0000 |
| $N = 128$ | 0.1057 | 0.0175 | 0.0175 | 0.1045 | 0.1054 | 0.0172 | −1.0755 | 1.0000 | 1.0000 |
| $N = 256$ | 0.1028 | 0.0124 | 0.0124 | 0.1027 | 0.1028 | 0.0122 | −0.1024 | 1.0000 | 1.0000 |
| $p = 128, q = 1, V_{rel}(\mathbf{P}) = 0.2$ | | | | | | | | | |
| $N = 4$ | 0.4184 | 0.1830 | 0.1695 | 0.3917 | 0.4178 | 0.0829 | −0.5103 | 0.8690 | 0.8286 |
| $N = 8$ | 0.2814 | 0.1198 | 0.1157 | 0.2677 | 0.2831 | 0.0873 | 1.4148 | 0.9962 | 0.9950 |
| $N = 16$ | 0.2350 | 0.0818 | 0.0805 | 0.2291 | 0.2352 | 0.0723 | 0.2642 | 1.0000 | 1.0000 |
| $N = 32$ | 0.2162 | 0.0569 | 0.0565 | 0.2138 | 0.2164 | 0.0530 | 0.2451 | 1.0000 | 1.0000 |
| $N = 64$ | 0.2078 | 0.0399 | 0.0398 | 0.2071 | 0.2081 | 0.0383 | 0.5487 | 1.0000 | 1.0000 |
| $N = 128$ | 0.2038 | 0.0281 | 0.0281 | 0.2035 | 0.2045 | 0.0278 | 1.7509 | 1.0000 | 1.0000 |
| $N = 256$ | 0.2019 | 0.0198 | 0.0198 | 0.2014 | 0.2018 | 0.0197 | −0.4400 | 1.0000 | 1.0000 |
| $p = 128, q = 1, V_{rel}(\mathbf{P}) = 0.4$ | | | | | | | | | |
| $N = 4$ | 0.5158 | 0.2421 | 0.2212 | 0.4971 | 0.5166 | 0.1339 | 0.4348 | 0.9574 | 0.9414 |
| $N = 8$ | 0.4318 | 0.1585 | 0.1528 | 0.4334 | 0.4314 | 0.1291 | −0.2045 | 0.9996 | 0.9996 |
| $N = 16$ | 0.4111 | 0.1083 | 0.1065 | 0.4105 | 0.4082 | 0.0999 | −2.0899 | 1.0000 | 1.0000 |
| $N = 32$ | 0.4046 | 0.0753 | 0.0747 | 0.4064 | 0.4055 | 0.0722 | 0.9047 | 1.0000 | 1.0000 |
| $N = 64$ | 0.4021 | 0.0528 | 0.0526 | 0.4013 | 0.4019 | 0.0518 | −0.2472 | 1.0000 | 1.0000 |
| $N = 128$ | 0.4010 | 0.0372 | 0.0371 | 0.4018 | 0.4007 | 0.0373 | −0.5842 | 1.0000 | 1.0000 |
| $N = 256$ | 0.4005 | 0.0263 | 0.0262 | 0.4014 | 0.4012 | 0.0259 | 2.0216 | 1.0000 | 1.0000 |
| $p = 128, q = 1, V_{rel}(\mathbf{P}) = 0.6$ | | | | | | | | | |
| $N = 4$ | 0.6313 | 0.2222 | 0.2051 | 0.6428 | 0.6302 | 0.1624 | −0.4981 | 0.9826 | 0.9758 |
| $N = 8$ | 0.5973 | 0.1455 | 0.1412 | 0.6166 | 0.5969 | 0.1405 | −0.1875 | 1.0000 | 1.0000 |
| $N = 16$ | 0.5963 | 0.0994 | 0.0981 | 0.6073 | 0.5965 | 0.0993 | 0.1776 | 1.0000 | 1.0000 |
| $N = 32$ | 0.5978 | 0.0691 | 0.0687 | 0.6027 | 0.5975 | 0.0708 | −0.2100 | 1.0000 | 1.0000 |
| $N = 64$ | 0.5988 | 0.0485 | 0.0484 | 0.6024 | 0.5991 | 0.0489 | 0.4259 | 1.0000 | 1.0000 |
| $N = 128$ | 0.5994 | 0.0342 | 0.0341 | 0.6012 | 0.5997 | 0.0340 | 0.6170 | 1.0000 | 1.0000 |
| $N = 256$ | 0.5997 | 0.0241 | 0.0241 | 0.6005 | 0.5996 | 0.0242 | −0.2387 | 1.0000 | 1.0000 |
| $p = 128, q = 1, V_{rel}(\mathbf{P}) = 0.8$ | | | | | | | | | |
| $N = 4$ | 0.7768 | 0.1386 | 0.1315 | 0.8248 | 0.7760 | 0.1528 | −0.3781 | 0.9942 | 0.9918 |
| $N = 8$ | 0.7831 | 0.0907 | 0.0892 | 0.8035 | 0.7804 | 0.1076 | −1.8237 | 1.0000 | 1.0000 |
| $N = 16$ | 0.7918 | 0.0620 | 0.0615 | 0.8028 | 0.7905 | 0.0683 | −1.4145 | 1.0000 | 1.0000 |
| $N = 32$ | 0.7961 | 0.0431 | 0.0430 | 0.8009 | 0.7957 | 0.0453 | −0.5783 | 1.0000 | 1.0000 |
| $N = 64$ | 0.7981 | 0.0302 | 0.0302 | 0.8012 | 0.7985 | 0.0313 | 1.0163 | 1.0000 | 1.0000 |
| $N = 128$ | 0.7991 | 0.0213 | 0.0213 | 0.8006 | 0.7994 | 0.0215 | 0.9895 | 1.0000 | 1.0000 |
| $N = 256$ | 0.7995 | 0.0150 | 0.0150 | 0.7999 | 0.7991 | 0.0152 | −1.8261 | 1.0000 | 1.0000 |

$ASD_K$: Asymptotic standard deviation of $V_{rel}(\mathbf{R})$ based on Konishi's (1979) theory (eq. 39).
$ASD_{PF}$: Approximate standard deviation of $V_{rel}(\mathbf{R})$ based on Pan & Frank's (2004) approach (eqs. 36–38).

(*continued*)

**Table S3.** (*continued*)

| | E[$V_{rel}$(**R**)] | $ASD_K$ | $ASD_{PF}$ | Median | Mean | ESD | *T* | Pow. 5% | Pow. 1% |
|---|---|---|---|---|---|---|---|---|---|
| $p = 128, q = 2, V_{rel}(\mathbf{P}) = 0.1$ | | | | | | | | | |
| $N = 4$ | 0.3759 | 0.0654 | 0.0791 | 0.3662 | 0.3749 | 0.0366 | −1.8246 | 0.8902 | 0.8370 |
| $N = 8$ | 0.2121 | 0.0428 | 0.0467 | 0.2074 | 0.2125 | 0.0350 | 0.6874 | 0.9988 | 0.9984 |
| $N = 16$ | 0.1508 | 0.0292 | 0.0305 | 0.1483 | 0.1505 | 0.0264 | −0.7843 | 1.0000 | 1.0000 |
| $N = 32$ | 0.1242 | 0.0203 | 0.0208 | 0.1230 | 0.1238 | 0.0193 | −1.7367 | 1.0000 | 1.0000 |
| $N = 64$ | 0.1118 | 0.0143 | 0.0144 | 0.1117 | 0.1121 | 0.0140 | 1.3424 | 1.0000 | 1.0000 |
| $N = 128$ | 0.1059 | 0.0100 | 0.0101 | 0.1057 | 0.1060 | 0.0100 | 0.8314 | 1.0000 | 1.0000 |
| $N = 256$ | 0.1029 | 0.0071 | 0.0071 | 0.1027 | 0.1029 | 0.0070 | 0.2004 | 1.0000 | 1.0000 |
| $p = 128, q = 2, V_{rel}(\mathbf{P}) = 0.2$ | | | | | | | | | |
| $N = 4$ | 0.4247 | 0.0857 | 0.1254 | 0.4087 | 0.4234 | 0.0644 | −1.3764 | 0.9832 | 0.9696 |
| $N = 8$ | 0.2874 | 0.0561 | 0.0685 | 0.2826 | 0.2883 | 0.0544 | 1.0797 | 1.0000 | 1.0000 |
| $N = 16$ | 0.2390 | 0.0383 | 0.0425 | 0.2370 | 0.2386 | 0.0398 | −0.6199 | 1.0000 | 1.0000 |
| $N = 32$ | 0.2185 | 0.0267 | 0.0281 | 0.2179 | 0.2185 | 0.0267 | 0.0300 | 1.0000 | 1.0000 |
| $N = 64$ | 0.2090 | 0.0187 | 0.0192 | 0.2091 | 0.2089 | 0.0192 | −0.4189 | 1.0000 | 1.0000 |
| $N = 128$ | 0.2044 | 0.0132 | 0.0134 | 0.2044 | 0.2044 | 0.0134 | −0.1665 | 1.0000 | 1.0000 |
| $N = 256$ | 0.2022 | 0.0093 | 0.0094 | 0.2023 | 0.2022 | 0.0094 | −0.1209 | 1.0000 | 1.0000 |
| $p = 128, q = 2, V_{rel}(\mathbf{P}) = 0.4$ | | | | | | | | | |
| $N = 4$ | 0.5560 | 0.0476 | 0.1975 | 0.5211 | 0.5574 | 0.1170 | 0.8814 | 0.9994 | 0.9992 |
| $N = 8$ | 0.4636 | 0.0312 | 0.0879 | 0.4472 | 0.4638 | 0.0740 | 0.1420 | 1.0000 | 1.0000 |
| $N = 16$ | 0.4296 | 0.0213 | 0.0439 | 0.4233 | 0.4293 | 0.0415 | −0.4751 | 1.0000 | 1.0000 |
| $N = 32$ | 0.4143 | 0.0148 | 0.0237 | 0.4124 | 0.4144 | 0.0236 | 0.3334 | 1.0000 | 1.0000 |
| $N = 64$ | 0.4071 | 0.0104 | 0.0138 | 0.4066 | 0.4072 | 0.0137 | 0.5258 | 1.0000 | 1.0000 |
| $N = 128$ | 0.4035 | 0.0073 | 0.0086 | 0.4033 | 0.4034 | 0.0087 | −0.5183 | 1.0000 | 1.0000 |
| $N = 256$ | 0.4017 | 0.0052 | 0.0056 | 0.4018 | 0.4018 | 0.0056 | 0.8376 | 1.0000 | 1.0000 |
| $p = 128, q = 4, V_{rel}(\mathbf{P}) = 0.1$ | | | | | | | | | |
| $N = 4$ | 0.3791 | 0.0304 | 0.0681 | 0.3725 | 0.3793 | 0.0325 | 0.3717 | 0.9568 | 0.9226 |
| $N = 8$ | 0.2152 | 0.0199 | 0.0329 | 0.2116 | 0.2148 | 0.0250 | −1.2699 | 1.0000 | 1.0000 |
| $N = 16$ | 0.1529 | 0.0136 | 0.0183 | 0.1519 | 0.1531 | 0.0164 | 0.8764 | 1.0000 | 1.0000 |
| $N = 32$ | 0.1254 | 0.0095 | 0.0111 | 0.1249 | 0.1253 | 0.0107 | −0.4537 | 1.0000 | 1.0000 |
| $N = 64$ | 0.1125 | 0.0066 | 0.0072 | 0.1123 | 0.1124 | 0.0072 | −0.9179 | 1.0000 | 1.0000 |
| $N = 128$ | 0.1062 | 0.0047 | 0.0049 | 0.1062 | 0.1061 | 0.0048 | −0.4628 | 1.0000 | 1.0000 |
| $N = 256$ | 0.1031 | 0.0033 | 0.0034 | 0.1030 | 0.1030 | 0.0034 | −1.6580 | 1.0000 | 1.0000 |
| $p = 128, q = 4, V_{rel}(\mathbf{P}) = 0.2$ | | | | | | | | | |
| $N = 4$ | 0.4456 | 0.0158 | 0.1209 | 0.4326 | 0.4435 | 0.0656 | −2.3155 | 0.9932 | 0.9874 |
| $N = 8$ | 0.3039 | 0.0103 | 0.0524 | 0.2988 | 0.3042 | 0.0415 | 0.5443 | 1.0000 | 1.0000 |
| $N = 16$ | 0.2485 | 0.0071 | 0.0250 | 0.2455 | 0.2486 | 0.0225 | 0.4644 | 1.0000 | 1.0000 |
| $N = 32$ | 0.2235 | 0.0049 | 0.0126 | 0.2217 | 0.2234 | 0.0122 | −0.4670 | 1.0000 | 1.0000 |
| $N = 64$ | 0.2115 | 0.0034 | 0.0067 | 0.2108 | 0.2115 | 0.0066 | −1.0065 | 1.0000 | 1.0000 |
| $N = 128$ | 0.2057 | 0.0024 | 0.0037 | 0.2056 | 0.2058 | 0.0037 | 0.4702 | 1.0000 | 1.0000 |
| $N = 256$ | 0.2029 | 0.0017 | 0.0022 | 0.2028 | 0.2028 | 0.0022 | −0.1830 | 1.0000 | 1.0000 |

$ASD_K$: Asymptotic standard deviation of $V_{rel}$(**R**) based on Konishi's (1979) theory (eq. 39).
$ASD_{PF}$: Approximate standard deviation of $V_{rel}$(**R**) based on Pan & Frank's (2004) approach (eqs. 36–38).

(*continued*)

**Table S3.** (*continued*)

| | $E[V_{rel}(\mathbf{R})]$ | $ASD_K$ | $ASD_{PF}$ | Median | Mean | ESD | $T$ | Pow. 5% | Pow. 1% |
|---|---|---|---|---|---|---|---|---|---|
| $p = 256, q = 1, V_{rel}(\mathbf{P}) = 0.1$ | | | | | | | | | |
| $N = 4$ | 0.3745 | 0.1128 | 0.1070 | 0.3559 | 0.3741 | 0.0484 | −0.6148 | 0.8268 | 0.7730 |
| $N = 8$ | 0.2108 | 0.0739 | 0.0719 | 0.1966 | 0.2094 | 0.0510 | −1.8791 | 0.9940 | 0.9892 |
| $N = 16$ | 0.1499 | 0.0505 | 0.0498 | 0.1435 | 0.1495 | 0.0415 | −0.7639 | 1.0000 | 1.0000 |
| $N = 32$ | 0.1237 | 0.0351 | 0.0349 | 0.1196 | 0.1227 | 0.0318 | −2.2709 | 1.0000 | 1.0000 |
| $N = 64$ | 0.1115 | 0.0246 | 0.0245 | 0.1101 | 0.1116 | 0.0234 | 0.2559 | 1.0000 | 1.0000 |
| $N = 128$ | 0.1057 | 0.0173 | 0.0173 | 0.1051 | 0.1057 | 0.0169 | −0.0901 | 1.0000 | 1.0000 |
| $N = 256$ | 0.1028 | 0.0122 | 0.0122 | 0.1022 | 0.1028 | 0.0122 | −0.3173 | 1.0000 | 1.0000 |
| $p = 256, q = 1, V_{rel}(\mathbf{P}) = 0.2$ | | | | | | | | | |
| $N = 4$ | 0.4184 | 0.1818 | 0.1682 | 0.3913 | 0.4174 | 0.0825 | −0.8586 | 0.9208 | 0.8878 |
| $N = 8$ | 0.2814 | 0.1190 | 0.1149 | 0.2651 | 0.2802 | 0.0856 | −0.9368 | 0.9996 | 0.9994 |
| $N = 16$ | 0.2350 | 0.0813 | 0.0800 | 0.2284 | 0.2351 | 0.0702 | 0.1706 | 1.0000 | 1.0000 |
| $N = 32$ | 0.2162 | 0.0565 | 0.0561 | 0.2128 | 0.2155 | 0.0527 | −0.8742 | 1.0000 | 1.0000 |
| $N = 64$ | 0.2078 | 0.0397 | 0.0395 | 0.2074 | 0.2079 | 0.0392 | 0.2748 | 1.0000 | 1.0000 |
| $N = 128$ | 0.2038 | 0.0279 | 0.0279 | 0.2041 | 0.2044 | 0.0272 | 1.4626 | 1.0000 | 1.0000 |
| $N = 256$ | 0.2019 | 0.0197 | 0.0197 | 0.2015 | 0.2020 | 0.0195 | 0.5562 | 1.0000 | 1.0000 |
| $p = 256, q = 1, V_{rel}(\mathbf{P}) = 0.4$ | | | | | | | | | |
| $N = 4$ | 0.5158 | 0.2411 | 0.2202 | 0.4975 | 0.5148 | 0.1320 | −0.5622 | 0.9706 | 0.9582 |
| $N = 8$ | 0.4318 | 0.1578 | 0.1522 | 0.4348 | 0.4343 | 0.1278 | 1.3865 | 1.0000 | 1.0000 |
| $N = 16$ | 0.4111 | 0.1078 | 0.1061 | 0.4118 | 0.4091 | 0.0995 | −1.4526 | 1.0000 | 1.0000 |
| $N = 32$ | 0.4046 | 0.0750 | 0.0744 | 0.4078 | 0.4052 | 0.0726 | 0.5664 | 1.0000 | 1.0000 |
| $N = 64$ | 0.4021 | 0.0526 | 0.0524 | 0.4033 | 0.4024 | 0.0522 | 0.4500 | 1.0000 | 1.0000 |
| $N = 128$ | 0.4010 | 0.0371 | 0.0370 | 0.3999 | 0.4004 | 0.0366 | −1.0479 | 1.0000 | 1.0000 |
| $N = 256$ | 0.4005 | 0.0262 | 0.0261 | 0.4011 | 0.4007 | 0.0260 | 0.6241 | 1.0000 | 1.0000 |
| $p = 256, q = 1, V_{rel}(\mathbf{P}) = 0.6$ | | | | | | | | | |
| $N = 4$ | 0.6313 | 0.2215 | 0.2045 | 0.6451 | 0.6291 | 0.1586 | −0.9820 | 0.9896 | 0.9844 |
| $N = 8$ | 0.5973 | 0.1450 | 0.1408 | 0.6166 | 0.5992 | 0.1353 | 0.9907 | 1.0000 | 1.0000 |
| $N = 16$ | 0.5963 | 0.0991 | 0.0978 | 0.6059 | 0.5966 | 0.0983 | 0.2590 | 1.0000 | 1.0000 |
| $N = 32$ | 0.5978 | 0.0689 | 0.0685 | 0.6044 | 0.5988 | 0.0696 | 1.0898 | 1.0000 | 1.0000 |
| $N = 64$ | 0.5988 | 0.0483 | 0.0482 | 0.6006 | 0.5977 | 0.0487 | −1.5654 | 1.0000 | 1.0000 |
| $N = 128$ | 0.5994 | 0.0340 | 0.0340 | 0.6009 | 0.5993 | 0.0344 | −0.2590 | 1.0000 | 1.0000 |
| $N = 256$ | 0.5997 | 0.0240 | 0.0240 | 0.6003 | 0.5997 | 0.0241 | 0.0156 | 1.0000 | 1.0000 |
| $p = 256, q = 1, V_{rel}(\mathbf{P}) = 0.8$ | | | | | | | | | |
| $N = 4$ | 0.7768 | 0.1383 | 0.1312 | 0.8208 | 0.7728 | 0.1551 | −1.8239 | 0.9960 | 0.9946 |
| $N = 8$ | 0.7831 | 0.0905 | 0.0890 | 0.8098 | 0.7857 | 0.1056 | 1.7052 | 1.0000 | 1.0000 |
| $N = 16$ | 0.7918 | 0.0618 | 0.0614 | 0.8031 | 0.7915 | 0.0681 | −0.3408 | 1.0000 | 1.0000 |
| $N = 32$ | 0.7961 | 0.0430 | 0.0429 | 0.8022 | 0.7963 | 0.0460 | 0.3738 | 1.0000 | 1.0000 |
| $N = 64$ | 0.7981 | 0.0302 | 0.0301 | 0.8004 | 0.7977 | 0.0305 | −0.9565 | 1.0000 | 1.0000 |
| $N = 128$ | 0.7991 | 0.0212 | 0.0212 | 0.8007 | 0.7993 | 0.0212 | 0.6642 | 1.0000 | 1.0000 |
| $N = 256$ | 0.7995 | 0.0150 | 0.0150 | 0.8002 | 0.7996 | 0.0153 | 0.4570 | 1.0000 | 1.0000 |

$ASD_K$: Asymptotic standard deviation of $V_{rel}(\mathbf{R})$ based on Konishi's (1979) theory (eq. 39).
$ASD_{PF}$: Approximate standard deviation of $V_{rel}(\mathbf{R})$ based on Pan & Frank's (2004) approach (eqs. 36–38).

(*continued*)

**Table S3.** (*continued*)

| | $E[V_{rel}(\mathbf{R})]$ | $ASD_K$ | $ASD_{PF}$ | Median | Mean | ESD | $T$ | Pow. 5% | Pow. 1% |
|---|---|---|---|---|---|---|---|---|---|
| $p = 256, q = 2, V_{rel}(\mathbf{P}) = 0.1$ | | | | | | | | | |
| $N = 4$ | 0.3759 | 0.0646 | 0.0773 | 0.3667 | 0.3751 | 0.0357 | −1.4520 | 0.9512 | 0.9198 |
| $N = 8$ | 0.2121 | 0.0423 | 0.0459 | 0.2078 | 0.2126 | 0.0340 | 0.9840 | 1.0000 | 1.0000 |
| $N = 16$ | 0.1508 | 0.0289 | 0.0301 | 0.1483 | 0.1500 | 0.0263 | −2.1534 | 1.0000 | 1.0000 |
| $N = 32$ | 0.1242 | 0.0201 | 0.0205 | 0.1229 | 0.1239 | 0.0189 | −1.1190 | 1.0000 | 1.0000 |
| $N = 64$ | 0.1118 | 0.0141 | 0.0142 | 0.1113 | 0.1119 | 0.0139 | 0.1824 | 1.0000 | 1.0000 |
| $N = 128$ | 0.1058 | 0.0099 | 0.0100 | 0.1055 | 0.1057 | 0.0099 | −0.9821 | 1.0000 | 1.0000 |
| $N = 256$ | 0.1029 | 0.0070 | 0.0070 | 0.1030 | 0.1031 | 0.0071 | 1.5009 | 1.0000 | 1.0000 |
| $p = 256, q = 2, V_{rel}(\mathbf{P}) = 0.2$ | | | | | | | | | |
| $N = 4$ | 0.4246 | 0.0853 | 0.1238 | 0.4108 | 0.4236 | 0.0622 | −1.1437 | 0.9914 | 0.9872 |
| $N = 8$ | 0.2874 | 0.0558 | 0.0678 | 0.2829 | 0.2871 | 0.0534 | −0.3519 | 1.0000 | 1.0000 |
| $N = 16$ | 0.2389 | 0.0382 | 0.0422 | 0.2382 | 0.2388 | 0.0385 | −0.2851 | 1.0000 | 1.0000 |
| $N = 32$ | 0.2184 | 0.0265 | 0.0279 | 0.2181 | 0.2180 | 0.0270 | −1.1649 | 1.0000 | 1.0000 |
| $N = 64$ | 0.2090 | 0.0186 | 0.0191 | 0.2090 | 0.2088 | 0.0189 | −0.6108 | 1.0000 | 1.0000 |
| $N = 128$ | 0.2044 | 0.0131 | 0.0133 | 0.2045 | 0.2046 | 0.0131 | 1.0852 | 1.0000 | 1.0000 |
| $N = 256$ | 0.2022 | 0.0093 | 0.0093 | 0.2023 | 0.2021 | 0.0092 | −0.5046 | 1.0000 | 1.0000 |
| $p = 256, q = 2, V_{rel}(\mathbf{P}) = 0.4$ | | | | | | | | | |
| $N = 4$ | 0.5555 | 0.0482 | 0.1958 | 0.5217 | 0.5576 | 0.1141 | 1.2655 | 0.9996 | 0.9996 |
| $N = 8$ | 0.4633 | 0.0315 | 0.0873 | 0.4466 | 0.4616 | 0.0721 | −1.6268 | 1.0000 | 1.0000 |
| $N = 16$ | 0.4294 | 0.0215 | 0.0437 | 0.4244 | 0.4307 | 0.0414 | 2.1603 | 1.0000 | 1.0000 |
| $N = 32$ | 0.4142 | 0.0150 | 0.0237 | 0.4130 | 0.4146 | 0.0231 | 1.2190 | 1.0000 | 1.0000 |
| $N = 64$ | 0.4070 | 0.0105 | 0.0139 | 0.4064 | 0.4069 | 0.0137 | −0.8165 | 1.0000 | 1.0000 |
| $N = 128$ | 0.4035 | 0.0074 | 0.0087 | 0.4034 | 0.4034 | 0.0088 | −1.0027 | 1.0000 | 1.0000 |
| $N = 256$ | 0.4017 | 0.0052 | 0.0057 | 0.4019 | 0.4018 | 0.0058 | 0.4936 | 1.0000 | 1.0000 |
| $p = 256, q = 4, V_{rel}(\mathbf{P}) = 0.1$ | | | | | | | | | |
| $N = 4$ | 0.3790 | 0.0302 | 0.0660 | 0.3724 | 0.3791 | 0.0312 | 0.1148 | 0.9894 | 0.9778 |
| $N = 8$ | 0.2152 | 0.0198 | 0.0320 | 0.2118 | 0.2148 | 0.0236 | −1.1456 | 1.0000 | 1.0000 |
| $N = 16$ | 0.1528 | 0.0135 | 0.0179 | 0.1515 | 0.1525 | 0.0157 | −1.3028 | 1.0000 | 1.0000 |
| $N = 32$ | 0.1254 | 0.0094 | 0.0110 | 0.1246 | 0.1252 | 0.0105 | −0.9418 | 1.0000 | 1.0000 |
| $N = 64$ | 0.1124 | 0.0066 | 0.0072 | 0.1125 | 0.1125 | 0.0071 | 1.0205 | 1.0000 | 1.0000 |
| $N = 128$ | 0.1062 | 0.0046 | 0.0048 | 0.1062 | 0.1062 | 0.0048 | −0.0593 | 1.0000 | 1.0000 |
| $N = 256$ | 0.1031 | 0.0033 | 0.0033 | 0.1031 | 0.1031 | 0.0033 | 1.2325 | 1.0000 | 1.0000 |
| $p = 256, q = 4, V_{rel}(\mathbf{P}) = 0.2$ | | | | | | | | | |
| $N = 4$ | 0.4449 | 0.0165 | 0.1188 | 0.4348 | 0.4454 | 0.0668 | 0.5683 | 0.9978 | 0.9972 |
| $N = 8$ | 0.3034 | 0.0108 | 0.0516 | 0.2979 | 0.3032 | 0.0401 | −0.3479 | 1.0000 | 1.0000 |
| $N = 16$ | 0.2482 | 0.0074 | 0.0247 | 0.2452 | 0.2483 | 0.0223 | 0.2351 | 1.0000 | 1.0000 |
| $N = 32$ | 0.2233 | 0.0051 | 0.0125 | 0.2217 | 0.2234 | 0.0122 | 0.4341 | 1.0000 | 1.0000 |
| $N = 64$ | 0.2115 | 0.0036 | 0.0067 | 0.2108 | 0.2115 | 0.0065 | 0.0211 | 1.0000 | 1.0000 |
| $N = 128$ | 0.2057 | 0.0025 | 0.0038 | 0.2055 | 0.2057 | 0.0039 | 0.8353 | 1.0000 | 1.0000 |
| $N = 256$ | 0.2028 | 0.0018 | 0.0023 | 0.2028 | 0.2028 | 0.0022 | 0.1085 | 1.0000 | 1.0000 |

$ASD_K$: Asymptotic standard deviation of $V_{rel}(\mathbf{R})$ based on Konishi's (1979) theory (eq. 39).
$ASD_{PF}$: Approximate standard deviation of $V_{rel}(\mathbf{R})$ based on Pan & Frank's (2004) approach (eqs. 36–38).

(*continued*)

**Table S3.** (*continued*)

| | $E[V_{rel}(\mathbf{R})]$ | $ASD_K$ | $ASD_{PF}$ | Median | Mean | ESD | $T$ | Pow. 5% | Pow. 1% |
|---|---|---|---|---|---|---|---|---|---|
| $p = 1024$, $q = 1$, $V_{rel}(\mathbf{P}) = 0.1$ | | | | | | | | | |
| $N = 4$ | 0.3745 | 0.1119 | 0.1058 | 0.3575 | 0.3740 | 0.0464 | −0.8598 | 0.9402 | 0.9184 |
| $N = 8$ | 0.2108 | 0.0733 | 0.0712 | 0.1994 | 0.2114 | 0.0517 | 0.8268 | 0.9992 | 0.9988 |
| $N = 16$ | 0.1499 | 0.0501 | 0.0494 | 0.1440 | 0.1498 | 0.0421 | −0.1942 | 1.0000 | 1.0000 |
| $N = 32$ | 0.1237 | 0.0348 | 0.0346 | 0.1195 | 0.1232 | 0.0317 | −1.1238 | 1.0000 | 1.0000 |
| $N = 64$ | 0.1115 | 0.0244 | 0.0243 | 0.1100 | 0.1116 | 0.0235 | 0.0669 | 1.0000 | 1.0000 |
| $N = 128$ | 0.1057 | 0.0172 | 0.0172 | 0.1046 | 0.1053 | 0.0167 | −1.4923 | 1.0000 | 1.0000 |
| $N = 256$ | 0.1028 | 0.0121 | 0.0121 | 0.1022 | 0.1027 | 0.0120 | −1.0207 | 1.0000 | 1.0000 |
| $p = 1024$, $q = 1$, $V_{rel}(\mathbf{P}) = 0.2$ | | | | | | | | | |
| $N = 4$ | 0.4184 | 0.1808 | 0.1671 | 0.3901 | 0.4175 | 0.0832 | −0.7299 | 0.9720 | 0.9636 |
| $N = 8$ | 0.2814 | 0.1184 | 0.1143 | 0.2681 | 0.2808 | 0.0852 | −0.4595 | 1.0000 | 1.0000 |
| $N = 16$ | 0.2350 | 0.0809 | 0.0795 | 0.2314 | 0.2361 | 0.0695 | 1.1295 | 1.0000 | 1.0000 |
| $N = 32$ | 0.2162 | 0.0563 | 0.0558 | 0.2130 | 0.2156 | 0.0525 | −0.8736 | 1.0000 | 1.0000 |
| $N = 64$ | 0.2078 | 0.0395 | 0.0393 | 0.2072 | 0.2083 | 0.0384 | 0.9543 | 1.0000 | 1.0000 |
| $N = 128$ | 0.2038 | 0.0278 | 0.0277 | 0.2026 | 0.2030 | 0.0273 | −2.2022 | 1.0000 | 1.0000 |
| $N = 256$ | 0.2019 | 0.0196 | 0.0196 | 0.2010 | 0.2013 | 0.0193 | −2.1799 | 1.0000 | 1.0000 |
| $p = 1024$, $q = 1$, $V_{rel}(\mathbf{P}) = 0.4$ | | | | | | | | | |
| $N = 4$ | 0.5158 | 0.2403 | 0.2195 | 0.4926 | 0.5118 | 0.1306 | −2.1637 | 0.9890 | 0.9848 |
| $N = 8$ | 0.4318 | 0.1573 | 0.1517 | 0.4266 | 0.4286 | 0.1287 | −1.7550 | 1.0000 | 1.0000 |
| $N = 16$ | 0.4111 | 0.1075 | 0.1057 | 0.4127 | 0.4111 | 0.1008 | −0.0454 | 1.0000 | 1.0000 |
| $N = 32$ | 0.4046 | 0.0748 | 0.0742 | 0.4072 | 0.4055 | 0.0726 | 0.8769 | 1.0000 | 1.0000 |
| $N = 64$ | 0.4021 | 0.0524 | 0.0522 | 0.4035 | 0.4024 | 0.0524 | 0.4578 | 1.0000 | 1.0000 |
| $N = 128$ | 0.4010 | 0.0369 | 0.0369 | 0.4006 | 0.4007 | 0.0367 | −0.5419 | 1.0000 | 1.0000 |
| $N = 256$ | 0.4005 | 0.0261 | 0.0260 | 0.4015 | 0.4007 | 0.0263 | 0.6336 | 1.0000 | 1.0000 |
| $p = 1024$, $q = 1$, $V_{rel}(\mathbf{P}) = 0.6$ | | | | | | | | | |
| $N = 4$ | 0.6313 | 0.2210 | 0.2040 | 0.6412 | 0.6285 | 0.1592 | −1.2475 | 0.9946 | 0.9932 |
| $N = 8$ | 0.5973 | 0.1447 | 0.1405 | 0.6157 | 0.5975 | 0.1355 | 0.1026 | 1.0000 | 1.0000 |
| $N = 16$ | 0.5963 | 0.0988 | 0.0976 | 0.6088 | 0.5965 | 0.0979 | 0.1669 | 1.0000 | 1.0000 |
| $N = 32$ | 0.5978 | 0.0688 | 0.0684 | 0.6035 | 0.5971 | 0.0703 | −0.6285 | 1.0000 | 1.0000 |
| $N = 64$ | 0.5988 | 0.0482 | 0.0481 | 0.6019 | 0.5985 | 0.0479 | −0.4679 | 1.0000 | 1.0000 |
| $N = 128$ | 0.5994 | 0.0340 | 0.0339 | 0.6006 | 0.5996 | 0.0335 | 0.3479 | 1.0000 | 1.0000 |
| $N = 256$ | 0.5997 | 0.0240 | 0.0240 | 0.6002 | 0.5995 | 0.0240 | −0.4863 | 1.0000 | 1.0000 |
| $p = 1024$, $q = 1$, $V_{rel}(\mathbf{P}) = 0.8$ | | | | | | | | | |
| $N = 4$ | 0.7768 | 0.1380 | 0.1309 | 0.8236 | 0.7733 | 0.1543 | −1.5756 | 0.9986 | 0.9980 |
| $N = 8$ | 0.7831 | 0.0903 | 0.0888 | 0.8088 | 0.7830 | 0.1074 | −0.0831 | 1.0000 | 1.0000 |
| $N = 16$ | 0.7918 | 0.0617 | 0.0613 | 0.8057 | 0.7930 | 0.0683 | 1.2443 | 1.0000 | 1.0000 |
| $N = 32$ | 0.7961 | 0.0429 | 0.0428 | 0.8014 | 0.7954 | 0.0454 | −0.9758 | 1.0000 | 1.0000 |
| $N = 64$ | 0.7981 | 0.0301 | 0.0301 | 0.8006 | 0.7981 | 0.0308 | 0.0710 | 1.0000 | 1.0000 |
| $N = 128$ | 0.7991 | 0.0212 | 0.0212 | 0.8006 | 0.7988 | 0.0220 | −0.7670 | 1.0000 | 1.0000 |
| $N = 256$ | 0.7995 | 0.0150 | 0.0150 | 0.7994 | 0.7989 | 0.0154 | −2.7764 | 1.0000 | 1.0000 |

$ASD_K$: Asymptotic standard deviation of $V_{rel}(\mathbf{R})$ based on Konishi's (1979) theory (eq. 39).
$ASD_{PF}$: Approximate standard deviation of $V_{rel}(\mathbf{R})$ based on Pan & Frank's (2004) approach (eqs. 36–38).

(*continued*)

**Table S3.** (*continued*)

| | E[$V_{rel}$(**R**)] | $ASD_K$ | $ASD_{PF}$ | Median | Mean | ESD | $T$ | Pow. 5% | Pow. 1% |
|---|---|---|---|---|---|---|---|---|---|
| $p$ = 1024, $q$ = 2, $V_{rel}$(**P**) = 0.1 | | | | | | | | | |
| $N$ = 4 | 0.3759 | 0.0640 | 0.0760 | 0.3678 | 0.3762 | 0.0350 | 0.7678 | 0.9934 | 0.9886 |
| $N$ = 8 | 0.2121 | 0.0419 | 0.0453 | 0.2076 | 0.2116 | 0.0327 | −1.0455 | 1.0000 | 1.0000 |
| $N$ = 16 | 0.1508 | 0.0286 | 0.0297 | 0.1485 | 0.1505 | 0.0259 | −0.8294 | 1.0000 | 1.0000 |
| $N$ = 32 | 0.1242 | 0.0199 | 0.0203 | 0.1231 | 0.1241 | 0.0192 | −0.5213 | 1.0000 | 1.0000 |
| $N$ = 64 | 0.1118 | 0.0140 | 0.0141 | 0.1119 | 0.1121 | 0.0138 | 1.5748 | 1.0000 | 1.0000 |
| $N$ = 128 | 0.1058 | 0.0098 | 0.0099 | 0.1058 | 0.1059 | 0.0096 | 0.5372 | 1.0000 | 1.0000 |
| $N$ = 256 | 0.1029 | 0.0069 | 0.0070 | 0.1027 | 0.1029 | 0.0069 | 0.0563 | 1.0000 | 1.0000 |
| $p$ = 1024, $q$ = 2, $V_{rel}$(**P**) = 0.2 | | | | | | | | | |
| $N$ = 4 | 0.4246 | 0.0850 | 0.1225 | 0.4104 | 0.4236 | 0.0623 | −1.1057 | 0.9990 | 0.9986 |
| $N$ = 8 | 0.2873 | 0.0556 | 0.0673 | 0.2833 | 0.2877 | 0.0534 | 0.4989 | 1.0000 | 1.0000 |
| $N$ = 16 | 0.2389 | 0.0380 | 0.0419 | 0.2370 | 0.2384 | 0.0382 | −0.9306 | 1.0000 | 1.0000 |
| $N$ = 32 | 0.2184 | 0.0264 | 0.0278 | 0.2186 | 0.2186 | 0.0271 | 0.4335 | 1.0000 | 1.0000 |
| $N$ = 64 | 0.2090 | 0.0185 | 0.0190 | 0.2086 | 0.2087 | 0.0186 | −1.0366 | 1.0000 | 1.0000 |
| $N$ = 128 | 0.2044 | 0.0131 | 0.0132 | 0.2048 | 0.2044 | 0.0130 | −0.1240 | 1.0000 | 1.0000 |
| $N$ = 256 | 0.2022 | 0.0092 | 0.0093 | 0.2025 | 0.2023 | 0.0092 | 0.9339 | 1.0000 | 1.0000 |
| $p$ = 1024, $q$ = 2, $V_{rel}$(**P**) = 0.4 | | | | | | | | | |
| $N$ = 4 | 0.5552 | 0.0486 | 0.1946 | 0.5181 | 0.5524 | 0.1116 | −1.7454 | 1.0000 | 1.0000 |
| $N$ = 8 | 0.4631 | 0.0318 | 0.0868 | 0.4460 | 0.4617 | 0.0722 | −1.3874 | 1.0000 | 1.0000 |
| $N$ = 16 | 0.4293 | 0.0217 | 0.0435 | 0.4240 | 0.4295 | 0.0407 | 0.3579 | 1.0000 | 1.0000 |
| $N$ = 32 | 0.4142 | 0.0151 | 0.0237 | 0.4120 | 0.4141 | 0.0240 | −0.1373 | 1.0000 | 1.0000 |
| $N$ = 64 | 0.4070 | 0.0106 | 0.0139 | 0.4064 | 0.4070 | 0.0137 | 0.2280 | 1.0000 | 1.0000 |
| $N$ = 128 | 0.4035 | 0.0075 | 0.0087 | 0.4038 | 0.4037 | 0.0087 | 2.2263 | 1.0000 | 1.0000 |
| $N$ = 256 | 0.4017 | 0.0053 | 0.0057 | 0.4017 | 0.4017 | 0.0057 | 0.1438 | 1.0000 | 1.0000 |
| $p$ = 1024, $q$ = 4, $V_{rel}$(**P**) = 0.1 | | | | | | | | | |
| $N$ = 4 | 0.3790 | 0.0301 | 0.0644 | 0.3724 | 0.3792 | 0.0307 | 0.4532 | 0.9996 | 0.9994 |
| $N$ = 8 | 0.2151 | 0.0197 | 0.0314 | 0.2122 | 0.2155 | 0.0237 | 1.1716 | 1.0000 | 1.0000 |
| $N$ = 16 | 0.1528 | 0.0134 | 0.0176 | 0.1515 | 0.1528 | 0.0159 | −0.0284 | 1.0000 | 1.0000 |
| $N$ = 32 | 0.1253 | 0.0094 | 0.0109 | 0.1248 | 0.1251 | 0.0106 | −1.6034 | 1.0000 | 1.0000 |
| $N$ = 64 | 0.1124 | 0.0066 | 0.0071 | 0.1123 | 0.1123 | 0.0070 | −1.7393 | 1.0000 | 1.0000 |
| $N$ = 128 | 0.1062 | 0.0046 | 0.0048 | 0.1061 | 0.1062 | 0.0048 | 0.4660 | 1.0000 | 1.0000 |
| $N$ = 256 | 0.1031 | 0.0033 | 0.0033 | 0.1031 | 0.1031 | 0.0033 | 0.1723 | 1.0000 | 1.0000 |
| $p$ = 1024, $q$ = 4, $V_{rel}$(**P**) = 0.2 | | | | | | | | | |
| $N$ = 4 | 0.4444 | 0.0171 | 0.1172 | 0.4343 | 0.4457 | 0.0657 | 1.4518 | 1.0000 | 1.0000 |
| $N$ = 8 | 0.3030 | 0.0112 | 0.0509 | 0.2979 | 0.3037 | 0.0405 | 1.2073 | 1.0000 | 1.0000 |
| $N$ = 16 | 0.2480 | 0.0076 | 0.0244 | 0.2455 | 0.2486 | 0.0220 | 1.7390 | 1.0000 | 1.0000 |
| $N$ = 32 | 0.2232 | 0.0053 | 0.0124 | 0.2220 | 0.2234 | 0.0120 | 0.8155 | 1.0000 | 1.0000 |
| $N$ = 64 | 0.2114 | 0.0037 | 0.0067 | 0.2109 | 0.2114 | 0.0065 | −0.6746 | 1.0000 | 1.0000 |
| $N$ = 128 | 0.2057 | 0.0026 | 0.0038 | 0.2054 | 0.2056 | 0.0038 | −1.4728 | 1.0000 | 1.0000 |
| $N$ = 256 | 0.2028 | 0.0019 | 0.0023 | 0.2028 | 0.2028 | 0.0023 | −0.1714 | 1.0000 | 1.0000 |

$ASD_K$: Asymptotic standard deviation of $V_{rel}$(**R**) based on Konishi's (1979) theory (eq. 39).
$ASD_{PF}$: Approximate standard deviation of $V_{rel}$(**R**) based on Pan & Frank's (2004) approach (eqs. 36–38).

(*continued*)

**Table S3.** (*continued*)

| | E[$V_{rel}$(**R**)] | ASD$_K$ | ASD$_{PF}$ | Median | Mean | ESD | $T$ | Pow. 5% | Pow. 1% |
|---|---|---|---|---|---|---|---|---|---|
| $p$ = 2, linearly decreasing λ ($V_{rel}$(**P**) = 0.1111) | | | | | | | | | |
| $N$ = 4 | 0.3793 | 0.3421 | 0.3103* | 0.3202 | 0.3782 | 0.3087 | −0.2446 | 0.0640 | 0.0134 |
| $N$ = 8 | 0.2185 | 0.2240 | 0.2096* | 0.1575 | 0.2187 | 0.2095 | 0.0654 | 0.1256 | 0.0354 |
| $N$ = 16 | 0.1593 | 0.1530 | 0.1461* | 0.1207 | 0.1601 | 0.1476 | 0.4067 | 0.2536 | 0.0924 |
| $N$ = 32 | 0.1339 | 0.1064 | 0.1035* | 0.1163 | 0.1362 | 0.1048 | 1.5246 | 0.4890 | 0.2662 |
| $N$ = 64 | 0.1222 | 0.0747 | 0.0736* | 0.1139 | 0.1227 | 0.0740 | 0.4641 | 0.7850 | 0.6022 |
| $N$ = 128 | 0.1166 | 0.0526 | 0.0522* | 0.1128 | 0.1173 | 0.0534 | 0.8838 | 0.9780 | 0.9248 |
| $N$ = 256 | 0.1138 | 0.0371 | 0.0370* | 0.1115 | 0.1136 | 0.0369 | −0.3938 | 0.9998 | 0.9986 |
| $p$ = 4, linearly decreasing λ ($V_{rel}$(**P**) = 0.0667) | | | | | | | | | |
| $N$ = 4 | 0.3607 | 0.1023 | 0.1479 | 0.3405 | 0.3602 | 0.1328 | −0.2992 | 0.0808 | 0.0214 |
| $N$ = 8 | 0.1881 | 0.0670 | 0.0836 | 0.1769 | 0.1876 | 0.0810 | −0.4529 | 0.1366 | 0.0428 |
| $N$ = 16 | 0.1221 | 0.0458 | 0.0514 | 0.1145 | 0.1217 | 0.0511 | −0.5361 | 0.3872 | 0.1528 |
| $N$ = 32 | 0.0932 | 0.0318 | 0.0337 | 0.0878 | 0.0925 | 0.0344 | −1.5091 | 0.8004 | 0.5156 |
| $N$ = 64 | 0.0796 | 0.0223 | 0.0230 | 0.0777 | 0.0801 | 0.0229 | 1.2955 | 0.9956 | 0.9664 |
| $N$ = 128 | 0.0731 | 0.0157 | 0.0160 | 0.0717 | 0.0729 | 0.0160 | −0.6557 | 1.0000 | 1.0000 |
| $N$ = 256 | 0.0699 | 0.0111 | 0.0112 | 0.0696 | 0.0702 | 0.0111 | 1.9461 | 1.0000 | 1.0000 |
| $p$ = 8, linearly decreasing λ ($V_{rel}$(**P**) = 0.0370) | | | | | | | | | |
| $N$ = 4 | 0.3485 | 0.0338 | 0.0800 | 0.3374 | 0.3479 | 0.0631 | −0.7111 | 0.0804 | 0.0214 |
| $N$ = 8 | 0.1679 | 0.0221 | 0.0409 | 0.1630 | 0.1672 | 0.0378 | −1.4542 | 0.1808 | 0.0612 |
| $N$ = 16 | 0.0974 | 0.0151 | 0.0225 | 0.0946 | 0.0971 | 0.0219 | −1.1963 | 0.4982 | 0.2516 |
| $N$ = 32 | 0.0661 | 0.0105 | 0.0133 | 0.0650 | 0.0664 | 0.0136 | 1.4437 | 0.9468 | 0.8070 |
| $N$ = 64 | 0.0513 | 0.0074 | 0.0084 | 0.0506 | 0.0513 | 0.0084 | 0.3549 | 1.0000 | 0.9998 |
| $N$ = 128 | 0.0441 | 0.0052 | 0.0056 | 0.0437 | 0.0440 | 0.0055 | −1.0232 | 1.0000 | 1.0000 |
| $N$ = 256 | 0.0405 | 0.0037 | 0.0038 | 0.0404 | 0.0406 | 0.0038 | 1.1287 | 1.0000 | 1.0000 |
| $p$ = 16, linearly decreasing λ ($V_{rel}$(**P**) = 0.0196) | | | | | | | | | |
| $N$ = 4 | 0.3415 | 0.0115 | 0.0427 | 0.3360 | 0.3417 | 0.0318 | 0.4905 | 0.0956 | 0.0280 |
| $N$ = 8 | 0.1563 | 0.0075 | 0.0210 | 0.1542 | 0.1564 | 0.0185 | 0.4621 | 0.2146 | 0.0758 |
| $N$ = 16 | 0.0831 | 0.0052 | 0.0109 | 0.0823 | 0.0832 | 0.0103 | 0.9491 | 0.5530 | 0.2952 |
| $N$ = 32 | 0.0502 | 0.0036 | 0.0060 | 0.0498 | 0.0503 | 0.0060 | 0.5576 | 0.9854 | 0.9070 |
| $N$ = 64 | 0.0347 | 0.0025 | 0.0035 | 0.0345 | 0.0346 | 0.0035 | −0.3910 | 1.0000 | 1.0000 |
| $N$ = 128 | 0.0271 | 0.0018 | 0.0021 | 0.0270 | 0.0271 | 0.0021 | −0.2154 | 1.0000 | 1.0000 |
| $N$ = 256 | 0.0233 | 0.0012 | 0.0014 | 0.0233 | 0.0233 | 0.0014 | −0.0289 | 1.0000 | 1.0000 |
| $p$ = 32, linearly decreasing λ ($V_{rel}$(**P**) = 0.0101) | | | | | | | | | |
| $N$ = 4 | 0.3375 | 0.0040 | 0.0219 | 0.3343 | 0.3376 | 0.0157 | 0.3965 | 0.0998 | 0.0242 |
| $N$ = 8 | 0.1497 | 0.0026 | 0.0106 | 0.1487 | 0.1496 | 0.0091 | −1.2013 | 0.2206 | 0.0844 |
| $N$ = 16 | 0.0751 | 0.0018 | 0.0053 | 0.0747 | 0.0750 | 0.0050 | −0.9852 | 0.6218 | 0.3566 |
| $N$ = 32 | 0.0415 | 0.0012 | 0.0028 | 0.0414 | 0.0415 | 0.0027 | −0.4252 | 0.9896 | 0.9576 |
| $N$ = 64 | 0.0255 | 0.0009 | 0.0015 | 0.0255 | 0.0255 | 0.0015 | 0.1871 | 1.0000 | 1.0000 |
| $N$ = 128 | 0.0178 | 0.0006 | 0.0009 | 0.0177 | 0.0178 | 0.0009 | 0.9912 | 1.0000 | 1.0000 |
| $N$ = 256 | 0.0139 | 0.0004 | 0.0005 | 0.0139 | 0.0139 | 0.0005 | −0.3046 | 1.0000 | 1.0000 |

ASD$_K$: Asymptotic standard deviation of $V_{rel}$(**R**) based on Konishi's (1979) theory (eq. 39).
ASD$_{PF}$: Approximate standard deviation of $V_{rel}$(**R**) based on Pan & Frank's (2004) approach (eqs. 36–38).
*: Exact standard deviations for $p$ = 2 are shown in these cells.

(*continued*)

**Table S3.** (*continued*)

| | $E[V_{rel}(\mathbf{R})]$ | $ASD_K$ | $ASD_{PF}$ | Median | Mean | ESD | $T$ | Pow. 5% | Pow. 1% |
|---|---|---|---|---|---|---|---|---|---|
| $p = 64$, linearly decreasing λ ($V_{rel}(\mathbf{P}) = 0.0051$) | | | | | | | | | |
| $N = 4$ | 0.3355 | 0.00140 | 0.0111 | 0.3341 | 0.3356 | 0.00789 | 0.7901 | 0.0940 | 0.0272 |
| $N = 8$ | 0.1464 | 0.00092 | 0.0053 | 0.1459 | 0.1464 | 0.00466 | −0.4444 | 0.2506 | 0.0858 |
| $N = 16$ | 0.0710 | 0.00063 | 0.0027 | 0.0708 | 0.0709 | 0.00247 | −1.4701 | 0.6356 | 0.3802 |
| $N = 32$ | 0.0370 | 0.00044 | 0.0014 | 0.0369 | 0.0370 | 0.00132 | 0.6883 | 0.9942 | 0.9640 |
| $N = 64$ | 0.0208 | 0.00031 | 0.0007 | 0.0208 | 0.0208 | 0.00070 | 1.0001 | 1.0000 | 1.0000 |
| $N = 128$ | 0.0129 | 0.00022 | 0.0004 | 0.0129 | 0.0129 | 0.00038 | −1.1839 | 1.0000 | 1.0000 |
| $N = 256$ | 0.0090 | 0.00015 | 0.0002 | 0.0090 | 0.0090 | 0.00022 | −2.0294 | 1.0000 | 1.0000 |
| $p = 128$, linearly decreasing λ ($V_{rel}(\mathbf{P}) = 0.0026$) | | | | | | | | | |
| $N = 4$ | 0.3344 | 0.00049 | 0.0056 | 0.3335 | 0.3343 | 0.00396 | −1.2815 | 0.1000 | 0.0318 |
| $N = 8$ | 0.1446 | 0.00032 | 0.0027 | 0.1445 | 0.1447 | 0.00231 | 0.6599 | 0.2520 | 0.0992 |
| $N = 16$ | 0.0688 | 0.00022 | 0.0013 | 0.0688 | 0.0688 | 0.00124 | −1.3645 | 0.6396 | 0.3790 |
| $N = 32$ | 0.0346 | 0.00015 | 0.0007 | 0.0346 | 0.0346 | 0.00065 | −0.4754 | 0.9924 | 0.9744 |
| $N = 64$ | 0.0183 | 0.00011 | 0.0003 | 0.0183 | 0.0183 | 0.00033 | −0.2605 | 1.0000 | 1.0000 |
| $N = 128$ | 0.0104 | 0.00008 | 0.0002 | 0.0104 | 0.0104 | 0.00018 | 0.5676 | 1.0000 | 1.0000 |
| $N = 256$ | 0.0065 | 0.00005 | 0.0001 | 0.0065 | 0.0065 | 0.00010 | 2.0255 | 1.0000 | 1.0000 |
| $p = 256$, linearly decreasing λ ($V_{rel}(\mathbf{P}) = 0.0013$) | | | | | | | | | |
| $N = 4$ | 0.3339 | 0.00017 | 0.00282 | 0.3334 | 0.3338 | 0.00197 | −1.4148 | 0.0930 | 0.0230 |
| $N = 8$ | 0.1438 | 0.00011 | 0.00134 | 0.1437 | 0.1438 | 0.00117 | 0.7633 | 0.2498 | 0.0956 |
| $N = 16$ | 0.0678 | 0.00008 | 0.00066 | 0.0677 | 0.0678 | 0.00062 | 2.7597 | 0.6702 | 0.4300 |
| $N = 32$ | 0.0334 | 0.00005 | 0.00033 | 0.0334 | 0.0334 | 0.00032 | −0.4550 | 0.9962 | 0.9750 |
| $N = 64$ | 0.0171 | 0.00004 | 0.00017 | 0.0171 | 0.0171 | 0.00016 | 0.6150 | 1.0000 | 1.0000 |
| $N = 128$ | 0.0091 | 0.00003 | 0.00009 | 0.0091 | 0.0091 | 0.00009 | 0.2258 | 1.0000 | 1.0000 |
| $N = 256$ | 0.0052 | 0.00002 | 0.00004 | 0.0052 | 0.0052 | 0.00004 | −0.0664 | 1.0000 | 1.0000 |
| $p = 1024$, linearly decreasing λ ($V_{rel}(\mathbf{P}) = 0.0003$) | | | | | | | | | |
| $N = 4$ | 0.3335 | 0.00002 | 0.00071 | 0.3334 | 0.3335 | 0.00049 | 0.6559 | 0.0972 | 0.0288 |
| $N = 8$ | 0.1431 | 0.00001 | 0.00034 | 0.1431 | 0.1431 | 0.00029 | −0.0088 | 0.2516 | 0.0904 |
| $N = 16$ | 0.0669 | 0.00001 | 0.00016 | 0.0669 | 0.0669 | 0.00016 | −0.5107 | 0.6618 | 0.4206 |
| $N = 32$ | 0.0326 | 0.00001 | 0.00008 | 0.0326 | 0.0326 | 0.00008 | 1.3157 | 0.9974 | 0.9772 |
| $N = 64$ | 0.0162 | 0.00000 | 0.00004 | 0.0162 | 0.0162 | 0.00004 | −0.9976 | 1.0000 | 1.0000 |
| $N = 128$ | 0.0082 | 0.00000 | 0.00002 | 0.0082 | 0.0082 | 0.00002 | −0.8058 | 1.0000 | 1.0000 |
| $N = 256$ | 0.0042 | 0.00000 | 0.00001 | 0.0042 | 0.0042 | 0.00001 | 0.4128 | 1.0000 | 1.0000 |

$ASD_K$: Asymptotic standard deviation of $V_{rel}(\mathbf{R})$ based on Konishi's (1979) theory (eq. 39).
$ASD_{PF}$: Approximate standard deviation of $V_{rel}(\mathbf{R})$ based on Pan & Frank's (2004) approach (eqs. 36–38).

(*continued*)

**Table S3.** (*continued*)

| | $E[V_{rel}(\mathbf{R})]$ | $ASD_K$ | $ASD_{PF}$ | Median | Mean | ESD | $T$ | Pow. 5% | Pow. 1% |
|---|---|---|---|---|---|---|---|---|---|
| $p = 2$, quadratically decreasing λ ($V_{rel}(\mathbf{P}) = 0.3600$) | | | | | | | | | |
| $N = 4$ | 0.4951 | 0.4434 | 0.3210* | 0.5073 | 0.4913 | 0.3196 | −0.8490 | 0.1268 | 0.0234 |
| $N = 8$ | 0.4006 | 0.2903 | 0.2490* | 0.3987 | 0.3998 | 0.2514 | −0.2231 | 0.3764 | 0.1568 |
| $N = 16$ | 0.3752 | 0.1983 | 0.1848* | 0.3766 | 0.3740 | 0.1853 | −0.4712 | 0.7346 | 0.4780 |
| $N = 32$ | 0.3665 | 0.1379 | 0.1335* | 0.3654 | 0.3654 | 0.1326 | −0.6016 | 0.9654 | 0.8998 |
| $N = 64$ | 0.3630 | 0.0968 | 0.0952* | 0.3640 | 0.3637 | 0.0954 | 0.4645 | 1.0000 | 0.9978 |
| $N = 128$ | 0.3615 | 0.0681 | 0.0676* | 0.3612 | 0.3622 | 0.0675 | 0.7336 | 1.0000 | 1.0000 |
| $N = 256$ | 0.3607 | 0.0481 | 0.0479* | 0.3615 | 0.3603 | 0.0482 | −0.5713 | 1.0000 | 1.0000 |
| $p = 4$, quadratically decreasing λ ($V_{rel}(\mathbf{P}) = 0.1911$) | | | | | | | | | |
| $N = 4$ | 0.4160 | 0.1703 | 0.1781 | 0.3858 | 0.4156 | 0.1516 | −0.1564 | 0.1596 | 0.0568 |
| $N = 8$ | 0.2766 | 0.1115 | 0.1111 | 0.2647 | 0.2764 | 0.1019 | −0.1091 | 0.4706 | 0.2268 |
| $N = 16$ | 0.2284 | 0.0762 | 0.0754 | 0.2182 | 0.2269 | 0.0722 | −1.4661 | 0.9412 | 0.7528 |
| $N = 32$ | 0.2085 | 0.0530 | 0.0526 | 0.2054 | 0.2086 | 0.0524 | 0.1041 | 1.0000 | 0.9992 |
| $N = 64$ | 0.1995 | 0.0372 | 0.0370 | 0.1969 | 0.1991 | 0.0367 | −0.8045 | 1.0000 | 1.0000 |
| $N = 128$ | 0.1953 | 0.0262 | 0.0261 | 0.1942 | 0.1954 | 0.0261 | 0.3599 | 1.0000 | 1.0000 |
| $N = 256$ | 0.1932 | 0.0185 | 0.0184 | 0.1928 | 0.1931 | 0.0182 | −0.2212 | 1.0000 | 1.0000 |
| $p = 8$, quadratically decreasing λ ($V_{rel}(\mathbf{P}) = 0.0980$) | | | | | | | | | |
| $N = 4$ | 0.3757 | 0.0545 | 0.1055 | 0.3645 | 0.3773 | 0.0786 | 1.4665 | 0.1776 | 0.0600 |
| $N = 8$ | 0.2114 | 0.0357 | 0.0537 | 0.2056 | 0.2111 | 0.0468 | −0.3972 | 0.5748 | 0.3076 |
| $N = 16$ | 0.1496 | 0.0244 | 0.0309 | 0.1462 | 0.1493 | 0.0289 | −0.8012 | 0.9890 | 0.9326 |
| $N = 32$ | 0.1227 | 0.0169 | 0.0193 | 0.1213 | 0.1226 | 0.0188 | −0.3686 | 1.0000 | 1.0000 |
| $N = 64$ | 0.1101 | 0.0119 | 0.0127 | 0.1094 | 0.1100 | 0.0126 | −0.5086 | 1.0000 | 1.0000 |
| $N = 128$ | 0.1040 | 0.0084 | 0.0087 | 0.1038 | 0.1040 | 0.0086 | −0.2554 | 1.0000 | 1.0000 |
| $N = 256$ | 0.1010 | 0.0059 | 0.0060 | 0.1008 | 0.1010 | 0.0060 | 0.4807 | 1.0000 | 1.0000 |
| $p = 16$, quadratically decreasing λ ($V_{rel}(\mathbf{P}) = 0.0496$) | | | | | | | | | |
| $N = 4$ | 0.3560 | 0.0200 | 0.0623 | 0.3485 | 0.3558 | 0.0395 | −0.3027 | 0.2010 | 0.0854 |
| $N = 8$ | 0.1786 | 0.0131 | 0.0296 | 0.1757 | 0.1785 | 0.0244 | −0.2391 | 0.6058 | 0.3646 |
| $N = 16$ | 0.1092 | 0.0089 | 0.0155 | 0.1079 | 0.1092 | 0.0144 | −0.0334 | 0.9916 | 0.9524 |
| $N = 32$ | 0.0783 | 0.0062 | 0.0088 | 0.0777 | 0.0783 | 0.0084 | −0.2148 | 1.0000 | 1.0000 |
| $N = 64$ | 0.0637 | 0.0044 | 0.0053 | 0.0634 | 0.0637 | 0.0052 | −0.0525 | 1.0000 | 1.0000 |
| $N = 128$ | 0.0565 | 0.0031 | 0.0034 | 0.0564 | 0.0565 | 0.0034 | −0.8445 | 1.0000 | 1.0000 |
| $N = 256$ | 0.0530 | 0.0022 | 0.0023 | 0.0530 | 0.0531 | 0.0023 | 1.9474 | 1.0000 | 1.0000 |
| $p = 32$, quadratically decreasing λ ($V_{rel}(\mathbf{P}) = 0.0249$) | | | | | | | | | |
| $N = 4$ | 0.3444 | 0.0071 | 0.0323 | 0.3400 | 0.3443 | 0.0204 | −0.1348 | 0.2064 | 0.0736 |
| $N = 8$ | 0.1605 | 0.0047 | 0.0149 | 0.1591 | 0.1605 | 0.0123 | −0.2153 | 0.5998 | 0.3796 |
| $N = 16$ | 0.0879 | 0.0032 | 0.0075 | 0.0873 | 0.0877 | 0.0068 | −1.3347 | 0.9948 | 0.9652 |
| $N = 32$ | 0.0553 | 0.0022 | 0.0040 | 0.0551 | 0.0554 | 0.0038 | 1.4091 | 1.0000 | 1.0000 |
| $N = 64$ | 0.0398 | 0.0016 | 0.0023 | 0.0397 | 0.0398 | 0.0022 | −0.7987 | 1.0000 | 1.0000 |
| $N = 128$ | 0.0323 | 0.0011 | 0.0014 | 0.0323 | 0.0323 | 0.0014 | 0.6379 | 1.0000 | 1.0000 |
| $N = 256$ | 0.0286 | 0.0008 | 0.0009 | 0.0286 | 0.0286 | 0.0009 | 1.7252 | 1.0000 | 1.0000 |

$ASD_K$: Asymptotic standard deviation of $V_{rel}(\mathbf{R})$ based on Konishi's (1979) theory (eq. 39).
$ASD_{PF}$: Approximate standard deviation of $V_{rel}(\mathbf{R})$ based on Pan & Frank's (2004) approach (eqs. 36–38).
*: Exact standard deviations for $p = 2$ are shown in these cells.

(*continued*)

**Table S3.** (*continued*)

| | $E[V_{rel}(\mathbf{R})]$ | $ASD_K$ | $ASD_{PF}$ | Median | Mean | ESD | $T$ | Pow. 5% | Pow. 1% |
|---|---|---|---|---|---|---|---|---|---|
| $p = 64$, quadratically decreasing λ ($V_{rel}(\mathbf{P}) = 0.0125$) | | | | | | | | | |
| $N = 4$ | 0.3391 | 0.00251 | 0.01660 | 0.3372 | 0.3392 | 0.01029 | 0.8681 | 0.2072 | 0.0882 |
| $N = 8$ | 0.1519 | 0.00165 | 0.00756 | 0.1512 | 0.1518 | 0.00621 | −0.3255 | 0.6328 | 0.3828 |
| $N = 16$ | 0.0774 | 0.00112 | 0.00371 | 0.0773 | 0.0775 | 0.00340 | 2.0673 | 0.9936 | 0.9736 |
| $N = 32$ | 0.0439 | 0.00078 | 0.00190 | 0.0438 | 0.0439 | 0.00181 | 0.0530 | 1.0000 | 1.0000 |
| $N = 64$ | 0.0279 | 0.00055 | 0.00102 | 0.0279 | 0.0279 | 0.00100 | 2.2327 | 1.0000 | 1.0000 |
| $N = 128$ | 0.0201 | 0.00039 | 0.00058 | 0.0201 | 0.0201 | 0.00057 | −2.0319 | 1.0000 | 1.0000 |
| $N = 256$ | 0.0163 | 0.00027 | 0.00035 | 0.0163 | 0.0163 | 0.00035 | 0.4608 | 1.0000 | 1.0000 |
| $p = 128$, quadratically decreasing λ ($V_{rel}(\mathbf{P}) = 0.0062$) | | | | | | | | | |
| $N = 4$ | 0.3362 | 0.00089 | 0.00838 | 0.3351 | 0.3362 | 0.00519 | 0.6706 | 0.2084 | 0.0942 |
| $N = 8$ | 0.1473 | 0.00058 | 0.00379 | 0.1471 | 0.1474 | 0.00304 | 0.3257 | 0.6450 | 0.4050 |
| $N = 16$ | 0.0720 | 0.00040 | 0.00184 | 0.0719 | 0.0720 | 0.00165 | −0.1768 | 0.9946 | 0.9700 |
| $N = 32$ | 0.0381 | 0.00028 | 0.00092 | 0.0380 | 0.0380 | 0.00087 | −1.2068 | 1.0000 | 1.0000 |
| $N = 64$ | 0.0219 | 0.00019 | 0.00048 | 0.0219 | 0.0219 | 0.00046 | −1.1815 | 1.0000 | 1.0000 |
| $N = 128$ | 0.0140 | 0.00014 | 0.00026 | 0.0140 | 0.0140 | 0.00025 | 0.1212 | 1.0000 | 1.0000 |
| $N = 256$ | 0.0101 | 0.00010 | 0.00014 | 0.0101 | 0.0101 | 0.00014 | −0.2988 | 1.0000 | 1.0000 |
| $p = 256$, quadratically decreasing λ ($V_{rel}(\mathbf{P}) = 0.0031$) | | | | | | | | | |
| $N = 4$ | 0.3347 | 0.00031 | 0.00421 | 0.3342 | 0.3347 | 0.00253 | −0.0296 | 0.2004 | 0.0746 |
| $N = 8$ | 0.1451 | 0.00020 | 0.00190 | 0.1449 | 0.1451 | 0.00153 | −0.8455 | 0.6272 | 0.3982 |
| $N = 16$ | 0.0693 | 0.00014 | 0.00091 | 0.0693 | 0.0693 | 0.00082 | −1.5371 | 0.9944 | 0.9700 |
| $N = 32$ | 0.0352 | 0.00010 | 0.00045 | 0.0351 | 0.0352 | 0.00044 | 0.3984 | 1.0000 | 1.0000 |
| $N = 64$ | 0.0189 | 0.00007 | 0.00023 | 0.0189 | 0.0189 | 0.00022 | −1.3213 | 1.0000 | 1.0000 |
| $N = 128$ | 0.0109 | 0.00005 | 0.00012 | 0.0109 | 0.0109 | 0.00012 | −2.1747 | 1.0000 | 1.0000 |
| $N = 256$ | 0.0070 | 0.00003 | 0.00006 | 0.0070 | 0.0070 | 0.00006 | 0.5172 | 1.0000 | 1.0000 |
| $p = 1024$, quadratically decreasing λ ($V_{rel}(\mathbf{P}) = 0.0008$) | | | | | | | | | |
| $N = 4$ | 0.3337 | 0.000039 | 0.00106 | 0.3336 | 0.3337 | 0.00062 | −0.0472 | 0.1970 | 0.0824 |
| $N = 8$ | 0.1434 | 0.000026 | 0.00048 | 0.1434 | 0.1434 | 0.00039 | 0.3587 | 0.6418 | 0.3862 |
| $N = 16$ | 0.0673 | 0.000017 | 0.00023 | 0.0673 | 0.0673 | 0.00020 | −1.8148 | 0.9940 | 0.9722 |
| $N = 32$ | 0.0330 | 0.000012 | 0.00011 | 0.0330 | 0.0330 | 0.00010 | 0.7257 | 1.0000 | 1.0000 |
| $N = 64$ | 0.0166 | 0.000009 | 0.00006 | 0.0166 | 0.0166 | 0.00005 | −1.0370 | 1.0000 | 1.0000 |
| $N = 128$ | 0.0086 | 0.000006 | 0.00003 | 0.0086 | 0.0086 | 0.00003 | −1.2065 | 1.0000 | 1.0000 |
| $N = 256$ | 0.0047 | 0.000004 | 0.00001 | 0.0047 | 0.0047 | 0.00001 | 0.0830 | 1.0000 | 1.0000 |

$ASD_K$: Asymptotic standard deviation of $V_{rel}(\mathbf{R})$ based on Konishi's (1979) theory (eq. 39).
$ASD_{PF}$: Approximate standard deviation of $V_{rel}(\mathbf{R})$ based on Pan & Frank's (2004) approach (eqs. 36–38).

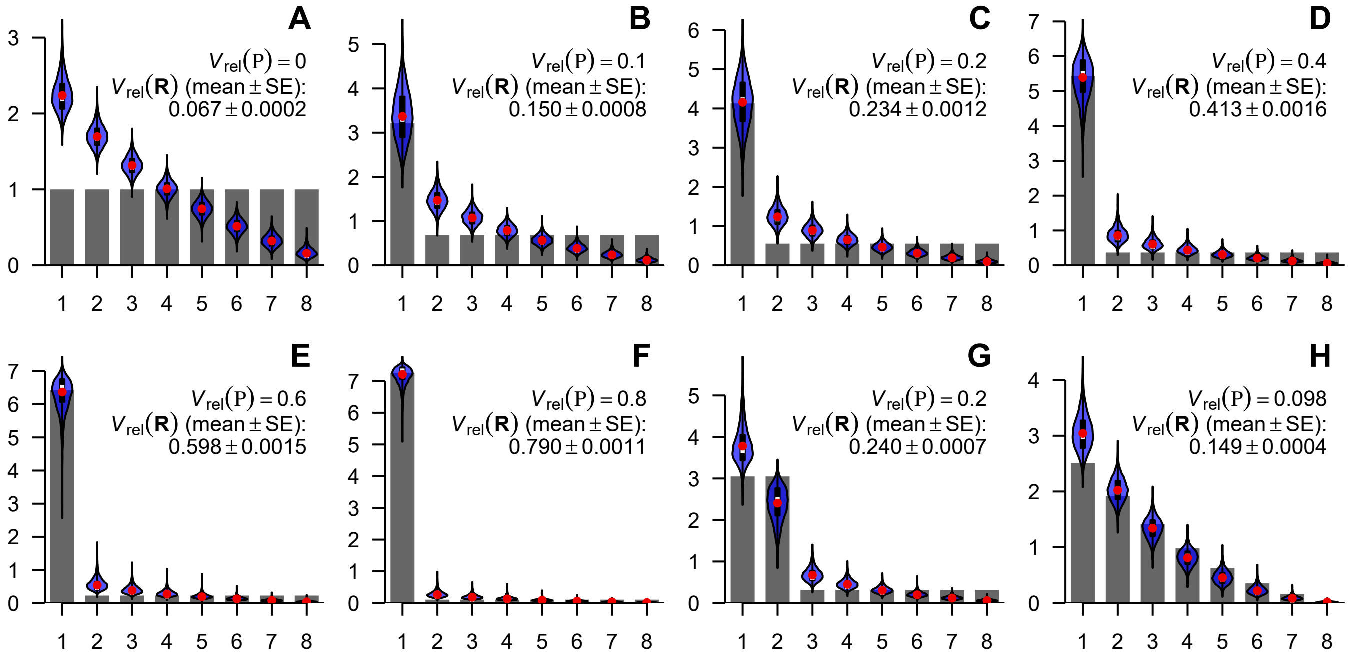


**Figure S1.** Selected population eigenvalue structures used in simulations and distributions of sample eigenvalues, examples for $p = 8$. The eigenvalues of population correlation matrix are shown as scree plots, and distributions of sample eigenvalues with $N = 16$ are shown as violin plots. The conditions are as in Figure 4. **A**, null condition; **B–G**, $q$-large λ conditions, $q = 1$ (**B–F**) or 2 (**G**), with $V_{rel}(\mathbf{\Sigma}) = 0.1, 0.2, 0.4, 0.6, 0.8$, and 0.2, respectively; **H**, quadratically decreasing λ condition. Red dots denote empirical means of sample eigenvalues, whereas white bars (mostly overlapping with red dots) denote medians. Thick black bars within violins denote interquartile ranges. Note different scales of vertical axes.

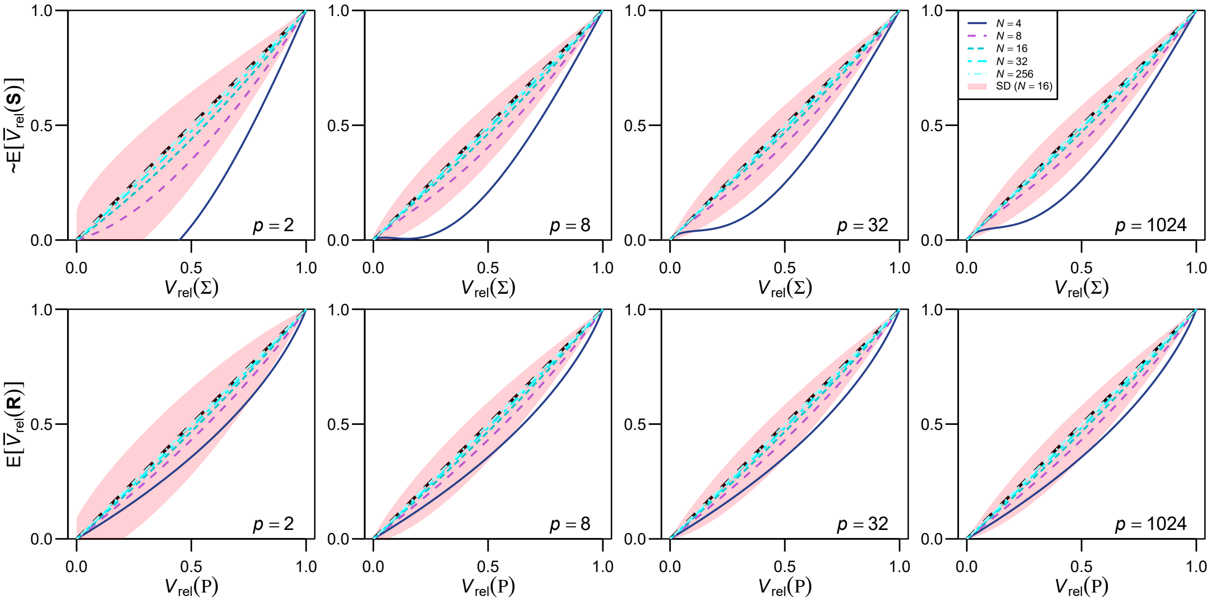


**Figure S2.** Profiles of the expectations of adjusted eigenvalue dispersion indices in selected conditions. The expectations of $\bar{V}_{rel}(\mathbf{S})$ (approximate; top row) and $\bar{V}_{rel}(\mathbf{R})$ (bottom row) are drawn with solid lines, for $p = 2$, 8, 32, and 1024 (from left to right) and for $N = 4$, 8, 16, 32, and 256. In all cases, $n = N - 1$. The breadth of one standard deviation at $N = 16$ is also shown around the mean profiles with pink fills; these are approximations except for $\bar{V}_{rel}(\mathbf{R})$ for $p = 2$ (variance of $\bar{V}_{rel}(\mathbf{R})$ is from eq. 39; eqs. 36–38 yielded similar values under these conditions). Note that actual distributions might be skewed unlike these fills. All profiles are from 1-large λ conditions as in Figure 2. The initial decrease of the E[$\bar{V}_{rel}(\mathbf{S})$] profiles in some cases seems to be an artifact of approximation, so the negative excursion in expectations should be ignored.

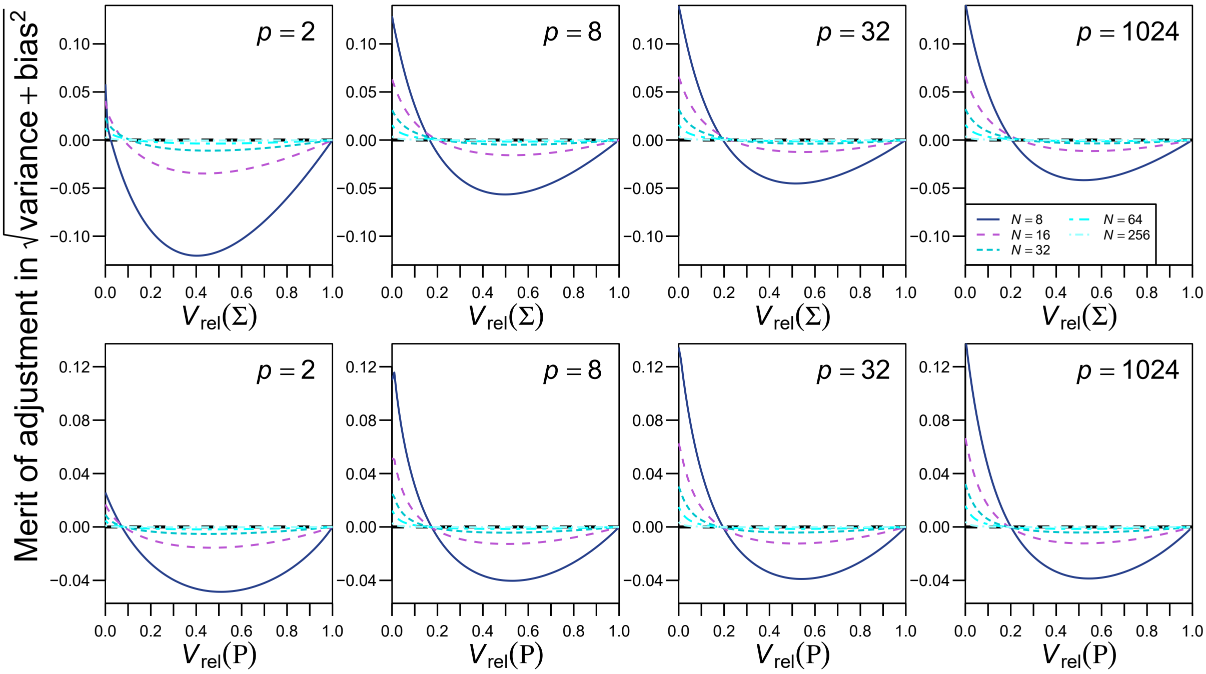


**Figure S3.** Merit of adjustment for eigenvalue dispersion indices. Square root of inaccuracy (= Var[$V_{\mathrm{rel}}(\mathbf{S})$] + {E[$V_{\mathrm{rel}}(\mathbf{S})$] − $V_{\mathrm{rel}}(\mathbf{\Sigma})$}$^2$, and so on) is compared between $\overline{V}_{\mathrm{rel}}(\mathbf{S})$ and $V_{\mathrm{rel}}(\mathbf{S})$ (top row), and between $\overline{V}_{\mathrm{rel}}(\mathbf{R})$ and $V_{\mathrm{rel}}(\mathbf{R})$ (bottom row) across varying parameter values, for $p$ = 2, 8, 32, and 1024 (from left to right) and for $N$ = 8, 16, 32, 64, and 256. In all cases, $n = N - 1$. Positive values correspond to smaller inaccuracy in the adjusted indices—merit of adjustment. Note that these profiles are approximations, except under the null conditions (left-hand end of each plot) and for $V_{\mathrm{rel}}(\mathbf{R})$ with $p$ = 2. Variance of $V_{\mathrm{rel}}(\mathbf{R})$ shown is from eq. 39; eqs. 36–38 yielded similar values under these conditions. Deflection of the profile near the null condition in some cases correspond to a discrepancy between exact and approximate results. All profiles are from 1-large λ conditions as in Figure 2.

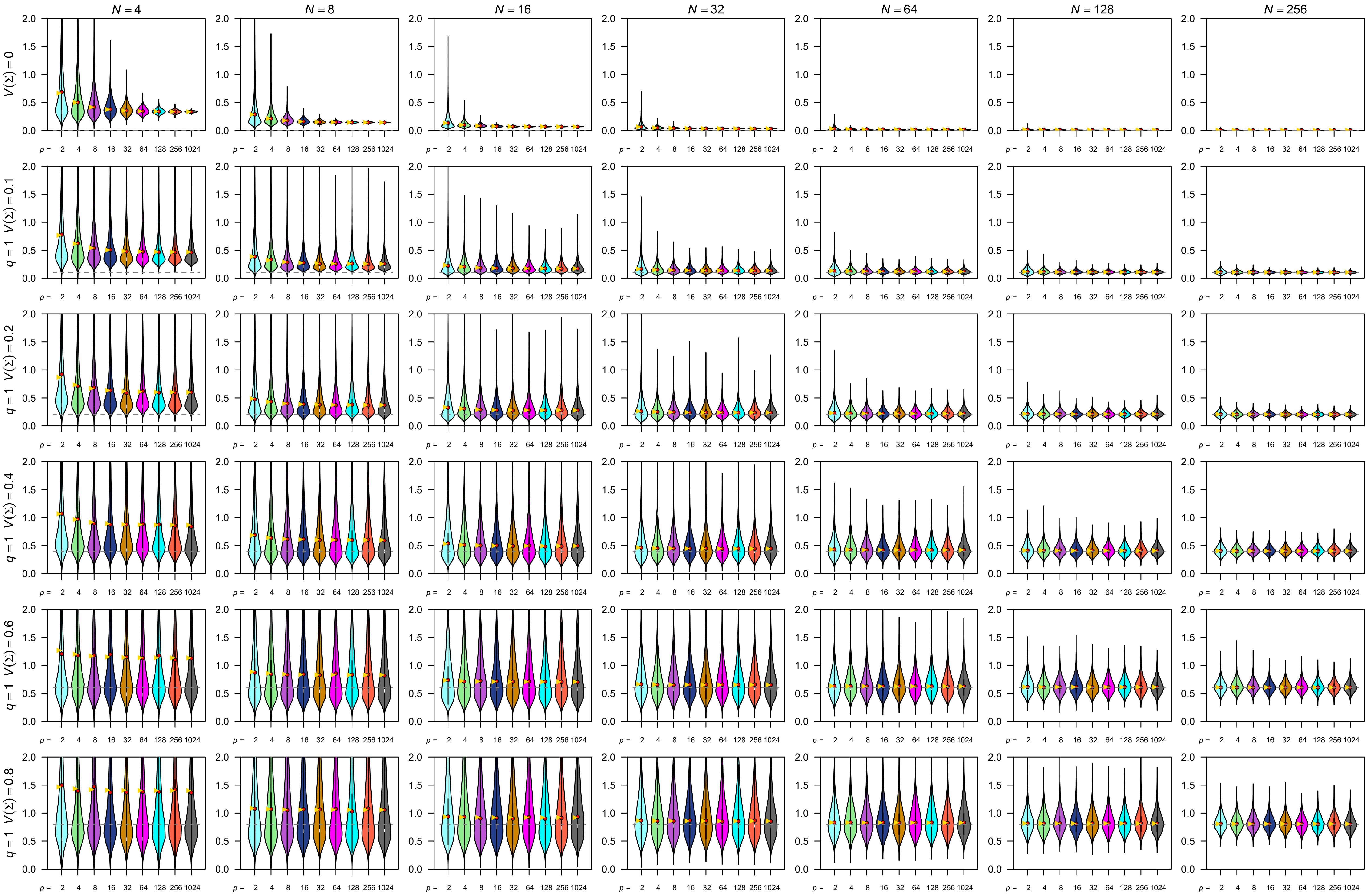

**Figure S4.** Simulation results for the eigenvalue variance of covariance matrix $V(\mathbf{S})$, $q = 1$. See Figure 5 for legends. Note that extreme values in some panels are cropped for visual clarity.

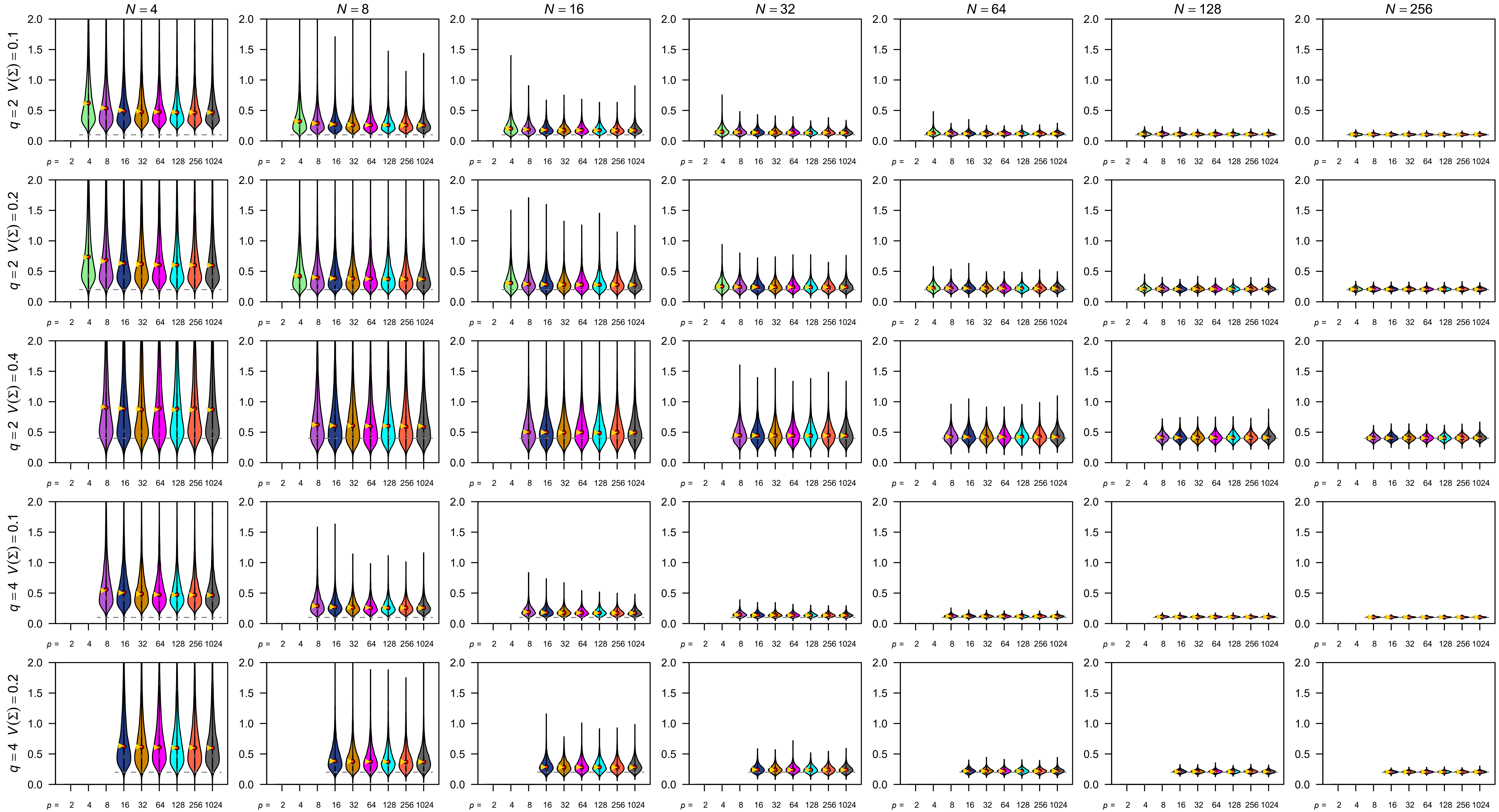


**Figure S5.** Simulation results for the eigenvalue variance of covariance matrix $V(\mathbf{S})$, $q = 2$ and 4. See Figure 5 for legends. Note that extreme values in some panels are cropped for visual clarity.

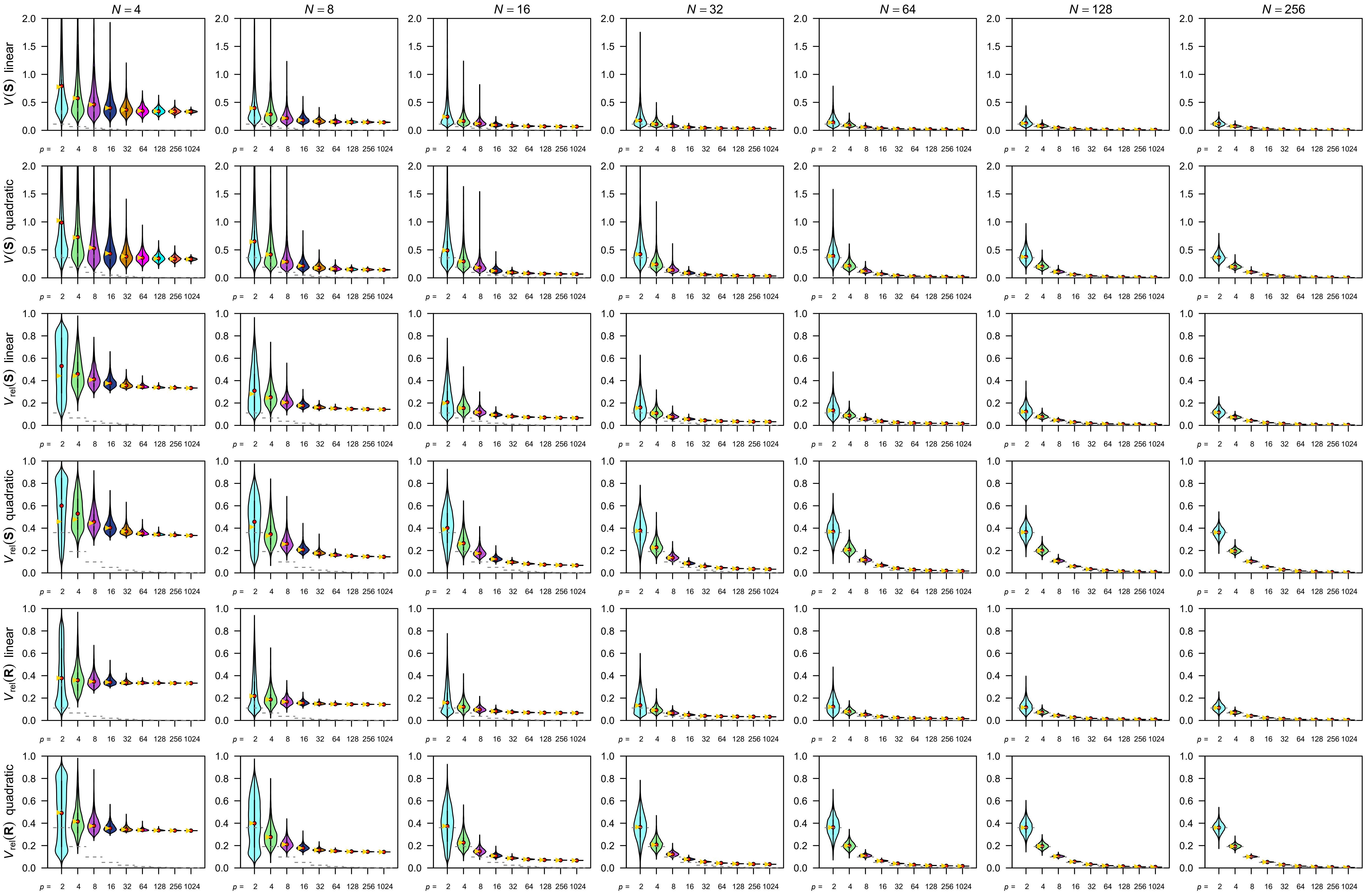


**Figure S6.** Simulation results, linearly- and quadratically decreasing λ conditions. Empirical distributions of simulated $V(\mathbf{S})$, $V_{\text{rel}}(\mathbf{S})$, and $V_{\text{rel}}(\mathbf{R})$ are shown in violin plots. See Figure 5 for legends. Note that extreme values in some panels for $V(\mathbf{S})$ are cropped for visual clarity.

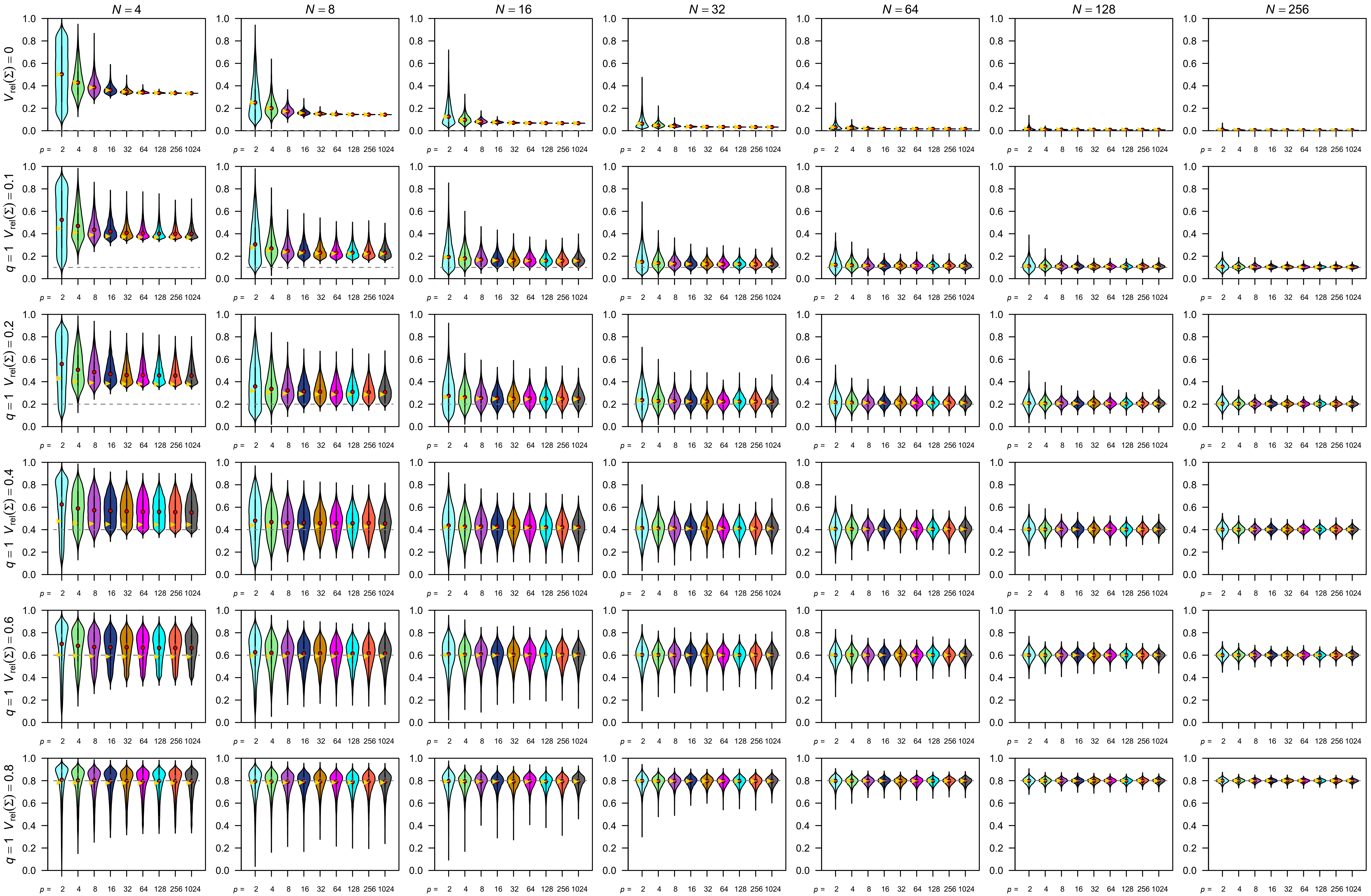


**Figure S7.** Simulation results for the relative eigenvalue variance of covariance matrix $V_{rel}(\mathbf{S})$, $q = 1$. See Figure 5 for legends.

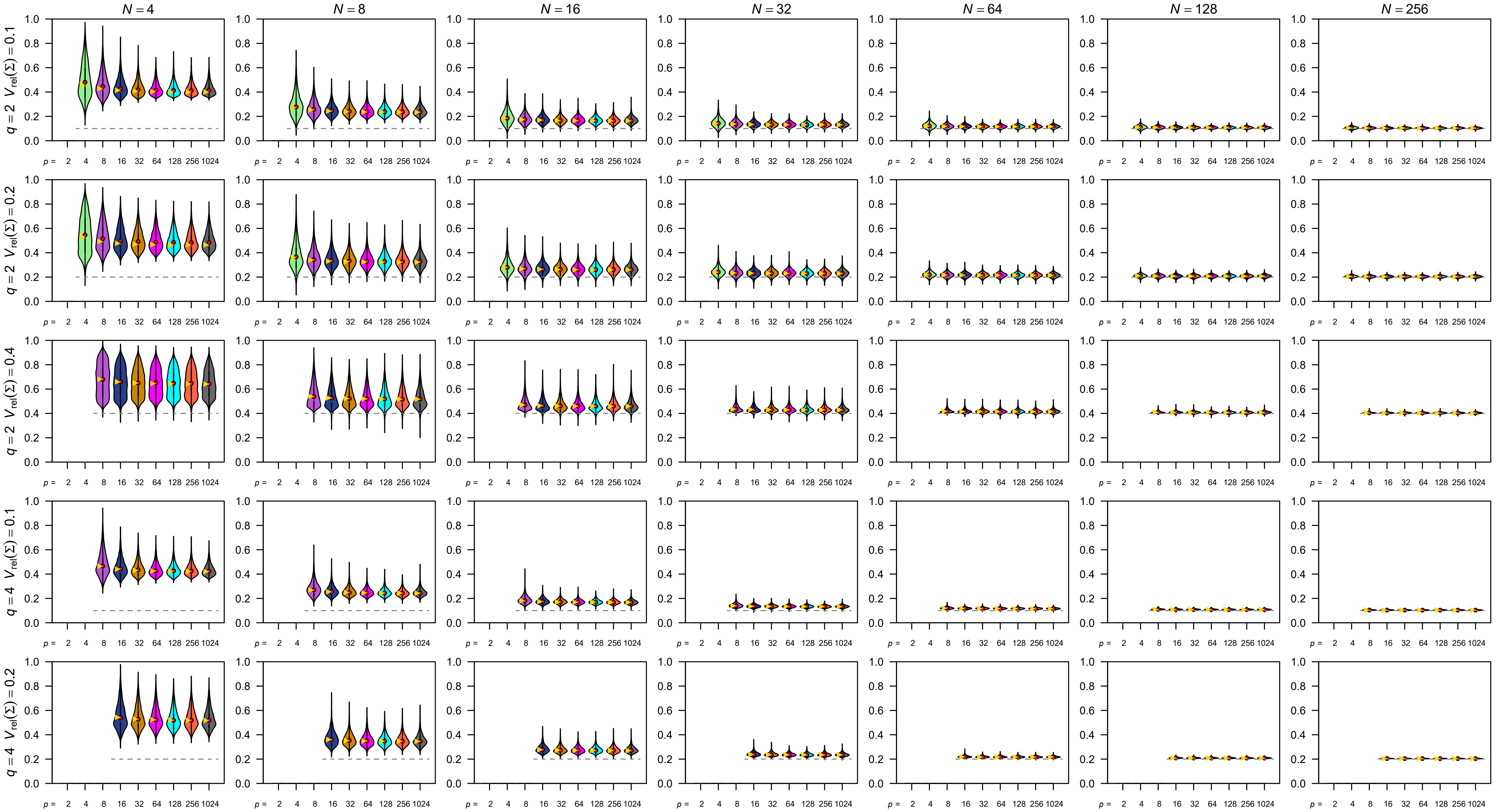


**Figure S8.** Simulation results for the relative eigenvalue variance of covariance matrix $V_{\text{rel}}(\mathbf{S})$, $q = 2$ and 4. See Figure 5 for legends.

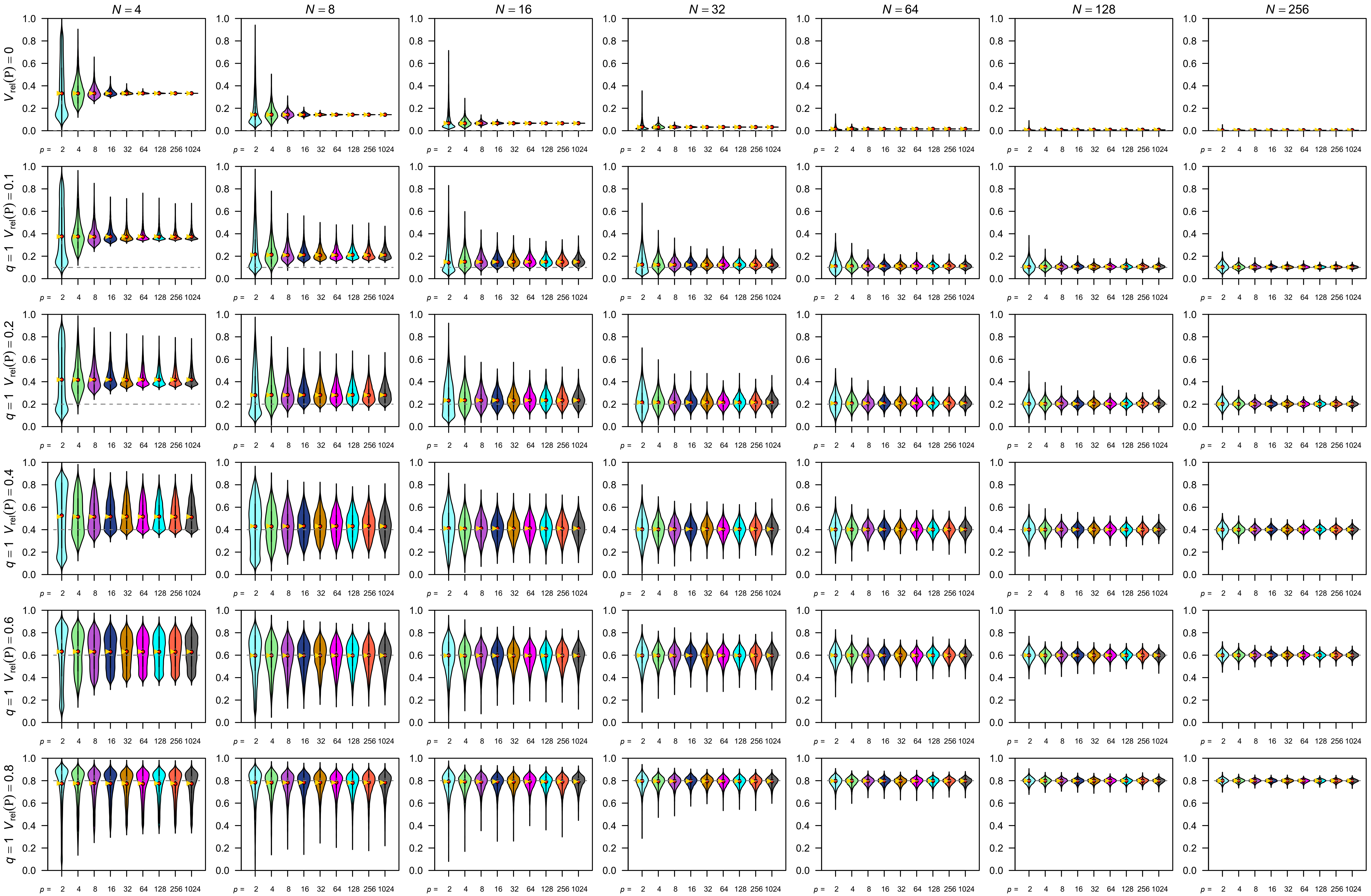


**Figure S9.** Simulation results for the relative eigenvalue variance of correlation matrix $V_{rel}(\mathbf{R})$, $q = 1$. See Figure 5 for legends. Note that a peak is usually present near 0 when $p = 2$ and $N \leq 8$, but this is invisible due to smoothing in the violin plots.

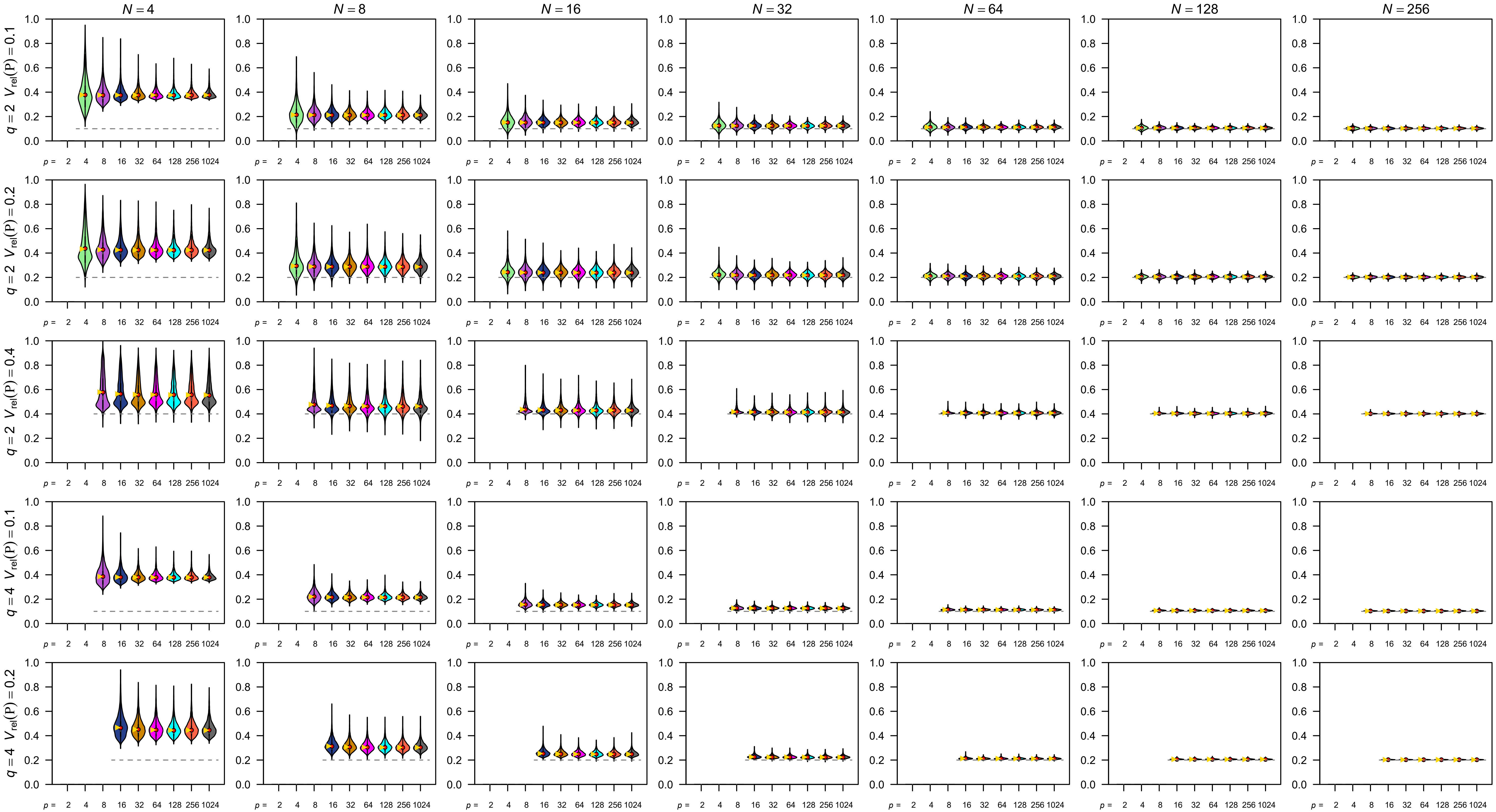


**Figure S10.** Simulation results for the relative eigenvalue variance of correlation matrix $V_{\mathrm{rel}}(\mathbf{R})$, $q = 2$ and 4. See Figure 5 for legends.
