## Supplemental codes for "Statistics of eigenvalue dispersion indices: quantifying the magnitude of phenotypic integration": _README.txt

###############################
# Supplementary material for
#    Statistics of eigenvalue dispersion indices:
#    quantifying the magnitude of phenotypic integration
#      by Junya Watanabe
# This package involves the following files:
# - codes.R: codes to be executed on R to reproduce the simulation results
# - functions.R: R functions used in simulations, as well as those to calculate
#                moments of eigenvalue dispersion indices
# - Cov_r2C.cpp: C++ code called in AVar.VRr_pfc defined in functions.R
#                for efficient calculation of Var[Vrel(R)]
#                to be used with the package Rcpp
###############################
