## Supplemental codes for "Statistics of eigenvalue dispersion indices: quantifying the magnitude of phenotypic integration": codes.R

```
###############################
# Supplementary material for
#    Statistics of eigenvalue dispersion indices:
#    quantifying the magnitude of phenotypic integration
#      by Junya Watanabe
# This file involves the code to reproduce the simulation results in the paper.
# The population covariance matrices and simulation results are created
# using functions defined in "functions.R".
# The simulations will take several hours.
# Version 2021.08.06
###############################

source("functions.R")
Ns <- c(4, 8, 16, 32, 64, 128, 256)
ps <- c(2, 4, 8, 16, 32, 64, 128, 256, 1024)
vrs <- c(0, 0.1, 0.2, 0.4, 0.6, 0.8)
qs <- c(1, 2, 4)
Evs <- "c(rep.int(1 + sqrt(vr) * sqrt((p - 1) * (p - q) / q), q), rep.int(1 - sqrt(vr) * sqrt(q * (p - 1) / (p - q)), p - q)) / sqrt(p - 1)"
Evs_l <- "2 * (p + 1 - seq_len(p)) / (p + 1) / sqrt(p - 1)"
Evs_q <- "(6 * (p + 1 - seq_len(p)) ^ 2) / ((p + 1) * (2 * p + 1)) / sqrt(p - 1)"

b <- 5000L

for(j in seq_along(ps)) {
    p <- ps[j]
    cat(paste("\nCommencing p =", p))
    for(k in seq_along(vrs)) {
        vr <- vrs[k]
        cat(paste("\n VR =", vr))
        for(l in seq_along(qs)) {
            q <- qs[l]
            if((vr == 0 && q > 1) || p <= q) next
            ev <- eval(parse(text = Evs))
            if(any(ev < 0)) next
            cat(paste(" q =", q))
            V <- GenCov(evalues = ev, evectors = "Givens",
                        scale = p / sqrt(p - 1), random = FALSE)
            cV <- if(sum(ev == 0) > 1) chol.qr(V) else chol.piv(V)
            assign(paste0("V_p", sprintf("%04.0f", p), "vr", sprintf("%02.0f", vr * 10), "_q", sprintf("%01.0f", q)), V)
            for(i in seq_along(Ns)) {
                N <- Ns[i]
                cat(paste(". N =", N))
                set.seed(123456 * i + 78901 * j + 2345 * k + 6789012 * l)
                invisible({gc(); gc()})
                assign(paste0("simn_n", sprintf("%03.0f", N), "p", sprintf("%04.0f", p), "vr", sprintf("%02.0f", vr * 10), "_q", sprintf("%01.0f", q)), Sim.VE(N = N, cV = cV, b = b, nv = 0))
                invisible({gc(); gc()})
            }
        }
    }
    cat("\n lin/qua")
    ev <- eval(parse(text = Evs_l))
    V <- GenCov(evalues = ev, evectors = "Givens",
                scale = p / sqrt(p - 1), random = FALSE)
    cV <- chol.piv(V)
    assign(paste0("V_p", sprintf("%04.0f", p), "_lindec"), V)
    for(i in seq_along(Ns)) {
        N <- Ns[i]
        invisible({gc(); gc()})
        set.seed(12345 * i + 67890 * j)
        assign(paste0("simn_n", sprintf("%03.0f", N), "p", sprintf("%04.0f", p), "_lindec"), Sim.VE(N = N, cV = cV, b = b, nv = 0))
        invisible({gc(); gc()})
    }
    ev <- eval(parse(text = Evs_q))
    V <- GenCov(evalues = ev, evectors = "Givens",
                scale = p / sqrt(p - 1), random = FALSE)
    cV <- chol.piv(V)
    assign(paste0("V_p", sprintf("%04.0f", p), "_quadec"), V)
    for(i in seq_along(Ns)) {
        N <- Ns[i]
        invisible({gc(); gc()})
        set.seed(123456 * i + 789012 * j)
        assign(paste0("simn_n", sprintf("%03.0f", N), "p", sprintf("%04.0f", p), "_quadec"), Sim.VE(N = N, cV = cV, b = b, nv = 0))
        invisible({gc(); gc()})
    }
    cat("\n")
}
```
