## Supplemental codes for "Statistics of eigenvalue dispersion indices: quantifying the magnitude of phenotypic integration": Cov_r2C.cpp

```
#include <Rcpp.h>
using namespace Rcpp;

/* *** Cov.r2C ***
* Supplementary material for
*    Statistics of eigenvalue dispersion indices:
*    quantifying the magnitude of phenotypic integration
*      by Junya Watanabe
* Version 2021.08.06
* Function to calculate sum of covariances of all non-redundant pairs of
* squared correlation coefficients.
* This is a C++ code to be used in within AVar.VRr_pfc with the package Rcpp:
* R> require(Rcpp); sourceCpp("[path/to/]Cov_r2.cpp")
* Arguments:
*  n: Degree(s) of freedom; numeric of length 1 or more
*  R: Correlation matrix; numeric of length p * p
*  E: Matrix of expectation of correlation coefficients corresponding to
*     the upper triangular of R; numeric of length p(p - 1) / 2 * ln
*     where ln is the length of n
* R and E are read as numeric vectors to save a modest amount of time
* Value:
*  numeric of length ln
*/

// [[Rcpp::export]]
NumericVector Cov_r2C(NumericVector n, NumericVector R, NumericVector E) {
    int ln = n.size();
    int p = sqrt(R.size()); // If R was read as NumericMatrix, int p = R.nrow()
    int nrE = p * (p - 1) / 2;
    double rij, rkl, rik, ril, rjk, rjl, C;
    double Eij, Ekl, Cijkl;
    NumericVector out(ln);
    for(int i = 0; i < p - 1; i++) {
        for(int j = i + 1; j < p; j++) {
            for(int k = i; k < p - 1; k++) {
                for(int l = ((i == k) ? (j + 1) : (k + 1)); l < p; l++) {
                    rij = R[i + j * p];  // If R was a NumericMatrix, R(i, j)
                    rkl = R[k + l * p];  // Similar for Eij below
                    rik = R[i + k * p];
                    rjl = R[j + l * p];
                    ril = R[i + l * p];
                    rjk = R[j + k * p];
                    C = rij * rkl * (rik * rik + ril * ril +
                                     rjk * rjk + rjl * rjl) / 2 +
                        rik * rjl + ril * rjk - (rij * rik * ril
                        + rij * rjk * rjl + rik * rjk * rkl + ril * rjl * rkl);
                    for(int m = 0; m < ln; m++) {
                        Eij = E[i + j * p - j * (2 * p - j + 1) / 2 + m * nrE];
                        Ekl = E[k + l * p - l * (2 * p - l + 1) / 2 + m * nrE];
                        Cijkl = C / n[m];
                        out[m] += (4 * Eij * Ekl + 2 * Cijkl) * Cijkl;
                    }
                }
            }
        }
    } // for speed, out is doubled after skipping redundant pairs of (rij, rkl)
    out = out * 2;
    return out;
}
```
