## Supplemental codes for "Statistics of eigenvalue dispersion indices: quantifying the magnitude of phenotypic integration": functions.R

```
###############################
# Supplementary material for
#    Statistics of eigenvalue dispersion indices:
#    quantifying the magnitude of phenotypic integration
#      by Junya Watanabe
# This file involves R functions required to reproduce results of the paper,
# as well as functions to calculate expectation and variance of eigenvalue
# dispersion indices.
# Partly depends on the following packages:
# - hypergeo: For evaluation of the hypergeometric function in Exv/Var.VRr
# - parallel: For parallel computation in AVar.VRr_pf/d (optional)
# - Rcpp:     For use of a C++ code in AVar.VRr_pfc (optional)
# Version 2021.08.06
###############################

##### Exv.VEv #####
# Function to calculate expectation of eigenvalue variance for an arbitrary
# covariance matrix E[V(S)] based on a Wishart matrix.
## Arguments
# L:       a numeric vector of population eigenvalues
# V:       population covariance matrix; assumed to be validly constructed;
#          required when L is not provided
# n:       degree(s) of freedom; numeric of length 1 or more
# divisor: either "UB" (default) or "ML", to decide the default value of m
# m:       divisor of cov matrix to be used ("n*" in the paper)
# drop_0:  logical, when TRUE, eigenvalues smaller than tol are dropped
# tol:     tolerance/threshold to be used with drop_0
## Value
# A numeric vector which is E[V(S)] corresponding to n
## Details
# n is degree(s) of freedom, not sample size(s).
# By default, the divisor of n (which gives the ordinary unbiased estimator)
# is used.  Additional arguments (...) are accommodated for compatibility with
# other functions with more options but ignored for this function.
Exv.VEv <- function(V, L, n = 100, divisor = c("UB", "ML"),
                    m = switch(divisor, UB = n, ML = n + 1), drop_0 = FALSE,
                    tol = .Machine$double.eps * 100, ...) {
    divisor <- match.arg(divisor)
    if(missing(L)) {
        L <- svd(V, nu = 0, nv = 0)$d
    }
    if(drop_0) {
        L <- L[L > tol]
    } else {
        L[L < tol] <- 0
    }
    p <- length(L)
    t1 <- sum(L)
    t2 <- sum(L ^ 2)
    n * ((p - n) * t1 ^ 2 + (p * n + p - 2) * t2) / (p ^ 2 * m ^ 2)
}

##### Exv.VRv #####
# Function to calculate expectation of relative eigenvalue variance for an
# arbitrary covariance matrix E[Vrel(S)] based on a Wishart matrix.
# This is an approximation based on the delta method, except under
# the null condition where the value coincides with the exact expectation.
# Usage is similar to Exv.VEv.
Exv.VRv <- function(V, L, n = 100, drop_0 = FALSE,
                    tol = .Machine$double.eps * 100, ...) {
    if(missing(L)) {
        L <- svd(V, nu = 0, nv = 0)$d
    }
    if(drop_0) {
        L <- L[L > tol]
    } else {
        L[L < tol] <- 0
    }
    p <- length(L)
    t1 <- sum(L)
    t2 <- sum(L ^ 2)
    t3 <- sum(L ^ 3)
    t4 <- sum(L ^ 4)
    EF <- (t1 ^ 2 + (n + 1) * t2) / (n * t1 ^ 2 + 2 * t2) -
          8 * ((n + 2) * (n - 1) * (3 * t4 * t1 ^ 2 - 2 * t3 * t2 * t1 - t2 ^ 3
                                    + n * t3 * t1 ^ 3 - n * t2 ^ 2 * t1 ^ 2)) /
          (n * ((n * t1 ^ 2 + 2 * t2)) ^ 3)
    (p * EF - 1) / (p - 1)
}

##### Exv.VEr #####
# Function to calculate expectation of eigenvalue variance for a correlation
# matrix E[V(R)] based on a Wishart matrix.
# This is just (p - 1) times Exv.VRr
Exv.VEr <- function(R, L, n = 100, tol = .Machine$double.eps * 100,
                    tol.hg = 0, maxiter.hg = 2000, ...) {
    if(missing(R)) {
        R <- GenCov(evalues = L, evectors = "Givens")
        if(L[2] != L[length(L)]) warning("R was generated from the eigenvalues provided \n  Expectation may vary even if eigenvalues are fixed")
    } else if(any(diag(R) != 1)) {
        R <- R / R[1, 1]
        if(any(diag(R) != 1)) stop("Provide a valid correlation matrix, or its eigenvalues")
        warning("R was scaled to have diagonal elements of unity")
    }
    p <- ncol(R)
    exv.VRr <- Exv.VRr(R = R, n = n, tol = tol,
                       tol.hg = tol.hg, maxiter.hg = maxiter.hg, ...)
    exv.VRr * (p - 1)
}

##### Exv.VRr #####
# Function to calculate expectation of relative eigenvalue variance for
# an arbitrary correlation matrix E[Vrel(R)] based on a Wishart matrix.
## Arguments
# R: a population correlation matrix; assumed to be validly constructed
# L: a vector of population eigenvalues
# n: degree(s) of freedom; numeric of length 1 or more
# tol, tol.hg, maxiter.hg: passed to Exv.r2; see description of that function
## Value
# A numeric vector which is E[Vrel(R)] corresponding to n
## Details
# As detailed in the paper, E[Vrel(R)] cannot be uniquely specified by
# the eigenvalues alone.  Hence, a full correlation matrix should preferably be
# provided.  Otherwise, a correlation matrix is constructed from the eigenvalues
# L provided using the function GenCov with randomly picked eigenvectors.
# When R is provided, some check is done: the matrix is scaled to have
# diagonals of 1; and if any of these are unequal, an error is returned.
# This function calls Exv.r2, which in turn calls hgf (both defined below).
## Dependency
# Depends on the package hypergeo
Exv.VRr <- function(R, L, n = 100, tol = .Machine$double.eps * 100,
                    tol.hg = 0, maxiter.hg = 2000, ...) {
    if(missing(R)) {
        R <- GenCov(evalues = L, evectors = "Givens")
        if(L[2] != L[length(L)]) warning("R was generated from the eigenvalues provided \n  Expectation may vary even if eigenvalues are fixed")
    } else if(any(diag(R) != 1)) {
        R <- R / R[1, 1]
        if(any(diag(R) != 1)) stop("Provide a valid correlation matrix, or its eigenvalues")
        warning("R was scaled to have diagonal elements of unity")
    }
    p <- ncol(R)
    R2 <- R[lower.tri(R)] ^ 2
    exv_r2 <- matrix(sapply(n, Exv.r2, x = R2, do.square = FALSE, tol = tol,
                            tol.hg = tol.hg, maxiter.hg = maxiter.hg),
                     ncol = length(n))
    2 * colSums(exv_r2) / (p * (p - 1))
}

##### Exv.VEav #####
# Function to calculate expectation of unbiased eigenvalue variance for an
# arbitrary covariance matrix E[V~(S)] based on a Wishart matrix.
# Of little practical use because this is just the population value V(Sigma).
# Usage is identical to Exv.VEv.
Exv.VEav <- function(V, L, n = 100, drop_0 = FALSE,
                     tol = .Machine$double.eps * 100, ...) {
    if(missing(L)) {
        L <- svd(V, nu = 0, nv = 0)$d
    }
    if(drop_0) {
        L <- L[L > tol]
    } else {
        L[L < tol] <- 0
    }
    p <- length(L)
    t1 <- sum(L)
    t2 <- sum(L ^ 2)
    t2 / p - t1 ^ 2 / p^2
}

##### Exv.VRav #####
# Function to calculate expectation of adjusted relative eigenvalue variance
# for an arbitrary covariance matrices E[V-rel(S)] based on a Wishart matrix.
# Usage is identical to Exv.VRv, which is called internally.
Exv.VRav <- function(V, L, n = 100, drop_0 = FALSE,
                     tol = .Machine$double.eps * 100, ...) {
    if(missing(L)) {
        L <- svd(V, nu = 0, nv = 0)$d
    }
    if(drop_0) {
        L <- L[L > tol]
    } else {
        L[L < tol] <- 0
    }
    p <- length(L)
    E <- Exv.VRv(L = L, n = n, drop_0 = drop_0, tol = tol, ...)
    En <- (p + 2) / (p * n + 2)
    1 - (1 - E) / (1 - En)
}

##### Exv.VRar #####
# Function to calculate expectation of adjusted relative eigenvalue variance
# for an arbitrary correlation matrices E[V-rel(R)] based on a Wishart matrix.
# Usage is identical to Exv.VRr, which is called internally.
Exv.VRar <- function(R, L, n = 100, tol = .Machine$double.eps * 100,
                     tol.hg = 0, maxiter.hg = 2000, ...) {
    if(missing(R)) {
        R <- GenCov(evalues = L, evectors = "Givens")
        if(L[2] != L[length(L)]) warning("R was generated from the eigenvalues provided \n  Expectation may vary even if eigenvalues are fixed")
    } else if(any(diag(R) != 1)) {
        R <- R / R[1, 1]
        if(any(diag(R) != 1)) stop("Provide a valid correlation matrix, or its eigenvalues")
        warning("R was scaled to have diagonal elements of unity")
    }
    p <- ncol(R)
    E <- Exv.VRr(R = R, n = n, tol = tol,
                 tol.hg = tol.hg, maxiter.hg = maxiter.hg, ...)
    En <- 1 / n
    1 - (1 - E) / (1 - En)
}

##### Var.VEv #####
# Function to calculate variance of eigenvalue variance for an arbitrary
# covariance matrix Var[V(S)] based on a Wishart matrix.
# Usage is similar to Exv.VEv.
Var.VEv <- function(V, L, n = 100, divisor = c("UB", "ML"),
                    m = switch(divisor, UB = n, ML = n + 1), drop_0 = FALSE,
                    tol = .Machine$double.eps * 100, ...) {
    divisor <- match.arg(divisor)
    if(missing(L)) {
        L <- svd(V, nu = 0, nv = 0)$d
    }
    if(drop_0) {
        L <- L[L > tol]
    } else {
        L[L < tol] <- 0
    }
    p <- length(L)
    t1 <- sum(L)
    t2 <- sum(L ^ 2)
    t3 <- sum(L ^ 3)
    t4 <- sum(L ^ 4)
    4 * n * ((2 * p^2 * n^2 + 5 * p^2 * n + 5 * p^2 - 12 * p * n - 12 * p + 12)
             * t4 + 4 * (p - n) * (p * n + p - 2) * t3 * t1 +
             (p^2 * n + p^2 - 4 * p + 2 * n) * t2^2 +
             2 * (p - n)^2 * t2 * t1^2) / (p ^ 4 * m ^ 4)
}

##### Var.VRv #####
# Function to calculate variance of relative eigenvalue variance for an
# arbitrary covariance matrix Var[Vrel(S)] based on a Wishart matrix.
# This is an approximation based on the delta method,
# except under the null condition where the exact variance is returned.
# Usage is similar to Exv.VRv.
Var.VRv <- function(V, L, n = 100, drop_0 = FALSE,
                    tol = .Machine$double.eps * 100, ...) {
    if(missing(L)) {
        L <- svd(V, nu = 0, nv = 0)$d
    }
    if(drop_0) {
        L <- L[L > tol]
    } else {
        L[L < tol] <- 0
    }
    p <- length(L)
    t1 <- sum(L)
    t2 <- sum(L ^ 2)
    t3 <- sum(L ^ 3)
    t4 <- sum(L ^ 4)
    if(isTRUE(all.equal(L / L[1], rep.int(1, p)))) {
        4 * p^2 * (p + 2) * (n + 2) * (n - 1) /
             (p - 1) / (p * n + 2)^2 / (p * n + 4) / (p * n + 6)
    } else {
        4 * p^2 / (p - 1)^2 * (n - 1) * (n + 2) *
            (- 4 * t4 * t2^2 - 4 * n * t4 * t2 * t1^2
             + (2 * n^2 + 3 * n - 6) * t4 * t1^4
             - 4 * (n - 1) * (n + 2) * t3 * t2 * t1^3
             + 2 * (n + 1) * t2^4 + 2 * n * (n + 1) * t2^3 * t1^2
             + n * t2^2 * t1^4) / n / (2 * t2 + n * t1^2)^4
    }
}

##### Var.VEr #####
# Function to calculate variance of eigenvalue variance for an arbitrary
# correlation matrix Var[V(R)] based on a Wishart matrix.
# This is just (p - 1)^2 times Var.VRr, which is called internally.
Var.VEr <- function(R, L, n = 100, ...) {
    if(missing(R)) {
        R <- GenCov(evalues = L, evectors = "Givens")
        if(L[2] != L[length(L)]) warning("R was generated from the eigenvalues provided \n  Expectation may vary even if eigenvalues are fixed")
    } else if(any(diag(R) != 1)) {
        R <- R / R[1, 1]
        if(any(diag(R) != 1)) stop("Provide a valid correlation matrix, or its eigenvalues")
        warning("R was scaled to have diagonal elements of unity")
    }
    p <- ncol(R)
    var.VRr <- Var.VRr(R = R, n = n, ...)
    var.VRr * (p - 1) ^ 2
}

##### Var.VRr #####
# Function to calculate variance of relative eigenvalue variance for an
# arbitrary correlation matrix Var[Vrel(R)] based on a Wishart matrix.
# Usage is similar to Exv.VRr.
## Details
# Returns the exact variance when p = 2 or under the null condition.
# Otherwise, asymptotic variance is returned with one of the functions
# defined below, as specified by the argument fun:
# pfd, pfv, pfc, pf are equivalent variants of Pan & Frank's (2004) approach
# (usually, pfd or pfc will be the choice);
# klv, kl, krv, kr are equivalent variants from Konishi's asymptotic theory
# (usually, klv will be the choice).
Var.VRr <- function(R, L, n = 100,
                    fun = c("pfd", "pfv", "pfc", "pf",
                            "klv", "kl", "krv", "kr"), ...) {
    fun <- match.arg(fun)
    if(missing(R)) {
        R <- GenCov(evalues = L, evectors = "Givens")
        if(L[2] != L[length(L)]) warning("R was generated from the eigenvalues provided \n  Expectation may vary even if eigenvalues are fixed")
    } else if(any(diag(R) != 1)) {
        R <- R / R[1, 1]
        if(any(diag(R) != 1)) stop("Provide a valid correlation matrix, or its eigenvalues")
        warning("R was scaled to have diagonal elements of unity")
    }
    p <- ncol(R)
    if(isTRUE(all.equal(R, diag(p)))) {
        4 * (n - 1) / (p * (p - 1) * n^2 * (n + 2))
    } else if(p == 2) {
        R2 <- R[lower.tri(R)] ^ 2
        var_r2 <- matrix(sapply(n, Var.r2, x = R2, do.square = FALSE),
                         ncol = length(n))
        4 * colSums(var_r2) / (p * (p - 1)) ^ 2
    } else {
        Fun <- switch(fun,
                      klv = AVar.VRr_klv, kl = AVar.VRr_kl, kr = AVar.VRr_kr,
                      krv = AVar.VRr_krv, pf = AVar.VRr_pf, pfv = AVar.VRr_pfv,
                      pfd = AVar.VRr_pfd, pfc = AVar.VRr_pfc)
        Fun(R = R, n = n, ...)
    }
}

##### AVar.VRr_pf #####
# Asymptotic and approximate variance of Vrel(R) based on
# Pan & Frank's (2004) approach.  Prototype version.
## Arguments
# R:         a population correlation matrix; assumed to be validly constructed
# L:         a vector of population eigenvalues
# n:         degree(s) of freedom; numeric of length 1 or more
# exv1.mode: Whether exact or asymptotic expression is used for E(r)
# var2.mode: Whether exact or asymptotic expression is used for Var(r^2)
# var1.mode: (At present, only "asymptotic" is accepted)
# order.exv1/order.var2: Used to specify the order of asymptotic expressions for
#                        E(r)/Var(r^2) when exv1.mode/var2.mode is "asymptotic";
#                        see descriptions of AExv.r1/AVar.r2 below
# mode:      Specifies the mode of iterations (see Details)
# (Below are used only when mode == "mclapply", "doParallel", or "parLapply")
# mc.cores:  Number of cores to be used.  When "auto" (default), set to
#            min(c(ceiling(p / 2), max.cores)), which seems to work well
# max.cores: Maximum number of cores to be used
# do.mcaffinity: Whether to run mcaffinity(), which seems required in
#                some Linux environments to assign threads to multiple cores
# affinity_mc:   Argument of mcaffinity() to specify assignment of threads
# cl:        A cluster object (made by makeCluster()); when already created,
#            one can be specified with this argument.  Otherwise, one is created
#            within this function, and is turned off on exit
## Details
# The algorithm involves evaluation of ~p^4 / 4 pairs of covariances between
# squared correlation coefficients.
# Options for mode:
# nested.for uses nested for loops, which is straifhgforward and RAM efficient.
# for.ind and lapply runs the iteration along an index vector to shorten
# computational time, although this does not seem to yield any much improvement.
# mclapply and parLapply parallelize this iteration by forking and socketing,
# respectively.  Note that the former doesn't work in the Windows environment.
# These depend on the package parallel.  See vignette("parallel") for details.
# The length of getInd(p) is about p^4 / 8 and that of B is about p^4 / 4
# therefore it takes RAM of ~2*p^4 bytes;
# this could be prohibitively large for large p.
## Dependency
# parallel (when mode == "mclapply" or "parLapply" is used)
AVar.VRr_pf <- function(R, L, n = 100, exv1.mode = c("exact", "asymptotic"),
                        var1.mode = "asymptotic",
                        var2.mode = c("exact", "asymptotic"),
                        order.exv1 = 2, order.var2 = 2,
                        mode = c("for.ind", "nested.for", "lapply",
                                 "mclapply", "parLapply"),
                        mc.cores = "auto", max.cores = detectCores(),
                        do.mcaffinity = TRUE,
                        affinity_mc = seq_len(max.cores), cl = NULL, ...) {
    exv1.mode <- match.arg(exv1.mode)
    var1.mode <- match.arg(var1.mode)
    var2.mode <- match.arg(var2.mode)
    mode <- match.arg(mode)
    getInds <- function(p){
        a <- seq_len(p)
        d <- digit(p)
        A <- outer(as.integer(10 ^ d) * a, a, "+")
        b <- t(A)[lower.tri(A)]
        B <- outer((100 ^ d) * b, b, "+")
        t(B)[lower.tri(B)]
    }
    parseInds <- function(x, d) {
        i <- (x %/% 1000 ^ d) %% 10 ^ d
        j <- (x %/% 100 ^ d) %% 10 ^ d
        k <- (x %/% 10 ^ d) %% 10 ^ d
        l <- x %% 10 ^ d
        c(i, j, k, l)
    }
    Cov_r2s <- function(I = NULL, pI) {
        if(!missing(I)) pI <- parseInds(I, d)
        i <- pI[1]
        j <- pI[2]
        k <- pI[3]
        l <- pI[4]
        Eij <- exv_r1[(j - 1) * p + i - (j - 1) * (2 * p - j + 2) / 2, ]
        Ekl <- exv_r1[(l - 1) * p + k - (l - 1) * (2 * p - l + 2) / 2, ]
        Cijkl <- v1fun(n = n, R = R, i = i, j = j, k = k, l = l)
        (4 * Eij * Ekl + 2 * Cijkl) * Cijkl
    }
    e1fun <- switch(exv1.mode,
                    exact = function(n, x) Exv.r1(n, x, ...),
                    asymptotic = function(n, x) AExv.r1(n, x,
                                                        order. = order.exv1))
    v1fun <- switch(var1.mode, ACov.r1)
    v2fun <- switch(var2.mode,
                    exact = function(n, x) Var.r2(n, x, do.square = TRUE, ...),
                    asymptotic = function(n, x) AVar.r2(n, x,
                                                        order. = order.var2))
    if(missing(R)) {
        R <- GenCov(evalues = L, evectors = "Givens")
        warning("R was generated from the eigenvalues provided \n  Expectation may vary even if eigenvalues are fixed")
    } else if(any(diag(R) != 1)) {
        R <- R / R[1, 1]
        if(any(diag(R) != 1)) stop("Provide a valid correlation matrix, or its eigenvalues")
        warning("R was scaled to have diagonal elements of unity")
    }
    p <- ncol(R)
    l_n <- length(n)
    R_u <- R[upper.tri(R)]
    var_r2 <- matrix(sapply(n, v2fun, x = R_u), ncol = l_n)
    exv_r1 <- matrix(sapply(n, e1fun, x = R_u), ncol = l_n)
    d <- digit(p)
    if(mode == "nested.for") {
        cov_r2 <- numeric(l_n)
        for(i in 1:(p - 2)) {
            for(j in (i + 1):p) {
                for(k in i:(p - 1)) {
                    for(l in (k + 1):p) {
                        if(i >= k && j >= l) next
                        Eij <- exv_r1[(j - 1) * p +
                                      i - (j - 1) * (2 * p - j + 2) / 2, ]
                        Ekl <- exv_r1[(l - 1) * p +
                                      k - (l - 1) * (2 * p - l + 2) / 2, ]
                        Cijkl <- sapply(n, v1fun, R = R,
                                        i = i, j = j, k = k, l = l)
                        cov_r2 <- cov_r2 + (4 * Eij * Ekl + 2 * Cijkl) * Cijkl
                    }
                }
            }
        }
    } else {
        Inds <- getInds(p)
        if(mode == "for.ind") {
            cov_r2 <- numeric(l_n)
            for(I in seq_along(Inds)) {
                cov_r2 <- cov_r2 + Cov_r2s(Inds[I])
            }
        } else {
            if(mode == "lapply") {
                cov_r2 <- lapply(Inds, Cov_r2s)
            } else {
                require(parallel)
                if(mc.cores == "auto") {
                    mc.cores <- pmin(ceiling(p / 2), max.cores)
                }
                if(mode == "mclapply") {
                    if(do.mcaffinity) try(invisible(mcaffinity(affinity_mc)),
                                          silent = TRUE)
                    cov_r2 <- mclapply(Inds, Cov_r2s, mc.cores = mc.cores)
                } else { # if(mode == "parLapply")
                    if(is.null(cl)) {
                        cl <- makeCluster(mc.cores)
                        on.exit(stopCluster(cl))
                        if(do.mcaffinity) try(clusterApply(cl,
                                              as.list(affinity_mc), mcaffinity),
                                              silent = TRUE)
                    }
                    cov_r2 <- parLapply(cl = cl, Inds, Cov_r2s)
                }
            }
            cov_r2 <- matrix(unlist(cov_r2), ncol = length(Inds))
            cov_r2 <- rowSums(cov_r2)
        }
    }
    ans <- colSums(var_r2) + 2 * cov_r2
    4 * ans / (p * (p - 1))^2
}

##### AVar.VRr_pfv #####
# Asymptotic and approximate variance of Vrel(R) based on
# Pan & Frank's (2004) approach.  Vectorized version.
# Usage is similar to AVar.VRr_pf.
# This is much faster than that, but requires a large RAM space as p grows.
AVar.VRr_pfv <- function(R, L, n = 100, exv1.mode = c("exact", "asymptotic"),
                       var1.mode = "asymptotic",
                       var2.mode = c("exact", "asymptotic"),
                       order.exv1 = 2, order.var2 = 2, ...) {
    exv1.mode <- match.arg(exv1.mode)
    var1.mode <- match.arg(var1.mode)
    var2.mode <- match.arg(var2.mode)
    rep_d <- function(x, from = 1, to = length(x)) {
        x <- x[seq.int(from, length(x))]
        sx <- seq_along(x)[seq_len(to - from + 1)]
        unlist(lapply(sx, function(i) x[-seq_len(i)]))
    }
    e1fun <- switch(exv1.mode,
                    exact = function(n, x) Exv.r1(n, x, ...),
                    asymptotic = function(n, x) AExv.r1(n, x,
                                                        order. = order.exv1))
    c1fun <- switch(var1.mode, function(n, Rij, Rkl, Rik, Rjl, Ril, Rjk) {
        A <- (Rij * Rkl * (Rik^2 + Ril^2 + Rjk^2 + Rjl^2) / 2 + Rik * Rjl +
             Ril * Rjk - (Rij * Rik * Ril + Rij * Rjk * Rjl + Rik * Rjk * Rkl +
                        Ril * Rjl * Rkl))
        outer(A, n, "/")
    })
    v2fun <- switch(var2.mode,
                    exact = function(n, x) Var.r2(n, x, do.square = TRUE, ...),
                    asymptotic = function(n, x) AVar.r2(n, x,
                                                        order. = order.var2))
    if(missing(R)) {
        R <- GenCov(evalues = L, evectors = "Givens")
        warning("R was generated from the eigenvalues provided \n  Expectation may vary even if eigenvalues are fixed")
    } else if(any(diag(R) != 1)) {
        R <- R / R[1, 1]
        if(any(diag(R) != 1)) stop("Provide a valid correlation matrix, or its eigenvalues")
        warning("R was scaled to have diagonal elements of unity")
    }
    p <- ncol(R)
    l_n <- length(n)
    R_u <- R[upper.tri(R)]
    var_r2 <- matrix(sapply(n, v2fun, x = R_u), ncol = l_n)
    d <- digit(p)
    a <- seq_len(p)
    A <- outer(as.integer(10 ^ d) * a, a, "+")
    b <- t(A)[lower.tri(A)]
    IJ <- rep.int(b, seq.int(length(b) - 1, 0))
    KL <- rep_d(b)
    Is <- as.integer((IJ %/% 10 ^ d) %% 10 ^ d)
    Js <- as.integer(IJ %% 10 ^ d)
    Ks <- as.integer((KL %/% 10 ^ d) %% 10 ^ d)
    Ls <- as.integer(KL %% 10 ^ d)
    rm(IJ, KL)
    exv_r1 <- matrix(sapply(n, e1fun, x = R_u), ncol = l_n)
    Eij <- exv_r1[(Js - 1) * p + Is - (Js - 1) * (2 * p - Js + 2) / 2, ]
    Ekl <- exv_r1[(Ls - 1) * p + Ks - (Ls - 1) * (2 * p - Ls + 2) / 2, ]
    Eij_Ekl <- Eij * Ekl
    rm(exv_r1, Eij, Ekl)
    Rij <- R[(Js - 1) * p + Is]
    Rkl <- R[(Ls - 1) * p + Ks]
    Rik <- R[(Ks - 1) * p + Is]
    Rjl <- R[(Ls - 1) * p + Js]
    Ril <- R[(Ls - 1) * p + Is]
    Rjk <- R[(Ks - 1) * p + Js]
    rm(Is, Js, Ks, Ls)
    Cijkl <- c1fun(n, Rij, Rkl, Rik, Rjl, Ril, Rjk)
    rm(Rij, Rkl, Rik, Rjl, Ril, Rjk)
    # cov_r2 <- 8 * Eij_Ekl * Cijkl + 4 * Cijkl ^ 2
    # ans <- colSums(var_r2) + colSums(cov_r2)
    cov_r2 <- 4 * (2 * crossprod(Eij_Ekl, Cijkl) + crossprod(Cijkl))
    ans <- colSums(var_r2) + diag(cov_r2)
    4 * ans / (p * (p - 1))^2
}

##### AVar.VRr_pfd #####
# Asymptotic and approximate variance of Vrel(R) based on
# Pan & Frank's (2004) approach; vectorized version modified
# Usage is similar to AVar.VRr_pf.
## Details
# This is an improvement over AVar.VRr_pfv.
# This function divides the index vector b into a list bd, and the calculations
# are done on each element of this list to save RAM space.
# This process takes some time when p is large (~10 sec for p = 1024, say).
# Alternatively, this list can be provided as the argument bd
# (which should exactly match the one to be generated).
# The argument max.size controls the maximum size of resulting vectors;
# at least max.size * (2 * length(n) + 6) * 8 bytes of RAM is required
# for storing temporary results (and more during computation);
# e.g., ~2e7 seems good for 16 GB RAM, ~4e8 for 256 GB
# However, performance does not seem to improve past 1e6-1e7 presumably because
# memory allocation takes substantial time for large objects.
# The iteration can be parallelized with modd == "mclapply" or "parLapply",
# but be careful about RAM limitations.
# When verbose == "yes" or "inline", the progress of iteration is printed.
## Dependency
# Depends on the package parallel if mode == "mclapply" or "parLapply" is used
AVar.VRr_pfd <- function(R, L, n = 100, exv1.mode = c("exact", "asymptotic"),
                         var1.mode = "asymptotic",
                         var2.mode = c("exact", "asymptotic"),
                         order.exv1 = 2, order.var2 = 2,
                         mode = c("for", "lapply", "mclapply", "parLapply"),
                         mc.cores = "auto", max.cores = detectCores(),
                         do.mcaffinity = TRUE,
                         affinity_mc = seq_len(max.cores), cl = NULL,
                         max.size = 2e6, bd = NULL,
                         verbose = c("yes", "inline", "no"), ...) {
    exv1.mode <- match.arg(exv1.mode)
    var1.mode <- match.arg(var1.mode)
    var2.mode <- match.arg(var2.mode)
    mode <- match.arg(mode)
    verbose <- match.arg(verbose)
    rep_d <- function(x, from = 1, to = length(x)) {
        x <- x[seq.int(from, length(x))]
        sx <- seq_along(x)[seq_len(to - from + 1)]
        unlist(lapply(sx, function(i) x[-seq_len(i)]))
    }
    bd2I1 <- function(bd) {
        lbd <- length(bd)
        q <- sapply(bd, length)
        qr <- q[seq.int(lbd, 1)]
        p1 <- cumsum(qr)[seq.int(lbd, 1)]
        p2 <- p1 - q + 1
        rbind(p1, p2)
    }
    e1fun <- switch(exv1.mode,
                    exact = function(n, x) Exv.r1(n, x, ...),
                    asymptotic = function(n, x) AExv.r1(n, x,
                                                        order. = order.exv1))
    c1fun <- switch(var1.mode, function(n, Rij, Rkl, Rik, Rjl, Ril, Rjk) {
        A <- (Rij * Rkl * (Rik^2 + Ril^2 + Rjk^2 + Rjl^2) / 2 + Rik * Rjl +
             Ril * Rjk - (Rij * Rik * Ril + Rij * Rjk * Rjl + Rik * Rjk * Rkl +
                        Ril * Rjl * Rkl))
        outer(A, n, "/")
    })
    v2fun <- switch(var2.mode,
                    exact = function(n, x) Var.r2(n, x, do.square = TRUE, ...),
                    asymptotic = function(n, x) AVar.r2(n, x,
                                                        order. = order.var2))
    disp <- switch(verbose,
        yes = function(i) cat(paste0(" ", i, "/", lbd, "; ", Sys.time(), "\n")),
        inline = function(i) {
            if(i %% 10^floor(log(lbd, 10) - 1) == 0) cat(paste0(" ", i, "."))
            else invisible(NULL)},
        function(i) invisible(NULL))
    Cov_r2m <- function(i) {
        disp(i)
        IJ <- rep.int(bd[[i]], seq.int(I1[1, i], I1[2, i]) - 1)
        KL <- rep_d(b, I2[1, i], I2[2, i])
        Is <- as.integer((IJ %/% 10 ^ d) %% 10 ^ d)
        Js <- as.integer(IJ %% 10 ^ d)
        Ks <- as.integer((KL %/% 10 ^ d) %% 10 ^ d)
        Ls <- as.integer(KL %% 10 ^ d)
        # rm(IJ, KL)
        Eij <- exv_r1[(Js - 1) * p + Is - (Js - 1) * (2 * p - Js + 2) / 2, ]
        Ekl <- exv_r1[(Ls - 1) * p + Ks - (Ls - 1) * (2 * p - Ls + 2) / 2, ]
        Eij_Ekl <- Eij * Ekl
        # rm(Eij, Ekl)
        Rij <- R[(Js - 1) * p + Is]
        Rkl <- R[(Ls - 1) * p + Ks]
        Rik <- R[(Ks - 1) * p + Is]
        Rjl <- R[(Ls - 1) * p + Js]
        Ril <- R[(Ls - 1) * p + Is]
        Rjk <- R[(Ks - 1) * p + Js]
        # rm(Is, Js, Ks, Ls)
        Cijkl <- c1fun(n, Rij, Rkl, Rik, Rjl, Ril, Rjk)
        # return(colSums(8 * Eij_Ekl * Cijkl + 4 * Cijkl ^ 2))
        return(4 * diag(2 * crossprod(Eij_Ekl, Cijkl) + crossprod(Cijkl)))
        # rm(Eij_Ekl, Cijkl)
    }
    if(missing(R)) {
        R <- GenCov(evalues = L, evectors = "Givens")
        warning("R was generated from the eigenvalues provided \n  Expectation may vary even if eigenvalues are fixed")
    } else if(any(diag(R) != 1)) {
        R <- R / R[1, 1]
        if(any(diag(R) != 1)) stop("Provide a valid correlation matrix, or its eigenvalues")
        warning("R was scaled to have diagonal elements of unity")
    }
    p <- ncol(R)
    l_n <- length(n)
    R_u <- R[upper.tri(R)]
    var_r2 <- matrix(sapply(n, v2fun, x = R_u), ncol = l_n)
    exv_r1 <- matrix(sapply(n, e1fun, x = R_u), ncol = l_n)
    d <- digit(p)
    if(is.null(bd)) {
        a <- seq_len(p)
        A <- outer(as.integer(10 ^ d) * a, a, "+")
        b <- t(A)[lower.tri(A)]
        bd <- divInd(b, Max = max.size)
    }
    I1 <- bd2I1(bd)
    I2 <- length(b) - I1 + 1
    lbd <- length(bd)
    cov_r2 <- numeric(l_n)
    if(verbose == "inline") cat(paste0(" Out of ", lbd, " iterations:"))
    if(mode == "for") {
        for(i in seq_along(bd)) {
            cov_r2 <- cov_r2 + Cov_r2m(i)
        }
        ans <- colSums(var_r2) + cov_r2
    } else {
        if(mode == "lapply") {
            cov_r2 <- lapply(seq_along(bd), Cov_r2m)
        } else {
            require(parallel)
            if(mc.cores == "auto") {
                mc.cores <- pmin(lbd, max.cores)
            }
            if(mode == "mclapply") {
                if(do.mcaffinity) try(invisible(mcaffinity(affinity_mc)),
                                      silent = TRUE)
                cov_r2 <- mclapply(seq_along(bd), Cov_r2m, mc.cores = mc.cores)
            } else if(mode == "parLapply") {
                if(is.null(cl)) {
                    cl <- makeCluster(mc.cores, outfile = "")
                    on.exit(stopCluster(cl))
                    if(do.mcaffinity) try(clusterApply(cl, as.list(affinity_mc),
                                          mcaffinity), silent = TRUE)
                }
                cov_r2 <- parLapply(cl = cl, seq_along(bd), Cov_r2m)
            }
        }
        cov_r2 <- matrix(unlist(cov_r2), l_n)
        ans <- colSums(var_r2) + rowSums(cov_r2)
    }
    if(verbose == "inline") cat("\n")
    4 * ans / (p * (p - 1))^2
}

##### AVar.VRr_pfc #####
# Asymptotic and approximate variance of Vrel(R) based on
# Pan & Frank's (2004) approach; with Rcpp
# Usage is similar to AVar.VRr_pf.
## Details
# This function makes use of C++ API via the package Rcpp for evaluation of
# the sum of covariance across pairs of squared correlation coefficients.
# This version works much faster than vectorized R codes (AVar.VRr_pfv/pfd).
# Note that the output can be slightly different from those of pure R codes
# (by the order of ~1e-9).
# When the function to be used (specified by cppfun) is not found,
# an attempt is made to sourceCpp() the .cpp file in the present directory.
# It is recommended to do, e.g., sourceCpp("Cov.r2C.cpp")
# to compile the C++ code beforehand as this typically takes several seconds.
## Dependency
# Depends on the package Rcpp
AVar.VRr_pfc <- function(R, L, n = 100, cppfun = "Cov_r2C",
                         exv1.mode = c("exact", "asymptotic"),
                         # var1.mode = "asymptotic",
                         var2.mode = c("exact", "asymptotic"),
                         order.exv1 = 2, order.var2 = 2, ...) {
    # cppfun <- match.arg(cppfun)
    exv1.mode <- match.arg(exv1.mode)
    # var1.mode <- match.arg(var1.mode)
    var2.mode <- match.arg(var2.mode)
    if(length(find(cppfun)) == 0) {
        require(Rcpp)
        sourceCpp(paste0(cppfun, ".cpp"))
    }
    c1fun <- switch(cppfun, Cov_r2C = Cov_r2C, Cov_r2V = Cov_r2V)
    e1fun <- switch(exv1.mode,
                    exact = function(n, x) Exv.r1(n, x, ...),
                    asymptotic = function(n, x) AExv.r1(n, x,
                                                        order. = order.exv1))
    v2fun <- switch(var2.mode,
                    exact = function(n, x) Var.r2(n, x, do.square = TRUE, ...),
                    asymptotic = function(n, x) AVar.r2(n, x,
                                                        order. = order.var2))
    if(missing(R)) {
        R <- GenCov(evalues = L, evectors = "Givens")
        warning("R was generated from the eigenvalues provided \n  Expectation may vary even if eigenvalues are fixed")
    } else if(any(diag(R) != 1)) {
        R <- R / R[1, 1]
        if(any(diag(R) != 1)) stop("Provide a valid correlation matrix, or its eigenvalues")
        warning("R was scaled to have diagonal elements of unity")
    }
    p <- ncol(R)
    l_n <- length(n)
    R_u <- R[upper.tri(R)]
    var_r2 <- matrix(sapply(n, v2fun, x = R_u), ncol = l_n)
    exv_r1 <- matrix(sapply(n, e1fun, x = R_u), ncol = l_n)
    cov_r2 <- c(c1fun(n, R, exv_r1))
    ans <- colSums(var_r2) + cov_r2
    4 * ans / (p * (p - 1))^2
}

##### digit #####
# Utility function to obtain the number of digit of an integer.
# For internal use in AVar.VRr_pfx functions (and others)
digit <- function(x) {
    as.integer(floor(log(x, 10))) + 1L
}

##### divInd #####
# Utility function to divide a vector of index to a list
# Mainly to be used internally in AVar.VRr_pfd
## Arguments
# b:      Vector to be divided into a list
# Max:    Maximum acceptable length of vector that results from the divided list
# Length: Length of the list generated
## Details
# The argument Max defines the maximum of length(bd[[i]]) in AVar.VRr_pfd,
# which defines the amount of RAM required (see description of that function).
# The number of steps (s) is defined as the maximum integer that satisfies
# the necessary condition: (p + p - s) * (s + 1) / 2 < Max
# Alternatively, Length could be specified to define the length of the list.
divInd <- function(b, Max = 2e6, Length = 2) {
    p <- length(b)
    if(missing(Max) && !missing(Length)) Max <- ceiling(nc / (Length - 1) - 1)
    ans <- list()
    while(p > 0) {
        if((2 * p - 1) ^ 2 + 8 * (p - Max) < 0) {
            s <- p - 1
        } else {
            s <- pmax(
                floor((2 * p - 1 - sqrt((2 * p - 1) ^ 2 + 8 * (p - Max))) / 2),
                0)
        }
        ans <- c(ans, list(b[1:(s + 1)]))
        b <- b[-(1:(s + 1))]
        p <- length(b)
    }
    return(ans)
}

##### hgf #####
# Wrapper for hypergeometric function with two upper and one lower arguments.
# This function utilizes the function genhypergeo from the package hypergeo;
# when it appears to have failed, then the function hypergeo is used.
# (For the present purpose, however, hyperg_2F1 of the package gsl may suffice.)
# Tries to reduce computational time by dropping duplicated values from x
# while trying to retain its original names and dim.
## Dependency
# Depends on the package hypergeo.
hgf <- function(a1, a2, b1, x, tol = 0, maxiter = 2000) {
    require(hypergeo)
    if(any(length(a1) > 1, length(a2) > 1, length(b1) > 1)) {
        stop("Only scalars are acceptable for a1, a2, and b1")
    }
    ox <- x
    dx <- dim(ox)
    nx <- names(ox)
    dim(ox) <- NULL
    names(ox) <- ox
    x <- unique(ox)
    names(x) <- ox[!duplicated(ox)]
    F <- Re(genhypergeo(c(a1, a2), b1, x, tol = tol, maxiter = maxiter))
    if(any(is.na(F) & !is.na(x))) F <- ifelse(is.na(F), Re(hypergeo(a1, a2, b1,
                                           x, tol = tol, maxiter = maxiter)), F)
    F <- F[names(ox)]
    names(F) <- nx
    dim(F) <- dx
    return(F)
}

##### Exv.r1 #####
# Calculates expectation of sample correlation coefficient(s).
## Arguments
# n: degree of freedom (not sample size), numeric of length 1
# x: population correlation coefficient, numeric of length 1 or more
# tol: tolerance used to judge wheter x is sufficiently close to 0 or +/- 1
#      (where numerical instability can happen)
# tol.hg, maxiter.hg: passed to hgf as tol and maxiter there
Exv.r1 <- function(n, x, tol = .Machine$double.eps * 100,
                   tol.hg = 0, maxiter.hg = 2000, ...) {
    C <- 2 / n * (gamma((n + 1) / 2) / gamma(n / 2)) ^ 2
    ifelse(abs(x) > 1 - tol, x, # When x ~ 1, the below may not converge
        ifelse(abs(x) < tol,
               0, # When x ~ 0
               C * x * hgf(1 / 2, 1 / 2, (n + 2) / 2, x ^ 2,
                           tol = tol.hg, maxiter = maxiter.hg)))
}

##### Exv.r2 #####
# Calculates expectation of sample correlation coefficient(s) squared.
# Usage is similar to Exv.r1.
## Arguments
# do.square: whether the provided x is to be squared before passed to hgf;
#            TRUE by default; set this FALSE when x is already squared.
Exv.r2 <- function(n, x, do.square = TRUE, tol = .Machine$double.eps * 100,
                   tol.hg = 0, maxiter.hg = 2000, ...) {
    if(do.square) x <- x ^ 2
    ifelse(abs(x) > 1 - tol, x,
        ifelse(abs(x) < tol,
               1 / n, # Same as below when x == 0
               {F <- hgf(1, 1, (n + 2) / 2, x,
                         tol = tol.hg, maxiter = maxiter.hg);
                1 - (n - 1) * (1 - x) / n * F}))
}

##### Exv.r3 #####
# Calculates expectation of sample correlation coefficient(s) ^ 3.
# Might be instable when the provided correlation coefficient x is small.
# Usage is similar to Exv.r1.
## Notes
# This expression is from Soper et al. (1917, eq. xxviii);
# that of Ghosh (1966) seems inaccurate for this moment.
Exv.r3 <- function(n, x, tol = .Machine$double.eps * 100,
                   tol.hg = 0, maxiter.hg = 2000, ...) {
    mu1_n0 <- Exv.r1(n, x, tol = tol, tol.hg = tol.hg, maxiter = maxiter.hg)
    mu1_n2 <- Exv.r1(n + 2, x, tol = tol, tol.hg = tol.hg, maxiter = maxiter.hg)
    mu1_n0 - n * (n - 1) * (mu1_n2 - mu1_n0)
}

##### Exv.r4 #####
# Calculates expectation of sample correlation coefficient(s) ^ 4.
# From Ghosh (1966, eq. 1).
# Usage is similar to Exv.r2.
Exv.r4 <- function(n, x, do.square = TRUE, tol = .Machine$double.eps * 100,
                   tol.hg = 0, maxiter.hg = 2000, ...) {
    if(do.square) x <- x ^ 2
    ifelse(abs(x) > 1 - tol, x,
        ifelse(abs(x) < tol,
               3 / n / (n + 2), # Same as below when x == 0
               {F <- hgf(1, 1, (n + 2) / 2, x, tol = tol.hg, max = maxiter.hg);
                Fd <- hgf(1, 2, (n + 4) / 2, x, tol = tol.hg, max = maxiter.hg);
                1 + (n - 1) * (n - 3) * (1 - x) / 2 / n * F -
                (n + 1) * (n - 1) * (1 - x) / 2 / (n + 2) * Fd}))
}

##### Var.r1 #####
# Calculates variance of sample correlation coefficient(s).
# Usage is similar to Exv.r1.
Var.r1 <- function(n, x, tol = .Machine$double.eps * 100,
                   tol.hg = 0, maxiter.hg = 2000, ...) {
    mu2_r1 <- Exv.r2(n, x, do.square = TRUE, tol = tol,
                     tol.hg = tol.hg, maxiter = maxiter.hg)
    mu1_r1 <- Exv.r1(n, x, tol = tol, tol.hg = tol.hg, maxiter = maxiter.hg)
    mu2_r1 - mu1_r1 ^ 2
}

##### Var.r2 #####
# Calculates variance of sample correlation coefficient(s) squared.
# Usage is similar to Exv.r2.
Var.r2 <- function(n, x, do.square = TRUE, tol = .Machine$double.eps * 100,
                   tol.hg = 0, maxiter.hg = 2000, ...) {
    if(do.square) x <- x ^ 2
    ifelse(abs(x) > 1 - tol, tol, # When x ~ 1, the below may not converge
        ifelse(abs(x) < tol,
               2 * (n - 1) / (n ^ 2 * (n + 2)), # When x ~ 0
               {F <- hgf(1, 1, (n + 2) / 2, x, tol = tol.hg, max = maxiter.hg);
                Fd <- hgf(1, 2, (n + 4) / 2, x, tol = tol.hg, max = maxiter.hg);
                (n - 1) * (n + 1) * (1 - x) / 2 / n *
                (F - n / (n + 2) * Fd - 2 * (n - 1) * (1 - x)
                                        / n / (n + 1) * F ^ 2)}))
}

##### Cm3.r1 #####
# Calculates the central third moment of sample correlation coefficient(s).
# Usage is similar to Exv.r1.
Cm3.r1 <- function(n, x, tol = .Machine$double.eps * 100,
                   tol.hg = 0, maxiter.hg = 2000, ...) {
    mu3_r1 <- Exv.r3(n = n, x = x, tol = tol, tol.hg = tol.hg, max = maxiter.hg)
    mu2_r1 <- Exv.r2(n, x, do.square = TRUE, tol = tol,
                     tol.hg = tol.hg, maxiter = maxiter.hg)
    mu1_r1 <- Exv.r1(n, x, tol = tol, tol.hg = tol.hg, maxiter = maxiter.hg)
    mu3_r1 - 3 * mu2_r1 * mu1_r1 + 2 * mu1_r1 ^ 3
}

##### Cm4.r1 #####
# Calculates the central fourth moment of sample correlation coefficient(s).
# Usage is similar to Exv.r1.
Cm4.r1 <- function(n, x, tol = .Machine$double.eps * 100,
                   tol.hg = 0, maxiter.hg = 2000, ...) {
    mu_r4 <- Exv.r4(n = n, x = x, tol = tol, tol.hg = tol.hg, max = maxiter.hg)
    mu_r3 <- Exv.r3(n = n, x = x, tol = tol, tol.hg = tol.hg, max = maxiter.hg)
    mu_r2 <- Exv.r2(n = n, x = x, tol = tol, tol.hg = tol.hg, max = maxiter.hg)
    mu_r1 <- Exv.r1(n = n, x = x, tol = tol, tol.hg = tol.hg, max = maxiter.hg)
    mu_r4 - 4 * mu_r3 * mu_r1 + 6 * mu_r2 * mu_r1 ^ 2 - 3 * mu_r1 ^ 4
}

##### AExv.r1 #####
# Calculates asymptotic expectation of sample correlation coefficient(s).
## Arguments
# n: degree of freedom (not sample size), numeric of length 1
# x: population correlation coefficient, numeric of length 1 or more
# order.:       order of approximation (exponent in O(m^r))
# return_terms: when TRUE, returns individual terms of different orders
## Details
# This is based on Ghosh's (1966) asymptotic expansion.
# Note that Ghosh (1966) used the symbol n for sample size.
# Allowd range of order. is 1-7;
# empirically, 2 seems most accurate except when n < 6 (where 3 works better)
# The argument return_terms is mainly for debugging,
# but could be used to track the order in derived statistics.
AExv.r1 <- function(n, x, order. = 2, return_terms = FALSE, ...) {
    if(order. > 7 || order. < 1) {
        order.old <- order.
        order. <- pmin(pmax(order., 1), 7)
        warning(paste("order. =", order.old, "is not allowed. order. =",
                      order., "was used."))
    }
    m <- n + 5
    lx <- length(x)
    rs <- sapply(x, function(z) z ^ (2 * seq.int(0, 6)))
    r2 <- rs[2, ]
    C <- matrix(c(
                3,        1,        0,        0,        0,        0,        0,
              121,       70,       25,        0,        0,        0,        0,
             6479,     4923,     2925,     1225,        0,        0,        0,
            86341,    77260,    58270,    38220,    19845,        0,        0,
          2290597,  2281659,  1972050,  1575350,  1157625,   800415,        0,
         30242033, 32443250, 30545375, 27171900, 23147775, 18523890, 19324305),
         nrow = 6, ncol = 7, byrow = TRUE)
    ms <- (m ^ seq.int(-1, -6) / c(4/3, 8, 64, 128, 512, 1024)) * 3
    tms <- ms[seq_len(order. - 1)] * C[seq_len(order. - 1), ] %*% rs
    tms <- rbind(1, tms)
    tms <- t(- x * (1 - r2) / 2 / m * t(tms))
    tms <- rbind(x, tms)
    rownames(tms) <- seq.int(0, order.)
    ans <- colSums(tms)
    if(return_terms){
        return(list(exp.value = ans, terms = tms))
    } else {
        return(ans)
    }
}

##### AVar.r1 #####
# Calculates asymptotic variance of sample correlation coefficient(s).
## Details
# Asymptotic expansion in Ghosh (1966). Usage is similar to AExv.r1.
# Allowd range of order. is 1-7;
# empirically, order. of 2 seems to work well for small-moderate n (<100 or so);
# for larger n, the difference tends to be negligible.
AVar.r1 <- function(n, x, order. = 2, return_terms = FALSE, ...) {
    if(order. > 7 || order. < 1) {
        order.old <- order.
        order. <- pmin(pmax(order., 1), 7)
        warning(paste("order. =", order.old, "is not allowed. order. =",
                      order., "was used."))
    }
    m <- n + 5
    lx <- length(x)
    rs <- sapply(x, function(z) z ^ (2 * seq.int(0, 6)))
    r2 <- rs[2, ]
    C <- matrix(c(
               14,       11,        0,        0,        0,        0,        0,
               98,      130,       75,        0,        0,        0,        0,
             2744,     4645,     4422,     2565,        0,        0,        0,
            19208,    37165,    44499,    40299,    26685,        0,        0,
           268912,   561743,   762882,   838182,   784770,   657135,        0,
          1882384,  4105472,  6005139,  7313100,  7924830,  7692660,  9365895),
          nrow = 6, ncol = 7, byrow = TRUE)
    ms <- (m ^ seq.int(-1, -6) / c(2, 2, 8, 8, 16, 16))
    tms <- ms[seq_len(order. - 1)] * C[seq_len(order. - 1), ] %*% rs
    tms <- rbind(1, tms)
    tms <- t((1 - r2) ^ 2 / m * t(tms))
    rownames(tms) <- seq.int(1, order.)
    ans <- colSums(tms)
    if(return_terms){
        return(list(exp.value = ans, terms = tms))
    } else {
        return(ans)
    }
}

##### ACm3.r1 #####
# Calculates asymptotic central third moment of
# sample correlation coefficient(s).
## Details
# Asymptotic expansion in Ghosh (1966). Usage is similar to AExv.r1.
# Allowd range of order. is 2-5; 3 seems best for moderate n (15-),
# while 4 may work better for smaller n.
## Note
# The validity of this asymptotic expression may need critical evaluation,
# as Ghosh's (1966) equation 1 (from which the result is supposedly drawn)
# seems inaccurate (see Soper et al. [1917: eq. xxviii]).
# The following functions depending on this might be inaccurate:
# AExv.r3, AExv.r4, AVar.r2
ACm3.r1 <- function(n, x, order. = 3, return_terms = FALSE, ...) {
    if(order. > 5 || order. < 2) {
        order.old <- order.
        order. <- pmin(pmax(order., 2), 5)
        warning(paste("order. =", order.old, "is not allowed. order. =",
                      order., "was used."))
    }
    m <- n + 5
    lx <- length(x)
    rs <- sapply(x, function(z) z ^ (2 * seq.int(0, 3)))
    r2 <- rs[2, ]
    C <- matrix(c(
               69,    88,     0,     0,
              797,  1691,  1560,     0,
            12325, 33147, 48099, 44109),
          nrow = 3, ncol = 4, byrow = TRUE)
    ms <- (m ^ seq.int(-1, -3) / c(1, 4 / 3, 8 / 3))
    tms <- ms[seq_len(order. - 2)] * C[seq_len(order. - 2), ] %*% rs
    tms <- rbind(6, tms)
    tms <- t(- x * (1 - r2) ^ 3 / m ^ 2 * t(tms))
    rownames(tms) <- seq.int(2, order.)
    ans <- colSums(tms)
    if(return_terms){
        return(list(exp.value = ans, terms = tms))
    } else {
        return(ans)
    }
}

##### ACm4.r1 #####
# Calculates asymptotic central fourth moment of
# sample correlation coefficient(s).
## Details
# Asymptotic expansion in Ghosh (1966). Usage is similar to AExv.r1.
# Allowd range of order. is 2-5;
# 3 looks good for moderate n (20-) especially when x is small;
# 4 is better for small n and large x (> 0.5)
ACm4.r1 <- function(n, x, order. = 3, return_terms = FALSE, ...) {
    if(order. > 5 || order. < 2) {
        order.old <- order.
        order. <- pmin(pmax(order., 2), 5)
        warning(paste("order. =", order.old, "is not allowed. order. =",
                      order., "was used."))
    }
    m <- n + 5
    lx <- length(x)
    rs <- sapply(x, function(z) z ^ (2 * seq.int(0, 3)))
    r2 <- rs[2, ]
    C <- matrix(c(
              12,    35,     0,     0,
             436,  2028,  3025,     0,
            3552, 20009, 46462, 59751),
          nrow = 3, ncol = 4, byrow = TRUE)
    ms <- (m ^ seq.int(-1, -3) / c(1, 4, 4))
    tms <- ms[seq_len(order. - 2)] * C[seq_len(order. - 2), ] %*% rs
    tms <- rbind(1, tms)
    tms <- t(3 * (1 - r2) ^ 4 / m ^ 2 * t(tms))
    rownames(tms) <- seq.int(2, order.)
    ans <- colSums(tms)
    if(return_terms){
        return(list(exp.value = ans, terms = tms))
    } else {
        return(ans)
    }
}

##### AExv.r2 #####
# Calculates asymptotic expectation of
# sample correlation coefficient(s) squared.
## Details
# Usage is similar to AExv.r1
# Allowd range of order. is 1-7;
# empirically, 2 seems most accurate except when n < 6 (where 3 works better)
AExv.r2 <- function(n, x, order. = 2, return_terms = FALSE, ...) {
    if(order. > 7 || order. < 1) {
        order.old <- order.
        order. <- pmin(pmax(order., 1), 7)
        warning(paste("order. =", order.old, "is not allowed. order. =",
                      order., "was used."))
    }
    m <- n + 5
    lx <- length(x)
    rs <- sapply(x, function(z) z ^ (2 * seq.int(0, 7)))
    r2 <- rs[2, ]
    C <- matrix(c(
               1,    -2,     0,      0,      0,      0,      0,       0,
               7,    -8,    -8,      0,      0,      0,      0,       0,
              49,   -26,   -56,    -48,      0,      0,      0,       0,
             343,   -32,  -272,   -384,   -384,      0,      0,       0,
            2401,   526,  -944,  -2016,  -2688,  -3840,      0,       0,
           16807,  7432,  -728,  -7680, -13440, -15360, -46080,       0,
          117649, 70774, 27544, -12048, -48000, -88320,  46080, -645120),
          nrow = 7, ncol = 8, byrow = TRUE)
    ms <- m ^ seq.int(-0, -6)
    tms <- ms[seq_len(order.)] * C[seq_len(order.), ] %*% rs
    tms <- t((1 - r2) / m * t(tms))
    tms <- rbind(r2, tms)
    rownames(tms) <- seq.int(0, order.)
    ans <- colSums(tms)
    if(return_terms){
        return(list(exp.value = ans, terms = tms))
    } else {
        return(ans)
    }
}


##### AExv.r3 #####
# Calculates asymptotic expectation of sample correlation coefficient(s) ^3.
## Details
# Usage is similar to AExv.r1
# Allowd range of order. is 1-5; 2 looks good.
AExv.r3 <- function(n, x, order. = 2, return_terms = FALSE, ...) {
    if(order. > 5 || order. < 1) {
        order.old <- order.
        order. <- pmin(pmax(order., 1), 5)
        warning(paste("order. =", order.old, "is not allowed. order. =",
                      order., "was used."))
    }
    m <- n + 5
    lx <- length(x)
    rs <- sapply(x, function(z) z ^ (2 * seq.int(0, 5)))
    r2 <- rs[2, ]
    C <- matrix(c(
              2,     -3,      0,      0,      0,        0,
             12,      1,    -25,      0,      0,        0,
            306,    371,   -100,  -1225,      0,        0,
           3112,   7011,   6175,   1225, -33075,        0,
          54486, 220811, 309080, 302330, 674730, -2401245),
          nrow = 5, ncol = 6, byrow = TRUE)
    ms <- m ^ seq.int(-0, -4) / c(2, 8/3, 16, 128/3, 256)
    tms <- ms[seq_len(order.)] * C[seq_len(order.), ] %*% rs
    tms <- t(3 * x * (1 - r2) / m * t(tms))
    tms <- rbind(x ^ 3, tms)
    rownames(tms) <- seq.int(0, order.)
    ans <- colSums(tms)
    if(return_terms){
        return(list(exp.value = ans, terms = tms))
    } else {
        return(ans)
    }
}

##### AExv.r4 #####
# Calculates asymptotic expectation of sample correlation coefficient(s) ^ 4.
## Details
# Usage is similar to AExv.r1
# Allowd range of order. is 1-5; 3 or 4 looks good for small x,
# while 2 is better for relatively large x (>0.4).
AExv.r4 <- function(n, x, order. = 2, return_terms = FALSE, ...) {
    if(order. > 5 || order. < 1) {
        order.old <- order.
        order. <- pmin(pmax(order., 1), 5)
        warning(paste("order. =", order.old, "is not allowed. order. =",
                      order., "was used."))
    }
    m <- n + 5
    lx <- length(x)
    rs <- sapply(x, function(z) z ^ (2 * seq.int(0, 6)))
    r2 <- rs[2, ]
    C <- matrix(c(
            0,    3,   -4,    0,     0,     0,      0,
            1,    1,   16,  -24,     0,     0,      0,
            6,   -9,   16,   88,  -128,     0,      0,
          109, -131,  -96,  144,  2688, -3200,      0,
          444, -291, -596, -464, -3200, 24960, -23040),
          nrow = 5, ncol = 7, byrow = TRUE)
    ms <- m ^ seq.int(-0, -4) / c(1/2, 1/3, 1/6, 1/3, 1/6)
    tms <- ms[seq_len(order.)] * C[seq_len(order.), ] %*% rs
    tms <- t((1 - r2) / m * t(tms))
    tms <- rbind(x ^ 4, tms)
    rownames(tms) <- seq.int(0, order.)
    ans <- colSums(tms)
    if(return_terms){
        return(list(exp.value = ans, terms = tms))
    } else {
        return(ans)
    }
}

##### AVar.r2 #####
# Calculates asymptotic variance of sample correlation coefficient(s) squared.
## Details
# Usage is similar to AExv.r1
# Allowd range of order. is 1-5.
# For moderate n (<100), 3 or 4 looks good for very small x,
# while 2 is better for relatively large x (>0.2).
# Slight overestimation seems to be common.
AVar.r2 <- function(n, x, order. = 2, return_terms = FALSE, ...) {
    if(order. > 5 || order. < 1) {
        order.old <- order.
        order. <- pmin(pmax(order., 1), 5)
        warning(paste("order. =", order.old, "is not allowed. order. =",
                      order., "was used."))
    }
    m <- n + 5
    lx <- length(x)
    rs <- sapply(x, function(z) z ^ (2 * seq.int(0, 5)))
    r2 <- rs[2, ]
    C <- matrix(c(
            0,    1,     0,     0,     0,     0,
            1,   -2,    26,     0,     0,     0,
           11,  -36,     8,   320,     0,     0,
           45,  -98,  -230,   -64,  2144,     0,
          323, -325, -1736, -2592, -6752, 32064),
          nrow = 5, ncol = 6, byrow = TRUE)
    ms <- m ^ seq.int(-0, -4) / c(1/4, 1/2, 1/2, 1/4, 1/4)
    tms <- ms[seq_len(order.)] * C[seq_len(order.), ] %*% rs
    tms <- t((1 - r2) ^ 2 / m * t(tms))
    rownames(tms) <- seq.int(1, order.)
    ans <- colSums(tms)
    if(return_terms){
        return(list(exp.value = ans, terms = tms))
    } else {
        return(ans)
    }
}

##### ACov.r1 #####
# Calculates asymptotic covariance between sample
# correlation coefficients from sample size and correlation matrix.
## Arguments
# n:          degree of freedom; numeric of length 1 or more
# R:          correlation matrix; assumed to be validly constructed
# ind1, ind2: indices for the two coefficients (vectors of length 2)
# i, j, k, l: alternative indices to directly specify the coefficients
## Details
# This asymptotic expression is as per, e.g., Olkin & Siotani (1976).
# When ind1 == ind2, it calculates the variance for the coefficient.
# When provided, i, j, k, l supercede the corresponding parts of ind1 or ind2.
# Alternatively, elements or R (rij, rkl, etc.) can be directly specified.
ACov.r1 <- function(n, R, ind1 = c(1, 2), ind2 = c(1, 2),
                    i = ind1[1], j = ind1[2], k = ind2[1], l = ind2[2],
                    rij = R[i, j], rik = R[i, k], ril = R[i, l],
                    rjk = R[j, k], rjl = R[j, l], rkl = R[k, l], ...) {
    (rij * rkl * (rik^2 + ril^2 + rjk^2 + rjl^2) / 2 + rik * rjl + ril * rjk -
     (rij * rik * ril + rij * rjk * rjl + rik * rjk * rkl + ril * rjl * rkl))/n
}

##### AVar.VRr_kl #####
# Asymptotic variance of Vrel(R) derived from Konishi (1979: corollary 2.2):
# Vrel(R) as a function of eigenvalues. Prototype version.
# Usage is similar to Exv.VRr.
# This is equivalent to, but much slower than, AVar.VRr_klv below
AVar.VRr_kl <- function(R, L, n = 100, ...) {
    if(missing(R)) {
        R <- GenCov(evalues = L, evectors = "Givens")
        warning("R was generated from the eigenvalues provided \n  Expectation may vary even if eigenvalues are fixed")
    } else if(any(diag(R) != 1)) {
        R <- R / R[1, 1]
        if(any(diag(R) != 1)) stop("Provide a valid correlation matrix, or its eigenvalues")
        warning("R was scaled to have diagonal elements of unity")
    }
    p <- ncol(R)
    svd.R <- svd(R, nu = 0)
    d <- svd.R$d
    V2 <- svd.R$v ^ 2
    R2 <- R ^ 2
    ans <- numeric(1)
    for(i in 1:p) {
        for(j in 1:p) {
            ans <- ans + d[i]^2 * d[j]^2 * drop(as.numeric(i == j)
                   - (d[i] + d[j]) * crossprod(V2[, i], V2[, j])
                   + crossprod(V2[, i], crossprod(R2, V2[, j])))
        }
    }
    8 * ans / (p * (p - 1)) ^ 2 / n
}

##### AVar.VRr_klv #####
# Asymptotic variance of Vrel(R) derived from Konishi (1979: corollary 2.2)
# Vectorized version.
# Usage is similar to Exv.VRr.
AVar.VRr_klv <- function(R, L, n = 100, ...) {
    if(missing(R)) {
        R <- GenCov(evalues = L, evectors = "Givens")
        warning("R was generated from the eigenvalues provided \n  Expectation may vary even if eigenvalues are fixed")
    } else if(any(diag(R) != 1)) {
        R <- R / R[1, 1]
        if(any(diag(R) != 1)) stop("Provide a valid correlation matrix, or its eigenvalues")
        warning("R was scaled to have diagonal elements of unity")
    }
    p <- ncol(R)
    svd.R <- svd(R, nu = 0)
    d <- svd.R$d
    d2 <- d ^ 2
    V2 <- svd.R$v ^ 2
    R2 <- R ^ 2
    G <- diag(p) - outer(d, d, "+") * crossprod(V2) +
         crossprod(V2, crossprod(R2, V2))
    F <- (d2 %*% t(d2)) * G
    ans <- sum(F)
    8 * ans / (p * (p - 1)) ^ 2 / n
}

##### AVar.VRr_kr #####
# Asymptotic variance of Vrel(R) derived from Konishi (1979: theorem 6.2):
# Vrel(R) as a function of correlation coefficients
# Usage is similar to Exv.VRr.
# Empirically, this gives the same result to AVar.VRr_kl/AVar.VRr_klv,
# but is much slower.
AVar.VRr_kr <- function(R, L, n = 100, ...) {
    if(missing(R)) {
        R <- GenCov(evalues = L, evectors = "Givens")
        warning("R was generated from the eigenvalues provided \n  Expectation may vary even if eigenvalues are fixed")
    } else if(any(diag(R) != 1)) {
        R <- R / R[1, 1]
        if(any(diag(R) != 1)) stop("Provide a valid correlation matrix, or its eigenvalues")
        warning("R was scaled to have diagonal elements of unity")
    }
    p <- ncol(R)
    ans <- numeric(1)
    for(i in 1:p) {
        for(j in (1:p)[-i]) {
            for(k in i:p) {
                for(l in (1:p)[-k]) {
                    ans <- ans + (R[j, k] - R[i, j] * R[i, k]) *
                                 (R[i, l] - R[i, k] * R[k, l]) *
                                 R[i, j] * R[k, l]
                }
            }
        }
    }
    16 * ans / (p * (p - 1)) ^ 2 / n
}

##### AVar.VRr_krv #####
# Asymptotic variance of Vrel(R) derived from Konishi (1979: theorem 6.2)
# Vectorized version; slightly faster for moderate p, but not particularly fast
# for large p as the number of elements to be summed is large.
AVar.VRr_krv <- function(R, L, n = 100, ...) {
    digit <- function(x) {
        i <- 1L
        while(x %/% 10 >= 1) {
            x <- x %/% 10
            i <- i + 1
        }
        return(i)
    }
    if(missing(R)) {
        R <- GenCov(evalues = L, evectors = "Givens")
        warning("R was generated from the eigenvalues provided \n  Expectation may vary even if eigenvalues are fixed")
    } else if(any(diag(R) != 1)) {
        R <- R / R[1, 1]
        if(any(diag(R) != 1)) stop("Provide a valid correlation matrix, or its eigenvalues")
        warning("R was scaled to have diagonal elements of unity")
    }
    p <- ncol(R)
    d <- digit(p)
    a <- seq_len(p)
    A <- outer(as.integer(10 ^ d) * a, a, "+")
    b <- A[lower.tri(A) | upper.tri(A)]
    B <- matrix(rep.int(b, length(b)), length(b))
    IJ <- t(B)[lower.tri(B)]
    KL <- B[lower.tri(B)]
    Is <- as.integer((IJ %/% 10 ^ d) %% 10 ^ d)
    Js <- as.integer(IJ %% 10 ^ d)
    Ks <- as.integer((KL %/% 10 ^ d) %% 10 ^ d)
    Ls <- as.integer(KL %% 10 ^ d)
    Rij <- R[(Js - 1) * p + Is]
    Rkl <- R[(Ls - 1) * p + Ks]
    Rik <- R[(Ks - 1) * p + Is]
    Rjl <- R[(Ls - 1) * p + Js]
    Ril <- R[(Ls - 1) * p + Is]
    Rjk <- R[(Ks - 1) * p + Js]
    ans <- (Rjk - Rij * Rik) * (Ril - Rik * Rkl) * Rij * Rkl
    ans <- sum(ans)
    16 * ans / (p * (p - 1)) ^ 2 / n
}

##### Var.VEav #####
# Function to calculate variance of unbiased eigenvalue variance for an
# arbitrary covariance matrix Var[V~(S)] based on a Wishart matrix.
# Usage is similar to Var.VEv.
Var.VEav <- function(V, L, n = 100, drop_0 = FALSE,
                     tol = .Machine$double.eps * 100, ...) {
    divisor <- match.arg(divisor)
    if(missing(L)) {
        L <- svd(V, nu = 0, nv = 0)$d
    }
    if(drop_0) {
        L <- L[L > tol]
    } else {
        L[L < tol] <- 0
    }
    p <- length(L)
    t1 <- sum(L)
    t2 <- sum(L ^ 2)
    t3 <- sum(L ^ 3)
    t4 <- sum(L ^ 4)
    4 * ((2 * p^2 * n^2 + 3 * p^2 * n - 6 * p^2 - 4 * p * n - 4) * t4 +
         4 * p * (n - 1) * (n + 2) * t3 * t1 +
         (p^2 * n + 4 * p + 2 * n + 2) * t2^2 +
         2 * (n - 1) * (n + 2) * t2 * t1^2) / (p ^ 4 * n * (n - 1) * (n + 2))
}

##### Var.VRav #####
# Function to calculate variance of adjusted relative eigenvalue variance for
# an arbitrary covariance matrix Var[V-rel(S)] based on a Wishart matrix.
# Usage is similar to Var.VRv, which is internally called.
Var.VRav <- function(V, L, n = 100, drop_0 = FALSE,
                 tol = .Machine$double.eps * 100, ...) {
    divisor <- match.arg(divisor)
    if(missing(L)) {
        L <- svd(V, nu = 0, nv = 0)$d
    }
    if(drop_0) {
        L <- L[L > tol]
    } else {
        L[L < tol] <- 0
    }
    p <- length(L)
    V <- Var.VRv(L = L, n = n, drop_0 = drop_0, tol = tol, ...)
    En <- (p + 2) / (p * n + 2)
    V / (1 - En)^2
}

##### Var.VRar #####
# Function to calculate variance of adjusted relative eigenvalue variance
# for an arbitrary correlation matrix Var[V-rel(R)] based on a Wishart matrix.
# Usage is similar to Var.VRr, which is internally called.
Var.VRar <- function(R, L, n = 100, tol = .Machine$double.eps * 100,
                     tol.hg = 0, maxiter.hg = 2000, ...) {
    if(missing(R)) {
        R <- GenCov(evalues = L, evectors = "Givens")
        if(L[2] != L[length(L)]) warning("R was generated from the eigenvalues provided \n  Expectation may vary even if eigenvalues are fixed")
    } else if(any(diag(R) != 1)) {
        R <- R / R[1, 1]
        if(any(diag(R) != 1)) stop("Provide a valid correlation matrix, or its eigenvalues")
        warning("R was scaled to have diagonal elements of unity")
    }
    p <- ncol(R)
    V <- Var.VRr(R = R, n = n, tol = tol,
                 tol.hg = tol.hg, maxiter.hg = maxiter.hg, ...)
    En <- 1 / n
    V / (1 - En)^2
}

##### GenCov #####
# Function to generate a covariance/correlation matrix of specified order p,
# shape, and (when applicable) relative variance of eigenvalues VR.
## Arguments
# p:        number of variables
# VR:       The relative eigenvalue variance of the generated cov matrix;
#           used only when shape = "q-large" or "elongated"
# scale.:   (Optional) Used to scale the trace of the generated cov matrix.
# shape:    Option to specify the shape of cov matrix;
#           ignored when evalues is provided.  Options are:
#           - q-large, q leading eigenvalues equally large,
#                      the rest equally small;
#           - elongate, special case of q-large with q forced to be 1;
#           - linearly_decreasing, eigenvalues linearly decreasing in magnitude;
#           - quadratically_decreasing, eigenvalues quadratically decreasing.
#           User-specified VR is ignored with the last two options.
# evalues:  Numeric vector of eigenvalues.
# evectors: Matrix of eigenvectors (when necessary transformed to orthonormal;
#           random vectors are added when the dimension is smaller).
#           Alternatively, option to specify how eigenvectors are generated.
#           Options assumed are:
#           - plain,  eigenvectors are taken as the identity matrix of order p.
#           - random, random eigenvectors are generated by QR decomposition
#                     of a random square matrix.
#           - MAP,    eigenvectors are constructed so that the resultant matrix
#                     is a correlation matrix with the specified eigenvalues
#                     (scaled so that the trace equals p);
#                     based on the iterative algorithm of Waller (2020).
#           - Givens, similar to MAP, but with the algorithm of
#                     Davies & Higham (2000) (based on codes of Waller [2020]);
#                     this is much faster and is guaranteed to converge.
#           A numeric matrix could be used to specify full or partial
#           eigenvectors for a cov matrix; random vectors are appended
#           when needed.  They are converted to be orthogonal when necessary.
# random_rotation: only relevant when evectors = "Givens";
#                  logical to specify whether the initial diagonal matrix of
#                  eigenvalues is randomly rotated or not
# pic.ind:  Only relevant when evectors = "Givens";
#           - random, the axes to be rotated are chosen randomly
#           - head/tail, the axes to be rotated are fixed to be the
#                        first/last one.
# tol:      Only relevant when evectors = "MAP"; numeric to specify the
#           tolerance against which convergence of eigenvalues is evaluated.
# maxits:   Only relevant when evectors = "MAP"; maximum number of iterations.
# Seed:     Random seed to be passed before random eigenvectors are generated.
# check:    Logical, whether to check and ensure all eigenvalues of the
#           resultant matrix are nonnegative.
GenCov <- function(p = length(evalues), VR = 0.5, scale. = NULL,
                   evalues = "auto", evectors = "plain",
                   shape = c("q-large", "elongate", "linearly_decreasing",
                             "quadratically_decreasing"), q = 1,
                   random_rotation = TRUE,
                   pick.ind = c("random", "head", "tail"),
                   tol = 1e-12, maxits = 5000L, Seed = NULL,
                   check = any(evalues == 0)) {
    if(missing(p) && missing(evalues)) stop("Provide either p or evalues")
    shape <- match.arg(shape)
    scale_unit <- function(X) {
       # apply(X, 2, function(x) x / drop(sqrt(crossprod(x))))
        sweep(X, 2, sqrt(diag(crossprod(X))), "/")
    }
    cbind_orth <- function(X, Y) {
        Y <- crossprod(diag(p) - tcrossprod(X), Y)
        Y <- scale_unit(Y)
        cbind(X, Y)
    }
    S_fromUL <- function(evec = evec, evalues = evalues) {
        te <- t(evec)
        crossprod(te * evalues, te)
    }
    if(missing(pick.ind) && !random_rotation) {
        pick.ind <- "head"
    } else {
        pick.ind <- match.arg(pick.ind)
    }
    sample1 <- switch(pick.ind,
                      random = function(x) x[ceiling(runif(1) * length(x))],
                      head = function(x) x[1],
                      tail = function(x) x[length(x)])
    signp <- function(x) {
        ans <- sign(x)
        ans[ans == 0] <- 1
        return(ans)
    }
    if(missing(evalues)) {
        evalues <- numeric(p)
        if(shape == "elongate") {
            if(!missing(q)) warning("user-specified q is ignored for shape = \"elongate\"")
            q <- 1
            shape <- "q-large"
        }
        if(shape == "q-large") {
            r <- sqrt(VR)
            evalues[1:q] <- rep.int(1 + r * sqrt((p - 1) * (p - q) / q),
                                    q) / sqrt(p - 1)
            evalues[(q + 1):p] <- rep.int(1 - r * sqrt(q * (p - 1) / (p - q)),
                                          p - q) / sqrt(p - 1)
            if(any(evalues <= 0)) {
                stop("Negative eigenvalues. Ensure VR < (p - q) / ((p - 1) * q)")
            }
        } else {
            if(!missing(VR)) {
                warning("In the selected method, VR is fixed for a given p.\n User-defined VR was ignored.")
            }
            if(shape == "linearly_decreasing") {
                evalues <- 2 * (p + 1 - seq_len(p)) / (p + 1) / sqrt(p - 1)
            } else if(shape == "quadratically_decreasing") {
                evalues <- (6 * (p + 1 - seq_len(p)) ^ 2) / ((p + 1) * (2 * p + 1)) /
                           sqrt(p - 1)
            }
        }
    }
    if(evectors[1] == "MAP" || evectors[1] == "Givens") {
        if(any(evalues < 0)) stop("Provide a vector of nonnegative eigenvalues")
        sum.eval <- sum(evalues)
        if(sum.eval != p) {
            evalues <- evalues * p / sum.eval
            if(missing(scale.)) warning("Eigenvalues were scaled to sum to p\n")
        }
    }
    if(evectors[1] == "Givens") {
        if(random_rotation) {
            if(!is.null(Seed)) set.seed(Seed)
            M <- matrix(rnorm(p ^ 2), nrow = p, ncol = p)
            evec <- qr.Q(qr(M))
        } else {
            evec <- diag(p)
        }
        A <- S_fromUL(evec, evalues)
        a <- diag(A)
        y <- which(a < 1)
        z <- which(a > 1)
        while(length(y) > 0 && length(z) > 0) {
            i <- sample1(y)
            j <- sample1(z)
            alpha <- sqrt(A[i, j] ^ 2 - (a[i] - 1) * (a[j] - 1))
            T <- (A[i, j] + signp(A[i, j]) * alpha) / (a[j] - 1)
            C <- 1 / sqrt(1 + T ^ 2)
            S <- T * C
            R <- matrix(c(C, -S, S, C), 2, 2)
            A[, c(i, j)] <- A[, c(i, j)] %*% R
            A[c(i, j), ] <- t(R) %*% A[c(i, j), ]
            A[i, i] <- 1
            a[i] <- 1
            a[j] <- A[j, j]
            y <- which(a < 1)
            z <- which(a > 1)
        }
        diag(A) <- 1
        A <- (A + t(A)) / 2
        if(check) {
            svd.A <- svd(A, nu = 0)
            if(any(svd.A$d < 0)) {
                A <- with(svd.A, S_fromUL(v, pmax(d, 0)))
            }
        }
        if(!is.null(scale.)) A <- A / p * scale.
        return(A)
    } else {
        if(evectors[1] == "plain") {
            evec <- diag(p)
        } else if(is.character(evectors)) {
            if(!is.null(Seed)) set.seed(Seed)
            M <- matrix(rnorm(p ^ 2), nrow = p, ncol = p)
            evec <- qr.Q(qr(M))
            if(evectors == "MAP") {
                Sstar <- S_fromUL(evec, evalues)
                i <- 0L
                while(i <= maxits) {
                    Su <- Sstar
                    diag(Su) <- 1
                    svd.Su <- svd(Su, nu = 0)
                    v <- svd.Su$v
                    d <- svd.Su$d
                    if(max(abs(evalues - d)) < tol) break
                    Sstar <- S_fromUL(v, evalues)
                    Sstar <- (Sstar + t(Sstar)) / 2
                    i <- i + 1L
                }
                if(i > maxits) warning(paste("\nEigenvectors for cor matrix failed to converge after",
                                             maxits, "iterations"))
                evec <- v
            }
        } else {
            evectors <- as.matrix(evectors)
            if(nrow(evectors) != p) {
                stop("User-specified eigenvectors should be a matrix with p rows")
            }
            q <- ncol(evectors)
            if(any(crossprod(evectors)[lower.tri(matrix(nrow = q,
                                                        ncol = q))] != 0)) {
                warning("User-specified eigenvectors were converted to be orthonormal")
                evec <- evectors[, 1, drop = FALSE]
                evec <- scale_unit(evec)
                for(i in 2:q) {
                    y <- evectors[, i, drop = FALSE]
                    evec <- cbind_orth(evec, y)
                }
            } else if(any(diag(crossprod(evectors)) != 1)) {
                warning("User-specified eigenvectors were scaled to be of unit length")
                evec <- scale_unit(evectors)
            }
            if(ncol(evec) < p) {
                while(ncol(evec) < p) {
                    y <- matrix(rnorm(p), p, 1)
                    evec <- cbind_orth(evec, y)
                }
            }
        }
        S <- S_fromUL(evec, evalues)
        S <- (S + t(S)) / 2
        if(check) {
            svd.S <- svd(S, nu = 0)
            if(any(svd.S$d < 0)) {
                S <- with(svd.S, S_fromUL(v, pmax(d, 0)))
            }
        }
        if(!is.null(scale.)) S <- S / sum(diag(S)) * scale.
        return(S)
    }
}

#### chol.piv #####
# Cholesky factorization with pivoting, for singular covariance matrices.
# crossprod(chol.piv(A)) will be equal to A,
# unless multiple 0 eigenvalues exist.
# Be careful when multiple eigenvalues are 0.
## Arguments
# A: a numeric matrix, assumed to be symmetric
## Values
# Upper triangular matrix which is the Cholesky factor of A.
chol.piv <- function(A) {
    cA <- suppressWarnings(chol(A, pivot = TRUE))
    cA <- cA[, order(attr(cA, "pivot"))]
    return(cA)
}

##### matsqrt #####
# Matrix square root with spectral or singular value decomposition,
# to be used within chol.qr below.
## Arguments
# A:      a numeric matrix, assumed to be symmetric
# method: either "svd" (default) or "eigen" to specify the function to be used.
#         Results should be identical, but svd seems faster.
## Values
# symmetric matrix whose cross-product is A.
matsqrt <- function(A, method = c("svd", "eigen")) {
    method <- match.arg(method)
    if(method == "svd") {
       svdA <- svd(A, nu = 0)
       tv <- t(svdA$v)
       d <- svdA$d
    } else {
        eigenA <- eigen(A)
        tv <- t(eigenA$vectors)
        d <- eigenA$values
    }
    sqrA <- crossprod(tv * sqrt(pmax(d, 0)), tv)
    return(sqrA)
}

##### chol.qr #####
# Cholesky factorization with QR decomposition, for singular
# covariance matrices.
# crossprod(chol.qr(A)) will be equal to A, even with multiple 0 eigenvalues.
# Much slower than chol.piv.
## Arguments
# A:      a numeric matrix, assumed to be symmetric
# method: to be passed to matsqrt.
## Values
# Upper triangular matrix which is the Cholesky factor of A
chol.qr <- function(A, method = c("svd", "eigen")) {
    method <- match.arg(method)
    sqrA <- matsqrt(A, method = method)
    QRsqrA <- qr(sqrA)
    cA <- qr.R(QRsqrA)
    return(cA)
}

##### chol.qrsvd #####
# Cholesky factorization with QR decomposition, for singular
# covariance matrices.
# crossprod(chol.qr(A)) will be equal to A.
chol.qrsvd <- function(A) {
    svdA <- svd(A, nu = 0)
    tv <- t(svdA$v)
    sqrA <- crossprod(tv * sqrt(pmax(svdA$d, 0)), tv)
    QRsqrA <- qr(sqrA)
    cA <- qr.R(QRsqrA)
    return(cA)
}

##### mc, scale2 #####
# Utility functions for mean-centering
mc <- function(x) {
    sweep(x, 2, colMeans(x))
}

scale2 <- function(x, center = TRUE, scale = FALSE) {
    ans <- scale(x, center = center, scale = scale)
    ans[, attr(ans, "scaled:scale") == 0] <- 0
    return(ans)
}

##### rmvn #####
# Function to generate multivariate normal variables.
## Arguments
# N:           number of observations / sample size
# p:           number of variables; ignored when V or cV is provided.
# s2:          population variance used for the spherical condition;
#              ignored when V or cV is provided.
# V:           population covariance matrix, assumed validly constructed;
#              by default a spherical covariance is used;
#              ignored when cV is provided.
# cV:          Cholesky factor of V; when multiple simulations are done,
#              cV can be specified instead of V for computational efficiency.
# mean:        population mean; default is 0.
# chol.method: method for Cholesky factorization; "default" calls chol,
#              "pivot" calls chol.piv, "qr" callse chol.qr.
## Details
# cV can be specified instead of V for computational efficiency, especially
# when multiple simulations are done.
## Values
# N * p matrix with the specified population mean and covariance.
rmvn <- function(N, p = 2L, s2 = 1, V = s2 * diag(p), cV = cholfun(V),
                 mean = NULL, chol.method = c("default", "pivot", "qr")) {
    chol.method <- match.arg(chol.method)
    if(missing(cV)) {
        cholfun <- switch(chol.method, qr = chol.qr, pivot = chol.piv, chol)
        cV <- cholfun(V)
    }
    p <- nrow(cV)
    X <- matrix(rnorm(p * N), N, p)
    X <- X %*% cV
    if (missing(mean)) mean <- rep_len(0, p)
    X <- sweep(X, 2, mean, "+")
    return(X)
}

##### VE #####
# Function to calculate eigenvalue variance V and
# relative eigenvalue variance Vrel of a covariance/correlation matrix.
## Arguments
# X:       data matrix from which cov/cor matrix is obtained.
# V:       covariance/correlation matrix
# L:       vector of eigenvalues
# center:  logical to specify whether sample-mean-centering should be done
# scale.:  logical to specify whether SD-scaling should be done
#          (that is, when TRUE, the analysis is on the correlation matrix)
# divisor: either "UB" (default) or "ML", to decide the default value of m
# m:       divisor of cov matrix to be used ("n*" in the paper)
# nv:      specify how many eigenvectors are to be retained; default 0
# sub:     vector to specify the range of eigenvalue indices to be involved;
#          used to exclude the subspace not specified.
# drop_0:  logical, when TRUE, eigenvalues smaller than tol are dropped
# tol:     tolerance to be used with drop_0
## Details
# V, Vrel, mean eigenvalue, and a vector of eigenvalues (L) are returned,
# along with a matrix of eigenvectors when nv > 0.
# Provide either a data matrix (X), cov matrix (V),
# or vector of eigenvalues (L).
# When X is given, the default divisor is sample size - 1.
# Give the vector sub to specify the subspace in which VE is to be calculated.
# Set drop_0 = TRUE if nil eigenvalues are to be omitted from calculation.
## Value
# A list containing the following
# $VE:    eigenvalue variance (V)
# $VR:    relative eigenvalue variance (Vrel)
# $meanL: mean (average) of the eigenvalues
# $L:     vector of eigenvalues
# $U:     matrix of eigenvectors, when nv > 0
VE <- function(X, V, L, center = TRUE, scale. = FALSE,
               divisor = c("UB", "ML"), m = switch(divisor, UB = N - 1, ML = N),
               nv = 0, sub = seq_len(length(L)),
               drop_0 = FALSE, tol = .Machine$double.eps * 100) {
    divisor <- match.arg(divisor)
    if(!missing(X)) {
        X <- scale2(X, center = center, scale = scale.)
        N <- nrow(X)
        p <- ncol(X)
        svd.X <- svd(X, nu = 0, nv = nv)
        L <- svd.X$d
        L <- (L ^ 2) / m
        L <- c(L, rep_len(0, max(p - length(L), 0)))
    } else if(!missing(V)){
        p <- ncol(V)
        if(scale.) V <- cov2cor(V)
        svd.X <- svd(V, nu = 0, nv = nv)
        L <- svd.X$d
    }
    L <- L[sub]
    if(drop_0) L <- L[L > tol]
    p <- length(L)
    mL <- mean(L)
    VE <- sum((L - mL) ^ 2) / p
    VR <- VE / ((p - 1) * mL ^ 2)
    ans <- list(VE = VE, VR = VR, meanL = mL, L = L)
    if(nv > 0) ans <- c(ans, list(U = svd.X$v))
    return(ans)
}

##### Sim.VE #####
# Function to simulate sample cov and cor matrices,
# retaining eigenvalues, eigenvectors (if nv > 0), and eigenvalue variance
# (both unstandardized [V] and standardized [Vrel]).
## Arguments
# b:          number of iterations
# X:          data matrix; when V and cV are missing, the sample covariance
#             matrix from X is used as the population covariance matrix in
#             simulations (parametric bootstrapping).
# V, cV, N:   passed to rmvn
# divisor, m, nv, drop_0, tol: passed to VE
# center, scale.: passed to VE,
#                 as well as used to construct V when X is provided
## Details
# This function generates normal variables using the function rmvn,
# retaining eigenvalues of the covariance and correlation matrices
# as well as eigenvalue dispersion indices obtained by the function VE
# (and optionally eigenvectors when nv > 0).
# Suffices v and r denote cov and cor matrices.
# Provide either data (X), cov matrix (V), or its Cholesky factor (cV).
# It is recommended to specify the sample size N, unless X is provided.
## Values
# A list containing the following contents:
#  $VEv: eigenvalue variance of cov matrix V(S) (b vector)
#  $VRv: relative eigenvalue variance of cov matrix Vrel(S) (b vector)
#  $Lv:  eigenvalues of cov matrix (p * b matrix)
#  $VEr: eigenvalue variance of cor matrix V(R) (b vector)
#  $VRr: relative eigenvalue variance of cor matrix Vrel(R) (b vector)
#  $Lr:  eigenvalues of cor matrix (p * b matrix)
# (the rest are optional, when nv > 0)
#  $Uv:  eigenvectors of cov matrix (p * nv * b array)
#  $Ur:  eigenvectors of cor matrix (p * nv * b array)
#  $Uv.org:  eigenvectors of population cov matrix (p * nv matrix)
#  $Ur.org:  eigenvectors of population cor matrix (p * nv matrix)
# (Uv.org and Ur.org are provided as references for signs of eigenvectors)
Sim.VE <- function(b = 100L, X, V, cV = cholfun(V), N = nrow(X),
                   divisor = c("UB", "ML"),
                   m = switch(divisor, UB = N - 1, ML = N),
                   center = TRUE, scale. = FALSE, sub = seq_len(ncol(cV)),
                   nv = 0, chol.method = c("default", "pivot", "qr"),
                   drop_0 = FALSE, tol = .Machine$double.eps * 100) {
    chol.method <- match.arg(chol.method)
    divisor <- match.arg(divisor)
    if(missing(V) && missing(cV) && !missing(X)) {
        X <- scale2(X, center = center, scale = scale.)
        V <- crossprod(X) / m
    }
    if(missing(cV)) {
        cholfun <- switch(chol.method, qr = chol.qr, pivot = chol.piv, chol)
        cV <- cholfun(V)
    } else if(!missing(V) || !missing(X)) {
        warning("The arguments X and/or V were ignored as cV was given")
    }
    p <- ncol(cV)
    Uv.org <- svd(cV, nu = 0, nv = nv)$v
    Ur.org <- svd(cov2cor(crossprod(cV)), nu = 0, nv = nv)$v
    ansv.L <- matrix(numeric(b * p), p, b)
    ansr.L <- matrix(numeric(b * p), p, b)
    ansv.VE <- numeric(b)
    ansv.VR <- numeric(b)
    ansr.VE <- numeric(b)
    ansr.VR <- numeric(b)
    ansv.U <- array(numeric(b * p * nv), dim = c(p, nv, b))
    ansr.U <- array(numeric(b * p * nv), dim = c(p, nv, b))
    dimnames(ansv.L)[[1]] <- paste0("L", seq_len(p))
    dimnames(ansr.L)[[1]] <- paste0("L", seq_len(p))
    dimnames(ansv.U)[[1]] <- paste0("v", seq_len(p))
    dimnames(ansr.U)[[1]] <- paste0("v", seq_len(p))
    i <- 1L
    while(i <= b) {
        Xi <- rmvn(cV = cV, N = N, chol.method = chol.method)
        ansv <- tryCatch(VE(Xi, center = center, scale. = FALSE, nv = nv,
                            sub = sub, divisor = divisor, m = m,
                            drop_0 = drop_0, tol = tol),
                         error = function(e) list(-1, numeric(p)))
        ansr <- tryCatch(VE(Xi, center = center, scale. = TRUE, nv = nv,
                            sub = sub, divisor = divisor, m = m,
                            drop_0 = drop_0, tol = tol),
                         error = function(e) list(-1, numeric(p)))
        if(ansv[[1]] < 0 || ansr[[1]] < 0) next
        ansv.VE[i] <- ansv$VE
        ansv.VR[i] <- ansv$VR
        ansr.VE[i] <- ansr$VE
        ansr.VR[i] <- ansr$VR
        ansv.L[, i] <- ansv$L
        ansr.L[, i] <- ansr$L
        ansv.U[, , i] <- ansv$U
        ansr.U[, , i] <- ansr$U
        i <- i + 1L
    }
    ans <- list(VEv = ansv.VE, VRv = ansv.VR, Lv = ansv.L, VEr = ansr.VE,
                VRr = ansr.VR, Lr = ansr.L)
    if(nv > 0) {
        for(i in seq_len(nv)) {
            Indv <- crossprod(Uv.org[, i], ansv.U[, i, ]) < 0
            Indr <- crossprod(Ur.org[, i], ansr.U[, i, ]) < 0
            ansv.U[, i, Indv] <- -ansv.U[, i, Indv]
            ansr.U[, i, Indr] <- -ansr.U[, i, Indr]
        }
        ans <- c(ans, list(Uv = ansv.U, Ur = ansr.U,
                           Uv.org = Uv.org, Ur.org = Ur.org))
    }
    invisible(ans)
}
```
